# Targeted degradation of USP7 in solid cancer cells reveals disparate effects of deubiquitinase inhibition vs. acute protein depletion

**DOI:** 10.1101/2025.07.01.662351

**Authors:** Nikolas Klink, Sebastian Urban, Johanna A. Seier, Bikash Adhikari, Martin P. Schwalm, Juliane Müller, Madeleine Dorsch, Farnusch Kaschani, Johannes Koch, Siska Führer, Markus Kaiser, Nina Schulze, Stefan Knapp, Elmar Wolf, Annette Paschen, Barbara M. Grüner, Malte Gersch

## Abstract

Proteolysis-targeting chimeras (PROTACs) co-op the ubiquitin system for targeted protein degradation, creating opportunities to interrogate cellular functions of proteins through “chemical knockdown”. However, matched pairs of protein degraders and inhibitors, that possess high specificity and chemical complementarity, for individual components of the ubiquitin system have remained scarce. This includes reagents to modulate activity and abundance of deubiquitinases (DUBs), which critically regulate ubiquitin-mediated signaling. Here, using an integrated chemical biology approach, we explored the cellular function of the DUB USP7 as a case study comparing inhibition and degradation of this DUB in melanoma and pancreatic cancer cells. Through the synthesis of a degrader library, we identified potent USP7 PROTACs for each cancer type, established BRET-based ternary complex formation and quantified degradation efficiency. USP7 degraders and their cognate inhibitor were subsequently employed to characterize treatment-induced phenotypic alterations. Proteomic and cellular analyses revealed that highly specific degradation of USP7 modulated both shared and distinct protein sets across cancer cell types, without impacting cell growth. Notably, cellular responses to USP7 degradation differed markedly from those to USP7 inhibition. Moreover, our data uncovered broad proteomic and metabolic changes induced by prolonged USP7 inhibitor treatment. Collectively, our work provides a chemical toolbox of comprehensively characterized reagents to distinguish on-target phenotypes which will aid the understanding of the role of USP7 in malignant diseases. More broadly, our data emphasize the importance of increased specificity via PROTAC-mediated degradation and the potential of this modality to distinguish catalytic from non-catalytic as well as cell-line specific functions of DUBs.

## Introduction

Proteolysis-targeting chimeras (PROTACs) have emerged as powerful tools for both therapeutic applications and mechanistic biology. These hetero-bifunctional molecules enable rapid targeted protein degradation by hijacking the endogenous ubiquitin-proteasome system and have demonstrated remarkable potential in overcoming some of the limitations associated with conventional small molecule inhibitors.^1,2^ Moreover, comparative studies assessing effects of targeted protein degradation vs. protein inhibition have revealed distinct outcomes facilitated by both modalities (e.g. for the transcriptional regulator BRD4)^3–5^ and enabled the discovery of non-catalytic protein functions (e.g. for the kinase Aurora A).^6,7^ However, such comparative analyses require matched pairs of well-characterized inhibitors and degraders, a requirement that remains largely unmet across most protein families.

The around 100 human deubiquitinating enzymes (DUBs) act as counterplayers of over 600 E3 ligases and represent a particularly compelling yet underexplored class of proteins where matched pairs of inhibitors and degraders are urgently needed.^8,9^ These enzymes serve as critical regulators of cellular protein homeostasis and ubiquitin-mediated signaling. DUBs have emerged as promising therapeutic targets in cancer, neurodegeneration, and inflammatory disorders.^10^ Despite their therapeutic potential, the development of selective DUB modulators has been challenging which has also hampered investigations into their cellular roles.^11^

Ubiquitin-specific protease 7 (USP7) represents one of the most extensively studied human deubiquitinases.^12^ USP7 has gathered significant attention as a regulator of key cellular processes including the p53-MDM2 axis (stabilizing both proteins depending on the cellular context), DNA damage response (DDR, stabilizing key DDR proteins such as MDC1 or TRIP12), and transcription (through PRC1 complexes as well as RYBP).^13–16^ The multifaceted functions of this enzyme and its involvement in various cancer types have established USP7 as an attractive target for drug development, with several inhibitors being explored in preclinical studies.^10,17–20^

Notably, while the targeted degradation of DUB proteins has been explored,^21–26^ studies describing matched pairs of well characterized inhibitors and PROTACs do not exist for DUBs. To address this unmet need, we selected USP7 as a case study and investigated inhibitor-PROTAC pairs in the biological context of pancreatic ductal adenocarcinoma (PDAC) and melanoma. USP7 is particularly suited to undergo a systematic comparative analysis of degradation vs. inhibition phenotypes for the following reasons:

i. USP7 possesses numerous substrates and exhibits many context-dependent functions across different cell types.^12,27–29^ This complexity demands well characterized and potent chemical tools to dissect its multifaceted biology.
ii. In addition to its well-characterized catalytic activities, non-catalytic functions of DUBs are increasingly appreciated.^30^ These include scaffold functions in chromatin regulation and protein complex assembly that are independent of deubiquitinase activity. A matched pair of USP7 inhibitor and degrader will thus provide valuable tools for dissecting these distinct functions.
iii. USP7 is highly expressed in many cancers. However, its role in solid cancers remains poorly understood. This includes PDAC and melanoma wherein high USP7 levels are associated with poor prognosis.^31–37^
iv. While therapeutic interest has driven the development of many USP7 inhibitors, recent reports showcased that widely used compounds like P22077 lack sufficient specificity for USP7.^11,38^ Although more specific USP7 inhibitors with different scaffolds as well as PROTACs targeting USP7 have been developed for application in leukemia cells^19,22,23,39^, matched inhibitor-PROTAC pairs suitable for comparative studies in solid tumors have remained absent.

Here, we employ an integrated chemical biology approach to provide a comprehensively characterized matched pair of USP7 modulators, enabling systematic comparison of protein inhibition versus degradation (**Fig. 1a**). Our work establishes a framework for understanding DUB biology through complementary chemical perturbation approaches and provides validated tools for the systematic investigation of USP7.

**Figure 1.**
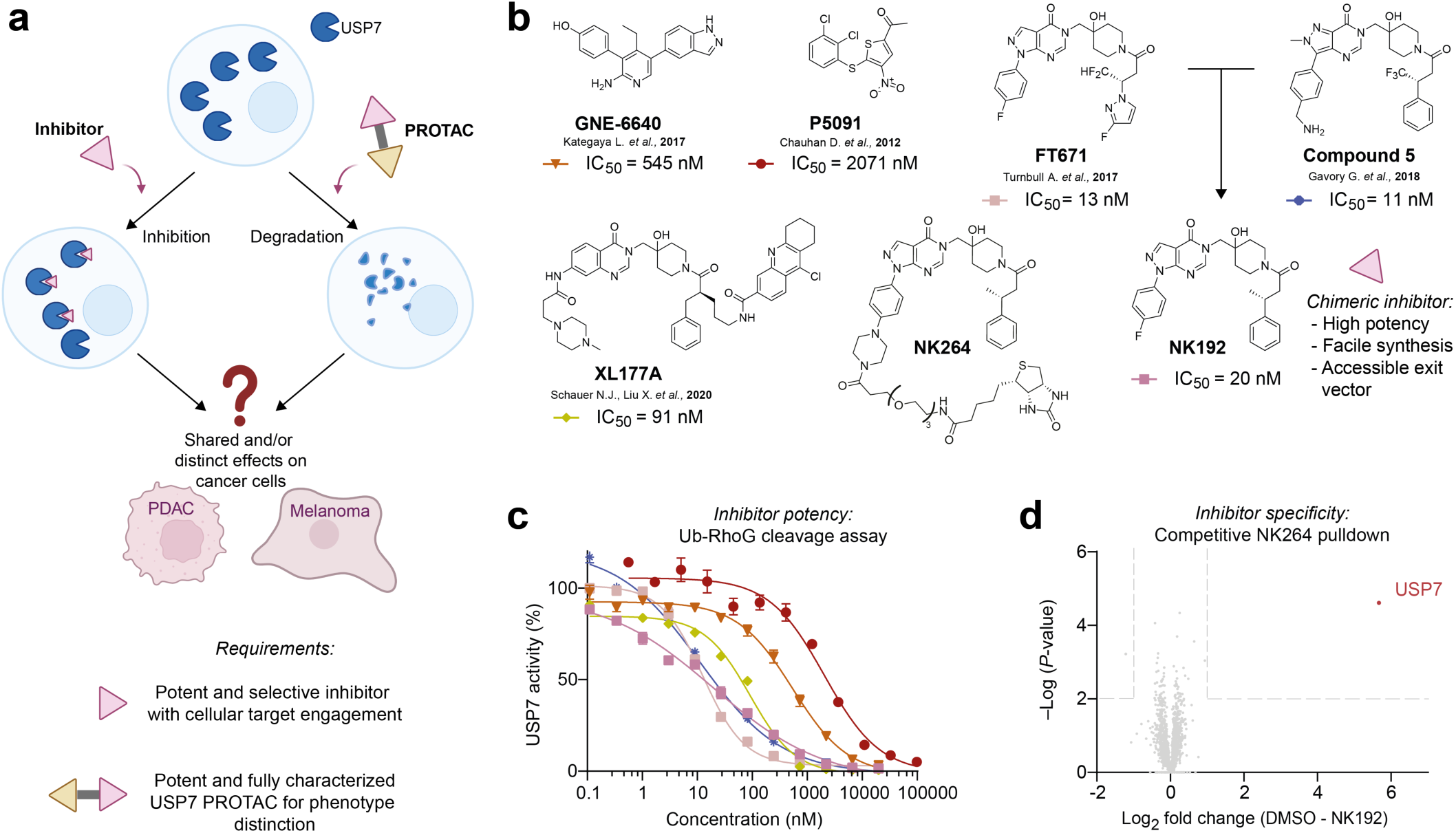
Towards a comparative analysis of small molecule-mediated USP7 inhibition versus USP7 degradation phenotypes. **a.** Outline of this study. Comparative analysis of USP7 inhibition or degradation in PDAC and melanoma will be carried out using customized inhibitors and degraders. **b.** Chemical structures of previously described USP7 inhibitors as well as of NK192 (chimera derived from potent USP7 inhibitors FT671 and Compound 5) which was used as USP7 ligand for the PROTAC library synthesis. Half-maximal inhibition (IC_50_) values were derived from data shown in panel c. NK264 represents a Biotin-functionalized variant of NK192, which was used for pulldown experiments shown in panel d. **c.** Ubiquitin-Rhodamine110Gly (Ub-RhoG) cleavage assay to determine inhibitory potency of compounds shown in b. Full length human USP7 was incubated with the respective compounds, the fluorogenic substrate was added and residual activity was read out through fluorescence measurements. **d.** Competitive pulldown experiment to assess proteome-wide specificity of NK192. Biotin-functionalized NK264 was immobilized on beads. Proteins enriched from Panc89 lysate either treated with DMSO or with NK192 (10 µM, 500x IC_50_) were quantified by mass spectrometry.

## Results

### NK192 is a potent USP7 inhibitor with proteome-wide specificity and a functional exit vector

To develop a validated, specific and potent USP7 inhibitor and degrader pair, we first assembled a small collection of previously reported, structurally diverse USP7 inhibitors (**Fig. 1b**).^17–19,29,40^ Using the fluorogenic substrate Ubiquitin-Rhodamine110Gly (Ub-RhoG), we confirmed their inhibitory potency on recombinant, full-length USP7 (**Fig. 1c**). We focused on FT671 and Compound 5, which both feature a hydroxypiperidine core, display high inhibitory potency in vitro and are suitable for cellular use.^17,19^ To facilitate the streamlined synthesis of USP7 degrader candidate molecules, we devised chimeric inhibitor NK192 which retains the 4-fluorophenyl-pyrazolo[3,4-*d*]pyrimidinone scaffold of FT671 and comprises (*R*)-3-phenylbutanoic acid as a fluorine-free equivalent of this motif in Compound 5. Chiral NK192 was readily prepared in four steps from commercially available starting materials (**Supporting Fig. 1a**) and retained the high inhibitory potency on USP7 (**Fig. 1c**).

Both FT671 and Compound 5 are highly specific for USP7 when assayed against related human USP family DUBs^17,19^, yet whether they can potently bind non-DUB proteins had not yet been investigated. We therefore set out to confirm the target specificity of our chimeric USP7 inhibitor NK192 on a proteome-wide scale. Inspection of the crystal structure of USP7 in complex with FT671 revealed that the 4-fluorophenyl group pointed away from the catalytic domain^19^ and suggested this site for chemical derivatization (**Supporting Fig. 1b**). By replacing the fluorine with a piperazine linker, we next prepared the biotin-functionalized, NK192-based probe NK264 **(Fig. 1b**) which still potently engaged with USP7 (**Supporting Fig. 1c**). To comprehensively assess cellular targets of NK192, we conducted a competitive pulldown experiment. Lysates of Panc89 cells were treated either with DMSO or with a high concentration of NK192 (10 µM, 500x IC_50_), followed by enrichment with NK264-functionalised streptavidin beads and mass spectrometric analysis. Our data revealed that NK192 exhibited very high specificity for USP7 (**Fig. 1d**) as USP7 was the only protein that was significantly competed away from the beads by NK192. These data established NK192 as a suitable, potent and specific ligand for the exploration of USP7 degraders to arrive at a chemically corresponding USP7 inhibitor and degrader pair. Moreover, the biotin probe yielded a functional exit vector in NK192 for the synthesis of bifunctional molecules.

### Identification of degraders for USP7 depletion in PDAC and melanoma

To arrive at a chemically diverse library of degrader candidate molecules, which would facilitate degradation of USP7 through induced proximity to the VHL E3 ligase^3^, we employed a modular synthesis approach (**Fig. 2a**). Starting from a bromo-substituted NK192-equivalent as the USP7 ligand, we appended various mono-protected bifunctional amine-containing heterocycles via Buchwald-Hartwig cross coupling reactions (**Fig. 2a**, step A). Following cleavage of the protecting groups, building blocks were either directly coupled to VHL ligand-linker conjugates (step B) or were further functionalized with linkers (step C), which were then fused to VHL ligand-linker conjugates (step D). After a first set of USP7 degrader candidates designed with long linear linkers did not yield promising degradation of USP7, we redirected our efforts toward a second-generation library of 15 compounds featuring mostly rigidified linkers (**Fig. 2b**). We aimed to limit flexibility between both ligands to enhance ternary complex formation and thereby degradation efficiency^41^. Synthesized compounds thus included a diverse set of hetero-, bi- and spiro-cyclic linkers, covering a wide variety of rigidity, geometry and linker length. Molecules comprising sterically demanding cubane and cyclohexyl groups directly adjacent to the VHL ligand building block were motivated by the presence of a hydrophobic pocket in the VHL E3 ligase which can be addressed by a cyclopropyl motif (e.g. in the VHL inhibitor VH298)^42^. Compounds were prepared with defined stereochemistry, purified by preparative reverse phase HPLC and subsequently used in cellular assays (see the Supporting Information for chemical synthesis and compound characterization data).

**Figure 2.**
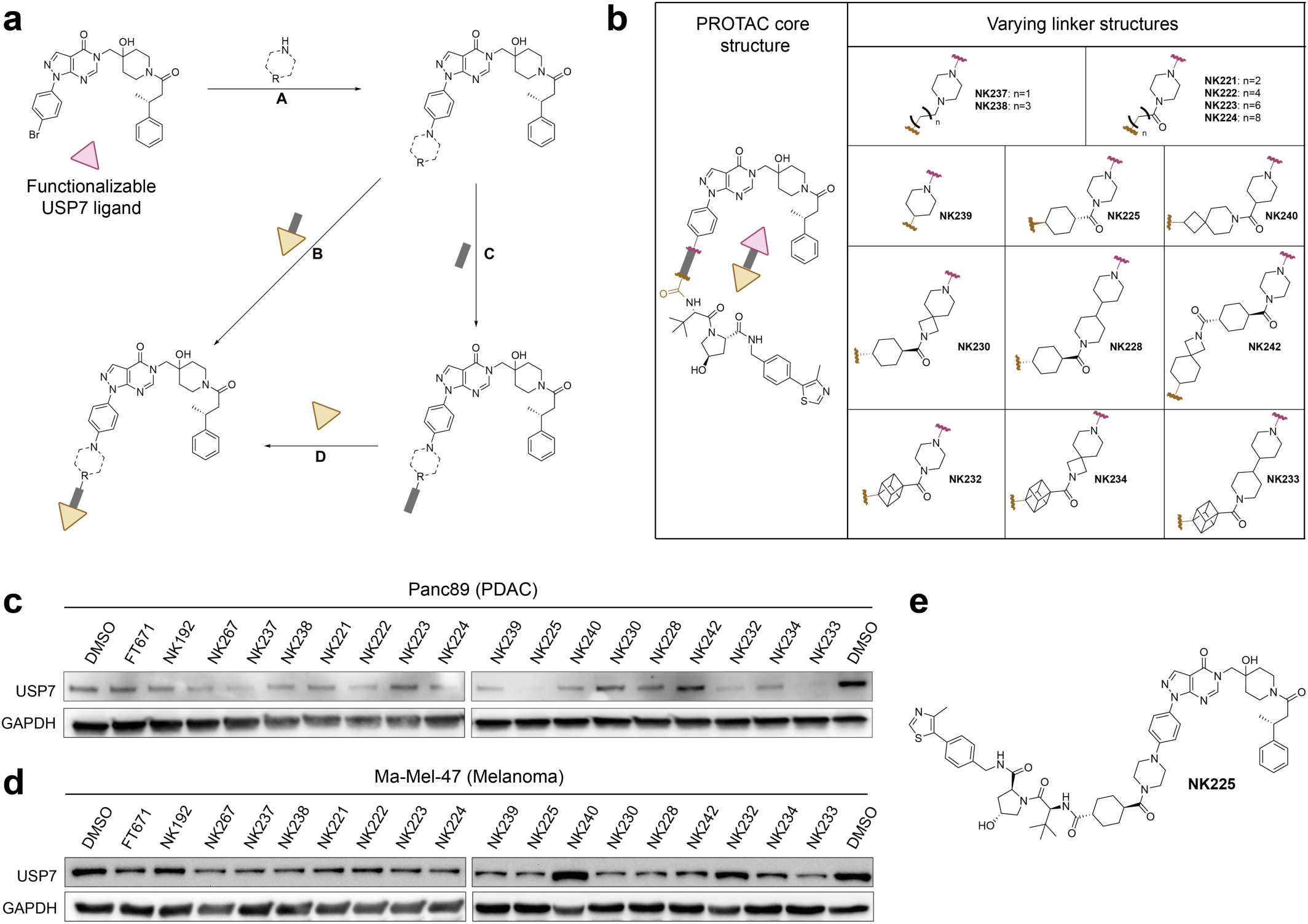
Design and synthesis of USP7 targeting PROTACs. **a.** Schematic of the synthesis of USP7-targeting degraders. Functionalized USP7 ligands (top left, pink triangle) were either coupled directly to VHL-linker conjugates (yellow triangle with grey stick, through steps A and B) or first further functionalized with linkers followed by coupling to the VHL ligand (yellow triangle, through steps A, C and D). See the Supporting Information for detailed chemical procedures. **b.** Chemical structures of USP7-targeting degrader library. A general structure of the PROTACs is shown on the left, consisting of a USP7 binding ligand (top) and E3 ligase binding ligand (bottom) separated by a linker. Linker attachment points to both ligands are indicated by waved lines (black: USP7, red: VHL). Linkers are shown in the table on the right. **c.-d.** Assessment of USP7 levels upon treatment with compounds in Panc89 cells (c) and Ma-Mel-47 cells (d). Cells were treated with 5 µM of indicated compounds for 24 h, and cell lysates were analyzed through Western blots with indicated antibodies. See the Supporting Information for uncropped blots. **e.** Chemical structure of NK225.

To identify degraders for USP7 depletion in PDAC and melanoma, we treated both Panc89 PDAC and Ma-Mel-47 melanoma cells with compounds for 24 hours and assessed USP7 protein levels by Western blot (**Fig. 2c-d**). We identified several PROTACs that effectively degraded USP7 in Panc89 cells, including NK225 (**Fig. 2e**) and NK233. While the degradation efficiency in Ma-Mel-47 was comparably lower, we also identified robust degradation of USP7 by NK225 and NK233. These compounds comprise a trans-configured cyclohexane-1,4-dicarboxylic acid (NK225) or cubane-1,4-dicarboxylic (NK233) as well as a piperazine (NK225) or 4,4’-bipiperidine (NK233). Taken together, these results showed that USP7 PROTACs with different linker geometries facilitated degradation of USP7 with notable efficiency differences between both investigated cell lines.

### Characterization of two improved USP7 degraders customized to PDAC and melanoma cell lines

Encouraged by these results, we set out to further optimize these two degraders. To this end, we added a benzylic (*S)-*methyl group to the VHL ligand, which has previously been shown to improve affinity towards VHL^43^. This modification resulted in the third-generation PROTACs NK250 (derived from NK225) and NK266 (derived from NK233) (**Fig. 3a**). Additionally, we synthesized an NK225-based negative control compound NK245 (**Fig. 3a**), which cannot bind VHL due to an inverted stereocenter at the hydroxyproline. Next, we treated both cell lines with these compounds and assessed USP7 levels by Western blot analysis (**Fig. 3b**). Both PROTACs NK250 and NK266, but not NK245, showed improved USP7 degradation in both cell lines. Consistently, cyclohexyl-PROTAC NK250 showed more complete degradation in Panc89 cells, whereas cubane-containing PROTAC NK266 showed higher potency in Ma-Mel-47 cells.

**Figure 3.**
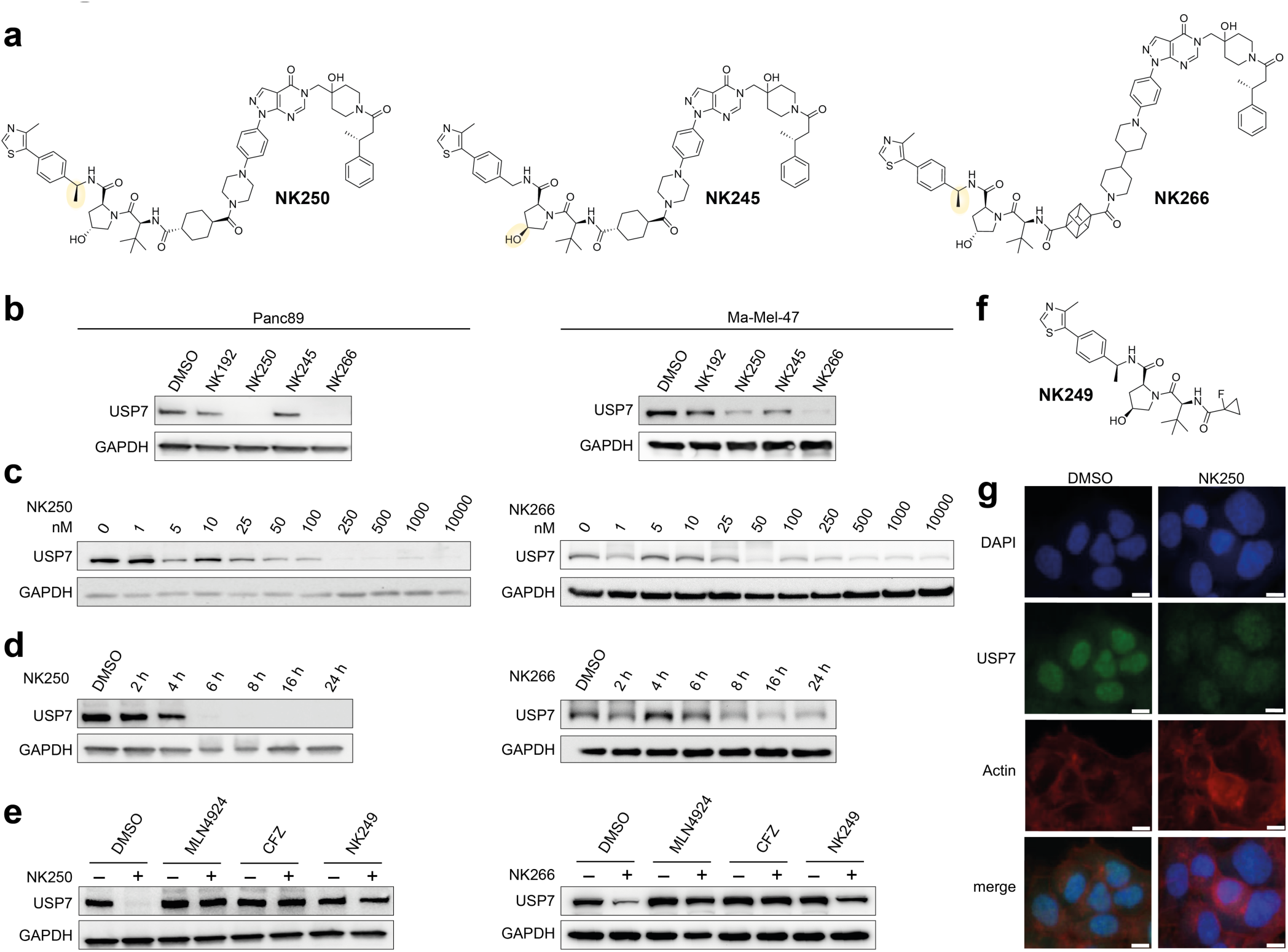
Characterization of two potent USP7 degraders. **a.** Chemical structures of improved VHL-targeting PROTACs NK250, NK266 and non-VHL binding control compound NK245. The additional methyl groups in the VHL ligand in NK250 and NK266 (leading to enhanced VHL binding) as well as the inversed hydroxyproline stereocenter in NK245 (abrogating VHL binding) are highlighted. **b.** USP7 degradation assay. Panc89 cells (left) or Ma-Mel-47 cells (right) were treated with 5 µM of indicated compounds for 24 h and analyzed by Western blot. See the Supporting Information for uncropped blots. **c.** Assessment of USP7 degradation efficiency. Panc89 cells (left) or Ma-Mel-47 cells (right) were treated with increasing concentrations of either NK250 (left) or NK266 (right). **d.** Determination of degradation kinetics. Panc89 cells (left) or Ma-Mel47 cells (right) were treated with of 1 µM of NK250 (left) and NK266 (right) for indicated times. **e.** Degradation rescue experiment. Panc89 cells (left) or Ma-Mel47 cells (right) were pretreated for 2 h with either DMSO, NEDDylation inhibitor MLN4924 (500 nM), proteasome inhibitor Carfilzomib (CFZ, 250 nM) or VHL ligand NK249 (10 µM) followed by 1 µM of NK250 or N266 for 20 h. **f.** Chemical structure of VHL ligand NK249. **g.** Orthogonal confirmation of USP7 depletion by immune fluorescence. Panc89 cells were treated with 1 µM NK250 for 24 h before staining for USP7 (green), actin (red), and with DAPI (blue). Scale bar: 10 µm.

To carry out an in-depth evaluation of both compounds, we assessed the degradation potency in a concentration-dependent manner (**Fig. 3c**). Therefore, we treated both Panc89 and Ma-Mel-47 cells for 24 hours with varying compound concentrations. We observed near complete degradation of USP7 with PROTAC NK250 in Panc89 cells already at a PROTAC concentration of 250 nM (**Fig. 3c**, left), whereas NK266 showed a strong reduction of USP7 levels at higher concentrations of around 500-1000 nM in Ma-Mel-47 cells (**Fig. 3c**, right). Notably, NK266 also showed similar degradation efficiency as NK250 in Panc89 cells, while NK250 was only able to fully degrade USP7 at 1 µM concentration in Ma-Mel-47 cells after 24 hours (**Supporting Fig. 2a**). Based on these data, we decided on a concentration of 1 µM compound for all following cell-based experiments and retained the use of NK250 for Panc89 and NK266 for Ma-Mel-47 cells.

To probe degradation kinetics, both cell lines were incubated with PROTACs, and USP7 levels were followed over 24 h (**Fig. 3d**). Treatment with NK250 resulted in rapid degradation of USP7 in Panc89 cells with protein depletion observed already after 6 hours. In contrast, NK266 showed potent degradation after 16 hours in Ma-Mel-47 cells (**Fig. 3d**), while the negative control NK245 did not reduce protein levels in both cells (**Supporting Fig. 2b**). To confirm the mode of action of our degraders, we performed rescue experiments in which we blocked the degradation machinery involved in VHL-mediated protein depletion (**Fig. 3e**). Consistently, pretreatment of cells with either the NEDDylation inhibitor MLN4924 (which abrogates VHL E3 ligase activity), the proteasome inhibitor carfilzomib or a tenfold excess of VHL ligand NK249 (**Fig. 3f**, to block VHL engagement) rescued USP7 degradation. In addition, we confirmed USP7 degradation by immune fluorescence imaging in Panc89 cells (**Fig. 3g**). We detected a signal for USP7 predominantly localized to the nucleus, consistent with literature on USP7 localization^44^, which was strongly reduced after treatment with NK250. Together, these data demonstrated potent depletion of USP7 by the PROTACs NK250 and NK266 and confirm their mode of action as Cullin-Ring-E3 ligase (CRL)-dependent degraders.

### Quantitative assessments of USP7 degradation efficiency and PROTAC-mediated ternary complex formation

We next aimed to quantitatively investigate the mechanism driving their distinct degradation kinetics. We first established an additional orthogonal approach to measure cellular degradation efficiencies based on the HiBiT technology (**Fig. 4a**).^45^ To this end, we created an MV4-11 cell line which stably expresses N-terminally HiBiT-tagged USP7. The 11 amino acid HiBiT-tag can bind the LgBiT polypeptide, which results in an active luciferase, generating a signal proportional to the amount of intact HiBiT-USP7 protein (**Fig. 4a**). We treated these cells with either NK250 or NK266 and observed dose-dependent reduction of USP7 levels after 6 and 24 hours (**Fig. 4b**). In line with the results from Western blots, NK250 showed higher potency towards HiBiT-USP7 than NK266, particularly at the 6-hour time point. Here, NK250 was able to achieve a maximum degradation efficiency (D_max_) of 87 %, while NK266 achieved a lower 74 % maximum degradation. Treating cells for 24 hours resulted in the same near complete degradation (D_max_ = 90-92 %). This was accompanied by low half-maximal degradation concentration values (DC_50_) values of 4 nM and 24 nM for NK250 and NK266, respectively, demonstrating excellent degradation efficiency.

**Figure 4.**
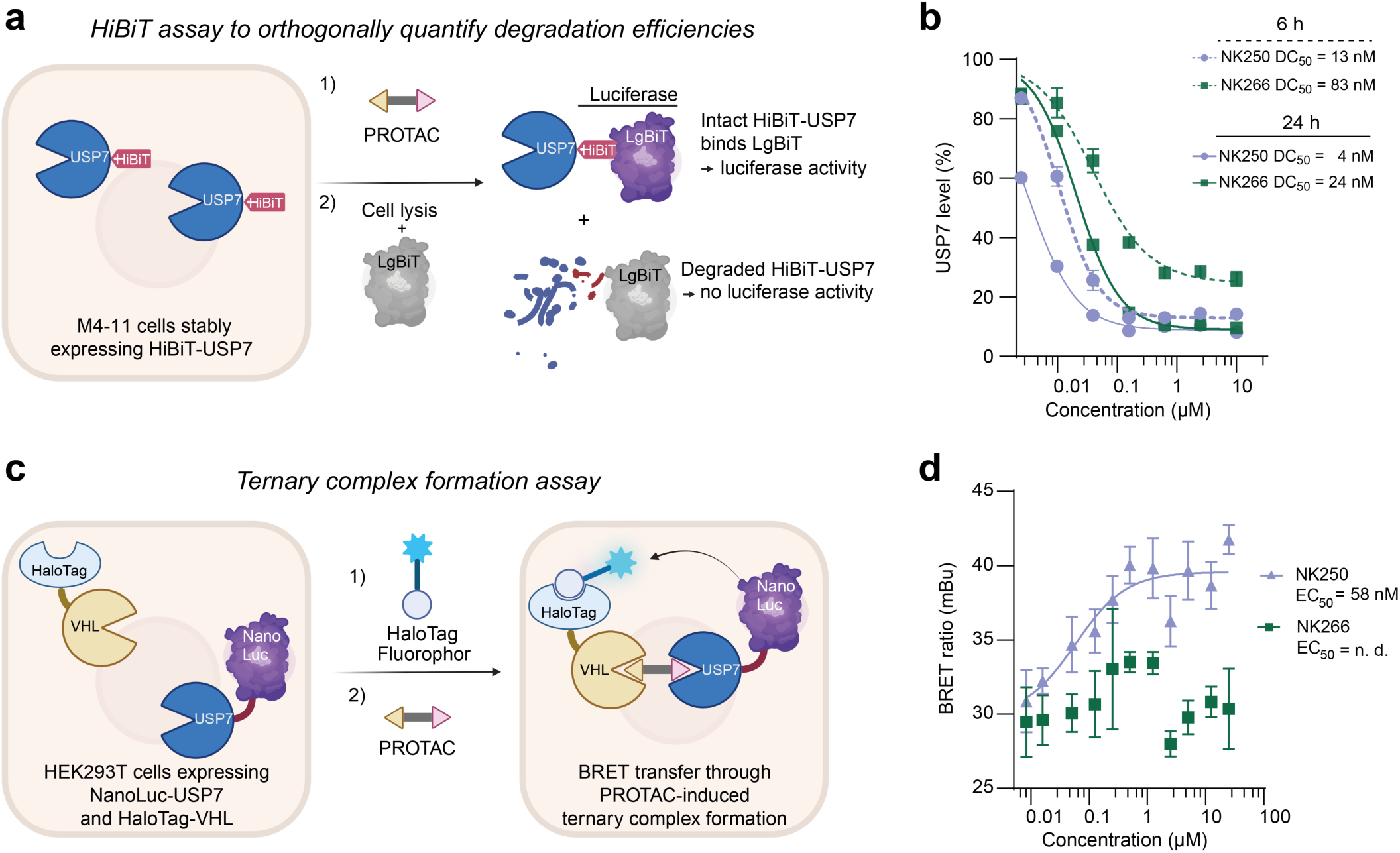
Quantitative assessments of USP7 degradation efficiency and ternary complex formation. **a.** Schematic representation of HiBiT endpoint degradation assay. **b.** HiBiT-based quantification of USP7 levels. MV4-11 cells stably expressing HiBiT-USP7 generated through lentiviral transduction were treated with PROTACs for either 6 or 24 h. Remaining HiBiT-USP7 could be detected through the luciferase signal using HiBiT Lytic Detection System. Data are shown as mean ± S.D. (N=3), normalized to DMSO treatment. Half-maximal degradation concentrations (DC_50_) derived from these data are given. **c.** Schematic representation of a cellular ternary complex formation assay. Halotag-VHL- and NanoLuc-USP7-expressing cells are sequentially treated with Halotag-Fluorophor and PROTAC. Compound-induced proximity is read out by a bioluminescence resonance energy transfer (BRET) signal as shown, demonstrating ternary complex formation. **d.** Cellular ternary complex formation assay. HEK293T cells overexpressing Halotag-VHL and NanoLuc-USP7 were treated with HaloTag NanoBRET 618 Ligand for 20 h, followed by NK250 or NK266 treatment at indicated concentrations for 2 h prior BRET measurement. The determined half-maximal ternary complex formation concentration (EC_50_) for NK250 is given. Data are shown as mean ± S.D. (N=4). mBu, milli BRET units.

Notably, we observed a partial reduction of USP7 levels with both the negative control PROTAC NK245 and the inhibitor NK192 (**Supporting Fig. 3a-b**). This phenomenon is consistent with the previously observed auto-regulation of USP7 ubiquitination^28,46^, by which the DUB can self-regulate its abundance through auto-deubiquitination. Importantly, PROTAC-mediated degradation depleted USP7 levels more potently (DC_50_ and D_max_) in all cell lines tested (**Fig. 2c-d**, **Fig. 3b, Supporting Fig. 3a-b**). Equivalent assays with second-generation compounds showed NK225 and NK233 to mimic NK250 and NK266 in degradation efficiency and kinetics with lower potency (**Supporting Fig. 3a-b**).

Next, we aimed to understand the molecular basis for the different degradation kinetics of cyclohexyl-(NK225, NK250) and cubane-containing (NK233, NK266) compounds. The formation of a stable ternary substrate-PROTAC-E3 ligase complex in cells is increasingly recognized as central for rapid targeted protein degradation.^47^ Therefore, we employed a cellular ternary complex formation assay to determine the ability of PROTACs to induce proximity between USP7 and VHL (**Fig. 4c**). We overexpressed both NanoLuc-USP7 and HaloTag-VHL in HEK293T cells and treated them with both a HaloTag-fluorophore and our PROTACs. Upon induction of a ternary complex, proximity between both proteins was recorded by measuring the resulting bioluminescence resonance energy transfer (BRET) from the NanoLuc luciferase to the VHL-located fluorophore.^48^ In this assay, only NK250 and NK225 were able to potently induce a stable ternary complex between the PROTAC, USP7 and VHL with half maximal effective concentrations (EC_50_) of 58 nM and 144 nM, respectively, while treatment with NK266 or N233 did not increase the BRET signal (**Fig. 4d, Supporting Fig. 3c**). Further, we verified that neither the negative control NK245 nor the VHL ligand VH298 induced a ternary complex. This potent induction of a stable ternary complex by NK250 but not NK266 provided a rational for the more rapid USP7 degradation facilitated by NK250 within 6 hours versus 16-24 hours by NK266. Collectively, these assays validated NK250 and NK266 as potent degraders of USP7, which enable the formation of validated and cell line-customized USP7 inhibitor and degrader pairs to interrogate cellular roles of USP7 in PDAC and melanoma.

### Proteome-wide analysis reveals strikingly disparate effects of USP7 modulation through inhibition versus degradation

Having established a chemical toolbox consisting of USP7-specific inhibitor and PROTAC pairs with chemical complementarity, we set out to examine consequences of USP7 modulation by directly comparing USP7 inhibition and degradation. For this we treated both Panc89 and Ma-Mel-47 cells with NK192 and PROTACs NK250/NK266 for 6, 24 or 72 hours and recorded changes in the cellular proteome by mass spectrometry. Analysis of the Panc89 samples through data-dependent acquisition (DDA) on an Orbitrap Fusion Lumos mass spectrometer resulted in quantitation of 4214 unique protein groups (**Supporting Fig. 4a-f**, **Supporting Tables 2-6**). The experiment demonstrated the exquisite specificity of NK250 for USP7 degradation as USP7 was the only protein significantly decreased after 6 h (**Supporting Fig. 4a**). Moreover, USP7 protein levels decreased further in a time-dependent manner, leading to a more than 100-fold reduction after three days (**Supporting Fig. 4b-c**). In contrast, USP7 inhibitor NK192 did not lead to USP7 protein level changes beyond the significance threshold (**Supporting Fig. 4d-g**). However, we were surprised to observe only very few other proteins changing in their abundance within 24 h, including the previously validated USP7 substrate TRIP12.^15^

To enable greater coverage, we thus adopted a data-independent acquisition (DIA) proteomics strategy on the same spectrometer.^49^ Re-analysis of the same Panc89 samples with a DIA protocol allowed the identification and quantitation of 8828 unique protein groups (**Fig. 5a-f**, **Supporting Tables 2,4**), thereby more than doubling the return of the DDA analysis. Analysis of the Ma-Mel-47 cell samples using DIA yielded quantitative information on 9344 proteins (**Fig. 5g-l**, **Supporting Tables 2,5**), demonstrating a comprehensive coverage of the cellular proteome in both cell lines. These DIA data were then analyzed further (**Fig. 5a-l**, **Supporting Fig. 5a-d**).

**Figure 5.**
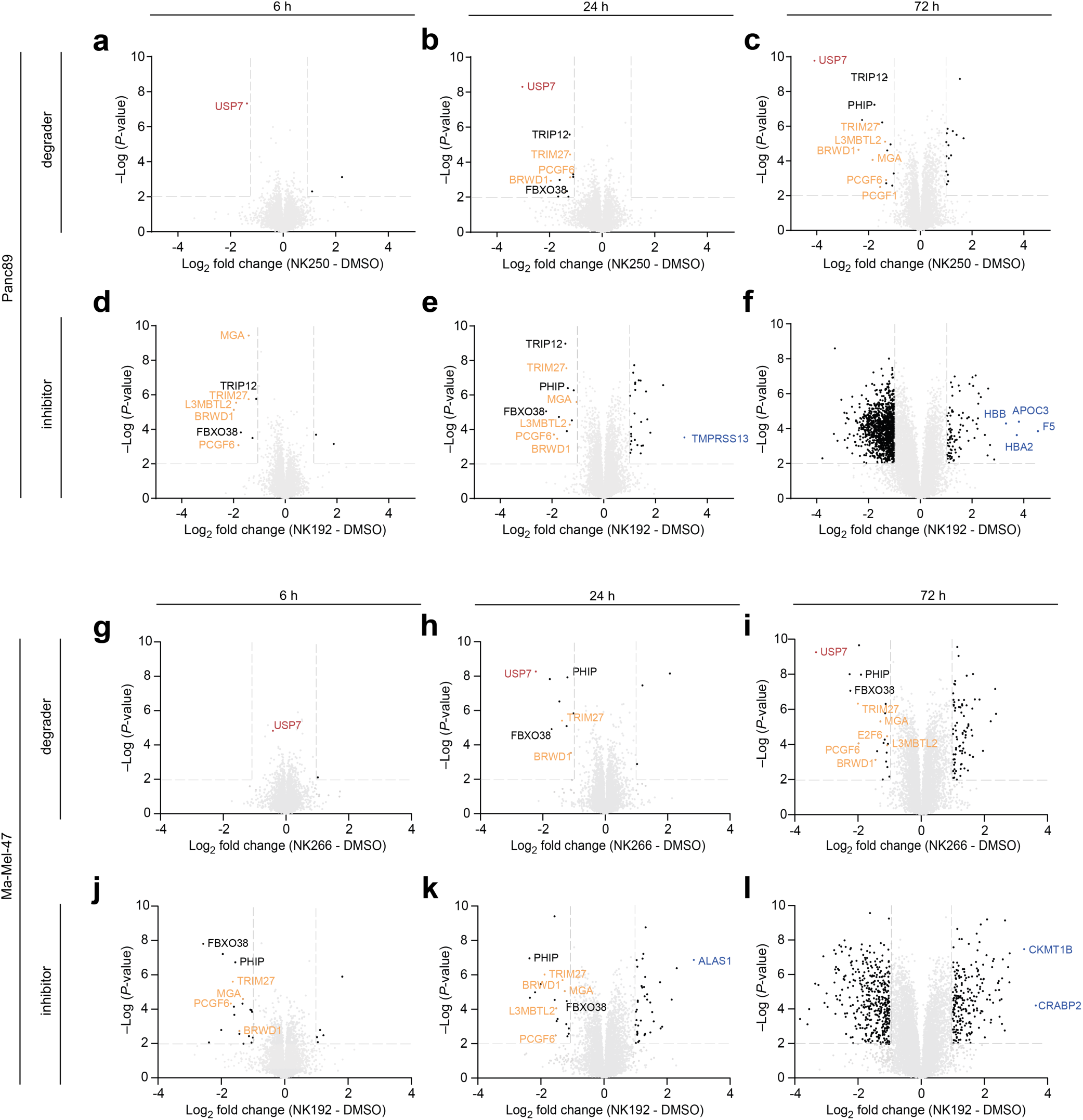
Proteome-wide analysis reveals strikingly disparate effects of USP7 modulation through inhibition versus degradation. a.-f. Proteomic analysis of Panc89 cells treated with PROTAC NK250 (a-c) or inhibitor NK192 (d-f) for indicated time points. Volcano blots report proteins identified by data-independent acquisition mass spectrometry in Panc89 cells treated with NK192 (5 µM) or NK250 (1 µM) for 6, 24 or 72 h. Proteins annotated as members of the non-canonical repressive complex 1.6 are highlighted in orange, chromatin bound E3 ligases are highlighted in black, the most upregulated proteins by USP7 inhibition in blue. **g.-l.** Proteomic analysis of Ma-Mel-47 cells treated with PROTAC NK266 (g-I, 1 µM) or inhibitor NK192 (j-l, 5 µM) for indicated time points.

As observed in the Panc89 DDA dataset, PROTAC treatment of Panc89 and Ma-Mel-47 cells resulted in significant and consistent USP7 depletion (**Supporting Fig. 4g)**. USP7 was the only protein that was significantly decreased in Panc89 cells after 6 h of NK250 treatment, further highlighting the specificity of our degrader (**Fig. 5a**). In accordance with the observations made through Western blot analysis, USP7 exhibited only minimal reduction in Ma-Mel-47 cells following 6 hours of NK266 treatment (**Fig. 5g**). After 24 h, we identified USP7 to be the most decreased protein, thereby showing that also NK266 potently and specifically, yet more slowly, degrades USP7 in melanoma cells (**Fig. 5h**).

Moreover, we identified several established USP7 substrates to be decreased in their abundance following PROTAC treatment in both cell lines (**Fig. 5a-c,g-i**, **Supporting Fig. 5a**), including chromatin-bound E3 ligases (TRIP12, TRIM27)^15^, members of the non-canonical Polycomb repressive complex 1.6 (PRC1.6) (MGA, L3MBLT2, BRWD1, PCGF6)^14^ and diverse other proteins including a helicase (DHX40) and E3 ligase substrate adaptors (FBXO38, PHIP)^27,50^. A smaller number of proteins were found to be increased following PROTAC treatment in both entities, which was particularly evident after 24 h of treatment (**Fig. 5b-c,h-i**). The limited number of regulated proteins shared by both cell lines (**Supporting Fig. 5a**), the majority of which are previously reported substrates of USP7, was unexpected given the near-complete depletion of this DUB, its multiple cellular roles and its ability to broadly regulate protein ubiquitination and protein stability.^14,19,20,27,28,46^ Importantly, the lack of regulation in these cell lines of several previously reported substrates (including MDM2 and FOXO4, which gave rise to USP7 modulation to be pursued as a broadly applicable anti-cancer strategy^13,19,20,22,29,40^), as well as the large number of proteins changed only in one of the two investigated cell lines strongly hint at pronounced cell-line-specific differences in USP7 functions. Our findings thereby underscore the value of selective depletion of USP7 to investigate substrates and downstream effects in the cellular system of interest.

Treatment of Panc89 or Ma-Mel-47 cells with USP7 inhibitor NK192 for 6 h or 24 h resulted in similar sets of proteins being regulated as observed with USP7-degrader-treated cells (**Fig. 5d-e,j-k**, **Supporting Fig. 5b-d**). These findings robustly highlight the role of USP7 in regulating the amounts of several E3 ligase proteins as well as of PRC1.6 complex members as a relatively small set of core substrates. Moreover, the pronounced differences between USP7 enzyme inhibition and protein degradation in both cell lines (**Supporting Fig. 5c-d**) stress the added value of degrader and inhibitor pairs to broadly uncover target-dependent cellular processes.

Strikingly, however, inhibitor treatment for 72 h induced drastic differences in the proteomes of Panc89 and Ma-Mel-47 cells. We found hundreds of proteins being either up- or downregulated (**Fig. 5f,l**), a finding that contrasts with the comparably low number of proteins affected by PROTAC treatment (**Fig. 5c,i**) across all time points. This large-scale, significant proteomic rewiring observed after 72 h of inhibitor treatment, but not with PROTAC treatment consistently in both cancer types, highlights the value of targeted protein degradation. The enhanced selectivity conferred through the PROTAC will greatly aid studying USP7’s functions and substrates with chemical modulators. While cellular effects specific to inhibited USP7-containing PRC complexes have been described^14^, the large scale of these changes and their induction of proteins not normally associated with either PDAC or melanoma cells (incl. the haemoglobin subunits HBB and HBA2) suggest a dysregulation through USP7-independent pleiotropic effects conferred by the inhibitor at longer treatment times.

### PROTAC-induced degradation of USP7 uncovered USP7-independent glucose dependency induced by USP7 inhibitors

Our proteomic data suggested that cells treated with USP7 inhibitor NK192 for prolonged times should show phenotypes distinct from USP7 depletion by the developed PROTACs. Indeed, when various PDAC and melanoma cell lines were treated with USP7 modulators, we observed a strong switch in the cell culture media color for the inhibitor-treated, but not for the degrader-treated samples. In line with the inhibitor-induced proteomic dysregulation observed after 3 days in both cell lines, these data pointed towards distinct phenotypic effects. To investigate this disparity systematically, we treated Panc89 cells with 1-10 µM of USP7 inhibitors FT671, NK192 and the PROTAC NK250 in standard complete DMEM medium containing either 25 mM (“high glucose”) or 5 mM (“low glucose”) D-glucose (**Fig. 6a**). In addition, cell growth was recorded after 72 h which was not affected in all conditions (**Supporting Fig. 6a**). Interestingly, the color change of the media occurred only in high glucose cell culture media, but not in low glucose media, indicating an acidification by increased nutrient consumption of inhibitor-treated cells. The same effect was observed in the melanoma cell line Ma-Mel-47 growing in more lightly colored RPMI 1640, when the standard media conditions with 11 mM D-glucose were compared to those with 5 mM D-glucose (**Supporting Fig. 6b-c**). Complete depletion of USP7 in PROTAC-treated cells within 72 hours for both Panc89 (**Fig. 6b**) and Ma-Mel-47 (**Supporting Fig. 6d**) confirmed that the observed phenotypic disparity was not due to insufficient USP7 degradation. Interestingly, while both inhibitors and their respective degraders did not (Panc89, **Supporting Fig. 6a**) or only mildly (Ma-Mel-47, **Supporting Fig. 6b**) affect cell viability (assessed through confluence) in high glucose media, inhibitor treatment but not PROTAC treatment in low glucose media strongly induced apoptosis in Panc89 (**Fig. 6c**) and Ma-Mel-47 cells (**Supporting Fig. 6e**). These data indicated a higher glucose dependency or glucose consumption only due to inhibitor treatment.

**Figure 6.**
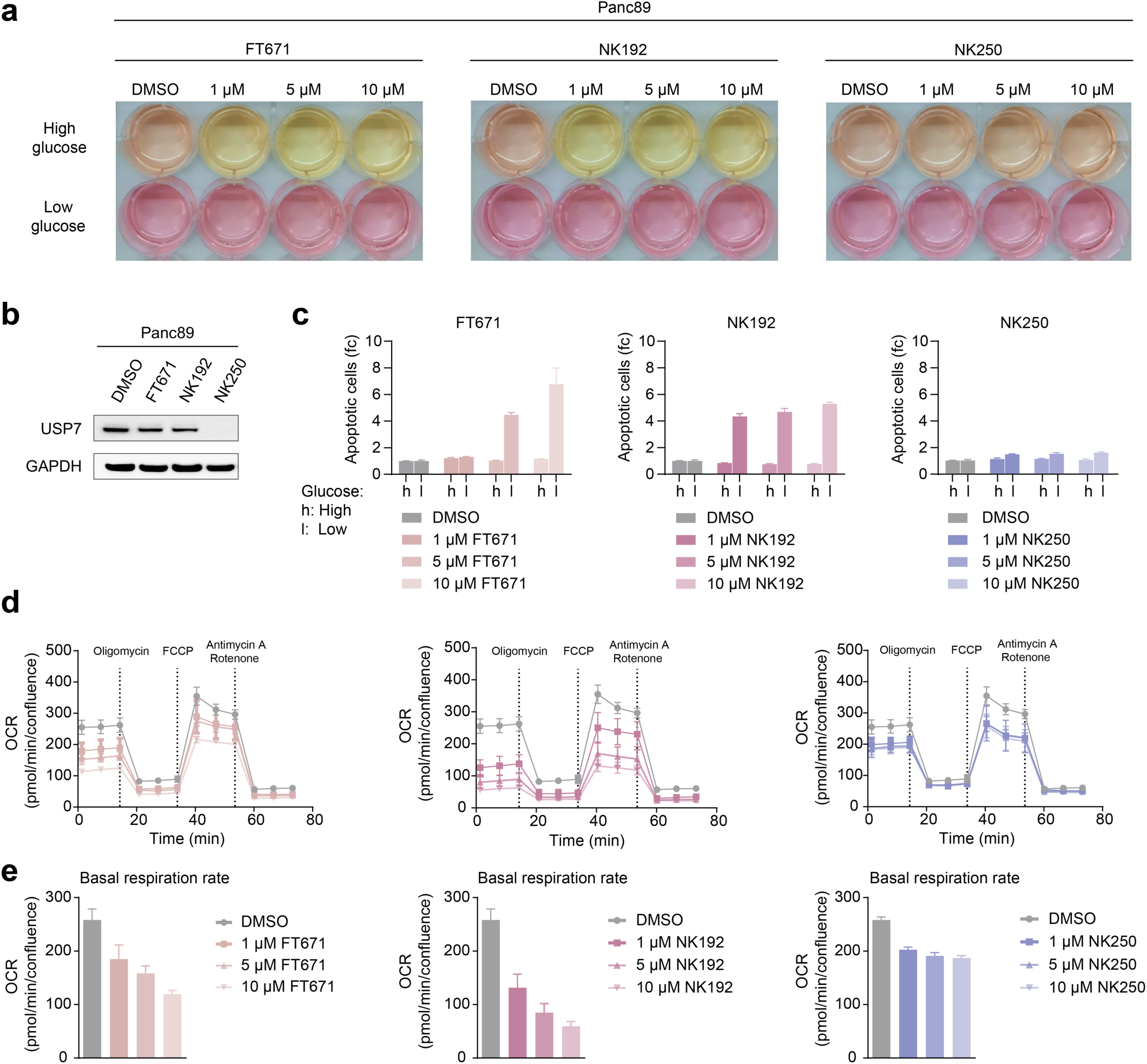
Disparate metabolic effects of USP7 inhibitor vs. PROTAC treatment. **a.** Images of Panc89 cells in culture medium containing 5 mM (low) or 25 mM (high) glucose, treated with indicated concentrations of FT671, NK192 or NK250 for 72 h. **b.** Western blot analysis of Panc89 cells treated with 1 µM of indicated compounds for 72 h. **c.** Apoptosis assay. Panc89 cells treated with compounds for 72 h in culture medium containing 5 mM (low) or 25 mM (high) glucose were analyzed by FACS for Annexin-V staining. Data were normalized to untreated controls; one representative experiment of at least three independent experiments with technical triplicates is shown. Data are shown as mean ± SEM. fc, fold change. The same DMSO samples are repeated for clarity in each graph. **d.** Seahorse mito stress test. Panc89 cells treated with indicated concentrations of FT671, NK192 or NK250 for 24 h, followed by measurements of their oxygen consumption rate (OCR). Three independent experiments were performed in technical quintuplicates, one representative is shown as mean ± SEM. **e.** Basal respiration rates of data shown in panel d, shown as mean ± SEM.

To further dissect this inhibitor-induced effect, we quantified cellular oxygen consumption rates (OCR) by seahorse assays. Whereas a degrader-mediated effect on OCR was much smaller and plateaued already at low doses where USP7 was fully degraded (**Fig. 6d-e, Supporting Fig. 6f-g**), USP7 inhibitors facilitated a persistent reduction of mitochondrial OCR in a dose-dependent manner. In both FT671- and NK192-treated cells, the basal respiration rate measured in confluence-adjusted OCR dropped far below 200 pmol/min/confluence in both cell lines (**Fig. 6d-e, Supporting Fig. 6f-g**). This reduced capacity of ATP production through oxidative phosphorylation is consistent with a more anaerobic glucose metabolism in inhibitor-treated cells. This metabolic switch in turn explains the glucose-dependent acidification seen in culture medium (**Fig. 6a**), whereas specific USP7 degradation upon PROTAC treatment in both PDAC and melanoma cells, through retained respiration capacity, did not induce glucose-dependency (**Fig. 6d, Supporting Fig. 6f**). Taken together, these data highlighted the value of highly specific USP7 degraders for distinguishing genuinely on-target USP7-mediated effects from possibly non-specific, metabolic phenotypes induced even by well-characterized inhibitors. These findings thus have important implications for the cellular use of USP7 inhibitors FT671 and NK192 under low glucose conditions or with prolonged treatment times. More broadly, our data showcases the advantages of the PROTAC approach for the dissection of cellular roles of a key deubiquitinase through providing highly specific degraders.

## Discussion

The development of matched inhibitor–degrader pairs marks a critical step forward in dissecting the multifaceted cellular roles of deubiquitinating enzymes (DUBs). Here, we present a comprehensive chemical biology framework for the interrogation of USP7 through PROTAC degraders and inhibitors. This dual-modality toolkit enables temporal and mechanistic analysis of USP7 functions across cancer types, while mitigating some of the limitations of classical inhibitor-based approaches.

VHL-based PROTACs NK250 and NK266 demonstrated potent and near-complete USP7 degradation in PDAC and melanoma cells, respectively. Degradation kinetics correlated with ternary complex formation, and the choice of VHL as E3 ligase proved strategic: unlike ligands for the Cereblon ligase^51^, ligands for VHL do not induce activation of the p53–p21 axis. Given that USP7 activity regulates p21 in some cell lines,^19,29^ CRBN-induced neosubstrate degradation could confound mechanistic interpretation. Such possibly confounding effects are thus excluded through the design of VHL-based degraders^23,39^.

A noteworthy observation was the modest reduction of USP7 protein levels upon prolonged inhibitor treatment, highlighting an inherent autoregulatory mechanism. This complicates attempts to ascribe phenotypes solely to catalytic inhibition, particularly beyond 24 hours of compound treatment where enzymatic blockade and protein loss co-occur. Such effects will need to be taken into account when distinguishing catalytic from scaffolding functions, for which our rapid degraders can be employed in future studies.

The proteomics data further illuminated the consequences of USP7 perturbation through inhibition versus degradation. While early timepoints revealed selective depletion of known USP7 substrate proteins, prolonged inhibitor treatment triggered expansive proteomic remodeling, concomitant with metabolic rewiring. In contrast, degraders produced narrower, on-target responses with limited changes to the proteome—supporting their specificity and utility in defining bona fide USP7-dependent pathways. This distinction is particularly relevant when revisiting phenotypes attributed to legacy inhibitors such as P5091 or P22077, now known to potently engage other DUBs in cells like USP10 and USP47.^11,38^

Notably, comparative proteomics across PDAC and melanoma revealed substantial cell-type-specific differences in USP7 substrate regulation. These findings reinforce emerging work suggesting that DUB functions are shaped by cell context and cell state.^52^ The exploration of USP7 as a drug target must therefore consider these contextual nuances when evaluating compound efficacy or potential resistance mechanisms.

In phenotypic assays, both NK250 and NK266 slightly reduced basal respiration and oxygen consumption rates, supporting a role for USP7 in bioenergetics. However, hydroxypiperidine-based inhibitors such as NK192 and FT671 induced metabolic collapse, glucose dependency, and widespread proteomic disturbances in the absence of USP7 depletion. While some effects may result from PRC1.1 complex perturbation in a USP7-inhibited state^14,53^, the extent and intensity of these phenotypes strongly argue for off-target engagement. This highlights the importance of systematic chemical compound validation and the superior fidelity of degrader-based approaches, particularly in mechanistic studies with longer compound treatment times.

Taken together, our results support a strategy for comprehensively investigating the cellular roles of USP7. Selective inhibitors remain valuable for rapid interrogation of catalytic function and substrate identification. However, for prolonged exposure or functional dissection beyond catalysis, high-fidelity degraders provide superior specificity and mechanistic resolution. The complementary application of these tools, as demonstrated here, enables nuanced investigation of USP7’s catalytic and non-enzymatic roles while minimizing confounding artifacts.

In summary, we introduce a rigorously validated inhibitor–degrader pair for USP7 that supports mechanistically resolved and time-adapted experimental design. These tools will not only refine our understanding of USP7 in cancer biology but also exemplify the broader potential of degrader–inhibitor complementarity in unraveling the complexity of the ubiquitin system.

## Acknowledgments

We are grateful to all members of the Paschen, Grüner and Gersch labs for discussions, advice, and reagents. We thank Jenny Borman, Finn Pfeffermann, Eva Hahn, Elisa Krosta and Nancy Meyer as well as the NMR and HRMS facilities of TU Dortmund University headed by Bastian Grabe and Sebastian Zühlke for excellent technical assistance. We are grateful to G. Peters for the gift of a USP7 plasmid (Addgene # 46753). Fig. 1a, 4a and 4c were created with BioRender. This work was funded by Deutsche Forschungsgemeinschaft (DFG, German Research Foundation, Project-ID 424228829 – SFB1430, to F.K., M.K., N.S., S.K., A.P., B.M.G, M.G.). Work in the Gersch lab is further supported by AstraZeneca, Merck KGaA, Pfizer Inc., and the Max Planck Society as part of the Chemical Genomics Centre III (CGCIII-352S, to M.G.), and by the DFG through an Emmy Noether Award (GE 3110/1-1 to M.G.). The Grüner lab is further supported by the DFG with an Emmy Noether Award (GR4575/1-1, GR4575/1-2, to B.M.G), the Paschen lab received additional funding from the German Cancer Aid (DKH: 70115300). Work in the Wolf lab was supported by grants from the German Research Foundation (DFG: WO 2108/2-1, TRR387, GRK 3085), the German Cancer Aid (DKH: TACTIC) and the European Research Council (ERC: PROTAC-PDAC: #101087045) to E.W. Work in the Knapp lab was further supported by the Structural Genomics Consortium (SGC, a registered charity (no: 1097737) that receives funds from Bayer AG, Boehringer Ingelheim, Bristol Myers Squibb, Genome Canada), by the EU/EFPIA/OICR/McGill/KTH/Diamond Innovative Medicines Initiative 2 Joint Undertaking (EUbOPEN grant 875510), Janssen, Pfizer and Takeda, as well as by the LOEWE Center Frankfurt Cancer Institute (FCI) funded by the Hessian Ministry of Higher Education, Research and the Arts (III L5-519/03/03.001-(0015)).

## Author Contributions

N.K., S.U., J.A.S., A.P., B.M.G. and M.G. jointly designed the study, planned experiments and analyzed data. N.K. performed chemical synthesis and inhibition assays. S.U., M.D. and J.A.S. performed cellular experiments with PDAC and melanoma cells, respectively. B.A. with support by J.M. and under supervision by E.W. generated HiBiT-USP7 cells and recorded HiBiT data. M.P.S. under supervision by S.K. recorded ternary complex formation data. F.K. and M.K. performed mass spectrometry. S.F. performed the pulldown assay. J.K. and N.S. performed cell imaging. N.K., S.U., J.A.S., A.P., B.M.G. and M.G. together assembled the manuscript with input from all authors.

## Methods

### Chemicals and chemical synthesis

Synthetic procedures and analytic data for compounds prepared for this work are shown in the Supporting Information. Chemicals, which were not synthesized for this work, were purchased from MedChemExpress (FT671: #HY-107985; XL177A: #HY-138793; USP7-In-3: #HY-112128; P5091: #HY-15667; GNE-6640: #HY-112937; Carfilzomib (CFZ): #HY-10455 and Pevonedistat/MLN4924: #HY-70062). All compounds (inhibitors, PROTACs and controls) were dissolved in DMSO, which was used as vehicle control.

### Cell culture

The melanoma cell line Ma-Mel-47 (established previously^54^ from metastatic lesions from patient tumor tissues after written informed consent) and the PDAC cell line Panc89 (RRID: CVCL_4056, also referenced as T3M-4^55^) were cultured in a humidified incubator with 5% CO_2_ at 37 °C in RPMI1640 (Ma-Mel-47) / DMEM (Panc89) medium containing 10% (v/v) fetal calf serum (FCS) and penicillin/streptomycin (all Gibco, Thermo Fisher Scientific). Studies on human material were approved by the institutional review board. Cells were confirmed to be negative for mycoplasma contamination. Testing for mycoplasma was performed by PCR monthly. To confirm cell identity, STR analysis was performed by the Microsynth Seqlab GmbH.

### PROTAC rescue experiments

Cells were treated with proteasome inhibitor Carfilzomib (CFZ, 250 nM), NEDDylation Inhibitor MLN4924 (500 nM) or VHL inhibitor NK249 (10 µM). After 2 h, PROTAC (1 µM) was added. Cells were analyzed after 20 h.

### Preparation of whole cell extracts

To prepare samples for Western blot analysis, cells were seeded in 6-well dishes the day before treatment to adhere at 70-80% confluency. Compounds were added to the culture media at working concentrations of 1 nM–10 µM and incubated between 2 and 72 h.

Cell pellets of all PDAC samples were lysed via adding a buffer containing 150 mM NaCl, 50 mM Tris/HCl (pH 7.4), 1% Triton X-100, 10 mM EDTA, 1 mM PMSF, 10 mM orthovanadate, and 1% aprotinin, incubating on ice for 1 h with vortexing every 15 minutes. Lysates were cleared of cell debris by centrifugation at 10000 ×g at 4 °C for 10 min. Cell pellets of melanoma samples were lysed using Cell Lysis Buffer #9803 (Cell Signaling Technology) and incubating for 15 min on ice. Lysates were then clarified of cell debris by centrifugation at 13000 ×g at 4 °C for 15 min. Thereafter, the total protein concentrations were either determined immediately or the protein extracts were stored at -20 or -80 °C. Pellets for samples shown in Fig. 6 and Supporting Fig. 6 were lysed in 120 µL lysis buffer (50 mM Tris pH 8.0, 150 mM NaCl, 1% (v/v) IGEPAL, 0.5% (w/v) Na-deoxycholate, 0.1% (w/v) SDS) supplemented with 1x EDTA-free inhibitor protease cocktail (cOmplete, Roche) for 15 min at 4 °C. 0.2 µL Turbonuclease (Sigma Aldrich) and 5 mM MgCl_2_ were then added and lysates were incubated for 15 min at 4 °C. Cell debris was separated by centrifugation at 20817 ×g for 15 min at 4 °C.

Protein concentrations of cell lysates were measured using the Pierce BCA Protein Assay (Thermo Fisher Scientific) or the Quick Start Bradford Protein Assay (Bio-Rad), following the manufacturers’ protocols. Measurements were recorded on Tecan Spark or Tecan Infinite M200 plate readers. Lysates were kept on ice throughout the entire process.

### SDS-Page, transfer and staining

To separate proteins via SDS-PAGE, 10-20 µg of protein in lysis buffer was diluted with SDS-PAGE loading buffer (for most blots containing DTT or 2-mercaptoethanol) and denatured at 95°C for 5 min. Protein separation was conducted at 95-110 V using the Mini-PROTEAN electrophoresis system (Bio-Rad). Proteins were then transferred to PVDF or nitrocellulose membranes using the Trans-Blot Turbo system (Bio-Rad) with the Trans-Blot Turbo RTA Kit for PVDF membranes. Prior to transfer, membranes were activated by soaking in 100% ethanol (for PVDF) or distilled water (for nitrocellulose) and proteins were transferred using the Mixed MW or the standard (1.0 A, 25 V, 30 min) protocol.

After transfer, membranes were blocked in 5% (w/v) milk/TBS-T for 1 h and then washed three times for 10 min with TBS-T. They were incubated overnight at 4°C with the primary antibody in 5% (w/v) milk/TBS-T or 1% (w/v) BSA/TBS-T. Membranes were then washed three times for 10 min with TBS-T and incubated with horseradish peroxidase (HRP)-conjugated secondary antibodies in a solution of 5% (w/v) BSA/TBS-T or 1% (w/v) milk/TBS-T for 1 h at room temperature. PBS-T was used for blots shown in Fig. 6 and Supporting Fig. 6.

Membranes were incubated with Clarity Western ECL substrate (Bio-Rad) for PDAC samples or Pierce ECL Western Blotting-Substrate (Thermo Fisher Scientific) for melanoma. Chemiluminescent signals were detected with a ChemiDoc Imaging System (Bio-Rad) for PDAC samples or with a ECL Chemostar Imager (Intas Science Imaging) for melanoma samples.

Antibodies used were purchased from Cell Signaling Technology (USP7: HAUSP (D17C6) XP Rabbit mAb #4833, 1:1000; GAPDH: GAPDH (14C10) Rabbit mAb #2118, 1:1000 or 1:5000; secondary: Anti-Rabbit IgG, HRP-linked #7074, 1:10000). The following antibodies were used for data shown in Fig. 6 and Supporting Fig. 6: USP7, Abcam, ab190183, 1:2000; GAPDH, Thermo Fisher, AM4300, 1:10000; secondary anti-mouse HRP-linked, Sigma, NXA931, 1:10000; secondary anti-rabbit HRP-linked, Sigma, GENA934, 1:5000).

### Seahorse mito stress test and confluence measurements

The Seahorse XF cell mito stress test was used to measure oxygen consumption rates (OCR). Cells (1.5×10^4^ per well) were seeded into cell culture microplates (Agilent) in culture medium containing either 1, 5 or 10 µM FT671, NK192, NK250 (Panc89), NK266 (Ma-Mel-47) or vehicle control 24 h prior to start of the assay. For normalization, cells were simultaneously seeded into 96-well plates (1.2×10^4^ per well), treated with compounds, and cell confluence was measured on the day of the assay using the NYONE automated cell imager (Synentec). To reveal key parameters of mitochondrial functionality, Oligomycin (1.5 µM), Carbonyl cyanide-4-trifluoromethoxyphenylhydrazone (FCCP; 1 µM), Rotenone (0.5 µM) and Antimycin A (0.5 µM) were loaded into ports of the XF sensor cartridge. After washing of the cells and calibration of the XF sensor cartridge plate, OCR values were determined by an XF96/XFe96 Seahorse Analyzer. Data analysis was conducted using the Wave software (Agilent).

### Annexin V staining and medium color switch

Annexin V staining was used to assess apoptosis of Panc89 and Ma-Mel-47 cells. Cells were seeded in a 12-well format and pre-treated with 1, 5 or 10 µM FT671, NK192, NK250 (Panc89), NK266 (Ma-Mel-47) or vehicle control for 24 h in standard culture medium conditions. Cells were then treated with the same compounds in high (DMEM: 25 mM; RPMI: 11 mM) or low (5 mM) glucose culture medium. After 72 h of compound treatment, cells were either harvested, stained with FITC Annexin V (1:30; BD Biosciences 556419) and DAPI (1:1000), and subsequently measured on a BD FACSCelesta Cell Analyzer (BD Biosciences) or a photo was taken to visualize the medium color switch. Data analysis of FACS data was performed with the FlowJo Software.

### Cellular ternary complex formation assay

The assay was performed as described step-by-step elsewhere.^48^ USP7 (NanoLuc-USP7) and VHL (HaloTag-VHL) were cloned in frame with N-terminal NanoLuc or HaloTag, respectively. For transfections, HEK293T cells were diluted in Opti-MEM medium without phenol red (Life Technologies) to 4×10^5^ cells/mL and 9 µL were pipetted into each well of a white 384-well plate (Greiner 781207). 1 mL transfection mix was prepared containing 30 µL FuGENE HD (Promega, E2312) and 8 µL of each plasmid, incubated 20 min at RT and 1 µL were pipetted into each well leading to the total assay volume of 10 µL. After incubation at 37 °C and 5% CO_2_ for 20 h, 40 nL HaloTag NanoBRET 618 Ligand (Promega) was added to the cells using an Echo acoustic dispenser (Labcyte) and the cells were incubated for an additional 20 h at 37 °C and 5% CO_2_. 2 h prior to BRET measurements, the compounds were titrated to the cells using an Echo acoustic dispenser and the cells were incubated further at 37 °C and 5% CO_2_ to allow complex formation. NanoBRET Nano-Glo substrate and Extracellular NanoLuc Inhibitor (Promega, N2540) were then added as per the manufacturer’s protocol, and filtered luminescence was recorded on a PHERAstar FSX plate reader (BMG Labtech) equipped with a luminescence filter pair (donor: 450 nm BP filter; acceptor: 610 nm LP filter). Data were analyzed using GraphPad Prism 10 software.

### HiBiT USP7 degradation assay

#### Cell culture

Human MV4-11 and HEK293 cells were cultured at 37 °C in 5% CO_2_ in RPMI-1640 and DMEM medium, respectively, each supplemented with 10% FBS and 1% penicillin/streptomycin.

#### Cloning

HiBiT-USP7 construct was cloned by extraction of USP7 fragment from plasmid pQHA-USP7 WT puroR (Addgene #46753) using appropriate restriction sites. The USP7 fragment was then ligated into the pRRL-PGK-HiBiT entry vector to generate the plasmid.

#### Cell line generation

Lentiviral infection was used to generate stable a MV4-11 HiBiT cell line. Lentivirus was produced using plasmids psPAX2, pMD2.G, and HiBiT-USP7 plasmid in HEK293 cells. MV4-11 cells were infected with filtered virus supernatant and selected after 48 h of infection for the generation of the stable MV4-11^HiBiT-USP^^7^ cell line.

#### HiBiT assay

The assay was performed as described previously.^6^ Briefly, MV4-11^HiBiT-USP^^7^ cells were seeded and treated with compounds for the indicated time. Nano-Glo HiBiT Lytic Detection System (Promega) was used for the assay and luminescence was measured on a Tecan Spark microplate reader (Tecan). DC_50_ values were calculated using the sigmoidal dose-response (four parameters) equation in GraphPad Prism and are given as the concentration where 50% of the total signal was remaining. D_max_ values are given as the maximum degradation observed at any concentration.

### Immunofluorescence staining and microscopy

Panc89 cells were seeded on 96-well plates (89626, Ibidi) and treated with either DMSO or compound for the indicated time and concentration. Cells were then fixed with 4% PFA in PBS for 20 min, permeabilized with 0.1% Triton X-100 in PBS for 10 min and incubated in blocking buffer containing 2% BSA in PBS for 1 h. Three 10 min washes with PBS were performed between different steps. Cells were treated with primary antibody anti-USP7 (ab190183, Abcam, rabbit, 1:500) in blocking solution for 1 h at RT. After three washes with PBS, the fluorescently labeled secondary antibody (goat anti-rabbit IgG labeled with Alexa Fluor Plus 488, A32731, Invitrogen, 1:1000) in blocking buffer was incubated together with rhodamine-phalloidin (R415, Invitrogen, 1:1000) and HCS CellMask deep red (H32721, Invitrogen, 1:5000) for 1 h at room temperature. Finally, cells were stained with DAPI (Invitrogen, 1 µg/ml) in PBS for 10 min and mounted with aqueous mounting medium (50001, Ibidi) for microscopy. Plates were imaged on a fully automated inverted EVOS M7000 microscope system (Invitrogen) with X-Apo 20x air objective (0.8 NA, Olympus, AMEP4906) using DAPI (AMEP4950), GFP (AMEP4951), RFP (AMEP4952), and Cy5 (AMEP4956) LED light cubes. Images were captured using a high-sensitivity 3.2 MP monochrome CMOS camera and were processed with Omero (v. web 5.28.0).

### USP7 inhibition assay

USP7 inhibitors in DMSO were diluted in UbR buffer (40 mM Tris-HCl pH 8, 150 mM NaCl, 1 mM TCEP, 0.1 mg/mL BSA, 0.01% Tween20) to a concentration of 80 μM (4x). This stock was then further diluted in an 11-times serial dilution in UbR buffer containing 4% DMSO (4x). To this was added the same volume of human USP7 full length protein^56^ (1.2 nM in UbR buffer, 4x). The resulting mixture was incubated for 15 min at rt after which 10 µL were added to a black 384 well low volume non-binding surface plate (Greiner 784900) in triplicates. The assay was initiated upon addition of 10 μL per well of Ubiquitin-Rhodamine110Gly (Ub-RhoG) (100 nM in UbR buffer, 2x). A TCEP-free UbR buffer was used for assaying P5091. Normalization was carried out using samples featuring either no inhibitor or no enzyme as respective controls. Fluorescence was measured on a Tecan Spark plate reader for 1 h in 2 min intervals at 25 °C (excitation = 492 nm, emission = 525 nm). Slopes of fluorescence over time curves were determined with Microsoft Excel through linear regression using the first 15 min of each measurement. Biochemical IC_50_ values were calculated in GraphPad Prism 10.4.1 using non-linear regression (4 parameters dose response).

### Competitive pulldown assay

Panc89 cells were lysed in 5 mL of chemoproteomics lysis buffer (50 mM Tris pH 7.5, 150 mM NaCl, 1% NP-40 Alternative, 5% Glycerol, 2 mM TCEP) supplemented with 2 µL Benzonase on ice for 2 h. Lysate was cleared by centrifugation for 15 min at 17 000 ×g, supernatant was transferred into a fresh tube and kept on ice. Protein concentration was determined using a DC-Assay kit (Bio-Rad). In parallel, streptavidin magnetic beads (30 µL per sample, Pierce #88817) were saturated with 1.1 equiv. of NK264 for 1 h at 4°C. Afterwards beads were washed 3 times using chemoproteomics washing buffer (10 mM Tris pH 7.5, 150 mM NaCl, 0.05% Tween20) and equally divided tubes. Afterwards, Panc89 lysate (500 µL containing 700 µg of protein) containing either DMSO or NK192 (10 µM) were added onto the beads, and the mixture was inverted overnight at 4°C. Next, the mixture was briefly centrifuged to gather the solution, beads were pelleted using a magnet and the supernatant was discarded. Beads were washed two times with chemoproteomics washing buffer and once with PBS. After each washing step, beads were pelleted using a magnet and the supernatant was discarded. Finally, bound proteins were eluted off the beads by addition of 50 µL of 2x SP3 buffer (2% SDS, 20 mM TCEP, 80 mM CAM, 100 mM Hepes pH 8.0) followed by heating to 90°C for 5 min with mixing. Afterwards, beads were separated using a magnet and the supernatant was transferred into a fresh tube. This elution step was repeated by adding 50 µL of 1x SP3 buffer (1% SDS, 10 mM TCEP, 40 mM CAM, 50 mM Hepes pH 8.0) onto the beads and heating at 90°C for 5 min. First and second eluates were united and frozen at -80°C until further processing.

### Sample preparation for enriched proteome samples

3 µL per sample of a 50 µg/µL 1:1 mixture of hydrophilic (#45152105050250) and hydrophobic (#65152105050250) carboxylate modified Sera-Mag SpeedBeads (Cytiva), that were washed twice with MS-grade water, were added. The next steps were carried out at room temperature unless noted otherwise. Afterwards, the samples were mixed shortly (1 min, 1000 rpm) and collected by short centrifugation (10 s, 200 ×g). Protein binding was induced by the addition of an equal volume of pure ethanol (10 min, 1000 rpm), the beads were collected by a brief centrifugation step (10 s, 200 ×g) and the plate was placed on a magnetic stand. Beads were allowed to bind for at least 5 min before the supernatant was removed. The beads were then taken up in 180 µL 80% ethanol and transferred to a fresh multiwell plate.

Subsequently the beads were washed four times with 180 µL 80% (v/v) ethanol prior to the addition of 100 µL digestion enzyme mix (0.6 μg of trypsin (V5111; Promega) and 0.6 µg LysC (125-05061; FUJIFILM Wako Pure Chemical) in 25 mM ammonium bicarbonate). Samples were incubated at 37 °C for 19 h while shaking (1300 rpm). On the next day, the samples were briefly centrifuged (10 s, 200 ×g) and placed on a magnet for 5 min. The clear solution containing the tryptic peptides was transferred to a fresh multiwell plate. The beads were taken up in 47 µL 25 mM ammonium bicarbonate and incubated while shaking (10 min, 1000 rpm). The plate was then placed on a magnetic stand and after 5 min the cleared supernatant was collected and combined with the recovered first peptide mix, followed by the addition of formic acid (FA) to a final concentration of 2% (v/v) for trypsin inactivation.

### Sample preparation for total proteome analysis

Panc89 and Ma-Mel cells were treated in quadruplicate with 5 µM NK192, 1 µM NK250 (PDAC), 1 µM NK266 (Melanoma), or DMSO for 6 h, 24 h or 72 h. Following treatment, cells were harvested, washed twice with PBS, and pellets were flash frozen in liquid nitrogen before being stored at -80 °C. Samples were prepared using the single-pot, solid-phase-enhanced sample-preparation (SP3) strategy.^57^ All buffers and solutions were prepared with mass spectrometry (MS)-grade water (Honeywell). Cell pellets were taken up in 100 µL 1x sample buffer (50 mM Hepes pH 8.0, 1% (wt/v) SDS, 1% (v/v) NP-40, 10 mM TCEP, 40 mM chloroacetamide) and samples were heated at 90°C for 5 min prior to sonication with a Bioruptor UCD-300 (Diagenode) for ten cycles of 1 min pulse and 30 s pause at high power. The samples were then supplemented with Benzonase (20 U/sample) and sonified for 5 min in a water bath (Bandelin) followed by 30 min incubation at 30 min.

The protein extracts were then centrifuged (20000 ×g, RT, 40 min) and the protein concentration of the cleared lysates was determined using the Pierce 660 nm Protein Assay Reagent (#22660; Thermo Scientific) with the Ionic Detergent Compatibility Reagent (#22663; Thermo Scientific) according to the manufacturers’ instructions. Next, 15 µg of total protein per sample in a volume of 100 µL sample buffer was transferred to a 500 µL 96-well plate and treated with 7 U of benzonase (#71206; Merck Millipore) in dilution buffer (20 mM Hepes pH 8.0, 2 mM MgCl_2_) at 37 °C for 30 min with agitation at 1500 rpm. After Benzonase treatment samples were heated to 90°C for 5 min and after cooling down to room temperature, iodoacetamide to a final concentration of 10 mM was added for complete alkylation of cysteines (37 °C, 15 min, 1000 rpm, in the dark).

Then, 3 µL of a 50 µg/µL 1:1 mixture of hydrophilic (#45152105050250) and hydrophobic (#65152105050250) carboxylate modified Sera-Mag SpeedBeads (Cytiva), that were washed twice with MS-grade water, were added to the samples. The next steps were carried out at room temperature unless noted otherwise. Afterwards, the samples were mixed shortly (1 min, 1000 rpm) and collected by short centrifugation (10 s, 200 ×g). Protein binding was induced by the addition of an equal volume of pure ethanol (10 min, 1000 rpm), the beads were collected by a brief centrifugation step (10 s, 200 ×g) and the plate was placed on a magnetic stand. Beads were allowed to bind for at least 5 min before the supernatant was removed. The beads were then taken up in 180 µL 80% ethanol and transferred to a fresh multiwell plate.

Subsequently the beads were washed four times with 180 µL 80% (v/v) ethanol prior to the addition of 100 µL digestion enzyme mix (0.6 μg of trypsin (V5111; Promega) and 0.6 µg LysC (125-05061; FUJIFILM Wako Pure Chemical) in 25 mM ammonium bicarbonate). Samples were incubated at 37 °C for 19 h while shaking (1300 rpm). On the next day, the samples were briefly centrifuged (10 s, 200 ×g) and placed on a magnet for 5 min. The clear solution containing the tryptic peptides was transferred to a fresh multiwell plate. The beads were taken up in 47 µL 25 mM ammonium bicarbonate and incubated while shaking (10 min, 1000 rpm). The plate was then placed on a magnetic stand and after 5 min the cleared supernatant was collected and combined with the recovered first peptide mix, followed by the addition of formic acid (FA) to a final concentration of 2% (v/v) for trypsin inactivation.

### Sample clean-up for LC-MS/MS

All digests were desalted on home-made C18 StageTips^58^ containing two layers of an octadecyl silica membrane (CDS Analytical). All centrifugation steps were carried out at room temperature. The StageTips were first activated and equilibrated by passing 50 μL of methanol (600 ×g, 2 min), 80% (v/v) acetonitrile (ACN) with 0.5% (v/v) FA (600 ×g, 2 min) and 0.5% (v/v) FA (600 ×g, 2 min) over the tips. Next, the acidified tryptic digests were passed over the tips (800 ×g, 3 min). The immobilized peptides were then washed with 50 μL and 25 μL 0.5% (v/v) FA (800 ×g, 3 min). Bound peptides were eluted from the StageTips by application of two rounds of 25 μL 80% (v/v) ACN with 0.5% (v/v) FA (800 ×g, 2 min). Peptide samples were then dried using a vacuum concentrator (Eppendorf) and the peptides were dissolved in 15 μL 0.1% (v/v) FA prior to analysis by MS.

### Mass spectrometry

LC-MS/MS analysis of peptide samples was performed on an Orbitrap Fusion Lumos mass spectrometer (Thermo Scientific) coupled to a Vanquish Neo ultra high-performance liquid chromatography (UHPLC) system (Thermo Scientific) that was operated in the one-column mode. The analytical column was a fused silica capillary (inner diameter: 75 μm, outer diameter: 360 µm, length: 28 cm; CoAnn Technologies) with an integrated sintered frit packed in-house with Kinetex 1.7 μm XB-C18 core shell material (Phenomenex). The analytical column was encased by a PRSO-V2 column oven (Sonation) and attached to a nanospray flex ion source (Thermo Scientific). The column oven temperature was set to 50 °C during sample loading and data acquisition. The LC was equipped with two mobile phases: solvent A (2% ACN and 0.2% FA, in water) and solvent B (80% ACN and 0.2% FA, in water). All solvents were of UHPLC grade (Honeywell). Peptides were directly loaded onto the analytical column with a maximum flow rate that would not exceed the set pressure limit of 950 bar (usually around 0.5 – 0.6 μL/min) and separated on the analytical column by running a 105 min gradient of solvent A and solvent B at a flow rate of 300 nL/min (start with 3% (v/v) B, gradient 3% to 6% (v/v) B for 5 min, gradient 6% to 29% (v/v) B for 70 min, gradient 29% to 42% (v/v) B for 15 min, gradient 42% to 100% (v/v) B for 5 min and 100% (v/v) B for 10 min).

The mass spectrometer was controlled by the Orbitrap Fusion Lumos Tune Application (version 4.1.4244) and operated using the Xcalibur software (version 4.7.69.37). The MS settings are provided in Supporting Table 1 (for measurements in data-dependent acquisition (DDA) mode; Fig. 1d, project: ACE_ ACE_0869; Supporting Fig. 4a-f, project: ACE_0882-DDA) and Supporting Table 2 (for measurements in data-independent acquisition (DIA) mode; Fig. 5, projects: ACE_0882-DIA and ACE_0883-DIA).

### Data processing and analysis of DIA mass spectrometry experiments

Recorded RAW data files were converted to the mzML file format using ProteoWizard^59^ (“peak Picking” (vendor MS level 1) as first filter, TPP compatibility, 64bit encoding precision, and index writing switched on) and analyzed with DIA-NN^60^ (version 1.8.1). DIA-NN was used in the library free mode. Spectral library generation is based on the Uniprot *H. sapiens* reference proteome (UP000005640_9606_OPPG; one protein per gene; 20606 entries, downloaded July 2023). For the search we selected “Trypsin/P” as protease with 2 missed cleavages allowed. As variable modifications we set “N-ter M excision”, “C carbamidomethylation”, “Ox(M)” and “Ac(N-term)”. The maximum number of variable modifications was set to 2. Peptide length was set from 7 – 30; Precursor charge range from 2 – 6. Precursor m/z range was set to 396 – 1004 and Fragment ion m/z range was set to 150 – 1800. Mass accuracy, MS1 accuracy and Scan window was kept at “0” (automatic). Unrelated runs, use isotopologues, MBR and no shared spectra were selected. Heuristic protein inference was based on “Genes”. Neural network classifier was set to “Single-pass mode”. Quantification strategy was set to “Robust LC (high precision) with cross-run normalization set to “RT & signal-dep.”. Library generation was set to “smart profiling” and Speed and RAM usage was set to “optimal results”. Precursor FDR was set to 1%.

For further data analysis and filtering of the DIA-NN output, the report.pg_matrix.tsv file which contains the normalized (MaxLFQ)^61^ protein group intensities was loaded into Perseus^62^ (version 1.6.10.0). The data was transformed to the log_2_(x) scale and biological replicates were combined into categorical groups to allow comparison of the different treatment groups. For the full proteome analysis, only protein groups (PGs) with a valid LFQ intensity in at least two out of four replicates in each categorical group were kept for further analysis. The log_2_-fold change in normalized protein group quantities between the different categorical groups was determined based on the mean LFQ intensities of replicate samples (relative quantification). To enable quantification, missing LFQ intensities were imputed from a normal distribution (width 0.3, down shift 1.8). The statistical significance of the difference in LFQ intensity was determined via a two-sided Student’s t-test. To compare and visualize significantly up- and down-regulated (–Log(*P*-value) >2 and Log_2_ fold change >1) proteins between both cancer entities and compound treatments of the mass spectrometry analysis, area-proportional Venn diagrams were used (https://biovenn.nl/).

### Data processing and analysis of DDA mass spectrometry experiments

RAW spectra were submitted to an Andromeda^63^ search in MaxQuant (version 2.5.2.0) using the default settings^64^. Label-free quantification and match-between-runs was activated.^61^ The MS/MS spectra data were searched against the Uniprot *H. sapiens* reference proteome (see above). All searches included a contaminants database search as implemented in MaxQuant with 245 entries. Andromeda searches allowed oxidation of methionine residues (16 Da) and acetylation of the protein N-terminus (42 Da). Carbamidomethylation on Cystein (57 Da) was selected as static modification. Enzyme specificity was set to “Trypsin/P”. The instrument type in Andromeda searches was set to Orbitrap and the precursor mass tolerance was set to ±20 ppm (first search) and ±4.5 ppm (main search). The MS/MS match tolerance was set to ±0.5 Da. The peptide spectrum match FDR and the protein FDR were set to 0.01 (based on target-decoy approach). For protein quantification unique and razor peptides were allowed. Modified peptides were allowed for quantification. The minimum score for modified peptides was 40. Label-free protein quantification was switched on, and unique and razor peptides were considered for quantification with a minimum ratio count of 2. Retention times were recalibrated based on the built-in nonlinear time-rescaling algorithm. MS/MS identifications were transferred between LC-MS/MS runs with the “match between runs” option in which the maximal match time window was set to 0.7 min and the alignment time window set to 20 min. The quantification is based on the “value at maximum” of the extracted ion current. At least two quantitation events were required for a quantifiable protein. Further analysis and filtering of the results was done in Perseus^62^ v1.6.10.0 as described above. Comparison of protein group quantities (relative quantification) between different runs is based solely on LFQ values as calculated by the MaxQuant MaxLFQ algorithm^61^.

## Data availability statement

Mass spectrometry raw data were deposited to the ProteomeXchange Consortium through the PRIDE partner repository under accession codes PXD063889, PXD063917, PXD063920 and PXD063913. Processed mass spectrometry data are enclosed as Supporting Table 3 (Fig. 1d), Supporting Table 4 (Fig. 5a-f), Supporting Table 5 (Fig. 5g-l) and Supporting Table 6 (Supporting Fig. 4a-f). Uncropped blots and chemical compound characterization data are provided in the Supporting Information.

## References

1. Hinterndorfer, M., Spiteri, V.A., Ciulli, A. & Winter, G.E. Targeted protein degradation for cancer therapy. Nat Rev Cancer (2025).

2. Bondeson, D.P. & Crews, C.M. Targeted Protein Degradation by Small Molecules. Annu Rev Pharmacol Toxicol 57, 107–123 (2017).

3. Zengerle, M., Chan, K.H. & Ciulli, A. Selective Small Molecule Induced Degradation of the BET Bromodomain Protein BRD4. ACS Chem Biol 10, 1770–7 (2015).

4. Lu, J., Qian, Y., Altieri, M., Dong, H., Wang, J., Raina, K., Hines, J., Winkler, J.D., Crew, A.P., Coleman, K. & Crews, C.M. Hijacking the E3 Ubiquitin Ligase Cereblon to Efficiently Target BRD4. Chem Biol 22, 755–63 (2015).

5. Burslem, G.M., Smith, B.E., Lai, A.C., Jaime-Figueroa, S., McQuaid, D.C., Bondeson, D.P., Toure, M., Dong, H., Qian, Y., Wang, J., Crew, A.P., Hines, J. & Crews, C.M. The Advantages of Targeted Protein Degradation Over Inhibition: An RTK Case Study. Cell Chem Biol 25, 67–77 e3 (2018).

6. Adhikari, B., Bozilovic, J., Diebold, M., Schwarz, J.D., Hofstetter, J., Schroder, M., Wanior, M., Narain, A., Vogt, M., Dudvarski Stankovic, N., Baluapuri, A., Schonemann, L., Eing, L., Bhandare, P., Kuster, B., Schlosser, A., Heinzlmeir, S., Sotriffer, C., Knapp, S. & Wolf, E. PROTAC-mediated degradation reveals a non-catalytic function of AURORA-A kinase. Nat Chem Biol 16, 1179–1188 (2020).

7. Wang, R., Ascanelli, C., Abdelbaki, A., Fung, A., Rasmusson, T., Michaelides, I., Roberts, K. & Lindon, C. Selective targeting of non-centrosomal AURKA functions through use of a targeted protein degradation tool. Commun Biol 4, 640 (2021).

8. Clague, M.J., Urbe, S. & Komander, D. Breaking the chains: deubiquitylating enzyme specificity begets function. Nat Rev Mol Cell Biol 20, 338–352 (2019).

9. Lange, S.M., Armstrong, L.A. & Kulathu, Y. Deubiquitinases: From mechanisms to their inhibition by small molecules. Mol Cell 82, 15–29 (2022).

10. Wertz, I.E. & Wang, X. From Discovery to Bedside: Targeting the Ubiquitin System. Cell Chem Biol 26, 156–177 (2019).

11. Ritorto, M.S., Ewan, R., Perez-Oliva, A.B., Knebel, A., Buhrlage, S.J., Wightman, M., Kelly, S.M., Wood, N.T., Virdee, S., Gray, N.S., Morrice, N.A., Alessi, D.R. & Trost, M. Screening of DUB activity and specificity by MALDI-TOF mass spectrometry. Nat Commun 5, 4763 (2014).

12. Pozhidaeva, A. & Bezsonova, I. USP7: Structure, substrate specificity, and inhibition. DNA Repair (Amst) 76, 30–39 (2019).

13. Sheng, Y., Saridakis, V., Sarkari, F., Duan, S., Wu, T., Arrowsmith, C.H. & Frappier, L. Molecular recognition of p53 and MDM2 by USP7/HAUSP. Nat Struct Mol Biol 13, 285–91 (2006).

14. Sijm, A., Atlasi, Y., van der Knaap, J.A., Wolf van der Meer, J., Chalkley, G.E., Bezstarosti, K., Dekkers, D.H.W., Doff, W.A.S., Ozgur, Z., van, I.W.F.J., Demmers, J.A.A. & Verrijzer, C.P. USP7 regulates the ncPRC1 Polycomb axis to stimulate genomic H2AK119ub1 deposition uncoupled from H3K27me3. Sci Adv 8, eabq7598 (2022).

15. Liu, X., Yang, X., Li, Y., Zhao, S., Li, C., Ma, P. & Mao, B. Trip12 is an E3 ubiquitin ligase for USP7/HAUSP involved in the DNA damage response. FEBS Lett 590, 4213–4222 (2016).

16. Zhang, X.W., Feng, N., Liu, Y.C., Guo, Q., Wang, J.K., Bai, Y.Z., Ye, X.M., Yang, Z., Yang, H., Liu, Y., Yang, M.M., Wang, Y.H., Shi, X.M., Liu, D., Tu, P.F. & Zeng, K.W. Neuroinflammation inhibition by small-molecule targeting USP7 noncatalytic domain for neurodegenerative disease therapy. Sci Adv 8, eabo0789 (2022).

17. Gavory, G., O’Dowd, C.R., Helm, M.D., Flasz, J., Arkoudis, E., Dossang, A., Hughes, C., Cassidy, E., McClelland, K., Odrzywol, E., Page, N., Barker, O., Miel, H. & Harrison, T. Discovery and characterization of highly potent and selective allosteric USP7 inhibitors. Nat Chem Biol 14, 118–125 (2018).

18. Kategaya, L., Di Lello, P., Rouge, L., Pastor, R., Clark, K.R., Drummond, J., Kleinheinz, T., Lin, E., Upton, J.P., Prakash, S., Heideker, J., McCleland, M., Ritorto, M.S., Alessi, D.R., Trost, M., Bainbridge, T.W., Kwok, M.C.M., Ma, T.P., Stiffler, Z., Brasher, B., Tang, Y., Jaishankar, P., Hearn, B.R., Renslo, A.R., Arkin, M.R., Cohen, F., Yu, K., Peale, F., Gnad, F., Chang, M.T., Klijn, C., Blackwood, E., Martin, S.E., Forrest, W.F., Ernst, J.A., Ndubaku, C., Wang, X., Beresini, M.H., Tsui, V., Schwerdtfeger, C., Blake, R.A., Murray, J., Maurer, T. & Wertz, I.E. USP7 small-molecule inhibitors interfere with ubiquitin binding. Nature 550, 534–538 (2017).

19. Turnbull, A.P., Ioannidis, S., Krajewski, W.W., Pinto-Fernandez, A., Heride, C., Martin, A.C.L., Tonkin, L.M., Townsend, E.C., Buker, S.M., Lancia, D.R., Caravella, J.A., Toms, A.V., Charlton, T.M., Lahdenranta, J., Wilker, E., Follows, B.C., Evans, N.J., Stead, L., Alli, C., Zarayskiy, V.V., Talbot, A.C., Buckmelter, A.J., Wang, M., McKinnon, C.L., Saab, F., McGouran, J.F., Century, H., Gersch, M., Pittman, M.S., Marshall, C.G., Raynham, T.M., Simcox, M., Stewart, L.M.D., McLoughlin, S.B., Escobedo, J.A., Bair, K.W., Dinsmore, C.J., Hammonds, T.R., Kim, S., Urbe, S., Clague, M.J., Kessler, B.M. & Komander, D. Molecular basis of USP7 inhibition by selective small-molecule inhibitors. Nature 550, 481–486 (2017).

20. Leger, P.R., Hu, D.X., Biannic, B., Bui, M., Han, X., Karbarz, E., Maung, J., Okano, A., Osipov, M., Shibuya, G.M., Young, K., Higgs, C., Abraham, B., Bradford, D., Cho, C., Colas, C., Jacobson, S., Ohol, Y.M., Pookot, D., Rana, P., Sanchez, J., Shah, N., Sun, M., Wong, S., Brockstedt, D.G., Kassner, P.D., Schwarz, J.B. & Wustrow, D.J. Discovery of Potent, Selective, and Orally Bioavailable Inhibitors of USP7 with In Vivo Antitumor Activity. J Med Chem 63, 5398–5420 (2020).

21. Tong, J., Watkins, J.M., Burke, J.M. & Kodadek, T. Assessing the Suitability of Deubiquitylases As Substrates For Targeted Protein Degradation. bioRxiv, 2025.06.13.659525 (2025).

22. Pei, Y., Fu, J., Shi, Y., Zhang, M., Luo, G., Luo, X., Song, N., Mi, T., Yang, Y., Li, J., Zhou, Y. & Zhou, B. Discovery of a Potent and Selective Degrader for USP7. Angew Chem Int Ed Engl 61, e202204395 (2022).

23. Xu, M., Fu, J., Pei, Y., Li, M., Kan, W., Yan, R., Xia, C., Ma, J., Wang, P., Zhang, Y., Gao, Y., Yang, Y., Zhou, Y., Li, J. & Zhou, B. Discovery of a Highly Potent, Selective and Efficacious USP7 Degrader for the Treatment of Acute Lymphoblastic Leukemia. J Med Chem 67, 13197–13216 (2024).

24. Murgai, A., Sosic, I., Gobec, M., Lemnitzer, P., Proj, M., Wittenburg, S., Voget, R., Gutschow, M., Kronke, J. & Steinebach, C. Targeting the deubiquitinase USP7 for degradation with PROTACs. Chem Commun (Camb) 58, 8858–8861 (2022).

25. Reinders, M., Kravic, B., Gahlot, P., van den Boom, J., Schulze, N., Levantovsky, S., Kleine, S., Kaiser, M., Kulathu, Y., Behrends, C. & Meyer, H. ATXN3 regulates lysosome regeneration after damage by targeting K48-K63-branched ubiquitin chains. bioRxiv, 2025.06.10.656592 (2025).

26. Yi, J., Li, H., Chu, B., Kon, N., Hu, X., Hu, J., Xiong, Y., Kaniskan, H.U., Jin, J. & Gu, W. Inhibition of USP7 induces p53-independent tumor growth suppression in triple-negative breast cancers by destabilizing FOXM1. Cell Death Differ 30, 1799–1810 (2023).

27. Bushman, J.W., Donovan, K.A., Schauer, N.J., Liu, X., Hu, W., Varca, A.C., Buhrlage, S.J. & Fischer, E.S. Proteomics-Based Identification of DUB Substrates Using Selective Inhibitors. Cell Chem Biol 28, 78–87 e3 (2021).

28. Steger, M., Demichev, V., Backman, M., Ohmayer, U., Ihmor, P., Muller, S., Ralser, M. & Daub, H. Time-resolved in vivo ubiquitinome profiling by DIA-MS reveals USP7 targets on a proteome-wide scale. Nat Commun 12, 5399 (2021).

29. Schauer, N.J., Liu, X., Magin, R.S., Doherty, L.M., Chan, W.C., Ficarro, S.B., Hu, W., Roberts, R.M., Iacob, R.E., Stolte, B., Giacomelli, A.O., Perera, S., McKay, K., Boswell, S.A., Weisberg, E.L., Ray, A., Chauhan, D., Dhe-Paganon, S., Anderson, K.C., Griffin, J.D., Li, J., Hahn, W.C., Sorger, P.K., Engen, J.R., Stegmaier, K., Marto, J.A. & Buhrlage, S.J. Selective USP7 inhibition elicits cancer cell killing through a p53-dependent mechanism. Sci Rep 10, 5324 (2020).

30. Campos Alonso, M. & Knobeloch, K.P. In the moonlight: non-catalytic functions of ubiquitin and ubiquitin-like proteases. Front Mol Biosci 11, 1349509 (2024).

31. Gu, J., Xiao, X., Zou, C., Mao, Y., Jin, C., Fu, D., Li, R. & Li, H. Ubiquitin-specific protease 7 maintains c-Myc stability to support pancreatic cancer glycolysis and tumor growth. J Transl Med 22, 1135 (2024).

32. Chen, H., Zhu, X., Sun, R., Ma, P., Zhang, E., Wang, Z., Fan, Y., Zhou, G. & Mao, R. Ubiquitin-specific protease 7 is a druggable target that is essential for pancreatic cancer growth and chemoresistance. Invest New Drugs 38, 1707–1716 (2020).

33. Desmond, M.E. & Schoenwolf, G.C. Evaluation of the roles of intrinsic and extrinsic factors in occlusion of the spinal neurocoel during rapid brain enlargement in the chick embryo. J Embryol Exp Morphol 97, 25–46 (1986).

34. Xiang, M., Liang, L., Kuang, X., Xie, Z., Liu, J., Zhao, S., Su, J., Chen, X. & Liu, H. Pharmacological inhibition of USP7 suppresses growth and metastasis of melanoma cells in vitro and in vivo. J Cell Mol Med 25, 9228–9240 (2021).

35. Granieri, L., Marocchi, F., Melixetian, M., Mohammadi, N., Nicoli, P., Cuomo, A., Bonaldi, T., Confalonieri, S., Pisati, F., Giardina, G., Bertalot, G., Bossi, D. & Lanfrancone, L. Targeting the USP7/RRM2 axis drives senescence and sensitizes melanoma cells to HDAC/LSD1 inhibitors. Cell Rep 40, 111396 (2022).

36. Gao, L., Zhu, D., Wang, Q., Bao, Z., Yin, S., Qiang, H., Wieland, H., Zhang, J., Teichmann, A. & Jia, J. Proteome Analysis of USP7 Substrates Revealed Its Role in Melanoma Through PI3K/Akt/FOXO and AMPK Pathways. Front Oncol 11, 650165 (2021).

37. Su, L., Wang, D., Purwin, T.J., Ran, S., Yang, Q., Zhang, Q. & Cai, W. Selective USP7 Inhibition Synergizes with MEK1/2 Inhibitor to Enhance Immune Responses and Potentiate Anti-PD-1 Therapy in NRAS-Mutant Melanoma. J Invest Dermatol (2025).

38. Weisberg, E.L., Schauer, N.J., Yang, J., Lamberto, I., Doherty, L., Bhatt, S., Nonami, A., Meng, C., Letai, A., Wright, R., Tiv, H., Gokhale, P.C., Ritorto, M.S., De Cesare, V., Trost, M., Christodoulou, A., Christie, A., Weinstock, D.M., Adamia, S., Stone, R., Chauhan, D., Anderson, K.C., Seo, H.S., Dhe-Paganon, S., Sattler, M., Gray, N.S., Griffin, J.D. & Buhrlage, S.J. Inhibition of USP10 induces degradation of oncogenic FLT3. Nat Chem Biol 13, 1207–1215 (2017).

39. Wittenburg, S., Zuleeg, M.R., Peter, K., Lemnitzer, P., Voget, R., Bricelj, A., Gobec, M., Dierlamm, N., Braun, M.B., Geiger, T.M., Heim, C., Stakemeier, A., Wagner, K.G., Nowak, R.P., Hartmann, M.D., Sosič, I., Gütschow, M., Krönke, J. & Steinebach, C. *ChemRxiv*, chemrxiv-2025-q39r1 (2025).

40. Chauhan, D., Tian, Z., Nicholson, B., Kumar, K.G., Zhou, B., Carrasco, R., McDermott, J.L., Leach, C.A., Fulcinniti, M., Kodrasov, M.P., Weinstock, J., Kingsbury, W.D., Hideshima, T., Shah, P.K., Minvielle, S., Altun, M., Kessler, B.M., Orlowski, R., Richardson, P., Munshi, N. & Anderson, K.C. A small molecule inhibitor of ubiquitin-specific protease-7 induces apoptosis in multiple myeloma cells and overcomes bortezomib resistance. Cancer Cell 22, 345–58 (2012).

41. Diehl, C.J. & Ciulli, A. Discovery of small molecule ligands for the von Hippel-Lindau (VHL) E3 ligase and their use as inhibitors and PROTAC degraders. Chem Soc Rev 51, 8216–8257 (2022).

42. Frost, J., Galdeano, C., Soares, P., Gadd, M.S., Grzes, K.M., Ellis, L., Epemolu, O., Shimamura, S., Bantscheff, M., Grandi, P., Read, K.D., Cantrell, D.A., Rocha, S. & Ciulli, A. Potent and selective chemical probe of hypoxic signalling downstream of HIF-alpha hydroxylation via VHL inhibition. Nat Commun 7, 13312 (2016).

43. Han, X., Wang, C., Qin, C., Xiang, W., Fernandez-Salas, E., Yang, C.Y., Wang, M., Zhao, L., Xu, T., Chinnaswamy, K., Delproposto, J., Stuckey, J. & Wang, S. Discovery of ARD-69 as a Highly Potent Proteolysis Targeting Chimera (PROTAC) Degrader of Androgen Receptor (AR) for the Treatment of Prostate Cancer. J Med Chem 62, 941–964 (2019).

44. Urbe, S., Liu, H., Hayes, S.D., Heride, C., Rigden, D.J. & Clague, M.J. Systematic survey of deubiquitinase localization identifies USP21 as a regulator of centrosome- and microtubule-associated functions. Mol Biol Cell 23, 1095–103 (2012).

45. Schwinn, M.K., Machleidt, T., Zimmerman, K., Eggers, C.T., Dixon, A.S., Hurst, R., Hall, M.P., Encell, L.P., Binkowski, B.F. & Wood, K.V. CRISPR-Mediated Tagging of Endogenous Proteins with a Luminescent Peptide. ACS Chem Biol 13, 467–474 (2018).

46. Fuhrer, S., Gallant, K., Kaschani, F., Kaiser, M., Janning, P., Waldmann, H. & Gersch, M. Small Molecule-Induced Alterations of Protein Polyubiquitination Revealed by Mass-Spectrometric Ubiquitome Analysis. Angew Chem Int Ed Engl, e202508916 (2025).

47. Roy, M.J., Winkler, S., Hughes, S.J., Whitworth, C., Galant, M., Farnaby, W., Rumpel, K. & Ciulli, A. SPR-Measured Dissociation Kinetics of PROTAC Ternary Complexes Influence Target Degradation Rate. ACS Chem Biol 14, 361–368 (2019).

48. Schwalm, M.P., Saxena, K., Muller, S. & Knapp, S. Luciferase- and HaloTag-based reporter assays to measure small-molecule-induced degradation pathway in living cells. Nat Protoc 19, 2317–2357 (2024).

49. Frohlich, K., Fahrner, M., Brombacher, E., Seredynska, A., Maldacker, M., Kreutz, C., Schmidt, A. & Schilling, O. Data-Independent Acquisition: A Milestone and Prospect in Clinical Mass Spectrometry-Based Proteomics. Mol Cell Proteomics 23, 100800 (2024).

50. Nie, L., Wang, C., Liu, X., Teng, H., Li, S., Huang, M., Feng, X., Pei, G., Hang, Q., Zhao, Z., Gan, B., Ma, L. & Chen, J. USP7 substrates identified by proteomics analysis reveal the specificity of USP7. Genes Dev 36, 1016–1030 (2022).

51. Fecteau, J.F., Corral, L.G., Ghia, E.M., Gaidarova, S., Futalan, D., Bharati, I.S., Cathers, B., Schwaederle, M., Cui, B., Lopez-Girona, A., Messmer, D. & Kipps, T.J. Lenalidomide inhibits the proliferation of CLL cells via a cereblon/p21(WAF1/Cip1)-dependent mechanism independent of functional p53. Blood 124, 1637–44 (2014).

52. Schenk, P., Devine, S.M., Cobbold, S.A., Ang, C.-S., Geoghegan, N.D., Calleja, D.J., Multari, D.H., Vaibhav, V., Lu, B.G.C., Klemm, T.A., Dagley, L.F., Lowes, K.N., Williamson, N., Eichhorn, P.J.A., Ng, A.P., Feltham, R. & Komander, D. Global analysis of cancer cell responses to USP9X inhibition. bioRxiv, 2025.06.18.660475 (2025).

53. Maat, H., Atsma, T.J., Hogeling, S.M., Rodriguez Lopez, A., Jaques, J., Olthuis, M., de Vries, M.P., Gravesteijn, C., Brouwers-Vos, A.Z., van der Meer, N., Datema, S., Salzbrunn, J., Huls, G., Baas, R., Martens, J.H.A., van den Boom, V. & Schuringa, J.J. The USP7-TRIM27 axis mediates non-canonical PRC1.1 function and is a druggable target in leukemia. iScience 24, 102435 (2021).

54. Thier, B., Zhao, F., Stupia, S., Bruggemann, A., Koch, J., Schulze, N., Horn, S., Coch, C., Hartmann, G., Sucker, A., Schadendorf, D. & Paschen, A. Innate immune receptor signaling induces transient melanoma dedifferentiation while preserving immunogenicity. J Immunother Cancer 10, e003863 (2022).

55. Sipos, B., Moser, S., Kalthoff, H., Torok, V., Lohr, M. & Kloppel, G. A comprehensive characterization of pancreatic ductal carcinoma cell lines: towards the establishment of an in vitro research platform. Virchows Arch 442, 444–52 (2003).

56. Schmidt, M., Grethe, C., Recknagel, S., Kipka, G.M., Klink, N. & Gersch, M. N-Cyanopiperazines as Specific Covalent Inhibitors of the Deubiquitinating Enzyme UCHL1. Angew Chem Int Ed Engl 63, e202318849 (2024).

57. Hughes, C.S., Moggridge, S., Muller, T., Sorensen, P.H., Morin, G.B. & Krijgsveld, J. Single-pot, solid-phase-enhanced sample preparation for proteomics experiments. Nat Protoc 14, 68–85 (2019).

58. Rappsilber, J., Mann, M. & Ishihama, Y. Protocol for micro-purification, enrichment, pre-fractionation and storage of peptides for proteomics using StageTips. Nat Protoc 2, 1896–906 (2007).

59. Chambers, M.C., Maclean, B., Burke, R., Amodei, D., Ruderman, D.L., Neumann, S., Gatto, L., Fischer, B., Pratt, B., Egertson, J., Hoff, K., Kessner, D., Tasman, N., Shulman, N., Frewen, B., Baker, T.A., Brusniak, M.Y., Paulse, C., Creasy, D., Flashner, L., Kani, K., Moulding, C., Seymour, S.L., Nuwaysir, L.M., Lefebvre, B., Kuhlmann, F., Roark, J., Rainer, P., Detlev, S., Hemenway, T., Huhmer, A., Langridge, J., Connolly, B., Chadick, T., Holly, K., Eckels, J., Deutsch, E.W., Moritz, R.L., Katz, J.E., Agus, D.B., MacCoss, M., Tabb, D.L. & Mallick, P. A cross-platform toolkit for mass spectrometry and proteomics. Nat Biotechnol 30, 918–20 (2012).

60. Demichev, V., Messner, C.B., Vernardis, S.I., Lilley, K.S. & Ralser, M. DIA-NN: neural networks and interference correction enable deep proteome coverage in high throughput. Nat Methods 17, 41–44 (2020).

61. Cox, J., Hein, M.Y., Luber, C.A., Paron, I., Nagaraj, N. & Mann, M. Accurate proteome-wide label-free quantification by delayed normalization and maximal peptide ratio extraction, termed MaxLFQ. Mol Cell Proteomics 13, 2513–26 (2014).

62. Tyanova, S., Temu, T., Sinitcyn, P., Carlson, A., Hein, M.Y., Geiger, T., Mann, M. & Cox, J. The Perseus computational platform for comprehensive analysis of (prote)omics data. Nat Methods 13, 731–40 (2016).

63. Cox, J., Neuhauser, N., Michalski, A., Scheltema, R.A., Olsen, J.V. & Mann, M. Andromeda: a peptide search engine integrated into the MaxQuant environment. J Proteome Res 10, 1794–805 (2011).

64. Cox, J. & Mann, M. MaxQuant enables high peptide identification rates, individualized p.p.b.-range mass accuracies and proteome-wide protein quantification. Nat Biotechnol 26, 1367–72 (2008).

