## Supporting Information for "Targeted degradation of USP7 in solid cancer cells reveals disparate effects of deubiquitinase inhibition vs. acute protein depletion"

#### 1. Supporting Figures

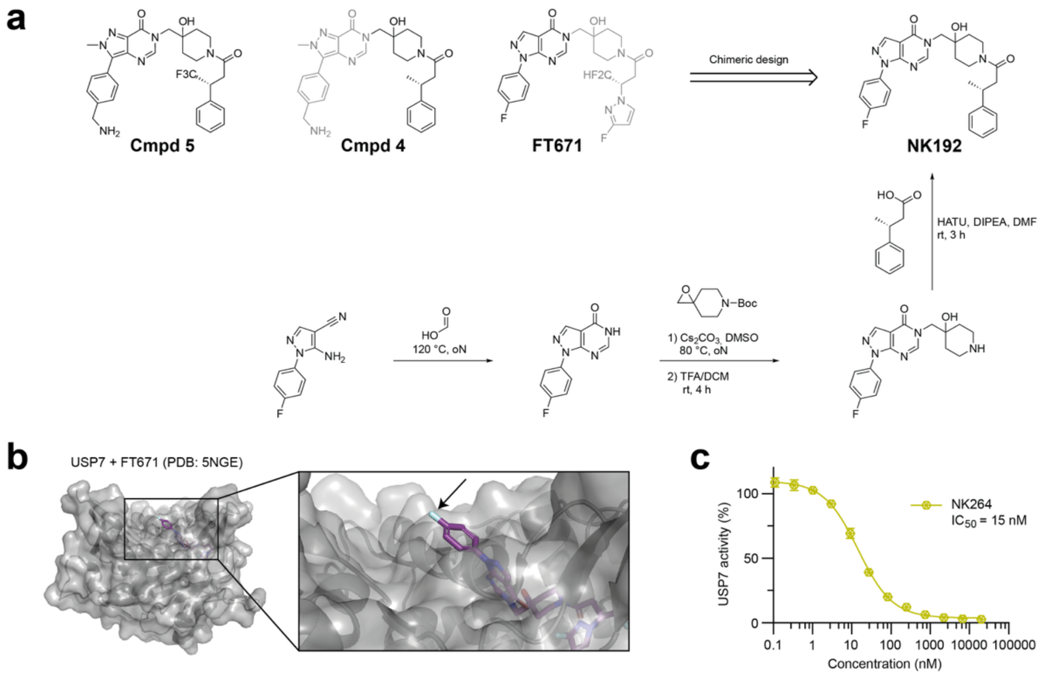

**Supporting Figure 1. Design of chimeric USP7 inhibitor NK192 and of derivatives.**

**a.** Chemical structures of previously described USP7 inhibitors Compound 5[1], Compound 4[1] and FT671[2] which were used for the design of chimeric USP7 inhibitor NK192. Its chemical synthesis is shown. Detailed synthesis procedures are shown in the chemistry methods section. **b.** Crystal structure of USP7 in complex with FT671 inhibitor (PDB: 5NGE).[2] A close-up view of the binding site is shown with the solvent-exposed 4-fluorophenyl group (which was used as exit vector for further functionalization) highlighted with an arrow. **c**. Ub-RhoG cleavage assay for biotin-functionalized compound NK264 (see Fig. 1b for its chemical structure). Data were recorded and analyzed as described for Fig. 1c.

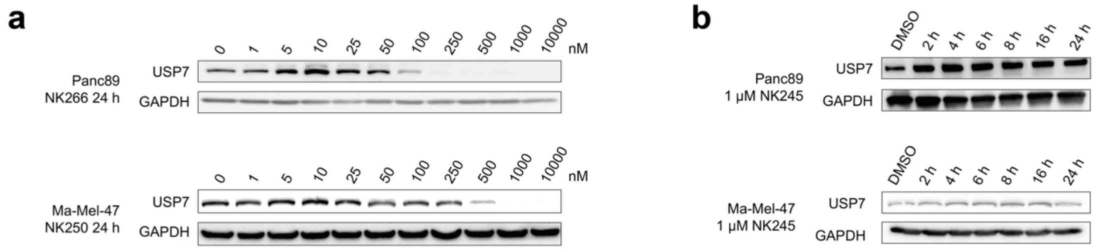

**Supporting Figure 2. E3 ligase engagement is required for USP7 degradation.**

**a.** Assessment of degradation efficiencies of PROTAC NK266 in Panc89 as well as of PROTAC NK250 in Ma-Mel-47 cells, which were treated with indicated compound concentrations for 24 h. **b.** Assays with negative control PROTAC NK245. Western blot analysis of Panc89 and Ma-Mel-47 cells treated with NK245 (1 µM) for indicated times. See Fig. 3a for the structure and rational of NK245.

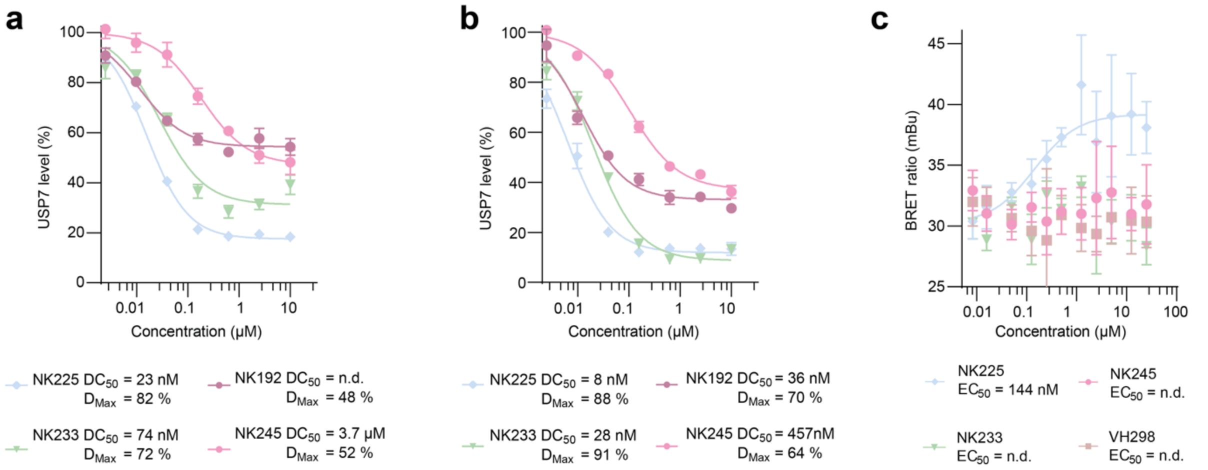

**Supporting Figure 3. Quantitative assessments of USP7 degradation efficiency and of ternary complex formation of first-generation PROTAC molecules and control compounds.**

**a.-b.** Assessment of USP7 degradation using HiBiT endpoint degradation assays in MV4-11 cells expressing HiBiT-USP7. Cells were treated for either 6 h (a) or 24 h (b) with NK192, NK225, NK233 or NK245 at varying concentrations. Data are shown as mean ± S.D. (N=3). Half-maximal degradation concentrations ($DC_{50}$) and maximal percentage of protein degradation values ($D_{Max}$) derived from these data are given below. **c.** Cellular ternary complex formation assay using PROTACs NK225 and NK233, as well as negative control PROTAC NK245 and VHL ligand VH298. The determined half-maximal ternary complex formation concentration ($EC_{50}$) for NK225 is given below. Data are shown as mean ± S.D. (N=4). mBu, milli BRET units.

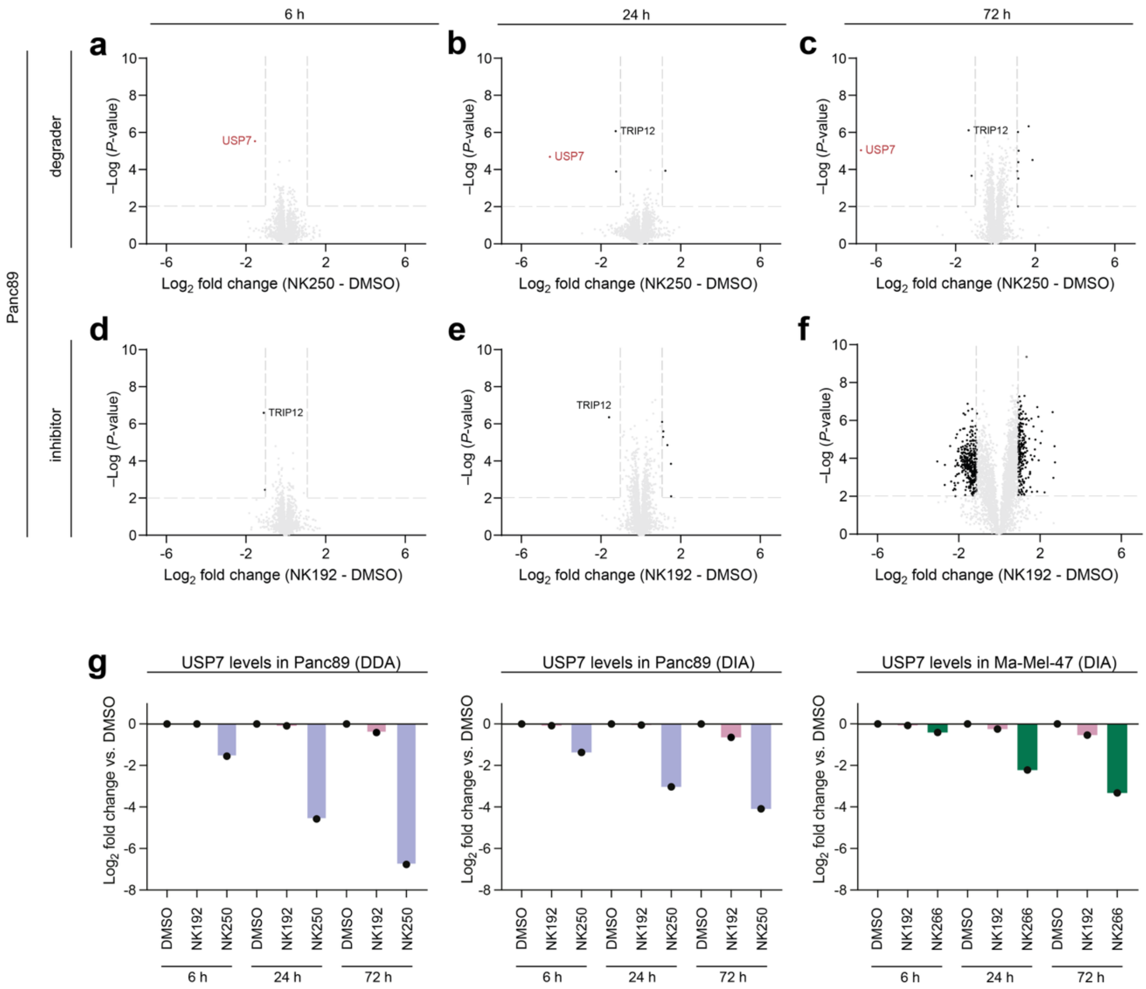

**Supporting Figure 4. Proteomic analysis of USP7 modulation in Panc89 cells reveals exceptional USP7 degradation specificity of NK250.**

**a.-f.** Proteomic analysis of Panc89 cells treated with PROTAC NK250 (a-c) or inhibitor NK192 (d-f) for indicated time points (6, 24 and 72 h). Volcano blots reporting quantitative analysis of proteins identified by mass spectrometry through data-dependent acquisition (DDA). Significantly regulated proteins are colored in black, USP7 is highlighted in red. See Fig. 5a-f for an analysis of the same samples through data-independent acquisition (DIA) mass spectrometry. **g.** $Log_2$ fold changes of USP7 protein levels compared to DMSO across all measured timepoints and for both DDA and DIA approaches. Shown is the average of the quadruplicates of each condition.

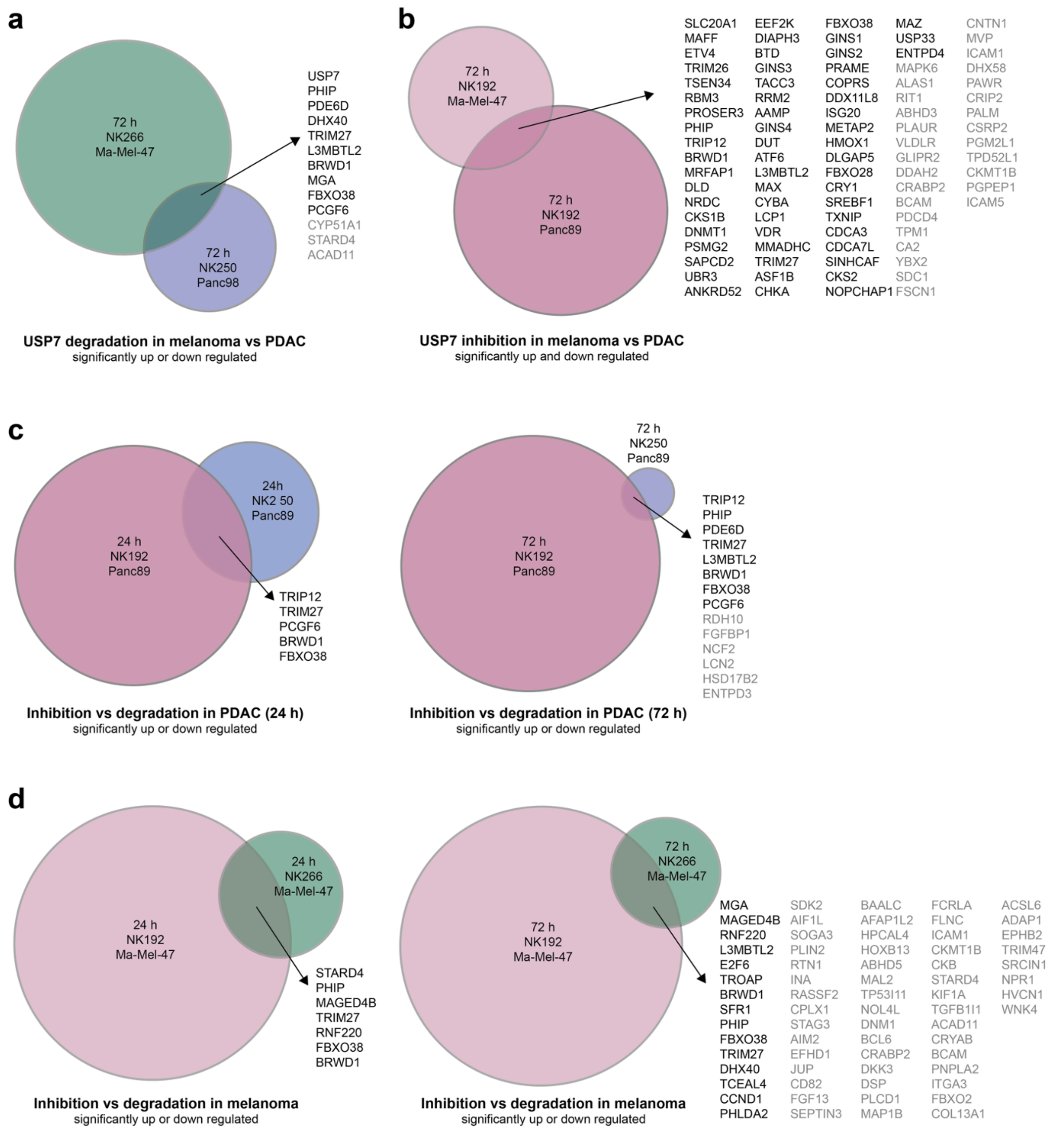

**Supporting Figure 5. Comparison of regulated proteins from DIA proteomic analysis in PDAC and melanoma cell lines.**

**a.** Venn diagram of significantly regulated proteins upon USP7 degradation with PROTACs NK266 in Ma-Mel-47 and NK250 in Panc89 after 72 h. Proteins with altered abundance in the same direction in both entities are listed, with up-regulated proteins in grey and down-regulated proteins in black. Cut-offs for significance (–Log (*P*-value) > 2) and fold change ($\log_2$ (condition vs control) > (-)1) were used as shown in the corresponding volcano plots in Fig. 5. **b.** Venn diagram of significantly regulated proteins upon USP7 inhibition with NK192 in Panc89 and Ma-Mel-47 after 72 h. Proteins with altered abundance in both cell lines are given as in panel a. **c.** Venn diagram of significantly regulated proteins upon USP7 degradation with PROTAC NK250 versus inhibition with NK192 in Panc89 cells after 24 h (left) or 72 h (right). **d.** Venn diagram of significantly regulated proteins upon USP7 degradation with PROTAC NK266 versus inhibition with NK192 in Ma-Mel-47 cells after 24 h (left) or 72 h (right).

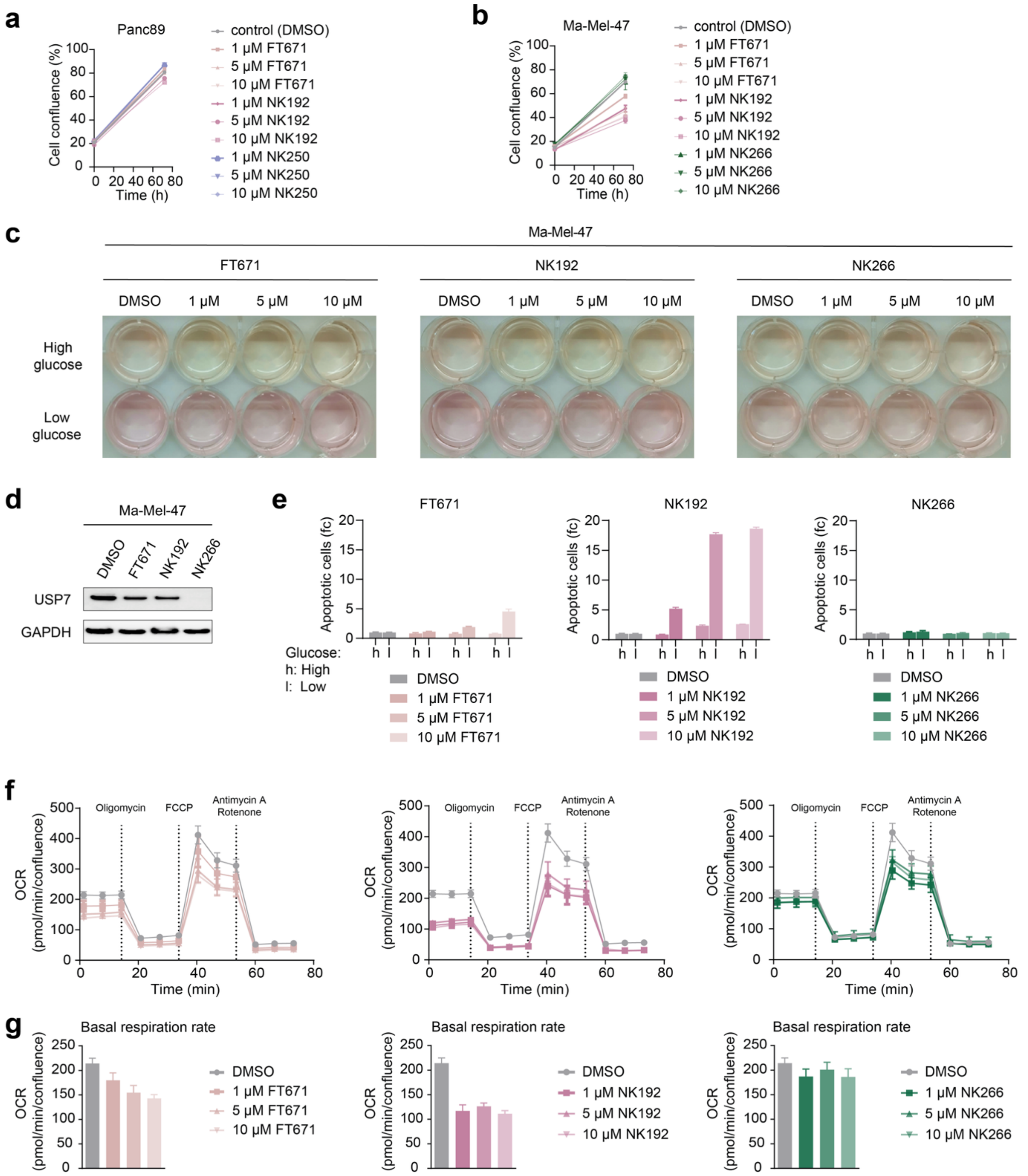

**Supporting Figure 6. Disparate effects of inhibitor vs. PROTAC treatment in Ma-Mel-47.**

**a.** Confluence of Panc89 cells treated with FT671, NK192 or NK250 at 1, 5 or 10 μM or DMSO after 72 h. **b.** Confluence of Ma-Mel-47 treated with indicated compounds after 72 h. **c.** Images of Ma-Mel-47 cells in culture medium containing 5 mM (low) or 11 mM (high) glucose, treated with indicated concentrations of FT671, NK192 or NK266 after 72 h. **d.** Western blot analysis of Ma-Mel-47 cells treated with 1 μM of indicated compounds for 72 h. **e.** Apoptosis assay. Ma-Mel-47cells treated with compounds for 72 h in culture medium containing 5 mM (low) or 11 mM (high) glucose were analyzed by FACS for Annexin-V staining. Data were analyzed as in Fig. 6c. The same DMSO samples are repeated for clarity in each graph. **f.** Seahorse mito stress test. Ma-Mel-47 cells treated with indicated concentrations of FT671, NK192 or NK266 for 24 h, followed by OCR measurements. Three independent experiments were performed in technical quintuplicates, one representative is shown as mean ± SEM. **g.** Basal respiration rates of data shown in panel f, shown as mean ± SEM.

#### 2. Supporting Tables

##### Supporting Table 1. Mass spectrometry settings for DDA analysis (ACE_0869 and ACE_0882-DDA).

| *general* | *MS1* | *MS2* | *Comments; special settings* |
|---|---|---|---|
| Tune v4.1.4244<br>Xcalibur v4.7.69.37<br>SII: 1.7.0.468<br>Gradient: 105 min | Analyzer: FT<br>Res.: 120000<br>SR: 375 - 1500<br>AGC: Standard<br>AGC abs.: 400000<br>AcT: 50 ms<br>RF: 30<br>SF: --<br>DDM: CT/3sec | Analyzer: IT<br>Res./ScR: -/rapid<br>SR: Auto<br>AGC: 300%<br>AGC abs.: 30000<br>AcT: 70 ms<br>CS: +2 to +7<br>IsM: Q<br>IsW: 1.6<br>Frag.: sHCD<br>NCE: 20, 30, 45 | classic orbitrap experiment: MS1 in Orbitrap at high resolution and data dependent MS2 also in Orbitrap high resolution. Dynamic exclusion enabled (exclude after n times=1; Exclusion duration (s)= 30; mass tolerance= ± 10 ppm)<br><br>Intensity Threshold: 5000<br>Ion transfer Tube Temp: 230 °C<br>Ion Source Voltage: 2500 V |

Note: **FT**= Fourier Transform (Orbitrap); **IT**= Iontrap; **Q**= Quadrupol; **Res.**= max. Resolution at 200 m/z (Lumos) or 400 m/z (Elite) [FWHM (full width at half maximum)]; **ScR**= scan rate for measurements in the IT; **SR**= scan range [m/z]; **AGC**= automatic gain control, max number of acquired ions per measurement; **AcT**= max. Ion acquisition time [ms]; **CS**= charge states used for fragmentation; **IsM**= Isolation mode (Q or IT), MS2 isolation and further is only done in IT; **IsW**= Isolation window [m/z], value followed by scan mode the isolation is based on (MS1, MS2 …) **Frag.**= Fragmentation method; **HCD**= Higher-energy collisional dissociation; **CID**= Collision-induced dissociation; **ETD**= Electron-transfer dissociation**; EThcD=** Electron-Transfer/Higher-Energy Collision Dissociation; **sHCD**= stepped HCD**; NCE**= normalized collision energy; **cycles**: number of MSn recorded or max cycle time; RF= RF Lens [%]; **SF**= Source Fragmentation [V]; **DDM**: Data dependent Mode (cycle time in seconds, CT/[s] or number of scans, NS); **NS**= Number of data dependent scans; **NSE**: Number of Scan Events; **WO**: Window overlap; **PMR**: Precursor Mass Range

**Supporting Table 2. Mass spectrometry settings for DIA analysis (ACE_0882-DIA and ACE_0883-DIA).**

| General | MS1 | MS2 | MS1 | MS2 | Comments; special settings |
|---|---|---|---|---|---|
| | Experiment 1 | Experiment 2 DIA 400-1000 with 8 mz window | Experiment 3 | Experiment 4 DIA with windows staggered | |
| Tune v4.1.4244<br>Xcalibur v4.7.69.37<br>SII: 1.7.0.468<br>Gradient: 105 min | Analyzer: FT<br>Res.: 60000<br>SR: 390 - 1010<br>AGC: Standard<br>AGC abs.: 400000<br>AcT: auto<br>RF: 30<br>SF: --<br>DDM: -/- | PMR: 400-1000<br>Analyzer: FT<br>Res./ScR: 15000/-<br>SR: 145-1450<br>AGC: 800%<br>AGC abs.: 400000<br>AcT: auto<br>CS: ---<br>IsM: Q<br>IsW: 8<br>WO: 0<br>NSE: 75<br>Frag.: HCD<br>NCE: 33 | Analyzer: FT<br>Res.: 60000<br>SR: 390 - 1010<br>AGC: Standard<br>AGC abs.: 400000<br>AcT: auto<br>RF: 30<br>SF: --<br>DDM: -/- | PMR: 396-1004<br>Analyzer: FT<br>Res./ScR: 30000/-<br>SR: 145-1450<br>AGC: 800%<br>AGC abs.: 400000<br>AcT: 60 ms<br>CS: ---<br>IsM: Q<br>IsW: 4<br>WO: 0<br>NSE: 152<br>Frag.: HCD<br>NCE: 33 | DIA orbitrap experiment. MS1 followed by fast DIA-MS2 followed by MS1 and slower DIA-MS2 with staggered windows.<br><br>Intensity Threshold: 5000<br>Ion transfer Tube Temp: 275 °C<br>Ion Source Voltage: 2300 V<br>Lock mass: 445.12002<br>Default charge State: +3 |

Note: See the footnote of Supporting Table 2 for a list of abbreviations.

Supporting Tables 3-6 are supplied as separate files.

#### 3. Chemical synthesis procedures

The chemicals and solvents used for this work were purchased from companies such as Activate Scientific, BLDpharm, Honeywell, Fisher Scientific, Merck, Roth, Sigma-Aldrich, TCI or VWR and were used without further purification.

Chemicals and solvents used for the synthesis are abbreviated as follows. EA: ethyl acetate, DCM: dichloromethane, MeOH: methanol, PE: petroleum ether, DMF: dimethylformamide, DMSO: dimethyl sulfoxide THF: tetrahydrofuran, TFA: trifluoroacetic acid, ACN: acetonitrile, $H_2O$: water, EtOH: ethanol.

Silica gel aluminum plates (silica gel 60 F254, Merck) were used for thin-layer chromatography. The detection was carried out using UV light at 254 nm or potassium permanganate as staining reagent.

A Pure C-850 FlashPrep system or a Pure-C-810 Flash system (Büchi) were used for automated column chromatographic purification. For preparative reverse phase column chromatographic purification, a VP125/21 Nucleodur C18 Gravity column (5 µm, Macherey Nagel) was used; for low pressure reverse phase column chromatographic purification various Ecoflex C18 columns (Büchi) were used. Compound purification by preparative HPLC was achieved with the following gradients: 0% ACN in $H_2O$ + 0.1% TFA over 6 min, 0-10% ACN in $H_2O$ + 0.1% TFA over 3 min, 10-55% ACN in $H_2O$ + 0.1% TFA over 30 min, 100% ACN + 0.1% TFA over 7 min) or: 0% ACN in $H_2O$ + 0.1% TFA over 6 min, 15-70% ACN in $H_2O$ + 0.1% TFA over 40 min, 100% ACN + 0.1% TFA over 7 min).

A 1260 series HPLC system (Agilent Technologies) with a ZORBAX Eclipse XDB column (C18 80 Å; 4.6 x 150 mm; 5 µm) was used for low resolution LC-MS analysis.

For high resolution mass spectrometry (HRMS) a 1260 Infinity II system (Agilent Technologies) using a G7129A autosampler, a G7116A column oven, a G7117C photodiode array detector and a G7111B quaternary pump system was used. The measurement was performed on a compact QTOF (Bruker Daltonics). The ionization mode was ESI (Electrospray ionization) with a source voltage of 4.5 kV.

The following devices from Bruker were used to record NMR spectra: AV 500 Avance III HD (500 MHz for $^{1}H$ and 125 MHz for $^{13}C$-NMR), AV 600 Avance III HD (600 MHz for $^{1}H$ and 151 MHz for $^{13}C$ NMR) and AV 700 Avance III HD (700 MHz for $^{1}H$ and 176 MHz for $^{13}C$ NMR).

The chemical shifts of all spectra are specified in ppm and the coupling constants $J$ are given in Hertz (Hz). Peaks of deuterated solvent were used as internal standards (DMSO-$d_6$: δ = 2.50 ppm / 39.52 ppm; $CDCl_3$: δ = 7.26 ppm / 77.16 ppm).

The multiplicities of the signals in the $^{1}H$ spectra are abbreviated as follows: s (singlet), d (doublet), dd (doublet of doublets), t (triplet), td (triplet of doublets), q (quartet), m (multiplet) and b (broad). Compounds including the (*R*)-3-phenylbutanoic acid moiety (such as NK192) showed splitting of some of the $^{13}C$ signals, as described before for molecules including this moiety[1], due to slowing exchanging conformers. In line with previously published annotations, signals showing this splitting are annotated as “d” as apparent doublet.

#### 3.1 Synthesis of a USP7 inhibitor

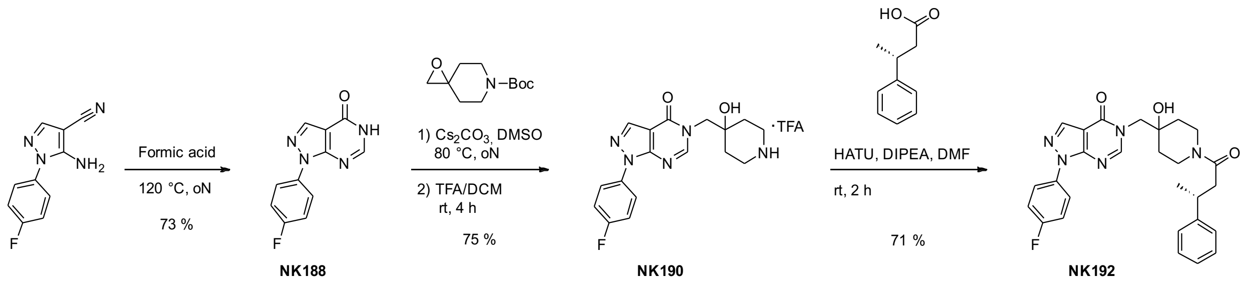

**Scheme 1.** Synthesis of NK192.

##### Synthesis of 1-(4-fluorophenyl)-1,5-dihydro-4*H*-pyrazolo[3,4-*d*]pyrimidin-4-one (NK188)

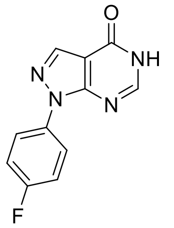

5-Amino-1-(4-fluorophenyl)-1*H*-pyrazole-4-carbonitrile (5.00 g, 24.7 mmol) was dissolved in 150 mL of formic acid. The mixture was heated to 120°C and stirred overnight. After completion, the mixture was cooled to rt for 15 min and then poured into 250 mL of ice-cold water. The mixture was filtered, and the precipitate was washed with ice-cold water (3 x 100 mL). The resulting solids were collected and dried under reduced pressure which yielded 1-(4-fluorophenyl)-1,5-dihydro-4*H*-pyrazolo[3,4-*d*]pyrimidin-4-one (NK188) (4.14 g, 18.0 mmol, 73% yield) as an off-white powder.

HRMS (m/z) $[M+H]^+$ calculated for $C_{11}H_8FN_4O^+$: 231.0667, found: 231.0675.

$^1$H NMR (500 MHz, DMSO-$d_6$) δ (ppm): 12.46 (s, 1H), 8.33 (s, 1H), 8.20 (s, 1H), 8.06 (dd, *J* = 8.9, 4.9 Hz, 2H), 7.42 (t, *J* = 8.8 Hz, 2H).

$^{13}$C NMR (126 MHz, DMSO-$d_6$) δ (ppm): 160.61 (d, *J* = 244.3 Hz), 157.17, 151.79, 148.93, 136.04, 134.62 (d, *J* = 2.9 Hz), 123.87 (d, *J* = 8.6 Hz), 116.07 (d, *J* = 22.8 Hz), 107.52.

##### Synthesis of 1-(4-fluorophenyl)-5-((4-hydroxypiperidin-4-yl)methyl)-1,5-dihydro-4*H*-pyrazolo[3,4-*d*]pyrimidin-4-one TFA salt (NK190)

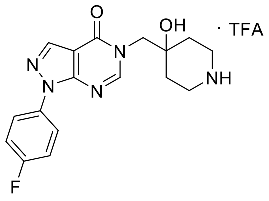

Step1:

NK188 (3.0 g, 13.0 mmol, 1.0 eq), *tert*-butyl 1-oxa-6-azaspiro[2.5]octane-6-carboxylate (4.17 g, 19.6 mmol, 1.5 eq), and cesium carbonate (12.74 g, 39.1 mmol, 3 eq) were added to a dry round bottom flask to which was added dry DMSO (30 mL). The reaction was heated to 80°C and stirred overnight. After completion, the mixture was diluted with water (100 mL) and then extracted with DCM (3x 100 mL). The combined organic layers were dried over $MgSO_4$ and concentrated under reduced pressure. The resulting oil was purified by reverse phase flash column chromatography (ACN/$H_2O$ + 0.1 % TFA) yielding *tert*-butyl 4-((1-(4-fluorophenyl)-4-oxo-1,4-dihydro-5*H*-pyrazolo[3,4-*d*]pyrimidin-5-yl)methyl)-4-hydroxypiperidine-1-carboxylate (NK189) (4.31 g, 9.73 mmol, 75 % yield) as an off-white powder.

HRMS (m/z) $[M+H]^+$ calculated for $C_{22}H_{26}FN_5NaO_4^+$: 466.1861, found: 466.1850.

$^1$H NMR (500 MHz, DMSO-$d_6$) δ (ppm): 8.37 (s, 1H), 8.34 (s, 1H), 8.09 – 8.05 (m, 2H), 7.45 – 7.40 (m, 2H), 4.90 (s, 1H), 4.03 (s, 2H), 3.66 (d, *J* = 13.1 Hz, 2H), 3.04 (s, 2H), 1.48 (ddd, *J* = 13.2, 11.2, 4.7 Hz, 2H), 1.39 (s, 9H), 1.38 – 1.33 (m, 2H).

$^{13}$C NMR (126 MHz, DMSO-$d_6$) δ (ppm): 160.61 (d, *J* = 244.3 Hz), 156.86, 153.82, 152.39, 151.08, 136.33, 134.53 (d, *J* = 2.8 Hz), 123.71 (d, *J* = 8.6 Hz), 116.14 (d, *J* = 23.0 Hz), 106.64, 78.52, 68.99, 53.05, 34.21, 28.08.

Step 2:

NK189 (4.31 g, 9.84 mmol) was dissolved in TFA/DCM (50:50, 30 mL) and stirred at rt for 3 h. After completion, the solvents were evaporated under reduced pressure. The resulting off-white solids of the product 1-(4-fluorophenyl)-5-((4-hydroxypiperidin-4-yl)methyl)-1,5-dihydro-4*H*-pyrazolo[3,4-*d*]pyrimidin-4-one TFA salt (NK190) (4.5 g, 9.84 mmol, quant. yield) were used for the next step without further purification.

HRMS (m/z) $[M+H]^+$ calculated for $C_{17}H_{19}FN_5O_2^+$: 344.1517, found: 344.1510.

**Synthesis of (*R*)-1-(4-fluorophenyl)-5-((4-hydroxy-1-(3-phenylbutanoyl)piperidin-4-yl)methyl)-1,5-dihydro-4*H*-pyrazolo[3,4-*d*]pyrimidin-4-one (NK192)**

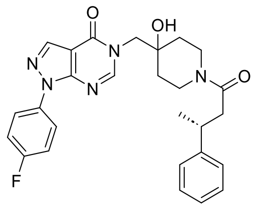

(*R*)-3-Phenylbutanoic acid (10.4 mg, 0.06 mmol, 1.2 eq), DIPEA (34.0 mg, 0.26 mmol, 5 eq) and HATU (26.03 mg, 0.07 mmol, 1.3 eq) were dissolved in DMF (1 mL) and stirred at rt for 10 minutes. To this was added NK190 (20.0 mg, 0.05 mmol, 1 eq) and the reaction was stirred for 2 h until completion. The solvent was evaporated under reduced pressure and the resulting crude product was directly purified by preparative reverse phase column chromatography. The product (*R*)-1-(4-fluorophenyl)-5-((4-hydroxy-1-(3-phenylbutanoyl)piperidin-4-yl)methyl)-1,5-dihydro-4*H*-pyrazolo[3,4-*d*]pyrimidin-4-one (NK192) (18.4 mg, 0.04 mmol, 71 % yield) was obtained as a white powder.

HRMS (m/z) $[M+Na]^+$ calculated for $C_{27}H_{28}FN_5NaO_3^+$: 512.2068, found: 512.2076.

$^1$H NMR (500 MHz, DMSO-$d_6$) δ (ppm): 8.37 (s, 1H), 8.33 (d, $J$ = 11.5 Hz, 1H), 8.09 – 8.05 (m, 2H), 7.45 – 7.41 (m, 2H), 7.26 (dd, $J$ = 8.1, 5.9 Hz, 4H), 7.16 (ddt, $J$ = 6.6, 5.0, 2.3 Hz, 1H), 4.91 (s, 1H), 4.09 – 3.98 (m, 2H), 3.95 (d, $J$ = 2.8 Hz, 1H), 3.65 (ddd, $J$ = 14.6, 9.4, 4.7 Hz, 1H), 3.25 – 3.12 (m, 2H), 2.91 – 2.83 (m, 1H), 2.65 – 2.53 (m, 2H), 1.57 – 1.24 (m, 4H), 1.20 (d, $J$ = 6.9 Hz, 3H).

$^{13}$C NMR (126 MHz, DMSO-$d_6$) δ (ppm): 169.13 (“d”, $J$ = 2.2 Hz), 160.62 (d, $J$ = 244.4 Hz), 156.82 (“d”, $J$ = 6.8 Hz), 152.36 (“d”, $J$ = 2.5 Hz), 151.08 (“d”, $J$ = 1.5 Hz), 146.64 ("d”, $J$ = 13.5 Hz), 136.33, 134.53 (d, $J$ = 3.0 Hz), 128.21 (“d”, $J$ = 4.5 Hz), 126.91 (“d”, $J$ = 3.7 Hz), 125.95 (“d”, $J$ = 5.0 Hz), 123.72 (d, $J$ = 8.5 Hz), 116.16 (d, $J$ = 23.1 Hz), 106.66, 69.11 (“d”, $J$ = 6.1 Hz), 53.06, 41.00 (“d”, $J$ = 14.5 Hz), 40.24 (“d”, $J$ = 3.2 Hz), 36.92, 36.12 (“d”, $J$ = 26.3 Hz), 34.89 (“d”, $J$ = 13.9 Hz), 34.17 (“d”, $J$ = 16.7 Hz), 21.99 (“d”, $J$ = 25.1 Hz).

#### 3.2 Synthesis of functionalized USP7 inhibitors

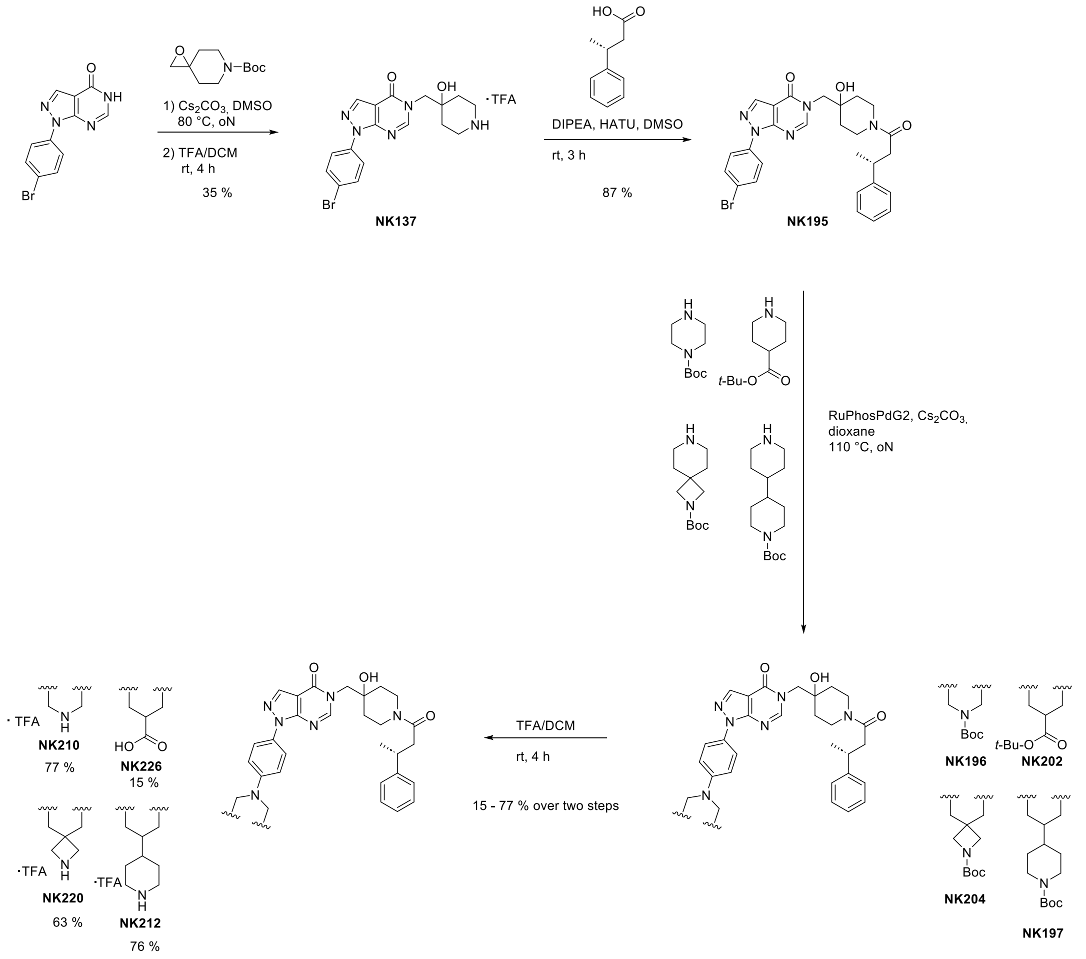

**Scheme 2.** Synthesis of USP7 inhibitors NK210, NK226, NK220 and NK212.

##### Synthesis of 1-(4-bromophenyl)-5-((4-hydroxypiperidin-4-yl)methyl)-1,5-dihydro-4*H*-pyrazolo[3,4-*d*]pyrimidin-4-one (NK137)

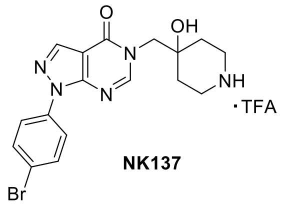

NK137

Step 1:

To a solution of 1-(4-bromophenyl)-1,5-dihydro-4H-pyrazolo[3,4-*d*]pyrimidin-4-one (3.00 g, 10.3 mmol, 1.0 eq) in dry DMF (25 mL) were added *tert*-butyl 1-oxa-6-azaspiro[2.5]octane-6-carboxylate (2.4 g, 11.4 mmol, 1.1 eq) and cesium carbonate (10.1 g, 31.0 mmol, 3.0 eq) at rt. The reaction was heated to 80 °C and stirred overnight. The mixture was then diluted with $H_2O$ (20 mL) and extracted with EA (3x 50 mL). The combined organic layers were dried over $MgSO_4$, and the solvent was removed under reduced pressure. The resulting crude product was purified by flash column chromatography (PE/EA) which yielded the product *tert*-butyl 4-((1-(4-bromophenyl)-4-oxo-1,4-dihydro-5*H*-pyrazolo[3,4-*d*]pyrimidin-5-yl)methyl)-4-hydroxypiperidine-1-carboxylate (NK135) as an off-white solid (1.8 g, 3.66 mmol, 35 %).

HRMS (m/z) $[M+Na]^+$ calculated for $C_{22}H_{26}BrN_5NaO_4^+$: 526.1060, found: 526.1065.

$^1$H NMR (500 MHz, DMSO-$d_6$) δ (ppm): 8.39 (s, 1H), 8.36 (s, 1H), 8.07 (d, $J$ = 8.9 Hz, 2H), 7.78 (d, $J$ = 8.8 Hz, 2H), 4.90 (s, 1H), 4.03 (d, $J$ = 2.0 Hz, 2H), 3.66 (d, $J$ = 13.0 Hz, 2H), 3.05 (s, 2H), 1.48 (ddd, $J$ = 13.4, 11.1, 4.6 Hz, 2H), 1.39 (s, 9H), 1.35 (s, 2H).

$^{13}$C NMR (126 MHz, DMSO-$d_6$) δ (ppm): 156.78, 153.81, 152.49, 151.29, 137.46, 136.66, 132.24, 123.08, 119.54, 106.96, 78.51, 68.98, 53.08, 34.20, 28.08.

Step 2:

NK135 (1.6 g, 3.2 mmol) was dissolved in TFA/DCM (20:80, 20 mL) and the reaction was stirred at rt for 4 h. After completion, the solvents were evaporated under reduced pressure which yielded 1-(4-bromophenyl)-5-((4-hydroxypiperidin-4-yl)methyl)-1,5-dihydro-4*H*-pyrazolo[3,4-*d*]pyrimidin-4-one (NK137) as an off-white solid (1.68 g, 3.2 mmol, quant. yield). The product was used for the next step without further purification.

HRMS (m/z) $[M+H]^+$ calculated for $C_{17}H_{19}BrN_5O_2^+$: 404.0717, found: 404.0709.

$^1H$ NMR (500 MHz, DMSO-$d_6$) δ (ppm): 8.61 (s, 1H), 8.40 (d, *J* = 8.7 Hz, 2H), 8.32 (s, 1H), 8.06 (d, *J* = 8.4 Hz, 2H), 7.78 (d, *J* = 8.4 Hz, 2H), 5.30 (s, 1H), 4.09 (s, 2H), 3.21 – 3.13 (m, 2H), 3.02 (t, *J* = 11.5 Hz, 2H), 1.83 – 1.73 (m, 2H), 1.58 (d, *J* = 14.3 Hz, 2H).

$^{13}C$ NMR (126 MHz, DMSO-$d_6$) δ (ppm): 156.81, 152.44, 151.28, 137.41, 136.67, 132.29, 123.11, 119.64, 106.94, 67.36, 52.80, 31.04.

**Synthesis of (*R*)-1-(4-bromophenyl)-5-((4-hydroxy-1-(3-phenylbutanoyl)piperidin-4-yl)methyl)-1,5-dihydro-4*H*-pyrazolo[3,4-*d*]pyrimidin-4-one (NK195)**

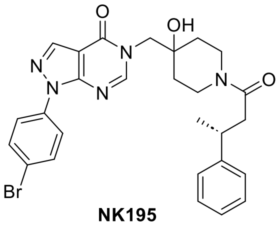

**NK195**

(*R*)-3-phenylbutanoic acid (587.9 mg, 3.58 mmol, 1.2 eq), DIPEA (1.54 g, 0.01 mol, 4.0 eq) and HATU (1.59 g, 4.18 mmol, 1.4 eq) in DMSO (10 mL) were added to a round bottom flask and were stirred at rt for 15 min. To this was added NK137 (1.31 g, 2.98 mmol, 1 eq). The reaction was stirred for 3 h at rt. The crude product was directly purified by reverse phase flash column chromatography (ACN/$H_2O$ + 0.1 % TFA). The product (*R*)-1-(4-bromophenyl)-5-((4-hydroxy-1-(3-phenylbutanoyl)piperidin-4-yl)methyl)-1,5-dihydro-4*H*-pyrazolo[3,4-*d*]pyrimidin-4-one (NK195) was obtained as an off-white solid (1.43 g, 2.6 mmol, 87 % yield).

HRMS (m/z) $[M+Na]^+$ calculated for $C_{27}H_{28}BrN_5NaO_3^+$: 572.1268, found: 572.1266.

$^1H$ NMR (600 MHz, DMSO-$d_6$) δ (ppm): 8.39 (s, 3H), 8.34 (d, *J* = 13.8 Hz, 3H), 8.06 (d, *J* = 8.9 Hz, 4H), 7.77 (d, *J* = 8.9 Hz, 4H), 7.26 – 7.24 (m, 10H), 7.15 (ddt, *J* = 8.4, 5.7, 2.0 Hz, 3H), 4.09 – 3.94 (m, 6H), 3.95 (d, *J* = 3.7 Hz, 3H), 3.69 – 3.62 (m, 22H), 3.25 – 3.14 (m, 5H), 2.92 – 2.83 (m, 3H), 2.66 – 2.52 (m, 4H), 1.57 – 1.30 (m, 10H), 1.20 (dd, *J* = 7.0, 1.3 Hz, 10H).

$^{13}C$ NMR (151 MHz, DMSO-$d_6$) δ (ppm): 169.12 ("d", *J* = 2.8 Hz), 156.75 ("d2, *J* = 8.1 Hz), 152.45 ("d", *J* = 3.2 Hz), 151.92 – 150.78 (m), 146.64 ("d", *J* = 16.5 Hz), 137.47, 136.66, 132.25, 128.21 ("d", *J* = 5.5 Hz), 126.90 ("d", *J* = 4.8 Hz), 125.95 ("d", *J* = 6.2 Hz), 123.09, 119.56, 106.98, 69.11 ("d", *J* = 7.2 Hz), 53.10 ("d", *J* = 2.5 Hz), 41.00 ("d", *J* = 17.5 Hz), 40.25 ("d", *J* = 4.5 Hz), 36.92, 36.12 ("d", *J* = 31.9 Hz), 34.88 ("d", *J* = 16.5 Hz), 34.16 ("d", *J* = 20.1 Hz), 21.99 ("d", *J* = 30.6 Hz).

**Synthesis of (*R*)-5-((4-hydroxy-1-(3-phenylbutanoyl)piperidin-4-yl)methyl)-1-(4-(piperazin-1-yl)phenyl)-1,5-dihydro-4*H*-pyrazolo[3,4-*d*]pyrimidin-4-one TFA salt (NK210)**

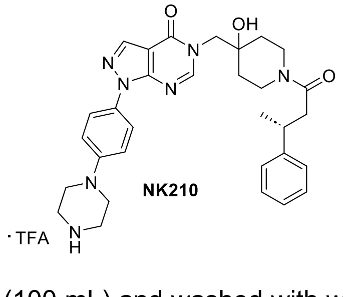

Step 1:

To a dry three necked round bottom flask under argon were added 1,4-dioxane (50 mL), NK195 (800 mg, 1.45 mmol, 1.0 eq), Boc-piperazine (379 mg, 2.03 mmol, 1.4 eq), cesium carbonate (1.42 g, 4.36 mmol, 3 eq) and RuPhosPd2G (112.9 mg, 0.15 mmol, 0.1 eq). The reaction was heated to 110 °C and stirred under argon atmosphere overnight. After completion, the mixture was filtered over Celite, and the solvents were removed under reduced pressure. The residue was resuspended with EA (100 mL) and washed with water and brine (3x 20 mL each). The organic layer was dried with $MgSO_4$, filtered and concentrated under reduced pressure. The resulting solids were then purified by reverse phase flash column chromatography (ACN/$H_2O$ + 0.1 % TFA) which yielded the product *tert*-butyl (*R*)-

4-(4-(5-((4-hydroxy-1-(3-phenylbutanoyl)piperidin-4-yl)methyl)-4-oxo-4,5-dihydro-1*H*-pyrazolo[3,4-*d*]pyrimidin-1-yl)phenyl)piperazine-1-carboxylate (NK196) as an off-white powder.

Step 2:

NK196 (730.0 mg, 1.11 mmol) was dissolved in TFA/DCM (50:50) and stirred at rt for 3 h. After completion, the solvents were evaporated under reduced pressure. The resulting product (*R*)-5-((4-hydroxy-1-(3-phenylbutanoyl)piperidin-4-yl)methyl)-1-(4-(piperazin-1-yl)phenyl)-1,5-dihydro-4*H*-pyrazolo[3,4-*d*]pyrimidin-4-one (NK210) (723 mg, 1.08 mmol, 77 % yield over two steps) was obtained as an off-white powder and used for the next steps without further purification.

HRMS (m/z) $[M+H]^+$ calculated for $C_{31}H_{38}N_7O_3^+$: 556.3031, found: 556.3040.

$^1$H NMR (600 MHz, DMSO-$d_6$) δ (ppm): 8.93 (s, 2H), 8.33 – 8.27 (m, 2H), 7.86 (d, *J* = 9.1 Hz, 2H), 7.26 (dd, *J* = 8.8, 6.2 Hz, 4H), 7.20 – 7.13 (m, 3H), 4.11 – 3.90 (m, 3H), 3.65 (td, *J* = 10.8, 9.1, 4.0 Hz, 1H), 3.43 (dd, *J* = 6.6, 3.8 Hz, 4H), 3.30 – 3.25 (m, 4H), 3.17 (dt, *J* = 14.5, 7.4 Hz, 2H), 2.91 – 2.82 (m, 1H), 2.67 – 2.54 (m, 2H), 1.59 – 1.24 (m, 4H), 1.20 (dd, *J* = 7.0, 1.3 Hz, 3H).

$^{13}$C NMR (151 MHz, DMSO-$d_6$) δ (ppm): 169.12 (“d”, *J* = 2.8 Hz), 156.91 (“d”, *J* = 8.1 Hz), 151.99 (“d”, *J* = 3.3 Hz), 150.64 (“d”, *J* = 2.1 Hz), 148.88, 146.64 (“d”, *J* = 16.4 Hz), 135.72, 130.58, 128.21 (“d”, *J* = 5.0 Hz), 126.91 (“d”, *J* = 3.9 Hz), 125.95 (“d”, *J* = 5.6 Hz), 122.88, 116.13, 106.33, 69.12 (“d”, *J* = 7.2 Hz), 52.99, 45.37, 42.64, 41.02 (“d”, *J* = 17.0 Hz), 40.25 (“d”, *J* = 5.9 Hz), 36.94, 36.13 (“d”, *J* = 31.6 Hz), 34.91 (“d”, *J* = 17.4 Hz), 34.17 (“d”, *J* = 20.4 Hz), 21.99 (“d”, *J* = 30.1 Hz).

**Synthesis of (*R*)-1-(4-([4,4'-bipiperidin]-1-yl)phenyl)-5-((4-hydroxy-1-(3-phenylbutanoyl)piperidin-4-yl)methyl)-1,5-dihydro-4*H*-pyrazolo[3,4-*d*]pyrimidin-4-one TFA salt (NK212)**

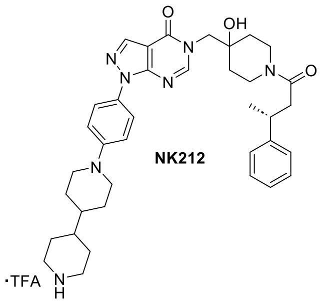

(*R*)-1-(4-([4,4'-Bipiperidin]-1-yl)phenyl)-5-((4-hydroxy-1-(3-phenylbutanoyl)piperidin-4-yl)methyl)-1,5-dihydro-4*H*-pyrazolo[3,4-*d*]pyrimidin-4-one TFA salt (NK212) was synthesized using the same method as described for NK210 (Scheme 2, exchanging *tert*-butyl piperazine-1-carboxylate for *tert*-butyl [4,4'-bipiperidine]-1-carboxylate).

HRMS (m/z) $[M+H]^+$ calculated for $C_{37}H_{48}N_7O_3^+$: 638.3813, found: 638.3806.

$^1$H NMR (600 MHz, DMSO-$d_6$) δ (ppm): 8.50 (d, *J* = 11.7 Hz, 1H), 8.30 (s, 1H), 8.28 (d, *J* = 14.1 Hz, 1H), 8.21 (d, *J* = 10.9 Hz, 1H), 7.84 (d, *J* = 8.7 Hz, 2H), 7.26 (dd, *J* = 8.6, 6.1 Hz, 4H), 7.19 (d, *J* = 8.5 Hz, 2H), 7.15 (ddd, *J* = 8.6, 6.4, 2.8 Hz, 1H), 4.08 – 3.98 (m, 2H), 3.98 – 3.90 (m, 1H), 3.83 – 3.78 (m, 2H), 3.70 – 3.61 (m, 1H), 3.30 (d, *J* = 12.3 Hz, 2H), 3.26 – 3.12 (m, 2H), 2.89 – 2.78 (m, 5H), 2.65 – 2.52 (m, 2H), 1.85 (d, *J* = 13.2 Hz, 2H), 1.80 (d, *J* = 10.0 Hz, 3H), 1.46 – 1.27 (m, 10H), 1.20 (dd, *J* = 7.0, 1.3 Hz, 3H).

$^{13}$C NMR (151 MHz, DMSO-$d_6$) δ (ppm): 169.11 (“d”, *J* = 2.8 Hz), 162.31, 156.90 (“d”, *J* = 8.1 Hz), 151.94 (“d”, *J* = 3.3 Hz), 150.62 (“d”, *J* = 2.0 Hz), 146.64 (“d”, *J* = 16.5 Hz), 135.69, 128.21 (“d”, *J* = 4.9 Hz), 126.90 (“d”, *J* = 4.0 Hz), 125.95 (“d”, *J* = 5.5 Hz), 122.89, 116.60 – 116.08 (m), 106.30, 69.12 (“d”, *J* = 7.2 Hz), 52.99, 49.51, 43.53, 41.01 (“d”, *J* = 16.7 Hz), 40.24 (“d”, *J* = 5.9 Hz), 37.75, 36.93, 36.12 (“d”, *J* = 31.3 Hz), 34.91 (“d”, *J* = 17.1 Hz), 34.17 (“d”, *J* = 20.2 Hz), 30.77, 28.00, 25.68, 21.99 (“d”, *J* = 30.1 Hz).

**Synthesis of (*R*)-1-(4-(2,7-diazaspiro[3.5]nonan-7-yl)phenyl)-5-((4-hydroxy-1-(3-phenylbutanoyl)piperidin-4-yl)methyl)-1,5-dihydro-4*H*-pyrazolo[3,4-*d*]pyrimidin-4-one TFA salt (NK220)**

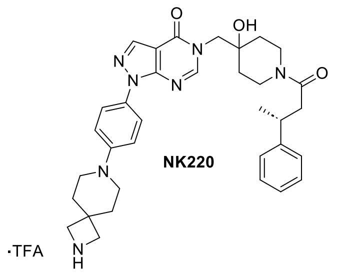

(*R*)-1-(4-(2,7-diazaspiro[3.5]nonan-7-yl)phenyl)-5-((4-hydroxy-1-(3-phenylbutanoyl)piperidin-4-yl)methyl)-1,5-dihydro-4*H*-pyrazolo[3,4-*d*]pyrimidin-4-one TFA salt **(**NK220) was synthesized using the same method as described for NK210 (Scheme 2, exchanging *tert*-butyl piperazine-1-carboxylate for *tert*-butyl 2,7-diazaspiro[3.5]nonane-2-carboxylate).

HRMS (m/z) $[M+H]^+$ calculated for $C_{34}H_{42}N_7O_3^+$: 596.3344, found: 596.3351.

$^1$H NMR (600 MHz, DMSO-*d*6) δ (ppm): 8.71 (s, 2H), 8.29 (d, *J* = 0.8 Hz, 1H), 8.27 (d, *J* = 14.1 Hz, 1H), 7.79 (d, *J* = 9.1 Hz, 2H), 7.26 (dd, *J* = 8.3, 6.1 Hz, 4H), 7.17 – 7.10 (m, 3H), 4.07 – 3.92 (m, 3H), 3.78 – 3.74 (m, 4H), 3.66 (t, *J* = 13.1 Hz, 1H), 3.27 – 3.15 (m, 6H), 2.87 (ddt, *J* = 17.6, 10.1, 4.2 Hz, 1H), 2.64 – 2.52 (m, 2H), 1.88 – 1.85 (m, 4H), 1.56 – 1.23 (m, 4H), 1.20 (d, *J* = 6.7 Hz, 3H).

$^{13}$C NMR (151 MHz, DMSO-*d*6) δ (ppm): 169.11 (“d”, *J* = 3.0 Hz), 156.91 (“d”, *J* = 8.1 Hz), 151.87 (“d”, *J* = 3.4 Hz), 150.51 (“d”, *J* = 2.1 Hz), 149.44, 146.64 (“d”, *J* = 16.3 Hz), 135.56, 129.47, 128.20 (“d2, *J* = 4.7 Hz), 126.90 (“d”, *J* = 3.8 Hz), 125.94 (“d”, *J* = 5.4 Hz), 122.88, 115.78, 106.23, 69.11 (“d”, *J* = 7.2 Hz), 54.79, 54.67, 52.97, 45.01, 41.00 (“d”, *J* = 16.9 Hz), 40.43, 40.23 (“d”, *J* = 6.5 Hz), 36.92, 36.22, 36.17, 36.01, 34.90 (“d”, *J* = 17.1 Hz), 34.16 (“d”, *J* = 20.0 Hz), 33.32, 21.99 (“d”, *J* = 29.8 Hz).

**Synthesis of (*R*)-1-(4-(5-((4-hydroxy-1-(3-phenylbutanoyl)piperidin-4-yl)methyl)-4-oxo-4,5-dihydro-1*H*-pyrazolo[3,4-*d*]pyrimidin-1-yl)phenyl)piperidine-4-carboxylic acid (NK226)**

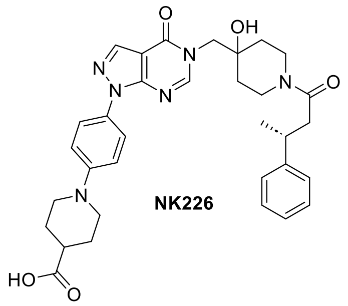

(*R*)-1-(4-(5-((4-hydroxy-1-(3-phenylbutanoyl)piperidin-4-yl)methyl)-4-oxo-4,5-dihydro-1*H*-pyrazolo[3,4-*d*]pyrimidin-1-yl)phenyl)piperidine-4-carboxylic acid **(**NK226) was synthesized using the same method as described for NK210 (Scheme 2, exchanging *tert*-butyl piperazine-1-carboxylate for *tert*-butyl piperidine-4-carboxylate) with an additional preparative reverse phase column chromatography (ACN/$H_2O$ + 0.1 % TFA) purification after Step 2.

HRMS (m/z) $[M+H]^+$ calculated for $C_{33}H_{39}N_6O_5^+$: 599.2976, found: 599.2977.

$^1$H NMR (700 MHz, DMSO-*d*6) δ (ppm): 8.30 (d, *J* = 1.0 Hz, 1H), 8.28 (d, *J* = 16.4 Hz, 1H), 7.84 (d, *J* = 8.5 Hz, 2H), 7.27 – 7.24 (m, 4H), 7.20 – 7.14 (m, 3H), 4.07 – 3.98 (m, 2H), 3.94 (d, *J* = 5.0 Hz, 1H), 3.72 (dt, *J* = 12.7, 3.9 Hz, 2H), 3.66 (t, *J* = 14.2 Hz, 1H), 3.25 – 3.14 (m, 2H), 2.96 – 2.84 (m, 3H), 2.64 – 2.57 (m, 2H), 1.95 (dd, *J* = 13.6, 3.7 Hz, 2H), 1.73 – 1.66 (m, 2H), 1.56 – 1.40 (m, 1H), 1.39 – 1.22 (m, 4H), 1.20 (dd, *J* = 6.9, 1.5 Hz, 3H).

$^{13}$C NMR (176 MHz, DMSO-*d*6) δ (ppm): 175.74, 169.11 (“d”, *J* = 3.4 Hz), 156.91 (“d”, *J* = 9.5 Hz), 151.93 (“d”, *J* = 3.7 Hz), 150.60 (“d”, *J* = 1.6 Hz), 147.12, 146.64 (“d”, *J* = 19.1 Hz), 135.66, 128.20 (“d”, *J* = 5.9 Hz), 126.90 (“d”, *J* = 4.7 Hz), 125.94 (“d”, *J* = 6.9 Hz), 122.86, 116.44, 116.40, 106.29, 69.11 (“d”, *J* = 8.5 Hz), 53.00, 48.51, 41.00 (“d”, *J* = 19.5 Hz), 40.43, 40.23 (“d”, *J* = 6.9 Hz), 36.92, 36.12 (“d”, *J* = 36.4 Hz), 34.89 (“d”, *J* = 19.3 Hz), 34.17 (“d”, *J* = 23.3 Hz), 27.24, 21.98 (“d”, *J* = 34.6 Hz).

#### 3.3 Synthesis of further functionalized USP7 inhibitors

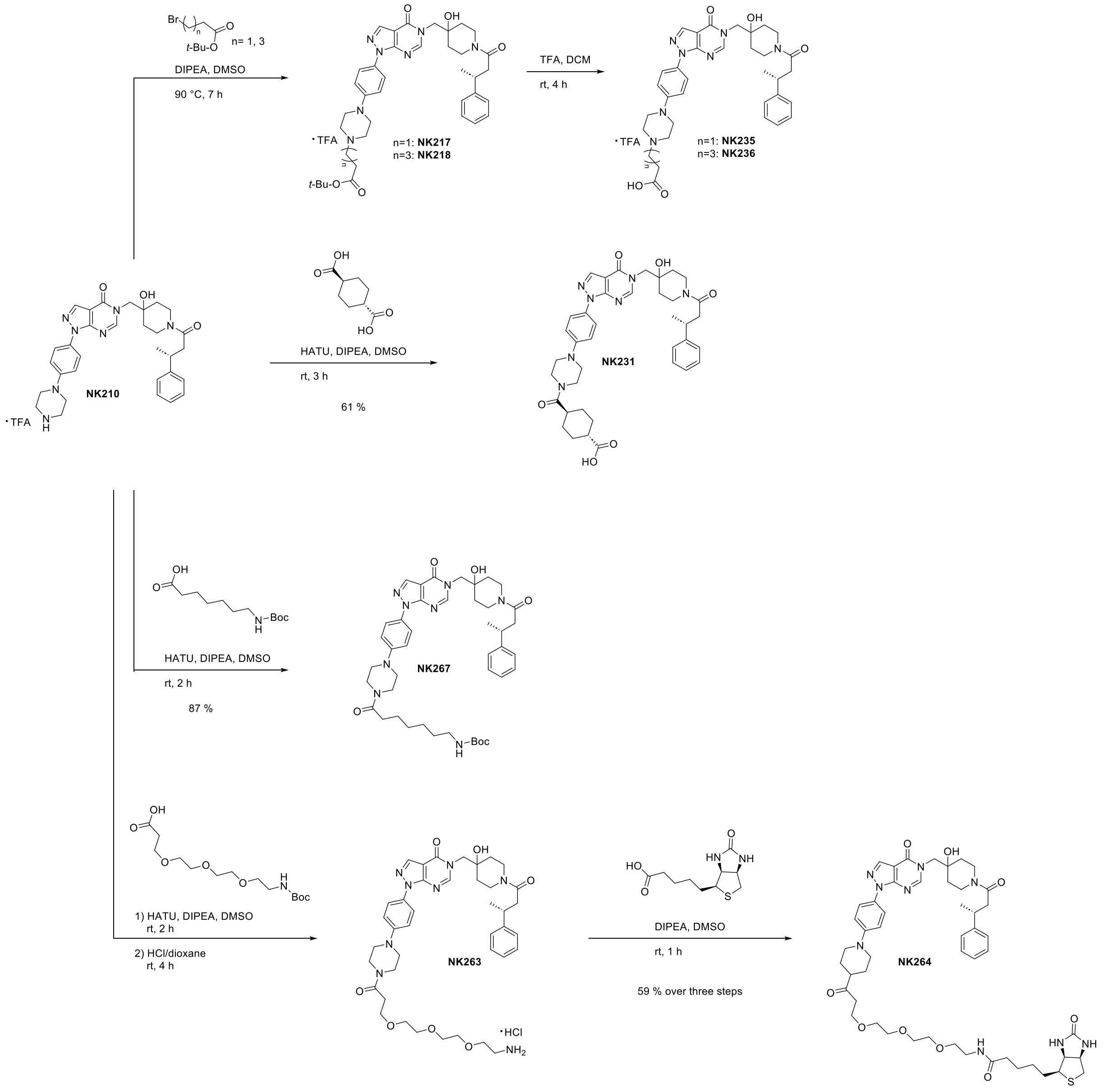

**Scheme 3.** Synthesis of further functionalized inhibitors NK235, NK236, NK231, NK267 and NK264.

##### Synthesis of (*R*)-3-(4-(4-(5-((4-hydroxy-1-(3-phenylbutanoyl)piperidin-4-yl)methyl)-4-oxo-4,5-dihydro-1*H*-pyrazolo[3,4-*d*]pyrimidin-1-yl)phenyl)piperazin-1-yl)propanoic acid TFA salt (NK235)

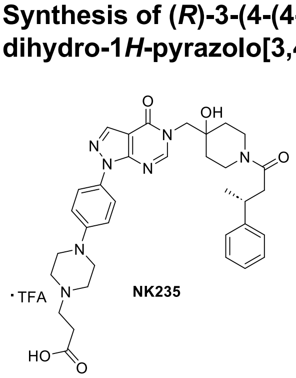

NK235

Step1:

NK210 (20.0 mg, 0.03 mmol, 1 eq), *tert*-butyl 3-bromopropanoate (12.5 mg, 0.06 mmol, 2 eq) and DIPEA (15.44 mg, 0.12 mmol, 4 eq) were dissolved in DMSO (1 mL) and stirred for 7 h at 90 °C. After cooling to rt, the crude mixture was directly purified by preparative reverse phase column chromatography (ACN/$H_2O$ + 0.1 % TFA). The fractions containing the product *tert*-butyl (*R*)-3-(4-(4-(5-((4-hydroxy-1-(3-phenylbutanoyl)piperidin-4-yl)methyl)-4-oxo-4,5-dihydro-1*H*-pyrazolo [3,4-*d*]pyrimidin-1-yl)phenyl)piperazin-1-yl)propanoate (NK217) were combined and most of the solvents were evaporated under reduced pressure. The product was then directly used for the next step.

Step 2:

The combined fractions from the previous step that contained NK217 were diluted with DCM/TFA (50:50, 2 mL) and stirred at rt for 4 h. After completion, the solvents were evaporated under reduced pressure which yielded the crude product (*R*)-3-(4-(4-(5-((4-hydroxy-1-(3-phenylbutanoyl)piperidin-4-yl)methyl)-4-oxo-4,5-dihydro-1*H*-pyrazolo[3,4-*d*]pyrimidin-1-yl)phenyl)piperazin-1-yl)propanoic acid TFA salt (NK235) which was used directly for the next step.

**Synthesis of (*R*)-5-(4-(4-(5-((4-hydroxy-1-(3-phenylbutanoyl)piperidin-4-yl)methyl)-4-oxo-4,5-dihydro-1*H*-pyrazolo[3,4-*d*]pyrimidin-1-yl)phenyl)piperazin-1-yl)pentanoic acid TFA salt (NK236)**

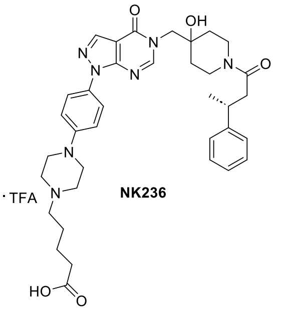

NK236

(*R*)-5-(4-(4-(5-((4-hydroxy-1-(3-phenylbutanoyl)piperidin-4-yl)methyl)-4-oxo-4,5-dihydro-1*H*-pyrazolo[3,4-*d*]pyrimidin-1-yl)phenyl)piperazin-1-yl)pentanoic acid TFA salt (NK236) was synthesized using the same method as described for NK235 (Scheme 3, exchanging *tert*-butyl 3-bromopropanoate for *tert*-butyl 5-bromopentanoate).

**Synthesis of (1*r*,4*r*)-4-(4-(4-(5-((4-hydroxy-1-((*R*)-3-phenylbutanoyl)piperidin-4-yl)methyl)-4-oxo-4,5-dihydro-1*H*-pyrazolo[3,4-*d*]pyrimidin-1-yl)phenyl)piperazine-1-carbonyl)cyclohexane-1-carboxylic acid (NK231)**

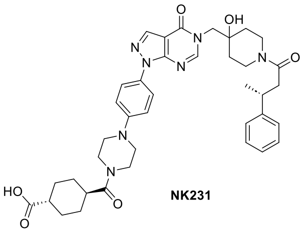

NK231

(1*r*,4*r*)-cyclohexane-1,4-dicarboxylic acid (19.8 mg, 0.11 mmol, 1.1 eq), HATU (51.7 mg, 0.14 mmol, 1.3 eq) and DIPEA (55.0 mg, 0.42 mmol, 4.0 eq) were dissolved in DMSO (1 mL) and stirred at rt for 15 min. To this was added NK210 (70.0 mg, 0.10 mmol, 1.0 eq) and the mixture was stirred at rt for 2 h. After completion, the crude product was directly purified by preparative reverse phase column chromatography (ACN/$H_2O$ + 0.1 % TFA). This yielded the product (1*r*,4*r*)-4-(4-(4-(5-((4-hydroxy-1-((*R*)-3-phenylbutanoyl)piperidin-4-yl)methyl)-4-oxo-4,5-dihydro-1*H*-pyrazolo[3,4-*d*]pyrimidin-1-yl)phenyl)-piperazine-1-carbonyl)cyclohexane-1-carboxylic acid (NK231) (41.8 mg, 0.06 mmol, 56 % yield) as a white powder.

HRMS (m/z) $[M+H]^+$ calculated for $C_{39}H_{48}N_7O_6^+$: 710.3661, found: 710.3653.

$^1$H NMR (700 MHz, DMSO-*d*$_6$) δ (ppm): 8.29 (d, *J* = 1.1 Hz, 1H), 8.28 (d, *J* = 16.5 Hz, 1H), 7.83 (d, *J* = 9.1 Hz, 2H), 7.28 – 7.24 (m, 4H), 7.17 – 7.12 (m, 3H), 4.08 – 3.92 (m, 3H), 3.70 – 3.60 (m, 5H), 3.27 – 3.14 (m, 6H), 2.90 – 2.83 (m, 1H), 2.67 – 2.52 (m, 3H), 2.18 (ddt, *J* = 11.5, 7.1, 3.8 Hz, 1H), 1.97 – 1.89 (m, 2H), 1.76 – 1.71 (m, 2H), 1.45 – 1.22 (m, 8H), 1.20 (dd, *J* = 7.0, 1.5 Hz, 3H).

$^{13}$C NMR (176 MHz, DMSO-*d*$_6$) δ (ppm): 176.54, 173.18, 169.11, 156.91 (“d”, *J* = 9.5 Hz), 151.90 (“d”, *J* = 4.2 Hz), 150.56, 149.62, 146.63 (“d”, *J* = 19.2 Hz), 135.60, 129.98, 128.20 (“d”, *J* = 5.7 Hz), 126.89 (“d”, *J* = 4.6 Hz), 125.94 (“d”, *J* = 6.6 Hz), 122.84 (“d”, *J* = 1.8 Hz), 115.78, 106.26, 69.11 (“d”, *J* = 8.3 Hz), 52.99, 48.45 (“d”, *J* = 107.6 Hz), 44.46, 42.00, 41.00 (“d”, *J* = 19.5 Hz), 40.78, 40.23 (“d”, *J* = 7.1 Hz), 38.31, 36.92, 36.11 (“d”, *J* = 36.3 Hz), 34.89 (“d”, *J* = 19.6 Hz), 34.17 (“d”, *J* = 23.6 Hz), 27.98 (“d”, *J* = 69.4 Hz), 21.98 (“d”, *J* = 34.7 Hz).

**Synthesis of *tert*-butyl (*R*)-(7-(4-(4-(5-((4-hydroxy-1-(3-phenylbutanoyl)piperidin-4-yl)methyl)-4-oxo-4,5-dihydro-1*H*-pyrazolo[3,4-*d*]pyrimidin-1-yl)phenyl)piperazin-1-yl)-7-oxoheptyl)carbamate (NK267)**

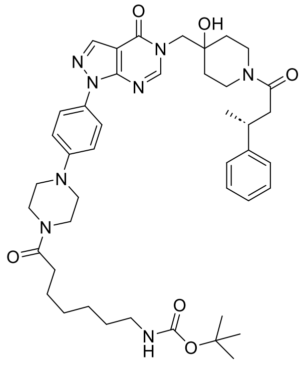

NK267

7-((*tert*-Butoxycarbonyl)amino)heptanoic acid (4.7 mg, 0.02 mmol, 1.3 eq), DIPEA (7.7 mg, 0.06 mmol, 4 eq) and HATU (7.4 mg, 0.02 mmol, 1.3 eq) were dissolved in DMSO (1 mL) and stirred for 15 min at rt. To this was added NK210 (10.0 mg, 0.01 mmol, 1 eq) and the mixture was stirred for 2 h at rt. After completion, the crude mixture was directly purified by preparative reverse phase column chromatography (ACN/$H_2O$ + 0.1 % TFA) which yielded the product *tert*-butyl (*R*)-(7-(4-(4-(5-((4-hydroxy-1-(3-phenylbutanoyl)piperidin-4-yl)methyl)-4-oxo-4,5-dihydro-1*H*-pyrazolo[3,4-*d*]pyrimidin-1-yl)phenyl)piperazin-1-yl)-7-oxoheptyl)carbamate (NK267) (10.2 mg, 0.01 mmol, 87 % yield) as a white solid.

HRMS (m/z) $[M+H]^+$ calculated for $C_{43}H_{59}N_8O_6^+$: 783.4552, found: 783.4570.

$^1$H NMR (500 MHz, DMSO-*d*$_6$) δ (ppm): 8.29 (s, 1H), 8.27 (d, *J* = 11.7 Hz, 1H), 7.82 (d, *J* = 9.1 Hz, 2H), 7.29 – 7.24 (m, 4H), 7.19 – 7.09 (m, 3H), 6.76 (t, *J* = 5.8 Hz, 1H), 4.90 (s, 1H), 4.09 – 3.91 (m, 3H), 3.71 – 3.58 (m, 5H), 3.26 – 3.12 (m, 6H), 2.89 (p, *J* = 5.8, 5.4 Hz, 3H), 2.66 – 2.52 (m, 2H), 2.35 (t, *J* = 7.5 Hz, 2H), 1.50 (p, *J* = 7.7 Hz, 4H), 1.37 (s, 9H), 1.35 – 1.23 (m, 8H), 1.20 (d, *J* = 6.9 Hz, 3H).

$^{13}$C NMR (126 MHz, DMSO-*d*$_6$) δ (ppm): 171.19, 169.58, 157.39 ("d", *J* = 6.8 Hz), 156.05, 152.37 ("d", *J* = 2.4 Hz), 151.03 ("d", *J* = 1.3 Hz), 150.16, 147.11 ("d", *J* = 13.7 Hz), 136.07, 130.37, 128.68 ("d", *J* = 4.1 Hz), 127.37 ("d", *J* = 3.4 Hz), 126.42 ("d", *J* = 4.6 Hz), 123.32, 116.19, 106.73, 77.76, 69.59 ("d", *J* = 5.3 Hz), 53.47, 48.71 ("d", *J* = 51.8 Hz), 45.09, 41.47 ("d", *J* = 13.3 Hz), 41.16, 40.73, 37.39, 36.59 (""d, *J* = 25.8 Hz), 35.36 ("d", *J* = 14.6 Hz), 34.64 ("d", *J* = 15.2 Hz), 32.68, 29.87, 28.96, 28.75, 26.61, 25.23, 22.46 ("d", *J* = 24.7 Hz).

**Synthesis of *N*-(2-(2-(2-(3-(4-(4-(5-((4-hydroxy-1-((*R*)-3-phenylbutanoyl)piperidin-4-yl)methyl)-4-oxo-4,5-dihydro-1*H*-pyrazolo[3,4-*d*]pyrimidin-1-yl)phenyl)piperazin-1-yl)-3-oxopropoxy)ethoxy)-ethoxy)ethyl)-5-((3*aS*,4*S*,6*aR*)-2-oxohexahydro-1*H*-thieno[3,4-*d*]imidazol-4-yl)pentanamide (NK264)**

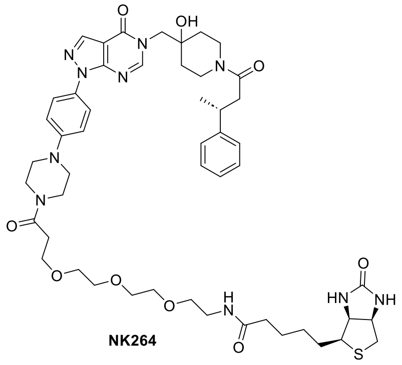

NK264

Step1:

2,2-dimethyl-4-oxo-3,8,11,14-tetraoxa-5-azaheptadecan-17-oic acid (8.6 mg, 0.03 mmol, 1.2 eq), DIPEA (11.6 mg, 0.09 mmol, 4 eq) and HATU (11.1 mg, 0.03 mmol, 1.3 eq) were dissolved in DMSO (1 mL) and stirred for 15 min at rt. To this was added NK210 (15.0 mg, 0.02 mmol, 1 eq) and the reaction was stirred for 2 h at rt. The crude mixture was directly purified by preparative reverse phase column chromatography (ACN/$H_2O$ + 0.1 % TFA). The fractions containing the product *tert*-butyl (*R*)-(2-(2-(2-(3-(4-(4-(5-((4-hydroxy-1-(3-phenylbutanoyl)piperidin-4-yl)methyl)-4-oxo-4,5-dihydro-1*H*-pyrazolo[3,4-*d*]pyrimidin-1-yl)phenyl)piperazin-1-yl)-3-oxopropoxy)ethoxy)ethoxy)ethyl)carbamate (NK262) were combined and most of the solvents were evaporated under reduced pressure. The product was then directly used for the next step without further purification.

Step 2:

The combined fractions from the previous step that contained product NK262 were diluted with HCl/dioxane (4 M, 5 mL) and stirred at rt for 4 h. After completion, the solvents were evaporated under reduced pressure which yielded the crude product (*R*)-1-(4-(4-(3-(2-(2-(2-aminoethoxy)ethoxy)ethoxy)-propanoyl)piperazin-1-yl)phenyl)-5-((4-hydroxy-1-(3-phenylbutanoyl)piperidin-4-yl)methyl)-1,5-dihydro-4*H*-pyrazolo[3,4-*d*]pyrimidin-4-one HCl salt (NK263) which was directly used for the next step.

Step 3:

NK263 (15.0 mg, 0.02 mmol, 1 eq) was dissolved in DMSO (1 mL). To this was added DIPEA (12.2 mg, 0.09 mmol, 5 eq) and 2,5-dioxopyrrolidin-1-yl-2,5-dioxopyrrolidin-1-yl-5-((3*aS*,4*S*,6*aR*)-2-oxohexahydro-1*H*-thieno[3,4-*d*]imidazol-4-yl)pentanoate (NHS-biotin, 6.4 mg, 0.02 mmol, 1 eq) and the reaction was stirred for 1 h at rt. The crude mixture was directly purified by preparative reverse phase column chromatography (ACN/$H_2O$ + 0.1 % TFA) which yielded *N*-(2-(2-(2-(3-(4-(4-(5-((4-hydroxy-1-((*R*)-3-phenylbutanoyl)piperidin-4-yl)methyl)-4-oxo-4,5-dihydro-1*H*-pyrazolo[3,4-*d*]pyrimidin-1-yl)phenyl)piperazin-1-yl)-3-oxopropoxy)ethoxy)ethoxy)ethyl)-5-((3*aS*,4*S*,6*aR*)-2-oxohexahydro-1*H*-thieno[3,4-*d*]imidazol-4-yl)pentanamide (NK264) (11.0 mg, 0.01 mmol, 59 % yield over three steps) as a white solid.

HRMS (m/z) $[M+Na]^+$ calculated for $C_{50}H_{68}N_{10}NaO_9S^+$: 1007.4784, found: 1007.4753.

$^1$H NMR (600 MHz, DMSO-$d_6$) δ (ppm): 8.31 – 8.26 (m, 2H), 7.85 – 7.80 (m, 3H), 7.26 (dd, *J* = 8.7, 6.2 Hz, 4H), 7.17 – 7.11 (m, 3H), 6.41 (s, 2H), 4.30 – 4.28 (m, 2H), 4.12 (dd, *J* = 7.7, 4.4 Hz, 2H), 4.07 – 4.03 (m, 3H), 3.98 (d, *J* = 13.9 Hz, 1H), 3.94 (d, *J* = 3.3 Hz, 2H), 3.67 – 3.63 (m, 6H), 3.51 (s, 8H), 3.38 (t, *J* = 5.9 Hz, 2H), 3.25 (t, *J* = 5.1 Hz, 2H), 3.18 (dt, *J* = 11.9, 5.9 Hz, 4H), 3.08 (ddd, *J* = 8.6, 6.2, 4.4 Hz, 1H), 2.89 – 2.84 (m, 1H), 2.81 (dd, *J* = 12.4, 5.1 Hz, 1H), 2.65 – 2.54 (m, 4H), 2.05 (t, *J* = 7.4 Hz, 2H), 1.62 – 1.25 (m, 10H), 1.20 (dd, *J* = 6.9, 1.2 Hz, 3H).

$^{13}$C NMR (151 MHz, DMSO-$d_6$) δ (ppm): 172.10, 169.10 ("d", *J* = 2.9 Hz), 168.91, 162.68, 156.91 ("d", *J* = 7.9 Hz), 151.91 ("d", *J* = 3.3 Hz), 150.55 ("d", *J* = 2.2 Hz), 149.58, 146.63 ("d", *J* = 16.7 Hz), 135.60, 129.93, 128.20 ("d", *J* = 5.0 Hz), 126.88, 125.94 ("d", *J* = 5.6 Hz), 122.85, 115.73, 106.25, 69.78, 69.72, 69.69, 69.56, 69.16, 69.09, 66.80, 61.02, 59.18, 55.41, 53.00 – 52.95 (m), 48.43, 47.97, 44.68, 41.00 ("d", *J* = 16.9 Hz), 40.69, 40.23 ("d", *J* = 5.8 Hz), 38.44, 36.92, 36.11 ("d", *J* = 31.2 Hz), 35.08, 34.89 ("d", *J* = 16.9 Hz), 34.17 ("d", *J* = 19.9 Hz), 32.78, 28.18, 28.03, 25.25, 21.98 ("d", *J* = 29.9 Hz).

#### 3.4 Synthesis of acid-functionalized VHL ligands

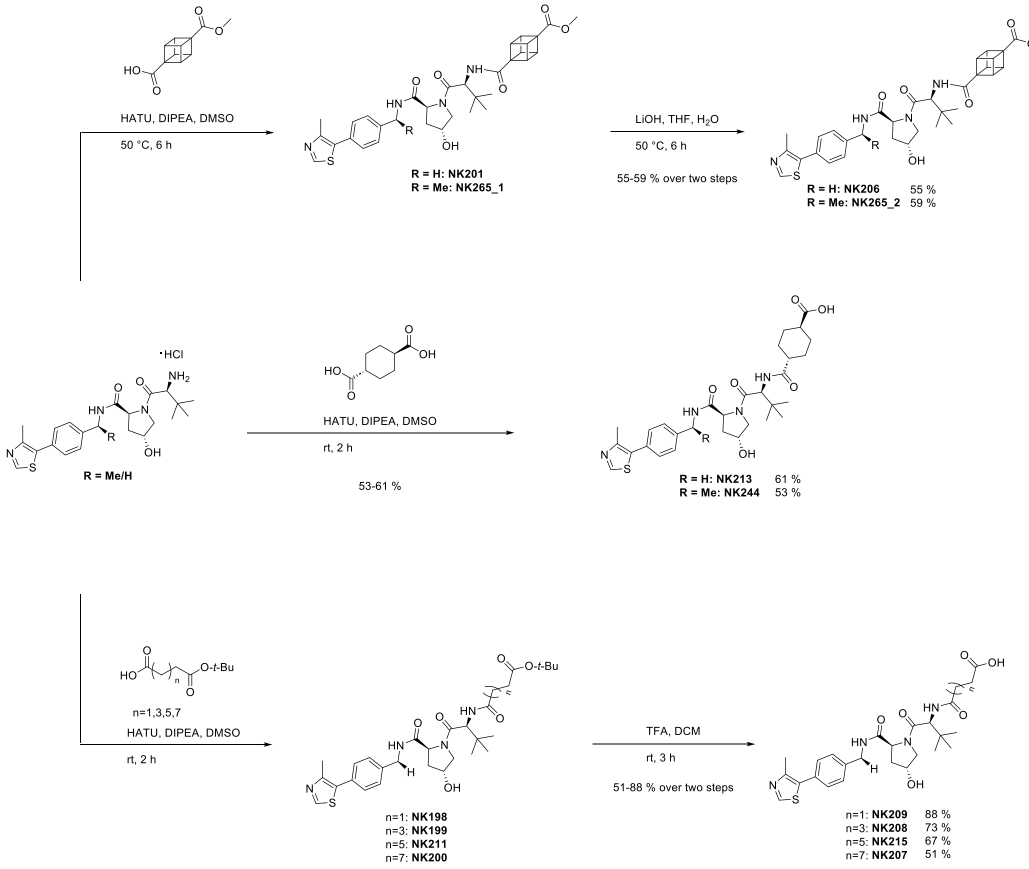

**Scheme 4.** Synthesis of acid-functionalized VHL-ligands NK206, NK265_2, NK213, NK244, NK209, NK208, NK215 and NK207.

##### Synthesis of (1*R*,2*R*,3*S*,8*S*)-4-(((*S*)-1-((2*S*,4*R*)-4-hydroxy-2-((4-(4-methylthiazol-5-yl)benzyl)carbamoyl)pyrrolidin-1-yl)-3,3-dimethyl-1-oxobutan-2-yl)carbamoyl)cubane-1-carboxylic acid (NK206)

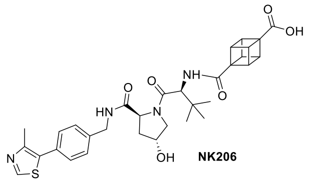

Step 1:

(2*r*,3*R*,4*s*,5*S*)-4-(Methoxycarbonyl)cubane-1-carboxylic acid (31.8 mg, 0.15 mmol, 1.2 eq), DIPEA (66.4 mg, 0.51 mmol, 4.0 eq) and HATU (63.5 mg, 0.17 mmol, 1.3 eq) were dissolved in DMSO (1 mL) and stirred for 15 min at 50 °C. To this was added (2*S*,4*R*)-1-((*S*)-2-amino-3,3-dimethylbutanoyl)-4-hydroxy-*N*-(4-(4-methylthiazol-5-yl)benzyl)pyrrolidine-2-carboxamide (60.0 mg, 0.13 mmol, 1.0 eq) and the reaction was stirred for 6 h at 50 °C. After completion, the crude product was cooled to rt and then directly purified by reverse phase flash column chromatography (ACN/$H_2O$ + 0.1 % TFA). After evaporation of the solvents by reduced pressure, the product methyl (1*R*,2*R*,3*S*,8*S*)-4-(((*S*)-1-((2*S*,4*R*)-4-hydroxy-2-((4-(4-methylthiazol-5-yl)benzyl)carbamoyl)pyrrolidin-1-yl)-3,3-dimethyl-1-oxobutan-2-yl)carbamoyl)cubane-1-carboxylate (NK201) was directly used for the next step without further purification.

Step 2:

(1*R*,2*R*,3*S*,8*S*)-4-(((*S*)-1-((2*S*,4*R*)-4-Hydroxy-2-((4-(4-methylthiazol-5-yl)benzyl)carbamoyl)pyrrolidin-1-yl)-3,3-dimethyl-1-oxobutan-2-yl)carbamoyl)cubane-1-carboxylate (NK201) (43.7 mg, 0.07 mmol, 1 eq) was dissolved in THF (0.2 mL) to which was added LiOH in $H_2O$ (2 M, 106 µL, 3 eq) and the reaction was stirred for 6 h at 50 °C. The reaction was quenched by the addition of 1 M HCl until a pH of 2-3 was achieved. After evaporation of the organic solvents by reduced pressure, the mixture was diluted with $H_2O$ (10 mL) and extracted with EA (3 x 20 mL). The combined organic layers were washed with brine

(1x 20 mL), dried with $MgSO_4$, filtered and concentrated under reduced pressure which yielded the product (1*R*,2*R*,3*S*,8*S*)-4-(((*S*)-1-((2*S*,4*R*)-4-hydroxy-2-((4-(4-methylthiazol-5-yl)benzyl)carbamoyl)-pyrrolidin-1-yl)-3,3-dimethyl-1-oxobutan-2-yl)carbamoyl)cubane-1-carboxylic acid (NK206) (42.4 mg, 0.07 mmol, 55 % yield) as an off-white powder.

HRMS (m/z) $[M+Na]^+$ calculated for $C_{32}H_{36}N_4NaO_6S^+$: 627.2248, found: 627.2246.

$^1$H NMR (700 MHz, DMSO-$d_6$) δ (ppm): 9.01 (s, 1H), 8.59 (t, *J* = 6.1 Hz, 1H), 7.68 (d, *J* = 9.4 Hz, 1H), 7.44 – 7.35 (m, 4H), 4.59 (d, *J* = 9.3 Hz, 1H), 4.42 (dt, *J* = 15.7, 7.1 Hz, 2H), 4.34 (tt, *J* = 4.4, 2.4 Hz, 1H), 4.22 (dd, *J* = 15.8, 5.4 Hz, 1H), 4.11 – 4.06 (m, 6H), 3.67 – 3.61 (m, 2H), 2.44 (s, 3H), 2.04 (dddd, *J* = 10.3, 7.6, 4.1, 1.9 Hz, 1H), 1.89 (ddd, *J* = 12.9, 8.7, 4.6 Hz, 1H), 0.94 (s, 9H).

$^{13}$C NMR (176 MHz, DMSO-$d_6$) δ (ppm): 172.63, 171.92, 170.19, 169.45, 151.59, 147.53, 139.55, 131.26, 129.56, 128.66, 127.45, 68.84, 58.74, 57.02, 56.41, 56.15, 55.14, 46.38, 45.98, 41.66, 37.94, 35.50, 26.44, 15.86.

**Synthesis of (1*R*,2*R*,3*S*,8*S*)-4-(((*S*)-1-((2*S*,4*R*)-4-hydroxy-2-(((*S*)-1-(4-(4-methylthiazol-5-yl)phenyl)ethyl)carbamoyl)pyrrolidin-1-yl)-3,3-dimethyl-1-oxobutan-2-yl)carbamoyl)cubane-1-carboxylic acid (NK265_2)**

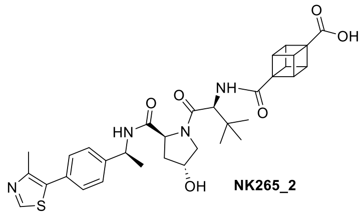

(1*R*,2*R*,3*S*,8*S*)-4-(((*S*)-1-((2*S*,4*R*)-4-hydroxy-2-(((*S*)-1-(4-(4-methylthiazol-5-yl)phenyl)ethyl)carbamoyl)pyrrolidin-1-yl)-3,3-dimethyl-1-oxobutan-2-yl)carbamoyl)cubane-1-carboxylic acid (NK265_2) was synthesized using the same procedure as described for NK206 (Scheme 4, exchanging (2*S*,4*R*)-1-((*S*)-2-amino-3,3-dimethylbutanoyl)-4-hydroxy-*N*-(4-(4-methylthiazol-5-yl)benzyl)pyrrolidine-2-carboxamide for (2*S*,4*R*)-1-((*S*)-2-amino-3,3-dimethylbutanoyl)-4-hydroxy-N-((*S*)-1-(4-(4-methylthiazol-5-yl)phenyl)ethyl)pyrrolidine-2-carboxamide).

HRMS (m/z) $[M+Na]^+$ calculated for $C_{33}H_{39}N_4O_6S^+$: 619.2585, found: 619.2586.

$^1$H NMR (500 MHz, DMSO-$d_6$) δ (ppm): 9.00 (s, 1H), 8.39 (d, *J* = 7.8 Hz, 1H), 7.57 (d, *J* = 9.3 Hz, 1H), 7.45 – 7.42 (m, 2H), 7.38 (d, *J* = 8.3 Hz, 2H), 4.98 – 4.89 (m, 1H), 4.60 – 4.53 (m, 1H), 4.43 (t, *J* = 8.1 Hz, 1H), 4.28 (dt, *J* = 4.2, 2.0 Hz, 1H), 4.14 – 4.02 (m, 6H), 3.59 (qd, *J* = 9.9, 9.1, 2.9 Hz, 2H), 2.45 (s, 3H), 2.04 – 1.94 (m, 1H), 1.79 (ddd, *J* = 13.0, 8.7, 4.6 Hz, 1H), 1.37 (d, *J* = 7.0 Hz, 3H), 0.94 (s, 9H).

$^{13}$C NMR (126 MHz, DMSO-$d_6$) δ (ppm): 172.64, 170.60, 170.11, 169.36, 151.57, 147.63, 144.65, 131.19, 129.64, 128.83, 126.41, 68.78, 58.65, 57.04, 56.37, 56.19, 55.18, 47.67, 46.35, 45.97, 37.71, 35.55, 26.50, 22.44, 15.93.

**Synthesis of (1*S*,4*r*)-4-(((*S*)-1-((2*S*,4*R*)-4-hydroxy-2-((4-(4-methylthiazol-5-yl)benzyl)carbamoyl)-pyrrolidin-1-yl)-3,3-dimethyl-1-oxobutan-2-yl)carbamoyl)cyclohexane-1-carboxylic acid (NK213)**

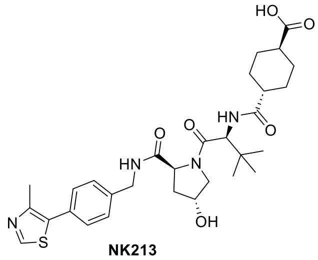

(1*r*,4*r*)-Cyclohexane-1,4-dicarboxylic acid (62.7 mg, 0.36 mmol, 1.7 eq), DIPEA (0.64 mmol, 83.0 mg, 4.0 eq) and HATU (97.7 mg, 0.26 mmol, 1.3 eq) were dissolved in DMSO (3 mL) and stirred for 15 min at rt. To this was added (2*S*,4*R*)-1-((*S*)-2-amino-3,3-dimethylbutanoyl)-4-hydroxy-N-(4-(4-methylthiazol-5-yl)benzyl)pyrrolidine-2-carboxamide (100.0 mg, 0.21 mmol, 1.0 eq) and the reaction was stirred for 2 h at rt. After completion, the crude product was directly purified by preparative reverse phase column chromatography (ACN/$H_2O$ + 0.1 % TFA). The product (1*S*,4*r*)-4-(((*S*)-1-((2*S*,4*R*)-4-hydroxy-2-((4-(4-methylthiazol-5-yl)benzyl)carbamoyl)-pyrrolidin-1-yl)-3,3-dimethyl-1-oxobutan-2-yl)carbamoyl)cyclohexane-1-carboxylic acid (NK213) (76.2 mg, 0.13 mmol, 61 % yield) was obtained as a white powder.

HRMS (m/z) $[M+Na]^+$ calculated for $C_{30}H_{40}N_4NaO_6S^+$: 607.2561, found: 607.2552.

$^{1}$H NMR (600 MHz, DMSO-$d_6$) δ (ppm): 9.00 (s, 1H), 8.56 (t, *J* = 6.1 Hz, 1H), 7.74 (d, *J* = 9.3 Hz, 1H), 7.43 – 7.35 (m, 4H), 4.51 (d, *J* = 9.4 Hz, 1H), 4.46 – 4.41 (m, 2H), 4.37 – 4.32 (m, 2H), 4.22 (dd, *J* = 15.8, 5.4 Hz, 1H), 3.68 – 3.60 (m, 2H), 2.44 (s, 3H), 2.34 (tt, *J* = 11.8, 3.5 Hz, 1H), 2.13 (ddd, *J* = 11.8, 8.3, 3.6 Hz, 1H), 2.05 – 1.99 (m, 1H), 1.89 (dq, *J* = 12.8, 4.7 Hz, 4H), 1.79 (dq, *J* = 11.6, 2.3 Hz, 1H), 1.69 – 1.65 (m, 1H), 1.39 – 1.21 (m, 4H), 0.93 (s, 9H).

$^{13}$C NMR (151 MHz, DMSO-$d_6$) δ (ppm): 176.62, 174.73, 171.97, 169.66, 151.55, 147.61, 139.55, 131.24, 129.59, 128.65, 127.44, 68.87, 58.73, 56.35, 56.11, 42.48, 41.91, 41.65, 37.93, 35.39, 29.08, 28.10, 27.87, 27.80, 26.37, 15.90.

**Synthesis of (1*S*,4*r*)-4-(((*S*)-1-((2*S*,4*R*)-4-hydroxy-2-(((*S*)-1-(4-(4-methylthiazol-5-yl)phenyl)ethyl)-carbamoyl)pyrrolidin-1-yl)-3,3-dimethyl-1-oxobutan-2-yl)carbamoyl)cyclohexane-1-carboxylic acid (NK244)**

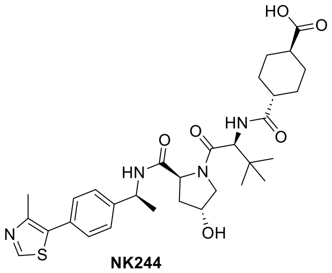

**NK244**

(1*S*,4*r*)-4-(((*S*)-1-((2*S*,4*R*)-4-Hydroxy-2-(((*S*)-1-(4-(4-methylthiazol-5-yl)phenyl)ethyl)-carbamoyl)pyrrolidin-1-yl)-3,3-dimethyl-1-oxobutan-2-yl)carbamoyl)cyclohexane-1-carboxylic acid (NK244) was synthesized as described for NK213 (Scheme 4, exchanging (2*S*,4*R*)-1-((*S*)-2-amino-3,3-dimethylbutanoyl)-4-hydroxy-*N*-(4-(4-methylthiazol-5-yl)benzyl)pyrrolidine-2-carboxamide for (2*S*,4*R*)-1-((*S*)-2-amino-3,3-dimethylbutanoyl)-4-hydroxy-N-((*S*)-1-(4-(4-methylthiazol-5-yl)phenyl)ethyl)pyrrolidine-2-carboxamide).

HRMS (m/z) $[M+H]^+$ calculated for $C_{31}H_{43}N_4O_6S^+$: 599.2898, found: 599.2896.

$^{1}$H NMR (700 MHz, DMSO-$d_6$) δ (ppm): 8.99 (s, 1H), 8.37 (d, *J* = 7.8 Hz, 1H), 7.68 (d, *J* = 9.3 Hz, 1H), 7.43 (d, *J* = 8.3 Hz, 2H), 7.38 (d, *J* = 8.3 Hz, 2H), 4.91 (p, *J* = 7.1 Hz, 1H), 4.48 (d, *J* = 9.3 Hz, 1H), 4.42 (t, *J* = 8.1 Hz, 1H), 4.28 (dt, *J* = 4.4, 2.0 Hz, 1H), 3.60 (s, 2H), 2.45 (s, 3H), 2.32 (td, *J* = 11.7, 5.9 Hz, 1H), 2.13 (ddt, *J* = 11.8, 8.3, 3.6 Hz, 1H), 2.02 – 1.99 (m, 1H), 1.91 – 1.87 (m, 2H), 1.78 (ddd, *J* = 12.6, 8.5, 4.4 Hz, 2H), 1.70 – 1.65 (m, 1H), 1.37 (d, *J* = 7.0 Hz, 4H), 1.35 – 1.25 (m, 4H), 0.93 (s, 9H).

$^{13}$C NMR (176 MHz, DMSO-$d_6$) δ (ppm): 176.76, 174.83, 170.78, 169.69, 151.66, 147.82, 144.81, 131.29, 129.79, 128.96, 126.52, 68.90, 58.71, 56.38, 56.33, 47.82, 42.64, 42.04, 37.84, 35.48, 29.14, 28.21, 28.00, 27.96, 26.56, 22.57, 16.08.

**Synthesis of 4-(((*S*)-1-((2*S*,4*R*)-4-hydroxy-2-((4-(4-methylthiazol-5-yl)benzyl)carbamoyl)-pyrrolidin-1-yl)-3,3-dimethyl-1-oxobutan-2-yl)amino)-4-oxobutanoic acid (NK209)**

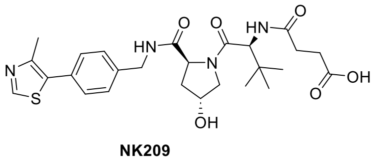

**NK209**

Step 1:

4-(*tert*-Butoxy)-4-oxobutanoic acid (26.9 mg, 0.15 mmol, 1.2 eq), DIPEA (66.4 mg, 0.51 mmol, 4.0 eq) and HATU (63.5 mg, 0.17 mmol, 1.3 eq) were dissolved in DMSO (1 mL) and stirred for 15 min at rt. To this was added (2*S*,4*R*)-1-((*S*)-2-amino-3,3-dimethylbutanoyl)-4-hydroxy-*N*-(4-(4-methylthiazol-5-yl)benzyl)pyrrolidine-2-carboxamide (60.0 mg, 0.13 mmol, 1 eq) and the reaction was stirred for 2 h at rt. After completion, the crude product was directly purified by preparative reverse phase column chromatography (ACN/$H_2O$ + 0.1 % TFA). After evaporation of the solvents under reduced pressure, the product *tert*-butyl 4-(((*S*)-1-((2*S*,4*R*)-4-hydroxy-2-((4-(4-methylthiazol-5-yl)benzyl)carbamoyl)pyrrolidin-1-yl)-3,3-dimethyl-1-oxobutan-2-yl)amino)-4-oxobutanoate (NK198) was directly used for the next step without further purification.

Step 2:

NK198 was dissolved in TFA/DCM (50:50, 2 mL) and stirred for 3 h at rt. After completion, the solvents were evaporated under reduced pressure. This yielded the product 4-(((*S*)-1-((2*S*,4*R*)-4-hydroxy-2-((4-(4-methylthiazol-5-yl)benzyl)carbamoyl)-pyrrolidin-1-yl)-3,3-dimethyl-1-oxobutan-2-yl)amino)-4-oxobutanoic acid (NK209) (60.3 mg, 0.11 mmol, 88 % yield) as a white powder.

HRMS (m/z) $[M+H]^+$ calculated for $C_{26}H_{35}N_4O_6S^+$: 531.2272, found: 531.2268.

$^1$H NMR (600 MHz, DMSO-$d_6$) δ (ppm): 9.00 (s, 1H), 8.57 (t, *J* = 6.0 Hz, 1H), 7.93 (d, *J* = 9.3 Hz, 1H), 7.43 – 7.38 (m, 4H), 4.53 (d, *J* = 9.4 Hz, 1H), 4.48 – 4.41 (m, 2H), 4.35 (dt, *J* = 4.4, 2.2 Hz, 1H), 4.22 (dd, *J* = 15.8, 5.4 Hz, 1H), 3.67 (dd, *J* = 10.5, 4.2 Hz, 1H), 3.61 (dt, *J* = 10.9, 1.7 Hz, 1H), 2.46 – 2.33 (m, 7H), 2.05 – 2.00 (m, 1H), 1.90 (ddd, *J* = 13.0, 8.6, 4.6 Hz, 1H), 0.93 (s, 9H).

$^{13}$C NMR (151 MHz, DMSO-$d_6$) δ (ppm): 173.86, 171.96, 170.88, 169.58, 151.54, 147.62, 139.55, 131.24, 129.60, 128.65, 127.44, 68.88, 58.72, 56.45, 56.36, 41.66, 37.93, 35.38, 29.74, 29.25, 26.37, 15.90.

**Synthesis of 6-(((*S*)-1-((2*S*,4*R*)-4-hydroxy-2-((4-(4-methylthiazol-5-yl)benzyl)carbamoyl)-pyrrolidin-1-yl)-3,3-dimethyl-1-oxobutan-2-yl)amino)-6-oxohexanoic acid (NK208)**

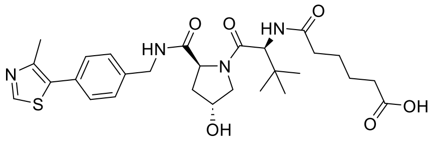

**NK208**

6-(((*S*)-1-((2*S*,4*R*)-4-hydroxy-2-((4-(4-methylthiazol-5-yl)benzyl)carbamoyl)-pyrrolidin-1-yl)-3,3-dimethyl-1-oxobutan-2-yl)amino)-6-oxohexanoic acid (NK208) was synthesized as described for NK209 (Scheme 4, exchanging 4-(*tert*-butoxy)-4-oxobutanoic acid for 6-(*tert*-butoxy)-6-oxohexanoic acid.

HRMS (m/z) $[M+H]^+$ calculated for $C_{28}H_{39}N_4O_6S^+$: 559.2585, found: 559.2579.

$^1$H NMR (700 MHz, DMSO-$d_6$) δ (ppm): 8.99 (s, 1H), 8.56 (t, *J* = 6.1 Hz, 1H), 7.86 (d, *J* = 9.4 Hz, 1H), 7.44 – 7.36 (m, 4H), 4.54 (d, *J* = 9.4 Hz, 1H), 4.45 – 4.40 (m, 2H), 4.35 (dt, *J* = 4.4, 2.0 Hz, 1H), 4.21 (dd, *J* = 15.8, 5.4 Hz, 1H), 3.69 – 3.61 (m, 2H), 2.44 (s, 3H), 2.26 (dt, *J* = 14.0, 7.2 Hz, 1H), 2.20 (td, *J* = 5.6, 4.5, 2.0 Hz, 2H), 2.14 – 2.10 (m, 1H), 2.03 (dddd, *J* = 12.3, 7.6, 2.8, 1.4 Hz, 1H), 1.90 (td, *J* = 8.4, 4.3 Hz, 1H), 1.54 – 1.44 (m, 4H), 0.93 (s, 9H).

$^{13}$C NMR (176 MHz, DMSO-$d_6$) δ (ppm): 174.39, 171.95, 171.87, 169.70, 151.51, 147.63, 139.54, 131.21, 129.60, 128.64, 127.43, 68.86, 58.69, 56.35, 56.31, 41.65, 37.93, 35.20, 34.56, 33.39, 26.38, 24.98, 24.16, 15.91.

**Synthesis of 8-(((*S*)-1-((2*S*,4*R*)-4-hydroxy-2-((4-(4-methylthiazol-5-yl)benzyl)carbamoyl)-pyrrolidin-1-yl)-3,3-dimethyl-1-oxobutan-2-yl)amino)-8-oxooctanoic acid (NK215)**

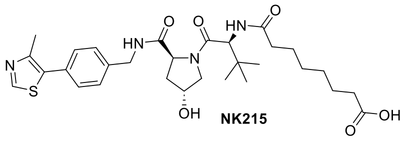

**NK215**

8-(((S)-1-((2*S*,4*R*)-4-hydroxy-2-((4-(4-methylthiazol-5-yl)benzyl)carbamoyl)-pyrrolidin-1-yl)-3,3-dimethyl-1-oxobutan-2-yl)amino)-8-oxooctanoic acid (NK215) was synthesized as described for NK209 (Scheme 4, exchanging 4-(*tert*-butoxy)-4-oxobutanoic acid for 8-(*tert*-butoxy)-8-oxooctanoic acid).

HRMS (m/z) $[M+H]^+$ calculated for $C_{30}H_{43}N_4O_6S^+$: 587.2898, found: 587.2892.

$^1$H NMR (700 MHz, DMSO-$d_6$) δ (ppm): 9.00 (s, 1H), 8.55 (t, *J* = 6.1 Hz, 1H), 7.84 (d, *J* = 9.4 Hz, 1H), 7.45 – 7.36 (m, 4H), 4.54 (d, *J* = 9.4 Hz, 1H), 4.45 – 4.41 (m, 2H), 4.35 (dt, *J* = 4.3, 2.0 Hz, 1H), 4.21 (dd, *J* = 15.8, 5.4 Hz, 1H), 3.73 – 3.61 (m, 2H), 2.44 (s, 3H), 2.25 (ddd, *J* = 14.1, 8.3, 7.0 Hz, 1H), 2.18 (td, *J* = 7.4, 5.1 Hz, 4H), 2.14 – 2.08 (m, 1H), 1.47 (d, *J* = 7.2 Hz, 4H), 1.25 (ddt, *J* = 11.5, 7.4, 4.2 Hz, 4H), 0.93 (s, 9H).

$^{13}$C NMR (176 MHz, DMSO-$d_6$) δ (ppm): 174.44, 172.06, 171.95, 169.71, 151.51, 147.62, 139.54, 131.21, 129.59, 128.64, 127.42, 68.85, 58.68, 56.33, 56.27, 41.64, 37.93, 35.20, 34.83, 33.57, 28.38, 28.22, 26.38, 25.30, 24.32, 15.90.

**Synthesis of 10-(((*S*)-1-((2*S*,4*R*)-4-hydroxy-2-((4-(4-methylthiazol-5-yl)benzyl)carbamoyl)-pyrrolidin-1-yl)-3,3-dimethyl-1-oxobutan-2-yl)amino)-10-oxodecanoic acid (NK207)**

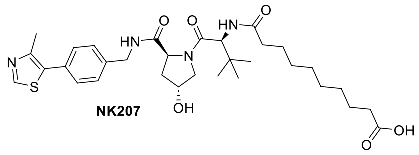

10-(((*S*)-1-((2*S*,4*R*)-4-hydroxy-2-((4-(4-methylthiazol-5-yl)benzyl)carbamoyl)-pyrrolidin-1-yl)-3,3-dimethyl-1-oxobutan-2-yl)amino)-10-oxodecanoic acid (NK207) was synthesized as described for NK209 (Scheme 4, exchanging 4-(*tert*-butoxy)-4-oxobutanoic acid for 10-(*tert*-butoxy)-10-oxodecanoic acid.

HRMS (m/z) $[M+H]^+$ calculated for $C_{32}H_{47}N_4O_6S^+$: 615.3211, found: 615.3201.

$^1$H NMR (600 MHz, DMSO-$d_6$) δ (ppm): 8.99 (s, 1H), 8.56 (t, $J$ = 6.1 Hz, 1H), 7.83 (d, $J$ = 9.3 Hz, 1H), 7.43 – 7.34 (m, 4H), 4.54 (d, $J$ = 9.4 Hz, 1H), 4.45 – 4.41 (m, 2H), 4.35 (dt, $J$ = 4.4, 2.1 Hz, 1H), 4.21 (dd, $J$ = 15.8, 5.3 Hz, 1H), 3.68 – 3.62 (m, 2H), 2.44 (s, 3H), 2.25 (ddd, $J$ = 14.1, 8.2, 7.1 Hz, 1H), 2.18 (t, $J$ = 7.4 Hz, 2H), 2.10 (ddd, $J$ = 14.2, 8.2, 6.2 Hz, 1H), 2.03 (dddd, $J$ = 11.8, 7.6, 3.0, 1.7 Hz, 1H), 1.90 (ddd, $J$ = 12.9, 8.6, 4.6 Hz, 1H), 1.47 (dq, $J$ = 14.7, 7.3 Hz, 4H), 1.26 – 1.21 (m, 10H), 0.93 (s, 9H).

$^{13}$C NMR (151 MHz, DMSO-$d_6$) δ (ppm): 174.49, 172.10, 171.95, 169.72, 162.31, 147.65, 139.54, 131.21, 129.60, 128.64, 127.43, 68.86, 58.69, 56.34, 56.27, 41.64, 40.43, 37.95, 35.21, 34.85, 33.67, 28.68, 28.63, 28.54, 26.38, 25.42, 24.48, 15.92.

#### 3.5 Synthesis of an amine-functionalized VHL ligand

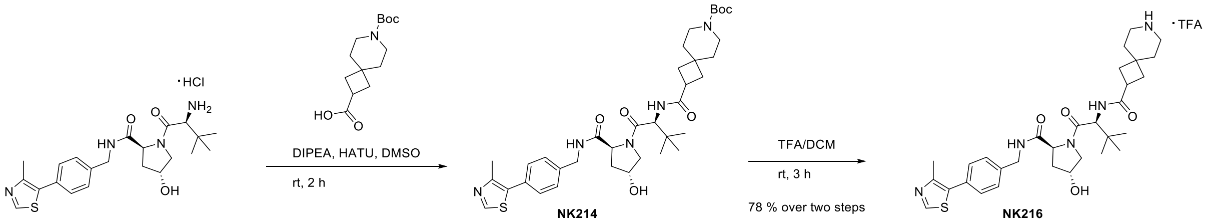

**Scheme 5.** Synthesis of amine-functionalized VHL ligand NK216.

##### Synthesis of *N*-((*S*)-1-((2*S*,4*R*)-4-hydroxy-2-((4-(4-methylthiazol-5-yl)benzyl)carbamoyl)pyrrolidin-1-yl)-3,3-dimethyl-1-oxobutan-2-yl)-7-azaspiro[3.5]nonane-2-carboxamide TFA salt (NK216)

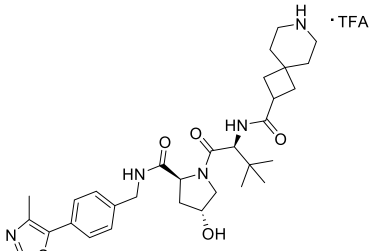

NK216

Step 1:

7-(*tert*-Butoxycarbonyl)-7-azaspiro[3.5]nonane-2-carboxylic acid (48.4 mg, 0.18 mmol, 1.2 eq), DIPEA (0.60 mmol, 77.486 mg, 4.0 eq) and HATU (74.09 mg, 0.19 mmol, 1.3 eq) were dissolved in DMSO (1 mL) and stirred for 15 min at rt. To this was added (2*S*,4*R*)-1-((*S*)-2-amino-3,3-dimethylbutanoyl)-4-hydroxy-*N*-(4-(4-methylthiazol-5-yl)benzyl)pyrrolidine-2-carboxamide (70.0 mg, 0.15 mmol, 1.0 eq) and the reaction was stirred at rt for 2 h. After completion, the crude product was directly purified by preparative reverse phase column chromatography (ACN/$H_2O$ + 0.1 % TFA). The product *tert*-butyl 2-(((*S*)-1-((2*S*,4*R*)-4-hydroxy-2-((4-(4-methylthiazol-5-yl)benzyl)carbamoyl)pyrrolidin-1-yl)-3,3-dimethyl-1-oxobutan-2-yl)carbamoyl)-7-azaspiro[3.5]nonane-7-carboxylate (NK214) was directly used for the next step.

Step 2:

NK214 was dissolved in TFA/DCM (50:50, 2 mL) and stirred for 3 h at rt. After completion, the solvents were evaporated under reduced pressure. This yielded the product *N*-((*S*)-1-((2*S*,4*R*)-4-hydroxy-2-((4-(4-methylthiazol-5-yl)benzyl)carbamoyl)pyrrolidin-1-yl)-3,3-dimethyl-1-oxobutan-2-yl)-7-azaspiro[3.5]nonane-2-carboxamide TFA salt (NK216) (76.9 mg, 0.11 mmol, 75 % yield) as a white powder.

HRMS (m/z) $[M+H]^+$ calculated for $C_{31}H_{44}N_5O_4S^+$: 582.3109, found: 582.3096.

$^1H$ NMR (700 MHz, DMSO-$d_6$) δ (ppm): 8.99 (s, 1H), 8.55 (t, *J* = 6.1 Hz, 1H), 8.29 (s, 2H), 7.74 (d, *J* = 9.3 Hz, 1H), 7.43 – 7.37 (m, 4H), 4.53 (d, *J* = 9.3 Hz, 1H), 4.42 (dt, *J* = 16.1, 7.5 Hz, 2H), 4.35 (dt, *J* = 4.4, 2.0 Hz, 1H), 4.21 (dd, *J* = 15.8, 5.4 Hz, 1H), 3.69 – 3.62 (m, 2H), 3.25 – 3.15 (m, 1H), 3.02 – 2.98 (m, 2H), 2.91 (s, 2H), 2.44 (s, 3H), 2.05 – 1.94 (m, 3H), 1.92 – 1.82 (m, 3H), 1.70 (t, *J* = 5.9 Hz, 2H), 1.64 (t, *J* = 5.9 Hz, 2H), 0.92 (s, 9H).

$^{13}C$ NMR (176 MHz, DMSO-$d_6$) δ (ppm): 173.90, 171.91, 169.56, 151.49, 147.68, 139.51, 131.18, 129.62, 128.64, 127.42, 68.87, 58.70, 56.45, 56.36, 41.64, 40.55, 40.43, 40.30, 37.98, 35.38, 34.58, 34.08, 32.70, 32.35, 31.34, 26.33, 15.92.

#### 3.6 Synthesis of PROTACs from acid functionalized USP7 ligands

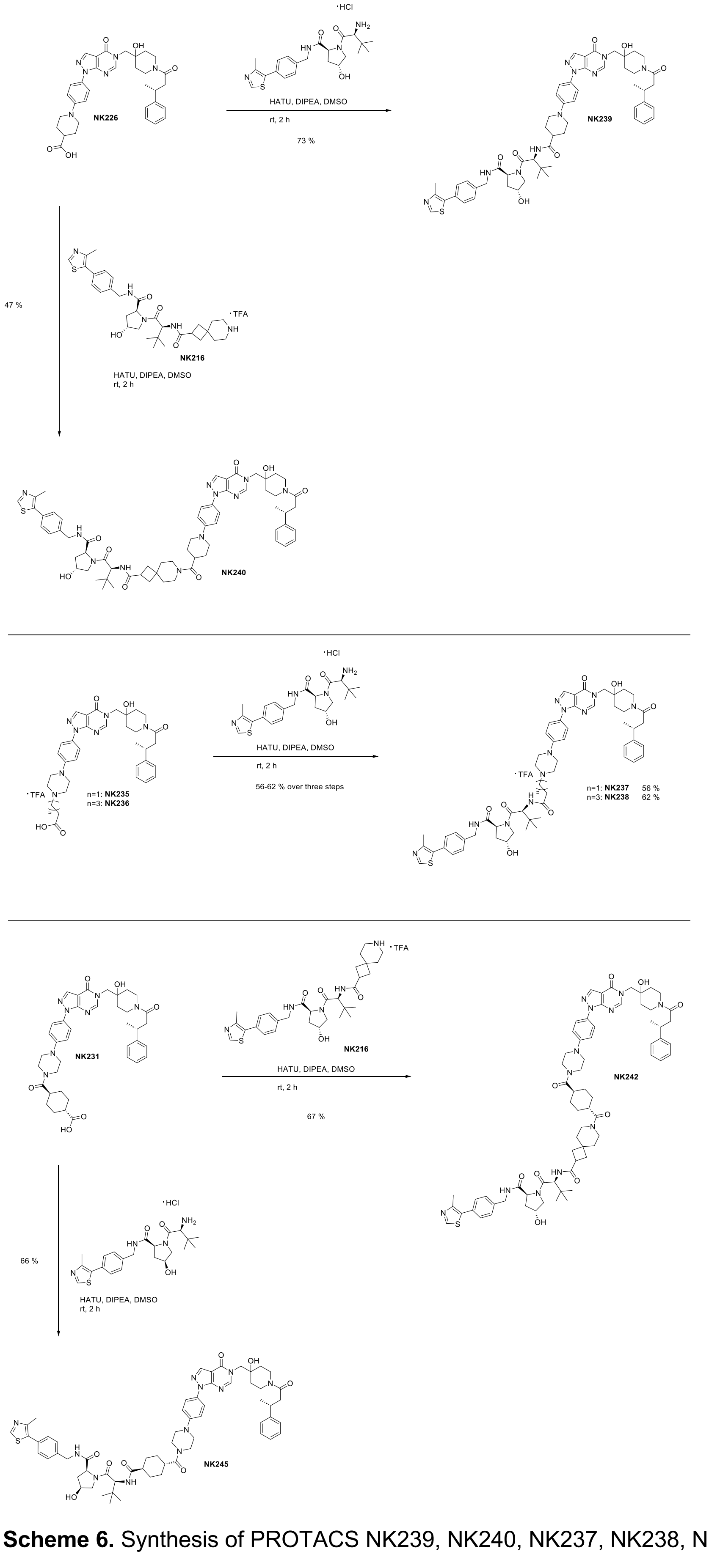

**Scheme 6.** Synthesis of PROTACS NK239, NK240, NK237, NK238, NK242 and of control NK245.

**Synthesis of 1-(4-(5-((4-hydroxy-1-((*R*)-3-phenylbutanoyl)piperidin-4-yl)methyl)-4-oxo-4,5-dihydro-1*H*-pyrazolo[3,4-*d*]pyrimidin-1-yl)phenyl)-*N*-((*S*)-1-((2*S*,4*R*)-4-hydroxy-2-((4-(4-methylthiazol-5-yl)benzyl)carbamoyl)pyrrolidin-1-yl)-3,3-dimethyl-1-oxobutan-2-yl)piperidine-4-carboxamide (NK239)**

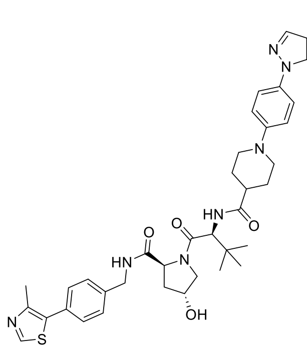

NK239

NK226 (9.9 mg, 0.02 mmol, 1.1 eq), HATU (7.4 mg, 0.02 mmol, 1.3 eq) and DIPEA (9.7 mg, 0,07 mmol, 5 eq) were dissolved in DMSO (1 mL) and stirred at rt for 15 min. To this was added (2*S*,4*R*)-1-((*S*)-2-amino-3,3-dimethyl-butanoyl)-4-hydroxy-*N*-(4-(4-methylthiazol-5-yl)benzyl)-pyrrolidine-2-carboxamide (7.0 mg, 0.01 mmol, 1 eq) and the mixture was stirred at rt for 2 h. After completion, the crude product was purified by preparative reverse phase column chromatography (ACN/$H_2O$ + 0.1 % TFA). After evaporation of the solvents under reduced pressure, the product 1-(4-(5-((4-hydroxy-1-((*R*)-3-phenylbutanoyl)-piperidin-4-yl)methyl)-4-oxo-4,5-dihydro-1*H*-pyrazolo[3,4-*d*]pyrimidin-1-yl)phenyl)-*N*-((*S*)-1-((2*S*,4*R*)-4-hydroxy-2-((4-(4-methylthiazol-5-yl)benzyl)carbamoyl)pyrrolidin-1-yl)-3,3-dimethyl-1-oxobutan-2-yl)piperidine-4-carboxamide (NK239) (11.1 mg, 0.01 mmol, 73 % yield) was obtained as a white powder.

HRMS (m/z) $[M+2H]^{2+}$ calculated for $C_{55}H_{68}N_{10}O_7S^{2+}$: 506.2491, found: 506.2477.

$^1$H NMR (600 MHz, DMSO-*d*$_6$) δ (ppm): 9.00 (s, 1H), 8.57 (t, *J* = 6.1 Hz, 1H), 8.30 (s, 1H), 8.28 (d, *J* = 14.1 Hz, 1H), 7.93 (d, *J* = 9.3 Hz, 1H), 7.83 (d, *J* = 8.6 Hz, 2H), 7.44 – 7.38 (m, 5H), 7.26 (dd, *J* = 8.8, 6.2 Hz, 5H), 7.21 – 7.14 (m, 3H), 4.55 (d, *J* = 9.4 Hz, 1H), 4.46 – 4.41 (m, 2H), 4.35 (dq, *J* = 4.4, 2.2 Hz, 1H), 4.22 (dd, *J* = 15.8, 5.5 Hz, 1H), 4.07 – 3.93 (m, 3H), 3.83 – 3.79 (m, 2H), 3.69 – 3.62 (m, 3H), 3.25 – 3.13 (m, 2H), 2.90 – 2.81 (m, 3H), 2.67 – 2.57 (m, 2H), 2.45 (s, 3H), 2.06 – 2.02 (m, 1H), 1.93 – 1.84 (m, 2H), 1.76 – 1.67 (m, 3H), 1.56 – 1.23 (m, 5H), 1.20 (dd, *J* = 7.0, 1.2 Hz, 3H), 0.95 (s, 9H).

$^{13}$C NMR (151 MHz, DMSO-*d*$_6$) δ (ppm): 173.96, 171.96, 169.60, 169.11 (“d”, *J* = 2.9 Hz), 156.91 (“d”, *J* = 8.1 Hz), 151.93 (“d”), 151.51, 150.60, 147.66, 146.64 (“d”, *J* = 16.6 Hz), 139.53, 135.66, 131.20, 129.62, 128.87 (“d”, *J* = 3.4 Hz), 128.65, 128.21 (“d”, *J* = 5.0 Hz), 128.04, 127.44, 126.90 (“d”, *J* = 3.9 Hz), 125.94 (“d”, *J* = 6.1 Hz), 122.87, 120.74, 106.29, 69.12 (“d”, *J* = 7.3 Hz), 68.89, 58.74, 56.41, 56.22, 53.01, 41.65, 41.00 (“d”, *J* = 16.8 Hz), 40.52, 40.23 (“d”, *J* = 5.9 Hz), 37.95, 36.92, 36.12 (“d”, *J* = 31.2 Hz), 35.44, 34.90 (“d”, *J* = 17.1 Hz), 34.18 (“d”, *J* = 20.0 Hz), 28.57, 27.19, 26.38, 21.99 (“d”, *J* = 29.9 Hz), 15.92.

**Synthesis of 7-(1-(4-(5-((4-hydroxy-1-((*R*)-3-phenylbutanoyl)piperidin-4-yl)methyl)-4-oxo-4,5-dihydro-1*H*-pyrazolo[3,4-*d*]pyrimidin-1-yl)phenyl)piperidine-4-carbonyl)-*N*-((*S*)-1-((2*S*,4*R*)-4-hydroxy-2-((4-(4-methylthiazol-5-yl)benzyl)carbamoyl)pyrrolidin-1-yl)-3,3-dimethyl-1-oxobutan-2-yl)-7-azaspiro[3.5]nonane-2-carboxamide (NK240)**

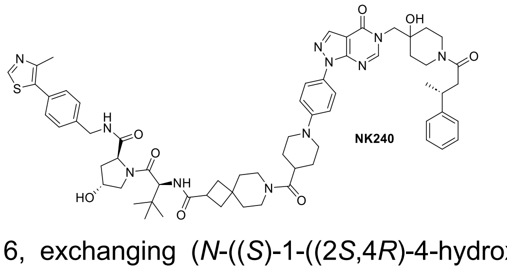

NK240

7-(1-(4-(5-((4-hydroxy-1-((*R*)-3-phenylbutanoyl)-piperidin-4-yl)methyl)-4-oxo-4,5-dihydro-1*H*-pyrazolo[3,4-*d*]pyrimidin-1-yl)phenyl)piperidine-4-carbonyl)-*N*-((*S*)-1-((2*S*,4*R*)-4-hydroxy-2-((4-(4-methylthiazol-5-yl)benzyl)carbamoyl)pyrrolidin-1-yl)-3,3-dimethyl-1-oxobutan-2-yl)-7-azaspiro[3.5]-nonane-2-carboxamide (NK240) was synthesized as described for NK239 (Scheme 6, exchanging (*N*-((*S*)-1-((2*S*,4*R*)-4-hydroxy-2-((4-(4-methylthiazol-5-yl)benzyl)carbamoyl)pyrrolidin-1-yl)-3,3-dimethyl-1-oxobutan-2-yl)-7-azaspiro[3.5]nonane-2-carboxamide (NK216) for (2*S*,4*R*)-1-((*S*)-2-amino-3,3-dimethyl-butanoyl)-4-hydroxy-*N*-(4-(4-methylthiazol-5-yl)benzyl)-pyrrolidine-2-carboxamide).

HRMS (m/z) $[M+2H]^{2+}$ calculated for $C_{64}H_{81}N_{11}O_8S^{2+}$: 581.7990, found: 581.7987.

H NMR (600 MHz, DMSO-*d*$_6$) δ (ppm): 8.99 (s, 1H), 8.56 (t, *J* = 6.1 Hz, 1H), 8.30 (s, 1H), 8.28 (d, *J* = 14.2 Hz, 1H), 7.81 (d, *J* = 8.4 Hz, 2H), 7.71 (t, *J* = 10.3 Hz, 1H), 7.43 – 7.36 (m, 4H), 7.26 (dd, *J* = 8.7, 6.2 Hz, 4H), 7.18 – 7.13 (m, 3H), 4.54 (d, *J* = 9.3 Hz, 1H), 4.43 (td, *J* = 8.8, 4.7 Hz, 2H), 4.36 (d, *J* = 3.8

Hz, 1H), 4.23 (d, $J$ = 5.5 Hz, 1H), 4.20 (d, $J$ = 5.4 Hz, 1H), 4.04 (d, $J$ = 6.8 Hz, 1H), 3.99 (s, 1H), 3.94 (d, $J$ = 3.5 Hz, 1H), 3.81 – 3.77 (m, 2H), 3.67 – 3.65 (m, 2H), 3.48 – 3.32 (m, 4H), 3.20 (dt, $J$ = 34.7, 8.4 Hz, 3H), 2.87 (q, $J$ = 17.4, 16.3 Hz, 4H), 2.64 – 2.54 (m, 2H), 2.44 (s, 6H), 2.06 – 1.81 (m, 6H), 1.70 (s, 4H), 1.59 – 1.31 (m, 8H), 1.20 (dd, $J$ = 6.9, 1.2 Hz, 3H), 0.93 (s, 9H).

$^{13}$C NMR (151 MHz, DMSO-$d_6$) δ (ppm): 174.55, 172.61, 172.42, 170.10, 169.58 (“d”, $J$ = 2.9 Hz), 157.39 (“d”, $J$ = 8.1 Hz), 152.37, 151.96, 151.03, 148.17, 147.11 (“d”, $J$ = 16.6 Hz), 139.99, 136.07, 131.65, 130.10, 129.12, 128.68 (“d”, $J$ = 4.9 Hz), 127.90, 127.38 (“d”, $J$ = 3.9 Hz), 126.42 (“d”, $J$ = 5.9 Hz), 123.38, 116.46, 116.37, 106.73, 69.59 (“d”, $J$ = 7.1 Hz), 69.36, 59.18, 56.92, 56.85, 53.48, 48.78, 42.11, 41.48 (“d”, $J$ = 17.1 Hz), 40.71 (“d”, $J$ = 5.7 Hz), 38.78, 38.43, 37.62, 37.40, 37.32, 36.59 (“d”, $J$ = 31.1 Hz), 35.83, 35.39 (“d”, $J$ = 11.6 Hz), 35.31, 34.65 (“d”, $J$ = 19.4 Hz), 34.51, 34.05, 32.17, 28.21, 26.82, 22.46 (“d”, $J$ = 29.8 Hz), 16.41.

**Synthesis of (2*S*,4*R*)-4-hydroxy-1-((*S*)-2-(3-(4-(4-(5-((4-hydroxy-1-((*R*)-3-phenylbutanoyl)-piperidin-4-yl)methyl)-4-oxo-4,5-dihydro-1*H*-pyrazolo[3,4-d]pyrimidin-1-yl)phenyl)piperazin-1-yl)propanamido)-3,3-dimethylbutanoyl)-*N*-(4-(4-methylthiazol-5-yl)benzyl)pyrrolidine-2-carboxamide TFA salt (NK237)**

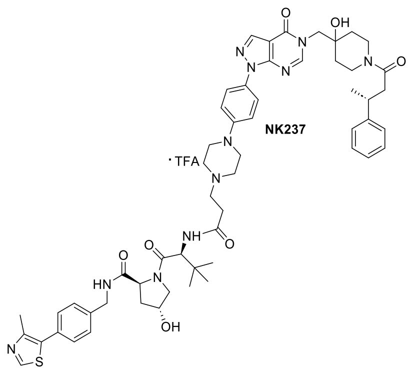

(2*S*,4*R*)-4-hydroxy-1-((*S*)-2-(3-(4-(4-(5-((4-hydroxy-1-((*R*)-3-phenylbutanoyl)-piperidin-4-yl)methyl)-4-oxo-4,5-di-hydro-1*H*-pyrazolo[3,4-d]pyrimidin-1-yl)phenyl)-piperazin-1-yl)propanamido)-3,3-dimethylbutanoyl)-*N*-(4-(4-methyl-thiazol-5-yl)benzyl)pyrrolidine-2-carboxamide TFA salt (NK237) was synthesized as described for NK239 (Scheme 6, exchanging (*R*)-1-(4-(5-((4-hydroxy-1-(3-phenylbutanoyl)piperidin-4-yl)methyl)-4-oxo-4,5-dihydro-1*H*-pyrazolo[3,4-*d*]pyrimidin-1-yl)phenyl)piperidine-4-carboxylic (NK226) acid for (*R*)-3-(4-(4-(5-((4-hydroxy-1-(3-phenylbutanoyl)piperidin-4-yl)methyl)-4-oxo-4,5-dihydro-1*H*-pyrazolo[3,4-*d*]pyrimidin-1-yl)phenyl)piperazin-1-yl)propanoic acid (NK235).

HRMS (m/z) $[M+2H]^{2+}$ calculated for $C_{56}H_{71}N_{11}O_7S^{2+}$: 520.7624, found: 520.7630.

$^{1}$H NMR (600 MHz, DMSO-$d_6$) δ (ppm): 9.62 (s, 1H), 9.00 (s, 1H), 8.59 (t, $J$ = 6.1 Hz, 1H), 8.37 – 8.26 (m, 3H), 7.89 – 7.84 (m, 2H), 7.46 – 7.37 (m, 4H), 7.27 (dd, $J$ = 8.4, 6.1 Hz, 4H), 7.21 – 7.14 (m, 3H), 4.59 (d, $J$ = 9.3 Hz, 1H), 4.47 (d, $J$ = 6.7 Hz, 1H), 4.45 (s, 1H), 4.38 (dt, $J$ = 4.2, 2.1 Hz, 1H), 4.23 (dd, $J$ = 15.8, 5.4 Hz, 1H), 4.09 – 3.93 (m, 6H), 3.72 – 3.55 (m, 6H), 3.43 (d, $J$ = 7.7 Hz, 2H), 3.20 (ddd, $J$ = 28.0, 12.6, 5.2 Hz, 4H), 3.07 (t, $J$ = 12.8 Hz, 1H), 2.91 – 2.85 (m, 1H), 2.80 (dt, $J$ = 14.7, 7.7 Hz, 1H), 2.65 – 2.54 (m, 2H), 2.45 (s, 3H), 2.07 – 2.03 (m, 1H), 1.93 (ddd, $J$ = 13.0, 8.8, 4.6 Hz, 1H), 1.45 – 1.26 (m, 4H), 1.21 (dd, $J$ = 6.9, 1.4 Hz, 3H), 0.98 (s, 9H).

$^{13}$C NMR (151 MHz, DMSO-$d_6$) δ (ppm): 172.32, 169.74, 169.58 (“d”, $J$ = 2.8 Hz), 169.16, 157.36 (“d”, $J$ = 8.2 Hz), 152.46, 151.97, 151.11, 148.84, 148.18, 147.11 (“d”, $J$ = 16.0 Hz), 139.97, 136.22, 131.63, 131.09, 130.13, 129.12, 128.68 (“d”, $J$ = 4.8 Hz), 127.89, 127.37 (“d”, $J$ = 4.0 Hz), 126.42 (“d”, $J$ = 5.1 Hz), 123.39, 116.56, 106.80, 69.60 (“d”, $J$ = 7.2 Hz), 69.36, 59.23, 57.18, 56.95, 53.46, 52.39, 51.29, 45.80, 42.12, 41.47 (“d”, $J$ = 17.2 Hz), 40.91, 40.71 (“d”, $J$ = 6.2 Hz), 38.52, 37.40, 36.69, 36.48, 35.92, 35.39 (“d”, $J$ = 17.3 Hz), 34.64 (“d”, $J$ = 20.0 Hz), 29.69, 26.84, 22.46 (“d”, $J$ = 29.8 Hz), 16.41.

**Synthesis of (2*S*,4*R*)-4-hydroxy-1-((*S*)-2-(5-(4-(4-(5-((4-hydroxy-1-((*R*)-3-phenylbutanoyl)-piperidin-4-yl)methyl)-4-oxo-4,5-dihydro-1*H*-pyrazolo[3,4-*d*]pyrimidin-1-yl)phenyl)piperazin-1-yl)pentanamido)-3,3-dimethylbutanoyl)-*N*-(4-(4-methylthiazol-5-yl)benzyl)pyrrolidine-2-carboxamide TFA salt (NK238)**

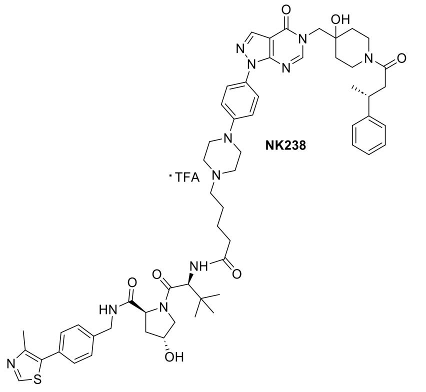

(2*S*,4*R*)-4-hydroxy-1-((*S*)-2-(5-(4-(4-(5-((4-hydroxy-1-((*R*)-3-phenylbutanoyl)-piperidin-4-yl)methyl)-4-oxo-4,5-dihydro-1*H*-pyrazolo[3,4-*d*]pyrimidin-1-yl)phenyl)piperazin-1-yl)pentanamido)-3,3-dimethylbutanoyl)-*N*-(4-(4-methylthiazol-5-yl)benzyl)pyrrolidine-2-carboxamide TFA salt (NK238) was synthesized as described for NK239 (Scheme 6, exchanging (*R*)-1-(4-(5-((4-hydroxy-1-(3-phenylbutanoyl)piperidin-4-yl)methyl)-4-oxo-4,5-dihydro-1*H*-pyrazolo[3,4-*d*]pyrimidin-1-yl)phenyl)piperidine-4-carboxylic acid (NK226) for (*R*)-5-(4-(4-(5-((4-hydroxy-1-(3-phenylbutanoyl)piperidin-4-yl)methyl)-4-oxo-4,5-dihydro-1*H*-pyrazolo[3,4-*d*]pyrimidin-1-yl)phenyl)piperazin-1-yl)pentanoic acid (NK236).

HRMS (m/z) $[M+2H]^{2+}$ calculated for $C_{58}H_{75}N_{11}O_7S^{2+}$: 534.7781, found: 534.7774.

$^1$H NMR (600 MHz, DMSO-*d*$_6$) δ (ppm): 9.48 (s, 1H), 8.98 (s, 1H), 8.56 (t, *J* = 6.1 Hz, 1H), 8.31 (s, 1H), 8.28 (d, *J* = 13.9 Hz, 1H), 7.95 (d, *J* = 9.3 Hz, 1H), 7.86 (d, *J* = 9.1 Hz, 2H), 7.44 – 7.36 (m, 4H), 7.26 (dd, *J* = 8.4, 6.1 Hz, 4H), 7.20 – 7.13 (m, 3H), 4.56 (d, *J* = 9.3 Hz, 1H), 4.43 (dt, *J* = 16.2, 7.1 Hz, 2H), 4.37 (dt, *J* = 4.5, 2.1 Hz, 1H), 4.22 (dd, *J* = 15.8, 5.5 Hz, 2H), 4.06 – 4.03 (m, 2H), 3.95 – 3.91 (m, 2H), 3.70 – 3.64 (m, 3H), 3.59 (d, *J* = 11.9 Hz, 2H), 3.24 (t, *J* = 11.3 Hz, 1H), 3.17 (dd, *J* = 9.3, 5.3 Hz, 5H), 3.08 – 3.00 (m, 2H), 2.90 – 2.83 (m, 1H), 2.65 – 2.54 (m, 2H), 2.44 (s, 3H), 2.33 (dt, *J* = 14.5, 7.3 Hz, 1H), 2.24 (dt, *J* = 14.5, 7.2 Hz, 1H), 2.07 – 2.02 (m, 1H), 1.91 (ddd, *J* = 12.9, 8.7, 4.6 Hz, 1H), 1.67 (p, *J* = 7.7 Hz, 2H), 1.59 – 1.52 (m, 2H), 1.39 – 1.23 (m, 4H), 1.20 (dd, *J* = 6.9, 1.4 Hz, 3H), 0.96 (s, 9H).

$^{13}$C NMR (151 MHz, DMSO-*d*$_6$) δ (ppm): 171.89, 171.60, 169.65, 169.10 (“d”, *J* = 2.8 Hz), 156.88 (“d”, *J* = 8.0 Hz), 151.98, 151.48, 150.64, 148.38, 147.70, 146.63 (“d”, *J* = 15.9 Hz), 139.49, 135.75, 131.15, 130.63, 129.64, 128.64, 128.20 (“d”, *J* = 4.7 Hz), 127.41, 126.90 (“d”, *J* = 4.1 Hz), 125.94 (“d”, *J* = 5.0 Hz), 122.92, 116.09, 106.32, 69.12 (“d”, *J* = 7.2 Hz), 68.88, 58.72, 56.45, 56.39, 55.25, 52.98, 50.68 (d, *J* = 11.5 Hz), 45.38, 41.64, 41.00 (“d”, *J* = 17.0 Hz), 40.43, 40.23 (“d”, *J* = 6.2 Hz), 38.04, 36.92, 36.11 (“d”, *J* = 31.1 Hz), 35.93, 35.27, 34.92 (“d”, *J* = 17.3 Hz), 34.23, 34.06, 26.40, 22.91, 22.44, 21.99 (“d”, *J* = 29.7 Hz), 15.93.

**Synthesis of 7-((1*r*,4*S*)-4-(4-(4-(5-((4-hydroxy-1-((*R*)-3-phenylbutanoyl)piperidin-4-yl)methyl)-4-oxo-4,5-dihydro-1*H*-pyrazolo[3,4-d]pyrimidin-1-yl)phenyl)piperazine-1-carbonyl)cyclohexane-1-carbonyl)-*N*-((*S*)-1-((2*S*,4*R*)-4-hydroxy-2-((4-(4-methylthiazol-5-yl)benzyl)carbamoyl)pyrrolidin-1-yl)-3,3-dimethyl-1-oxobutan-2-yl)-7-azaspiro[3.5]nonane-2-carboxamide (NK242)**

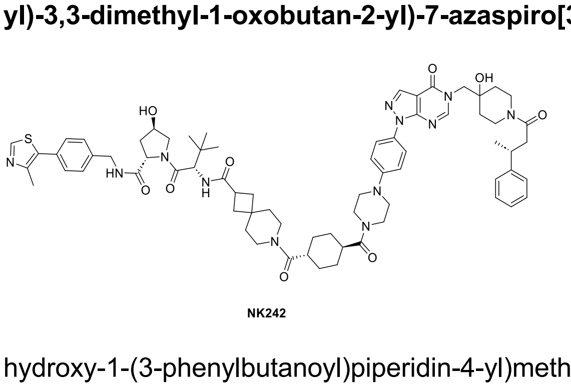

7-((1*r*,4*S*)-4-(4-(4-(5-((4-hydroxy-1-((*R*)-3-phenylbutanoyl)piperidin-4-yl)methyl)-4-oxo-4,5-dihydro-1*H*-pyrazolo[3,4-*d*]pyrimidin-1-yl)phenyl)piperazine-1-carbonyl)cyclohexane-1-carbonyl)-*N*-((*S*)-1-((2*S*,4*R*)-4-hydroxy-2-((4-(4-methyl-thiazol-5-yl)benzyl)carbamoyl)-pyrrolidin-1-yl)-3,3-dimethyl-1-oxobutan-2-yl)-7-azaspiro[3.5]nonane-2-carboxamide (NK242) was synthesized as described for NK239 (Scheme 6, exchanging (*R*)-1-(4-(5-((4-hydroxy-1-(3-phenylbutanoyl)piperidin-4-yl)methyl)-4-oxo-4,5-dihydro-1*H*-pyrazolo[3,4-*d*]pyrimidin-1-yl)phenyl)piperidine-4-carboxylic acid (NK226) for (1r,4r)-4-(4-(4-(5-((4-hydroxy-1-((R)-3-phenylbutanoyl)piperidin-4-yl)methyl)-4-oxo-4,5-dihydro-1*H*-pyrazolo[3,4-*d*]pyrimidin-1-yl)phenyl)piperazine-1-carbonyl)cyclohexane-1-carboxylic acid (NK231) and *N*-((*S*)-1-((2*S*,4*R*)-4-hydroxy-2-((4-(4-methylthiazol-5-yl)benzyl)carbamoyl)pyrrolidin-1-yl)-3,3-dimethyl-1-oxobutan-2-yl)-7-azaspiro[3.5]nonane-2-carboxamide (NK216) for 2*S*,4*R*)-1-((*S*)-2-amino-3,3-dimethyl-butanoyl)-4-hydroxy-*N*-(4-(4-methylthiazol-5-yl)benzyl)-pyrrolidine-2-carboxamide.

HRMS (m/z) $[M+2H]^{2+}$ calculated for $C_{70}H_{90}N_{12}O_9S^{2+}$: 673.3332, found: 673.3322.

$^{1}$H NMR (600 MHz, DMSO-$d_6$) δ (ppm): 8.99 (s, 1H), 8.56 (t, *J* = 6.1 Hz, 1H), 8.30 – 8.26 (m, 2H), 7.82 (d, *J* = 9.0 Hz, 2H), 7.70 (dd, *J* = 9.8, 5.2 Hz, 1H), 7.43 – 7.37 (m, 4H), 7.26 (dd, *J* = 8.7, 6.1 Hz, 4H), 7.17 – 7.12 (m, 3H), 4.53 (d, *J* = 9.3 Hz, 1H), 4.42 (td, *J* = 8.7, 4.7 Hz, 2H), 4.35 (d, *J* = 3.6 Hz, 1H), 4.21 (dd, *J* = 15.8, 5.5 Hz, 1H), 4.05 (q, *J* = 8.6, 7.0 Hz, 1H), 3.98 (d, *J* = 14.0 Hz, 1H), 3.94 (d, *J* = 3.5 Hz, 1H), 3.67 (dd, *J* = 10.9, 4.6 Hz, 6H), 3.62 (p, *J* = 3.3, 2.9 Hz, 1H), 3.41 (d, *J* = 10.5 Hz, 2H), 3.31 (s, 2H), 3.26– 3.13 (m, 8H), 2.89 – 2.84 (m, 1H), 2.66 – 2.56 (m, 2H), 2.44 (s, 3H), 2.03 (ddd, *J* = 10.2, 8.1, 2.4 Hz, 1H), 1.99 – 1.89 (m, 3H), 1.82 (d, *J* = 8.0 Hz, 2H), 1.68 (d, *J* = 26.6 Hz, 3H), 1.51 (d, *J* = 48.0 Hz, 8H), 1.39 – 1.24 (m, 6H), 1.20 (dd, *J* = 7.0, 1.2 Hz, 3H), 0.92 (s, 9H).

$^{13}$C NMR (151 MHz, DMSO-$d_6$) δ (ppm): 174.08, 173.29, 172.96, 172.94, 171.94, 169.62, 169.10 (“d”, *J* = 2.7 Hz), 156.91 (“d”, *J* = 7.9 Hz), 151.91 (“d”, *J* = 2.8 Hz), 151.48, 150.67 – 150.49 (“d”), 149.66, 147.69, 146.63 (“d”, *J* = 16.5 Hz), 139.51, 135.60, 131.17, 129.93, 129.62, 128.64, 128.20 (“d”, *J* = 5.0 Hz), 127.42, 126.90 (“d”, *J* = 3.9 Hz), 125.94 (“d”, *J* = 5.7 Hz), 122.84, 115.74, 106.25, 69.11 (“d”, *J* = 7.3 Hz), 68.88, 58.70, 56.43, 56.37, 53.55, 52.99, 48.75, 48.12, 44.48, 41.63, 41.00 (“d”, *J* = 16.8 Hz), 40.79, 40.23 (“d”, *J* = 5.7 Hz), 38.92, 38.56 (“d”, *J* = 10.0 Hz), 37.94, 36.92, 36.11 (“d”, *J* = 31.2 Hz), 35.34, 34.90, 34.84, 34.17 (d, *J* = 20.1 Hz), 34.04, 33.55, 31.68, 28.22, 28.18, 26.34, 21.99 (“d”, *J* = 29.9 Hz), 18.07, 16.73, 15.93, 12.44.

**Synthesis of (2*S*,4*S*)-4-hydroxy-1-((*S*)-2-((1*r*,4*S*)-4-(4-(4-(5-((4-hydroxy-1-((*R*)-3-phenylbutanoyl)piperidin-4-yl)methyl)-4-oxo-4,5-dihydro-1*H*-pyrazolo[3,4-*d*]pyrimidin-1-yl)phenyl)piperazine-1-carbonyl)cyclohexane-1-carboxamido)-3,3-dimethylbutanoyl)-*N*-(4-(4-methylthiazol-5-yl)benzyl)pyrrolidine-2-carboxamide (NK245)**

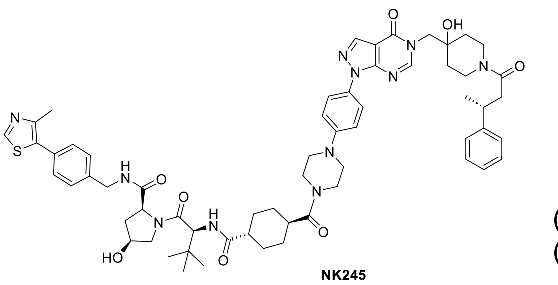

NK245

(2*S*,4*S*)-4-hydroxy-1-((*S*)-2-((1*r*,4*S*)-4-(4-(4-(5-((4-hydroxy-1-((*R*)-3-phenylbutanoyl)piperidin-4-yl)methyl)-4-oxo-4,5-dihydro-1*H*-pyrazolo[3,4-*d*]-pyrimidin-1-yl)phenyl)piperazine-1-carbonyl)-cyclohexane-1-carboxamido)-3,3-dimethylbutanoyl)-N-(4-(4-methylthiazol-5-yl)benzyl)pyrrolidine-2-carboxamide (NK245) was synthesized as described for NK239 (Scheme 6, exchanging (*R*)-1-(4-(5-((4-hydroxy-1-(3-phenylbutanoyl)piperidin-4-yl)methyl)-4-oxo-4,5-dihydro-1*H*-pyrazolo[3,4-*d*]pyrimidin-1-yl)phenyl)piperidine-4-carboxylic acid (NK226) for (1*r*,4*r*)-4-(4-(4-(5-((4-hydroxy-1-((*R*)-3-phenylbutanoyl)piperidin-4-yl)methyl)-4-oxo-4,5-dihydro-1*H*-pyrazolo[3,4-*d*]pyrimidin-1-yl)phenyl)piperazine-1-carbonyl)cyclohexane-1-carboxylic acid (NK231) and 2*S*,4*S*)-1-((*S*)-2-amino-3,3-dimethyl-butanoyl)-4-hydroxy-*N*-(4-(4-methylthiazol-5-yl)benzyl)-pyrrolidine-2-carboxamide for (2*S*,4*R*)-1-((*S*)-2-amino-3,3-dimethyl-butanoyl)-4-hydroxy-*N*-(4-(4-methylthiazol-5-yl)benzyl)-pyrrolidine-2-carboxamide .

HRMS (m/z) $[M+2H]^{2+}$ calculated for $C_{61}H_{77}N_{11}O_8S^{2+}$: 561.7833, found: 561.7828.

$^{1}$H NMR (700 MHz, DMSO-$d_6$) δ (ppm): 8.92 (s, 1H), 8.58 (t, *J* = 6.1 Hz, 1H), 8.23 (d, *J* = 1.1 Hz, 1H), 8.21 (d, *J* = 16.5 Hz, 1H), 7.76 (d, *J* = 9.0 Hz, 2H), 7.68 (d, *J* = 8.8 Hz, 1H), 7.33 (q, *J* = 8.4 Hz, 4H), 7.21 – 7.18 (m, 4H), 7.10 – 7.05 (m, 3H), 4.40 – 4.36 (m, 2H), 4.30 (dd, *J* = 8.6, 6.1 Hz, 1H), 4.19 (dd, *J* = 15.7, 5.5 Hz, 1H), 4.15 (d, *J* = 5.6 Hz, 1H), 4.00 – 3.94 (m, 2H), 3.93 – 3.88 (m, 1H), 3.89 – 3.83 (m, 1H), 3.55 (s, 6H), 3.38 (dd, *J* = 10.1, 5.3 Hz, 1H), 3.18 – 3.07 (m, 6H), 2.82 – 2.77 (m, 1H), 2.57 – 2.50 (m, 2H), 2.38 (s, 3H), 2.27 (ddd, *J* = 12.6, 8.5, 5.7 Hz, 1H), 1.72 – 1.58 (m, 5H), 1.48 – 1.24 (m, 8H), 1.14 (d, *J* = 6.9 Hz, 3H), 0.89 (s, 9H).

$^{13}$C NMR (176 MHz, DMSO-$d_6$) δ (ppm): 175.62, 173.77, 172.97, 170.41, 169.58 (“d”, *J* = 3.1 Hz), 157.39 (“d”, *J* = 8.7 Hz), 152.39, 151.97, 151.04, 150.14, 148.20, 147.11 (“d”, *J* = 19.3 Hz), 139.68, 136.09 (“d”, *J* = 5.5 Hz), 131.62, 130.41, 130.20, 129.14, 128.68 (“d”, *J* = 5.9 Hz), 127.93, 127.37 (“d”, *J* = 4.7 Hz), 126.42 (“d”, *J* = 6.3 Hz), 123.32, 116.21, 106.74, 69.61 (“d”, *J* = 3.9 Hz), 69.57, 58.99, 56.90, 56.07, 53.47, 49.20, 48.53, 44.95, 43.03, 42.26, 41.42, 41.25 (“d”, *J* = 3.1 Hz), 40.71 (“d”, *J* = 6.5 Hz), 38.85, 37.39, 36.59 (“d”, *J* = 36.1 Hz), 35.32, 34.65 (“d”, *J* = 22.2 Hz), 29.43, 28.88, 28.64, 28.28, 26.84, 22.46 (“d”, *J* = 34.6 Hz), 16.41.

#### 3.7 Synthesis of VHL-alkylcarboxy-functionalized PROTACs

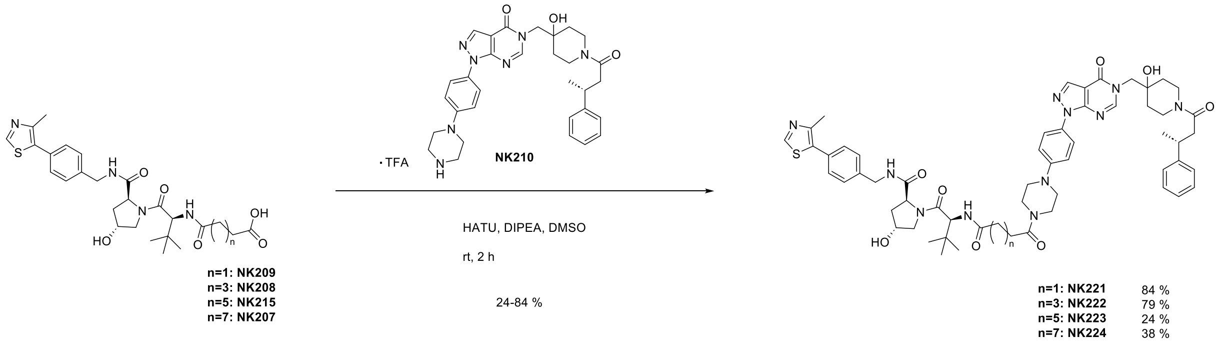

**Scheme 7.** Synthesis scheme of PROTACs NK221, NK222, NK223 and NK224.

##### Synthesis of (2*S*,4*R*)-4-hydroxy-1-((*S*)-2-(4-(4-(4-(5-((4-hydroxy-1-((*R*)-3-phenylbutanoyl)-piperidin-4-yl)methyl)-4-oxo-4,5-dihydro-1*H*-pyrazolo[3,4-*d*]pyrimidin-1-yl)phenyl)piperazin-1-yl)-4-oxobutanamido)-3,3-dimethylbutanoyl)-*N*-(4-(4-methylthiazol-5-yl)benzyl)pyrrolidine-2-carboxamide (NK221)

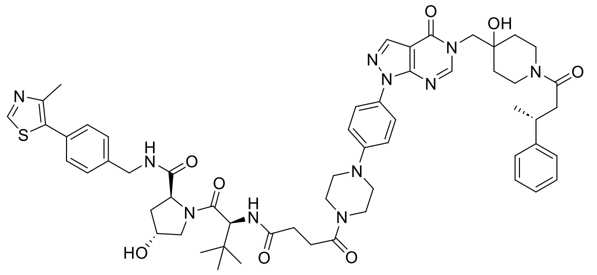

NK221

4-(((*S*)-1-((2*S*,4*R*)-4-Hydroxy-2-((4-(4-methylthiazol-5-yl)benzyl)carbamoyl)pyrrolidin-1-yl)-3,3-dimethyl-1-oxobutan-2-yl)amino)-4-oxobutanoic acid (NK209) (9.3 mg, 0.02 mmol, 1.1 eq), HATU (7.4 mg, 0.02 mmol, 1.3 eq) and DIPEA (7.2 mg, 0.06 mmol, 4 eq) were dissolved in DMSO (1 mL) and stirred at rt for 15 min. To this was added (*R*)-5-((4-hydroxy-1-(3-phenylbutanoyl)piperidin-4-yl)methyl)-1-(4-(piperazin-1-yl)phenyl)-1,5-dihydro-4*H*-pyrazolo[3,4-*d*]pyrimidin-4-one (NK210) and the reaction was stirred for 2 h at rt. After completion, the crude product was purified by preparative reverse phase column chromatography (ACN/$H_2O$ + 0.1 % TFA). After evaporation of the solvents under reduced pressure, the product (2*S*,4*R*)-4-hydroxy-1-((*S*)-2-(4-(4-(4-(5-((4-hydroxy-1-((*R*)-3-phenylbutanoyl)-piperidin-4-yl)methyl)-4-oxo-4,5-dihydro-1*H*-pyrazolo[3,4-*d*]pyrimidin-1-yl)phenyl)piperazin-1-yl)-4-oxobutanamido)-3,3-dimethylbutanoyl)-*N*-(4-(4-methylthiazol-5-yl)benzyl)pyrrolidine-2-carboxamide (NK221) (13.4 mg, 0.01 mmol, 84 %) was obtained as a white solid.

HRMS (m/z) $[M+2H]^{2+}$ calculated for $C_{57}H_{71}N_{11}O_8S^{2+}$: 534.7599, found: 534.7592.

$^{1}$H NMR (500 MHz, DMSO-$d_6$) δ (ppm): 8.99 (s, 1H), 8.56 (t, *J* = 6.1 Hz, 1H), 8.29 (s, 1H), 8.28 (d, *J* = 11.9 Hz, 1H), 7.93 (d, *J* = 9.3 Hz, 1H), 7.82 (d, *J* = 9.2 Hz, 2H), 7.40 (q, *J* = 8.4 Hz, 4H), 7.26 (dd, *J* = 7.4, 5.6 Hz, 4H), 7.18 – 7.11 (m, 3H), 4.53 (d, *J* = 9.3 Hz, 1H), 4.47 – 4.40 (m, 2H), 4.35 (tt, *J* = 4.4, 2.3 Hz, 1H), 4.22 (dd, *J* = 15.8, 5.5 Hz, 1H), 4.08 – 3.93 (m, 3H), 3.69 – 3.59 (m, 7H), 3.29 – 3.12 (m, 6H), 2.90 – 2.85 (m, 1H), 2.64 – 2.54 (m, 5H), 2.45 (s, 3H), 2.43 – 2.40 (m, 1H), 2.06 – 2.00 (m, 1H), 1.90 (ddd, *J* = 12.9, 8.6, 4.6 Hz, 1H), 1.57 – 1.24 (m, 4H), 1.20 (d, *J* = 6.9 Hz, 3H), 0.94 (s, 9H).

$^{13}$C NMR (126 MHz, DMSO-$d_6$) δ (ppm): 171.94, 171.28, 170.05, 169.59, 169.10, 156.91 („d“, *J* = 6.7 Hz), 151.90, 151.50, 150.55, 149.62, 147.65, 146.64 („d“, *J* = 13.6 Hz), 139.53, 135.60, 131.20, 129.91, 129.61, 128.65, 128.20 („d“, *J* = 4.0 Hz), 127.43, 126.90 („d“, *J* = 3.2 Hz), 125.96, 122.86, 115.72, 106.26, 69.12 („d“, *J* = 6.0 Hz), 68.89, 58.72, 56.45, 56.30, 53.00, 48.15 („d“, *J* = 46.9 Hz), 44.48, 41.65, 40.91 („d“, *J* = 10.0 Hz), 40.23 („d“, *J* = 4.5 Hz), 37.94, 36.92, 36.12 („d“, *J* = 25.9 Hz), 35.37, 34.90 („d“, *J* = 15.6 Hz), 34.18 („d“, *J* = 17.3 Hz), 30.14, 27.97, 26.39, 21.99 („d“, *J* = 24.8 Hz), 15.92.

**Synthesis of (2*S*,4*R*)-4-hydroxy-1-((*S*)-2-(6-(4-(4-(5-((4-hydroxy-1-((*R*)-3-phenylbutanoyl)-piperidin-4-yl)methyl)-4-oxo-4,5-dihydro-1*H*-pyrazolo[3,4-*d*]pyrimidin-1-yl)phenyl)piperazin-1-yl)-6-oxohexanamido)-3,3-dimethylbutanoyl)-*N*-(4-(4-methylthiazol-5-yl)benzyl)pyrrolidine-2-carboxamide (NK222)**

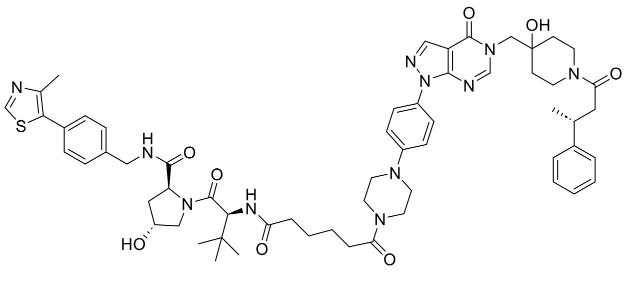

NK222

(2*S*,4*R*)-4-Hydroxy-1-((*S*)-2-(6-(4-(4-(5-((4-hydroxy-1-((*R*)-3-phenylbutanoyl)-piperidin-4-yl)methyl)-4-oxo-4,5-dihydro-1*H*-pyrazolo[3,4-*d*]pyrimidin-1-yl)phenyl)piperazin-1-yl)-6-oxohexanamido)-3,3-dimethylbutanoyl)-*N*-(4-(4-methylthiazol-5-yl)benzyl)pyrrolidine-2-carboxamide (NK222) was synthesized as described for NK221 (Scheme 7, exchanging 4-(((*S*)-1-((2*S*,4*R*)-4-hydroxy-2-((4-(4-methylthiazol-5-yl)benzyl)carbamoyl)pyrrolidin-1-yl)-3,3-dimethyl-1-oxobutan-2-yl)amino)-4-oxobutanoic acid (NK209) for 6-(((*S*)-1-((2*S*,4*R*)-4-hydroxy-2-((4-(4-methylthiazol-5-yl)benzyl)carbamoyl)-pyrrolidin-1-yl)-3,3-dimethyl-1-oxobutan-2-yl)amino)-6-oxohexanoic acid (NK208)).

HRMS (m/z) $[M+2H]^{2+}$ calculated for $C_{59}H_{75}N_{11}O_8S^{2+}$: 548.7755 Found: 548.7752.

$^1$H NMR (700 MHz, DMSO-*d*$_6$) δ (ppm): 8.98 (s, 1H), 8.55 (t, *J* = 6.1 Hz, 1H), 8.29 (d, *J* = 1.1 Hz, 1H), 8.27 (d, *J* = 16.5 Hz, 1H), 7.87 (d, *J* = 9.3 Hz, 1H), 7.83 – 7.81 (m, 2H), 7.42 (d, *J* = 8.2 Hz, 2H), 7.38 (d, *J* = 8.3 Hz, 2H), 7.27 – 7.24 (m, 4H), 7.15 (td, *J* = 6.3, 3.6 Hz, 1H), 7.13 – 7.11 (m, 2H), 4.55 (d, *J* = 9.4 Hz, 1H), 4.45 – 4.41 (m, 2H), 4.35 (dt, *J* = 4.4, 2.1 Hz, 1H), 4.21 (dd, *J* = 15.8, 5.4 Hz, 1H), 4.07 – 3.93 (m, 3H), 3.66 (dd, *J* = 12.6, 8.5 Hz, 4H), 3.62 (t, *J* = 5.3 Hz, 2H), 3.24 (t, *J* = 5.4 Hz, 2H), 3.20 – 3.14 (m, 4H), 2.90 – 2.83 (m, 1H), 2.63 – 2.53 (m, 3H), 2.44 (s, 3H), 2.37 (dt, *J* = 7.6, 3.3 Hz, 2H), 2.30 (dt, *J* = 10.5, 7.1 Hz, 1H), 2.17 – 2.12 (m, 1H), 2.05 – 2.01 (m, 1H), 1.90 (ddd, *J* = 12.9, 8.6, 4.6 Hz, 1H), 1.56 – 1.53 (m, 2H), 1.45 – 1.22 (m, 3H), 1.20 (dd, *J* = 6.9, 1.5 Hz, 3H), 0.93 (s, 9H).

$^{13}$C NMR (176 MHz, DMSO-*d*$_6$) δ (ppm): 171.96, 171.94, 170.62, 169.70, 169.10 (“d”, *J* = 3.0 Hz), 151.88, 151.46, 150.54, 149.68, 147.70, 146.64 (“d”, *J* = 19.7 Hz), 139.51, 135.59, 131.16, 129.89, 129.62, 128.63, 128.20 (“d”, *J* = 5.9 Hz), 127.41, 126.90 (“d”, *J* = 4.7 Hz), 125.94 (“d”, *J* = 6.6 Hz), 122.85, 122.84, 115.72, 106.25, 69.11 (“d”, *J* = 8.3 Hz), 68.86, 58.68, 56.36, 56.29, 53.00, 48.23 (“d”, *J* = 73.8 Hz), 44.60, 41.64, 40.99 (“d”, *J* = 18.7 Hz), 40.71, 40.69, 40.23 (“d”, *J* = 6.9 Hz), 40.02, 37.95, 36.97 – 36.86 (m), 36.11 (“d”, *J* = 36.5 Hz), 35.22, 34.69, 34.17 (“d”, *J* = 23.4 Hz), 31.98, 26.39, 25.22, 24.43, 21.99 (“d”, *J* = 34.8 Hz), 15.94.

**Synthesis of (2*S*,4*R*)-4-hydroxy-1-((*S*)-2-(8-(4-(4-(5-((4-hydroxy-1-((*R*)-3-phenylbutanoyl)-piperidin-4-yl)methyl)-4-oxo-4,5-dihydro-1*H*-pyrazolo[3,4-*d*]pyrimidin-1-yl)phenyl)piperazin-1-yl)-8-oxooctanamido)-3,3-dimethylbutanoyl)-*N*-(4-(4-methylthiazol-5-yl)benzyl)pyrrolidine-2-carboxamide (NK223)**

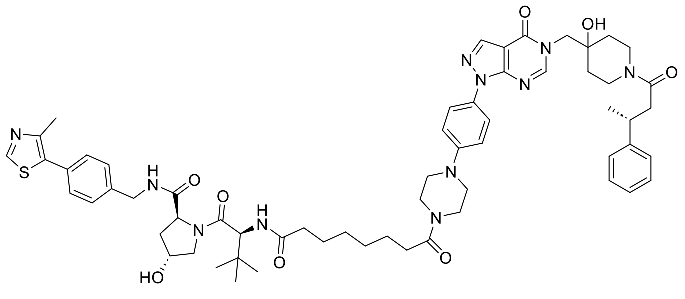

NK223

(2*S*,4*R*)-4-Hydroxy-1-((*S*)-2-(8-(4-(4-(5-((4-hydroxy-1-((*R*)-3-phenylbutanoyl)-piperidin-4-yl)methyl)-4-oxo-4,5-dihydro-1*H*-pyrazolo[3,4-*d*]pyrimidin-1-yl)phenyl)piperazin-1-yl)-8-oxooctanamido)-3,3-dimethylbutanoyl)-*N*-(4-(4-methylthiazol-5-yl)benzyl)pyrrolidine-2-carboxamide (NK223) was synthesized as described for NK221 (Scheme 7, exchanging 4-(((*S*)-1-((2*S*,4*R*)-4-hydroxy-2-((4-(4-methylthiazol-5-yl)benzyl)carbamoyl)pyrrolidin-1-yl)-3,3-dimethyl-1-oxobutan-2-yl)amino)-4-oxobutanoic acid (NK209) for 8-(((*S*)-1-((2*S*,4*R*)-4-hydroxy-2-((4-(4-methylthiazol-5-yl)benzyl)carbamoyl)-pyrrolidin-1-yl)-3,3-dimethyl-1-oxobutan-2-yl)amino)-8-oxooctanoic acid (NK215).

HRMS (m/z) [M+2H]$^{2+}$ calculated for $C_{61}H_{77}N_{11}O_8S^{2+}$: 562.7912, found: 562.7907.

$^{1}$H NMR (600 MHz, DMSO-$d_6$) δ (ppm): 8.98 (s, 1H), 8.56 (t, *J* = 6.1 Hz, 1H), 8.29 (d, *J* = 0.8 Hz, 1H), 8.28 (d, *J* = 14.2 Hz, 1H), 7.85 (d, *J* = 9.4 Hz, 1H), 7.82 (d, *J* = 9.1 Hz, 2H), 7.42 – 7.36 (m, 4H), 7.27 – 7.23 (m, 4zH), 7.16 – 7.11 (m, 3H), 4.54 (d, *J* = 9.4 Hz, 1H), 4.45 – 4.40 (m, 2H), 4.35 (dt, *J* = 4.3, 2.0 Hz, 1H), 4.21 (dd, *J* = 15.8, 5.5 Hz, 2H), 4.05 – 4.02 (m, 3H), 3.93 (d, *J* = 3.7 Hz, 2H), 3.69 – 3.64 (m, 2H), 3.61 (t, *J* = 5.1 Hz, 4H), 3.25 – 3.22 (m, 2H), 3.17 (q, *J* = 7.5, 5.7 Hz, 3H), 2.90 – 2.83 (m, 1H), 2.64 – 2.54 (m, 3H), 2.44 (s, 3H), 2.34 (t, *J* = 7.5 Hz, 2H), 2.12 (ddd, *J* = 14.2, 8.1, 6.3 Hz, 1H), 2.03 (ddd, *J* = 11.3, 7.7, 2.3 Hz, 1H), 1.50 (p, *J* = 8.7, 8.2 Hz, 4H), 1.32 – 1.23 (m, 8H), 1.20 (d, *J* = 6.8 Hz, 3H), 0.93 (s, 9H).

$^{13}$C NMR (151 MHz, DMSO-$d_6$) δ (ppm): 172.57, 172.43, 171.20, 170.19, 169.58 („d“, *J* = 2.7 Hz), 152.37, 151.94, 151.02, 150.14, 148.17, 147.11 (“d”, *J* = 16.6 Hz), 139.99, 136.07, 131.65, 130.23 (“d”, *J* = 42.3 Hz), 129.11, 128.70, 127.89, 127.38 (“d”, *J* = 4.0 Hz), 126.42 (“d”, *J* = 5.6 Hz), 123.32, 116.19, 106.73, 69.59 (“d”, *J* = 7.5 Hz), 69.34, 59.16, 56.79 (“d”, *J* = 12.1 Hz), 53.47, 48.71 (“d”, *J* = 62.5 Hz), 45.09, 42.11, 41.48 (“d”, *J* = 16.9 Hz), 41.16, 40.91, 40.69, 38.43, 37.39, 36.59 (“d”, *J* = 31.3 Hz), 35.68, 35.43, 35.32, 34.65 (“d”, *J* = 20.7 Hz), 32.69, 28.99, 26.86, 25.83, 25.17, 22.46 (“d”, *J* = 30.1 Hz), 16.41.

**Synthesis of (2*S*,4*R*)-4-hydroxy-1-((*S*)-2-(10-(4-(4-(5-((4-hydroxy-1-((*R*)-3-phenylbutanoyl)-piperidin-4-yl)methyl)-4-oxo-4,5-dihydro-1*H*-pyrazolo[3,4-*d*]pyrimidin-1-yl)phenyl)piperazin-1-yl)-10-oxodecanamido)-3,3-dimethylbutanoyl)-*N*-(4-(4-methylthiazol-5-yl)benzyl)pyrrolidine-2-carboxamide (NK224)**

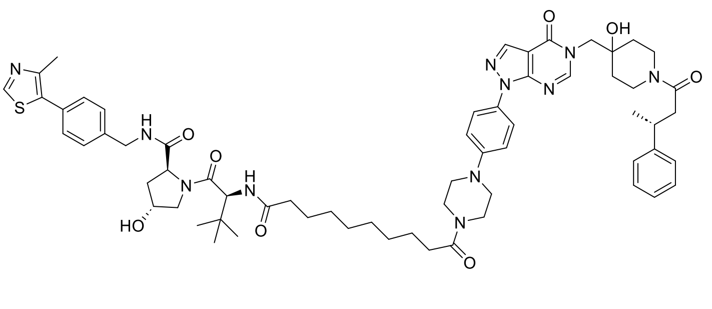

**NK224**

(2*S*,4*R*)-4-Hydroxy-1-((*S*)-2-(10-(4-(4-(5-((4-hydroxy-1-((*R*)-3-phenylbutanoyl)-piperidin-4-yl)methyl)-4-oxo-4,5-dihydro-1*H*-pyrazolo[3,4-*d*]pyrimidin-1-yl)phenyl)piperazin-1-yl)-10-oxodecanamido)-3,3-dimethylbutanoyl)-*N*-(4-(4-methylthiazol-5-yl)benzyl)pyrrolidine-2-carboxamide (NK224) was synthesized as described for NK221 (Scheme 7, exchanging 4-(((*S*)-1-((2*S*,4*R*)-4-hydroxy-2-((4-(4-methylthiazol-5-yl)benzyl)carbamoyl)pyrrolidin-1-yl)-3,3-dimethyl-1-oxobutan-2-yl)amino)-4-oxobutanoic acid (NK209) for 10-(((*S*)-1-((2*S*,4*R*)-4-hydroxy-2-((4-(4-methylthiazol-5-yl)benzyl)carbamoyl)-pyrrolidin-1-yl)-3,3-dimethyl-1-oxobutan-2-yl)amino)-10-oxodecanoic acid (NK207).

HRMS (m/z) [M+2H]$^{2+}$ calculated for $C_{63}H_{83}N_{11}O_8S^{2+}$ 576.8068, found: 576.8056.

$^{1}$H NMR (700 MHz, DMSO-$d_6$) δ (ppm): 8.98 (s, 1H), 8.55 (t, *J* = 6.1 Hz, 1H), 8.29 (s, 1H), 8.28 (d, *J* = 16.4 Hz, 1H), 7.84 – 7.81 (m, 3H), 7.42 – 7.37 (m, 4H), 7.28 – 7.24 (m, 4H), 7.16 – 7.11 (m, 3H), 4.54 (d, *J* = 9.4 Hz, 1H), 4.46 – 4.40 (m, 2H), 4.35 (dt, *J* = 4.3, 2.0 Hz, 1H), 4.21 (dd, *J* = 15.8, 5.5 Hz, 1H), 4.06 – 3.92 (m, 4H) 3.68 (d, *J* = 5.8 Hz, 2H), 3.66 (s, 2H), 3.61 (t, *J* = 5.3 Hz, 4H), 3.23 (d, *J* = 6.7 Hz, 1H), 3.20 – 3.15 (m, 1H), 2.87 (t, *J* = 10.9 Hz, 1H), 2.64 – 2.56 (m, 2H), 2.44 (s, 3H), 2.34 (t, *J* = 7.5 Hz, 2H), 2.25 (dt, *J* = 14.7, 7.6 Hz, 1H), 2.11 (ddd, *J* = 14.2, 8.1, 6.2 Hz, 1H), 2.05 – 2.01 (m, 1H), 1.90 (ddd, *J* = 12.9, 8.5, 4.6 Hz, 1H), 1.51 (p, *J* = 7.4 Hz, 4H), 1.32 – 1.23 (m, 12H), 1.20 (dd, *J* = 7.0, 1.5 Hz, 3H), 0.93 (s, 9H).

$^{13}$C NMR (176 MHz, DMSO-$d_6$) (ppm): δ 172.08, 171.93, 170.73, 169.70, 169.09 („d“, *J* = 3.8 Hz), 157.96 („d“, *J* = 34.4 Hz), 151.87, 151.45, 149.66, 147.69, 139.50, 135.58, 131.16, 129.88, 129.62, 128.62, 128.19 („d“, *J* = 5.4 Hz), 127.41, 126.89 („d“, *J* = 4.7 Hz), 126.34, 125.93 („d“, *J* = 6.2 Hz), 122.84, 115.70, 114.82, 106.25, 68.97 („d“, *J* = 42.5 Hz), 58.67, 56.33, 56.25, 52.98, 48.43, 48.02, 46.87, 44.61, 41.63, 40.99 („d“, *J* = 18.7 Hz), 40.68, 40.22 („d“, *J* = 7.1 Hz), 37.94, 36.91, 36.10 („d“, *J* = 36.3 Hz), 35.19, 34.85, 32.24, 28.80 („d“, *J* = 7.6 Hz), 28.69, 28.63, 26.37, 25.42, 24.78, 21.98 („d“, *J* = 34.8 Hz), 16.68, 15.93.

#### 3.8 Synthesis of VHL-cyclohexyl-functionalized PROTACs

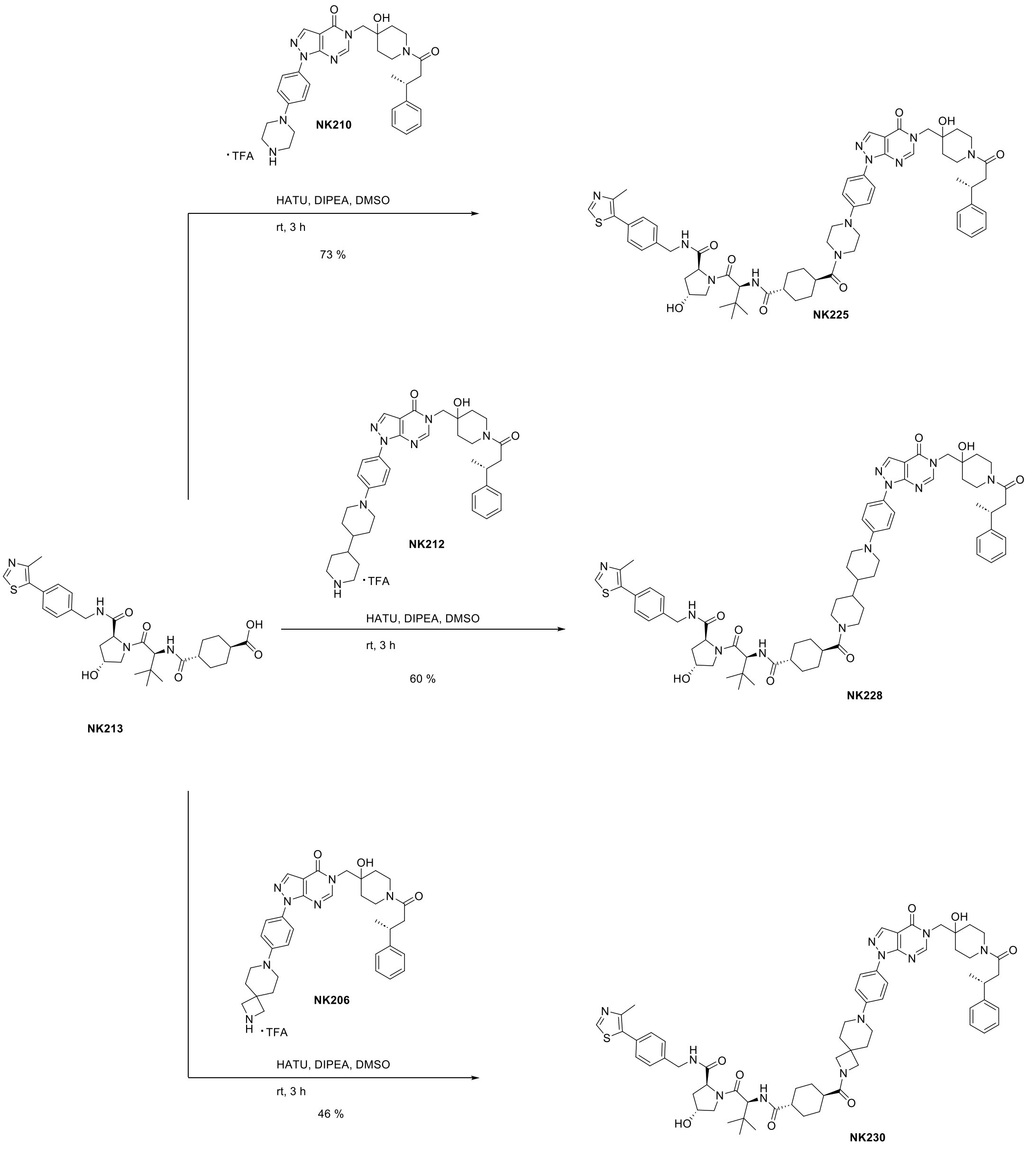

**Scheme 8.** Synthesis scheme of cyclohexyl-linker PROTACs NK225, NK228 and NK230.

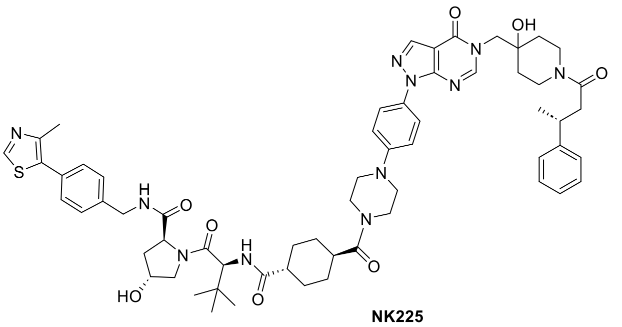

NK225

**Synthesis of (2*S*,4*R*)-4-hydroxy-1-((*S*)-2-((1*r*,4*S*)-4-(4-(4-(5-((4-hydroxy-1-((*R*)-3-phenylbutanoyl)-piperidin-4-yl)methyl)-4-oxo-4,5-dihydro-1*H*-pyrazolo[3,4-*d*]pyrimidin-1-yl)phenyl)piperazine-1-carbonyl)cyclohexane-1-carboxamido)-3,3-dimethylbutanoyl)-*N*-(4-(4-methylthiazol-5-yl)benzyl)pyrrolidine-2-carboxamide (NK225)**

NK213 (10.5 mg, 0.02 mmol, 1.2 eq), HATU (7.4 mg, 0.02 mmol, 1.3 eq) and DIPEA (7.7 mg, 0.06 mmol, 4.0 eq) were dissolved in DMSO (1 mL) and stirred at rt for 15 min. To this was added NK210 (10.0 mg, 0.01 mmol, 1.0 eq) and the mixture was stirred at rt for 2 h. After completion, the crude product was directly purified by preparative reverse phase column chromatography (ACN/$H_2O$ + 0.1 % TFA). After evaporation of the solvents under reduced pressure, the product (2*S*,4*R*)-4-hydroxy-1-((*S*)-2-

((1*r*,4*S*)-4-(4-(4-(5-((4-hydroxy-1-((*R*)-3-phenylbutanoyl)-piperidin-4-yl)methyl)-4-oxo-4,5-dihydro-1*H*-pyrazolo[3,4-*d*]pyrimidin-1-yl)phenyl)piperazine-1-carbonyl)cyclohexane-1-carboxamido)-3,3-dimethylbutanoyl)-*N*-(4-(4-methylthiazol-5-yl)benzyl)pyrrolidine-2-carboxamide (NK225) (12.2 mg, 0.01 mmol, 73 % yield) was obtained as a white powder.

HRMS (m/z) $[M+2H]^{2+}$ calculated for $C_{61}H_{77}N_{11}O_8S^{2+}$: 561.7833, found: 561.7823.

$^1$H NMR (600 MHz, DMSO-*d*$_6$) δ (ppm): 9.00 (s, 1H), 8.58 (t, *J* = 6.1 Hz, 1H), 8.30 (s, 1H), 8.28 (d, *J* = 14.1 Hz, 1H), 7.83 (d, *J* = 9.0 Hz, 2H), 7.75 (d, *J* = 9.3 Hz, 1H), 7.45 – 7.37 (m, 4H), 7.27 (dd, *J* = 8.6, 6.1 Hz, 4H), 7.19 – 7.11 (m, 3H), 4.55 (s, 2H), 4.47 – 4.42 (m, 4H), 4.36 (dt, *J* = 4.3, 1.9 Hz, 1H), 4.23 (dd, *J* = 15.8, 5.5 Hz, 1H), 4.09 – 4.01 (m, 1H), 4.01 – 3.93 (m, 2H), 3.70 – 3.59 (m, 6H), 3.27 – 3.15 (m, 6H), 2.91 – 2.83 (m, 1H), 2.66 – 2.57 (m, 2H), 2.45 (s, 3H), 2.07 – 2.02 (m, 1H), 1.91 (ddd, *J* = 12.9, 8.6, 4.6 Hz, 1H), 1.83 – 1.79 (m, 1H), 1.71 (q, *J* = 15.1, 12.8 Hz, 3H), 1.54 – 1.30 (m, 8H), 1.21 (dd, *J* = 7.0, 1.2 Hz, 3H), 0.94 (s, 9H).

$^{13}$C NMR (151 MHz, DMSO-*d*$_6$) δ (ppm): 175.29, 173.78, 172.43, 170.12, 169.58 (“d”, *J* = 2.9 Hz), 157.39 (“d”, *J* = 8.1 Hz), 152.38 (“d”, *J* = 3.0 Hz), 151.98, 151.03 (“d”, *J* = 1.7 Hz), 150.12, 148.14, 147.11 (“d”, *J* = 16.5 Hz), 140.00, 136.08, 131.67, 130.42, 130.09, 129.12, 128.68 (“d”, *J* = 5.0 Hz), 127.91, 127.37 (“d”, *J* = 4.1 Hz), 126.42 (“d”, *J* = 5.5 Hz), 123.32, 116.23, 106.74, 69.59 (“d”, *J* = 7.2 Hz), 69.36, 59.19, 56.82, 56.54, 53.47, 49.23, 48.59, 44.95, 43.16, 42.12, 41.47 (“d”, *J* = 17.3 Hz), 41.25, 40.71 (“d”, *J* = 6.2 Hz), 38.87, 38.42, 37.39, 36.59 (“d”, *J* = 31.1 Hz), 35.92, 35.37 (“d”, *J* = 17.1 Hz), 34.65 (“d”, *J* = 19.5 Hz), 29.62, 28.91, 28.63, 28.19, 26.84, 22.46 (“d”, *J* = 29.9 Hz), 16.40.

**Synthesis of (2*S*,4*R*)-4-hydroxy-1-((*S*)-2-((1*r*,4*S*)-4-(1'-(4-(5-((4-hydroxy-1-((*R*)-3-phenylbutanoyl)piperidin-4-yl)methyl)-4-oxo-4,5-dihydro-1*H*-pyrazolo[3,4-*d*]pyrimidin-1-yl)phenyl)-[4,4'-bipiperidine]-1-carbonyl)cyclohexane-1-carboxamido)-3,3-dimethylbutanoyl)-*N*-(4-(4-methylthiazol-5-yl)benzyl)pyrrolidine-2-carboxamide (NK228)**

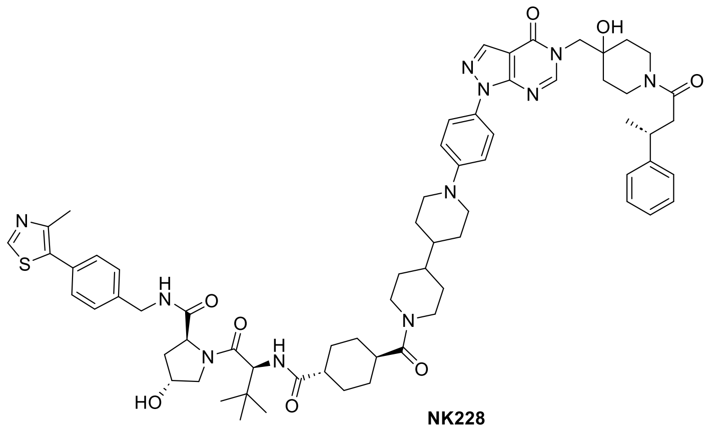

NK228

(2*S*,4*R*)-4-Hydroxy-1-((*S*)-2-((1*r*,4*S*)-4-(1'-(4-(5-((4-hydroxy-1-((*R*)-3-phenylbutanoyl)piperidin-4-yl)methyl)-4-oxo-4,5-dihydro-1*H*-pyrazolo[3,4-*d*]pyrimidin-1-yl)phenyl)-[4,4'-bipiperidine]-1-carbonyl)cyclohexane-1-carboxamido)-3,3-dimethylbutanoyl)-*N*-(4-(4-methylthiazol-5-yl)benzyl)pyrrolidine-2-carboxamide (NK228) was synthesized as described for NK239 (Scheme 8, exchanging (*R*)-5-((4-hydroxy-1-(3-phenylbutanoyl)piperidin-4-yl)methyl)-1-(4-(piperazin-1-yl)phenyl)-1,5-dihydro-4*H*-pyrazolo[3,4-*d*]pyrimidin-4-one (NK210) for (*R*)-1-(4-([4,4'-bipiperidin]-1-yl)phenyl)-5-((4-hydroxy-1-(3-phenylbutanoyl)piperidin-4-yl)methyl)-1,5-dihydro-4*H*-pyrazolo[3,4-*d*]pyrimidin-4-one TFA salt (NK212).

HRMS (m/z) $[M+2H]^{2+}$ calculated for $C_{67}H_{87}N_{11}O_8S^{2+}$: 602.8225, found: 602.8226.

$^1$H NMR (600 MHz, DMSO-*d*$_6$) δ (ppm): 9.00 (s, 1H), 8.57 (t, *J* = 6.1 Hz, 1H), 8.30 (s, 1H), 8.28 (d, *J* = 14.1 Hz, 1H), 7.82 (d, *J* = 8.5 Hz, 2H), 7.74 (d, *J* = 9.2 Hz, 1H), 7.44 – 7.37 (m, 4H), 7.29 – 7.24 (m, 4H), 7.18 – 7.14 (m, 3H), 4.53 (d, *J* = 9.3 Hz, 1H), 4.46 – 4.41 (m, 3H), 4.35 (td, *J* = 4.3, 2.2 Hz, 1H), 4.22 (dd, *J* = 15.8, 5.5 Hz, 1H), 4.08 – 3.96 (m, 4H), 3.94 (d, *J* = 3.9 Hz, 1H), 3.81 (d, *J* = 12.0 Hz, 2H), 3.70 – 3.65 (m, 5H), 3.26 – 3.13 (m, 3H), 2.91 (dt, *J* = 42.8, 12.9 Hz, 2H), 2.76 (s, 1H), 2.64 – 2.56 (m, 2H), 2.45 (s, 3H), 2.36 (dt, *J* = 14.0, 3.0 Hz, 1H), 2.06 – 2.01 (m, 1H), 1.91 (ddd, *J* = 12.9, 8.4, 4.6 Hz, 1H), 1.82 – 1.65 (m, 8H), 1.57 – 1.24 (m, 13H), 1.21 (d, *J* = 6.8 Hz, 3H), 0.94 (s, 9H).

$^{13}$C NMR (151 MHz, DMSO-*d*$_6$) δ (ppm): 174.83, 172.81, 171.96, 169.65, 169.10 (“d”, *J* = 2.7 Hz), 156.91 (“d”, *J* = 8.0 Hz), 151.91, 151.89, 151.48, 150.89, 150.56, 147.70, 146.63 (“d”, *J* = 16.4 Hz), 139.51, 135.61, 131.17, 129.63, 128.86, 128.64, 128.21 (“d”, *J* = 4.8 Hz), 127.43, 126.90 (“d”, *J* = 4.0 Hz), 125.94

(“d”, $J$ = 5.6 Hz), 122.88, 106.26, 69.12 (“d”, $J$ = 7.2 Hz), 68.87, 58.71, 56.20 (“d”, $J$ = 42.5 Hz), 52.99, 45.12, 42.72, 42.46, 41.64, 41.44, 41.00 (“d”, $J$ = 16.5 Hz), 40.43, 40.23 (“d”, $J$ = 6.0 Hz), 38.51, 37.94, 36.92, 36.12 (“d”, $J$ = 31.1 Hz), 35.44, 35.38, 34.90 (“d”, $J$ = 16.4 Hz), 34.18 (“d”, $J$ = 19.5 Hz), 29.97, 28.86, 28.33, 26.36, 21.99 (“d”, $J$ = 29.9 Hz), 15.97, 15.94.

**Synthesis of (2*S*,4*R*)-4-hydroxy-1-((*S*)-2-((1*r*,4*S*)-4-(7-(4-(5-((4-hydroxy-1-((*R*)-3-phenylbutanoyl)-piperidin-4-yl)methyl)-4-oxo-4,5-dihydro-1*H*-pyrazolo[3,4-*d*]pyrimidin-1-yl)phenyl)-2,7-diazaspiro[3.5]nonane-2-carbonyl)cyclohexane-1-carboxamido)-3,3-dimethylbutanoyl)-*N*-(4-(4-methylthiazol-5-yl)benzyl)pyrrolidine-2-carboxamide (NK230)**

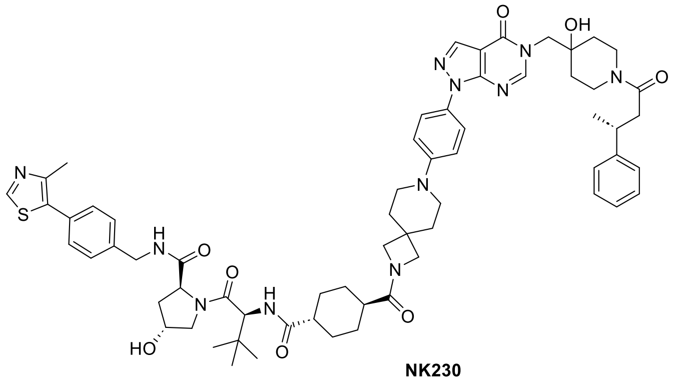

NK230

(2*S*,4*R*)-4-Hydroxy-1-((*S*)-2-((1*r*,4*S*)-4-(7-(4-(5-((4-hydroxy-1-((*R*)-3-phenylbutanoyl)-piperidin-4-yl)methyl)-4-oxo-4,5-dihydro-1*H*-pyrazolo[3,4-*d*]pyrimidin-1-yl)phenyl)-2,7-diazaspiro[3.5]nonane-2-carbonyl)-cyclohexane-1-carboxamido)-3,3-dimethylbutanoyl)-*N*-(4-(4-methylthiazol-5-yl)benzyl)pyrrolidine-2-carboxamide (NK230) was synthesized as described for NK239 (Scheme 8, exchanging (*R*)-5-((4-hydroxy-1-(3-phenylbutanoyl)piperidin-4-yl)methyl)-1-(4-(piperazin-1-yl)phenyl)-1,5-dihydro-4*H*-pyrazolo[3,4-*d*]pyrimidin-4-one (NK210) for (*R*)-1-(4-(2,7-diazaspiro[3.5]nonan-7-yl)phenyl)-5-((4-hydroxy-1-(3-phenylbutanoyl)piperidin-4-yl)methyl)-1,5-dihydro-4*H*-pyrazolo[3,4-*d*]pyrimidin-4-one TFA salt (NK220).

HRMS (m/z) $[M+2H]^{2+}$ calculated for $C_{64}H_{81}N_{11}O_8S^{2+}$: 581.7990, found: 581.7978.

$^1H$ NMR (700 MHz, DMSO-$d_6$) δ (ppm): 8.99 (s, 1H), 8.56 (t, $J$ = 6.1 Hz, 1H), 8.31 – 8.25 (m, 2H), 7.81 (d, $J$ = 8.6 Hz, 2H), 7.74 (d, $J$ = 9.3 Hz, 1H), 7.44 – 7.37 (m, 4H), 7.28 – 7.23 (m, 4H), 7.15 (ddd, $J$ = 8.6, 6.3, 2.4 Hz, 3H), 4.52 (d, $J$ = 9.4 Hz, 1H), 4.46 – 4.40 (m, 2H), 4.35 (tt, $J$ = 4.5, 2.5 Hz, 2H), 4.22 (dd, $J$ = 15.8, 5.5 Hz, 1H), 4.08 – 4.01 (m, 2H), 4.00 – 3.93 (m, 2H), 3.91 (s, 2H), 3.69 – 3.57 (m, 5H), 3.24 (d, $J$ = 22.3 Hz, 4H), 3.19 – 3.14 (m, 1H), 2.90 – 2.83 (m, 1H), 2.64 – 2.52 (m, 2H), 2.45 (s, 3H), 2.39 – 2.33 (m, 1H), 2.20 – 2.14 (m, 1H), 2.06 – 2.00 (m, 1H), 1.90 (ddd, $J$ = 12.9, 8.6, 4.6 Hz, 1H), 1.81 (dq, $J$ = 18.8, 9.8, 7.7 Hz, 4H), 1.68 (q, $J$ = 13.7, 11.1 Hz, 2H), 1.56 – 1.24 (m, 9H), 1.20 (dd, $J$ = 6.9, 1.5 Hz, 3H), 0.93 (s, 9H).

$^{13}C$ NMR (176 MHz, DMSO-$d_6$) δ (ppm): 174.83, 174.78, 171.94, 169.64, 169.10 (“d”, $J$ = 3.1 Hz), 156.91 (“d”, $J$ = 9.1 Hz), 151.88 (“d”, $J$ = 3.3 Hz), 151.48, 150.54, 147.67, 146.63 (“d”, $J$ = 19.2 Hz), 139.51, 135.58, 131.18, 129.62, 128.64, 128.20 (“d”, $J$ = 5.6 Hz), 127.43, 126.89 (“d”, $J$ = 4.6 Hz), 125.93 (“d”, $J$ = 6.8 Hz), 122.87, 116.09, 114.72, 106.24, 69.11 (“d”, $J$ = 8.6 Hz), 68.87, 59.12, 58.71, 56.90, 56.32, 56.07, 52.99, 42.52, 41.64, 41.00 (“d”, $J$ = 19.0 Hz), 40.23 (“d”, $J$ = 7.6 Hz), 38.20, 37.94, 36.91, 36.11 (“d”, $J$ = 36.1 Hz), 35.41, 34.89 (“d”, $J$ = 20.3 Hz), 34.04, 32.80, 29.13, 27.73, 27.45, 26.35, 21.98 (“d”, $J$ = 34.4 Hz), 15.92.

#### 3.9 Synthesis of VHL-cubane-functionalized PROTACs

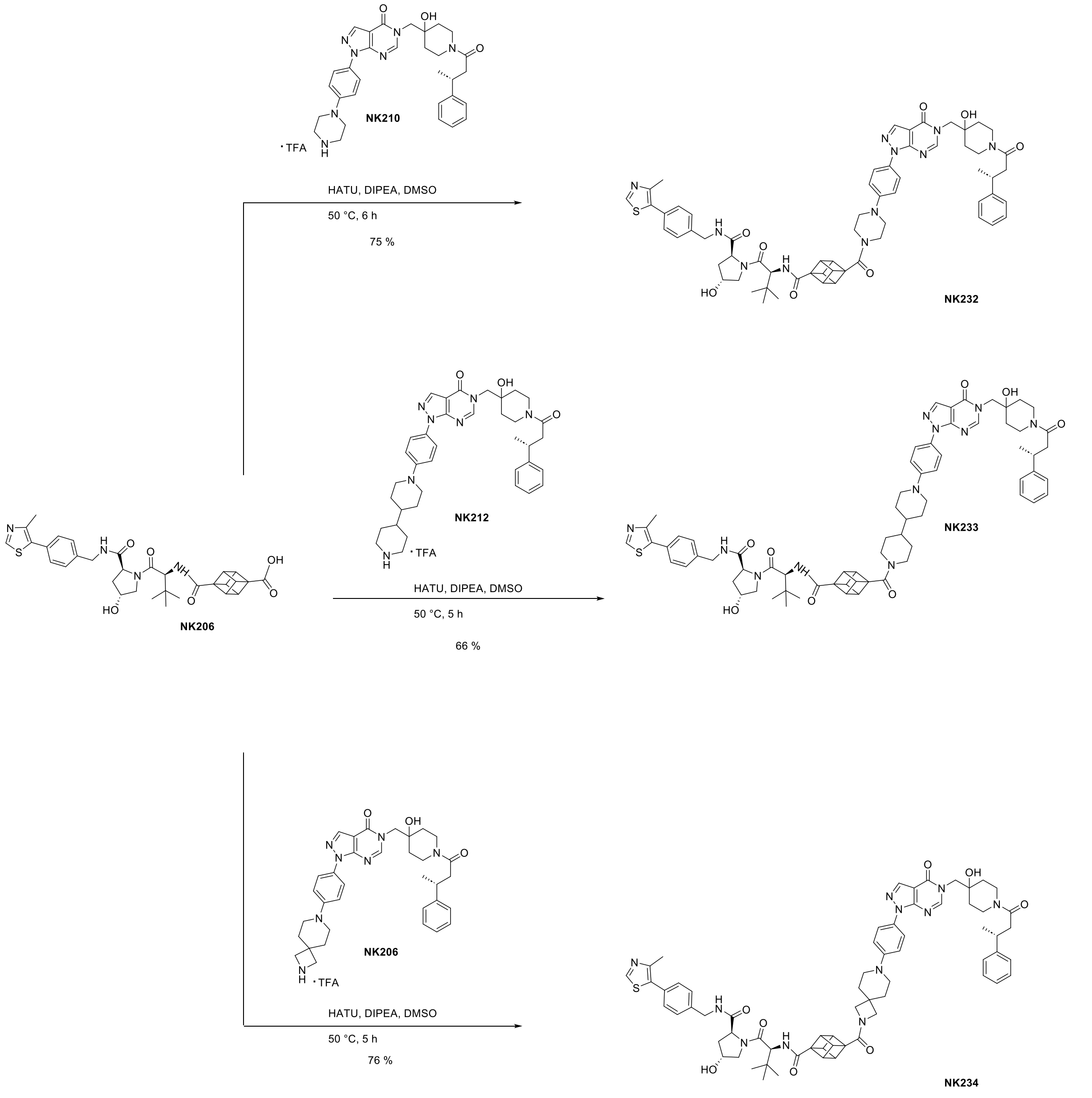

**Scheme 9.** Synthesis of VHL-cubane-functionalized PROTACs NK232, NK233 and NK234.

##### Synthesis of (2*S*,4*R*)-4-hydroxy-1-((*S*)-2-(4-(4-(4-(5-((4-hydroxy-1-((*R*)-3-phenylbutanoyl)-piperidin-4-yl)methyl)-4-oxo-4,5-dihydro-1*H*-pyrazolo[3,4-*d*]pyrimidin-1-yl)phenyl)piperazine-1-carbonyl)cubane-1-carboxamido)-3,3-dimethylbutanoyl)-*N*-(4-(4-methylthiazol-5-yl)benzyl)pyrrolidine-2-carboxamide (NK232)

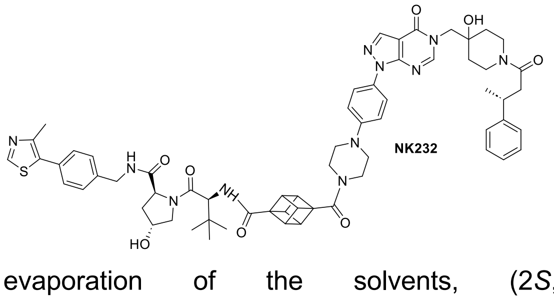

NK206 (12.6 mg, 0.02 mmol, 1.2 eq), HATU (7.4 mg, 0.02 mmol, 1.3 eq) and DIPEA (7.7 mg, 0.06 mmol, 4.0 eq) were dissolved in DMSO (1 mL) and stirred at 50 °C for 15 min. To this was added NK210 (10.0 mg, 0.01 mmol, 1.0 eq) and the mixture was stirred at 50 °C for 5 h. After completion, the crude product was directly purified by preparative reverse phase column chromatography (ACN/$H_2O$ + 0.1 % TFA). After evaporation of the solvents, (2*S*,4*R*)-4-hydroxy-1-((*S*)-2-(4-(4-(4-(5-((4-hydroxy-1-((*R*)-3-phenylbutanoyl)-piperidin-4-yl)methyl)-4-oxo-4,5-dihydro-1*H*-pyrazolo[3,4-*d*]pyrimidin-1-

yl)phenyl)piperazine-1-carbonyl)cubane-1-carboxamido)-3,3-dimethylbutanoyl)-*N*-(4-(4-methylthiazol-5-yl)benzyl)pyrrolidine-2-carboxamide (NK232) (12.8 mg, 0.01 mmol, 75 % yield) was obtained as a white powder.

HRMS (m/z) $[M+2H]^{2+}$ calculated for $C_{63}H_{73}N_{11}O_8S^{2+}$: 571.7667, found: 571.7667.

$^1$H NMR (500 MHz, DMSO-*d*$_6$) δ (ppm): 8.93 (s, 1H), 8.50 (t, *J* = 6.1 Hz, 1H), 8.23 (s, 1H), 8.21 (d, *J* = 11.9 Hz, 1H), 7.76 (d, *J* = 9.1 Hz, 2H), 7.68 (d, *J* = 9.3 Hz, 1H), 7.38 – 7.30 (m, 4H), 7.19 (dd, *J* = 7.3, 5.6 Hz, 4H), 7.08 (ddt, *J* = 10.3, 8.0, 3.0 Hz, 3H), 4.55 (d, *J* = 9.4 Hz, 1H), 4.40 – 4.32 (m, 2H), 4.29 (tt, *J* = 4.4, 2.5 Hz, 1H), 4.16 (dd, *J* = 15.8, 5.6 Hz, 1H), 4.08 (s, 6H), 4.02 – 3.85 (m, 3H), 3.65 – 3.51 (m, 5H), 3.27 (dt, *J* = 38.2, 4.9 Hz, 4H), 3.13 (dt, *J* = 25.2, 5.9 Hz, 4H), 2.85 – 2.76 (m, 1H), 2.58 – 2.47 (m, 2H), 2.38 (s, 3H), 2.01 – 1.95 (m, 1H), 1.84 (ddd, *J* = 13.0, 8.7, 4.6 Hz, 1H), 1.38 – 1.16 (m, 4H), 1.14 (d, *J* = 6.9 Hz, 3H), 0.89 (s, 9H).

$^{13}$C NMR (126 MHz, DMSO-*d*$_6$) δ (ppm): 172.39, 170.77, 169.94, 169.58 ("d", *J* = 1.8 Hz), 169.13, 157.39 ("d", *J* = 6.8 Hz), 152.39, 152.00, 151.04 ("d"), 150.08, 148.13, 147.11 ("d", *J* = 13.7 Hz), 139.99, 136.09, 131.68, 130.47, 130.10, 129.14, 128.68 ("d", *J* = 4.1 Hz), 127.92, 127.37 ("d", *J* = 3.3 Hz), 126.42 ("d", *J* = 4.7 Hz), 123.32, 116.35, 106.75, 69.59 ("d", *J* = 6.0 Hz), 69.35, 59.23, 57.43, 56.90, 56.67, 56.48, 53.47, 49.06, 48.52, 46.49, 44.50, 42.14, 41.48 ("d", *J* = 13.5 Hz), 41.26, 40.71 ("d", *J* = 4.8 Hz), 38.43, 37.40, 36.59 ("d", *J* = 26.0 Hz), 36.00, 35.37 ("d", *J* = 15.0 Hz), 34.65 ("d", *J* = 17.9 Hz), 26.92, 22.46 ("d", *J* = 25.0 Hz), 16.39.

**Synthesis of (2*S*,4*R*)-4-hydroxy-1-((*S*)-2-(4-(1'-(4-(5-((4-hydroxy-1-((*R*)-3-phenylbutanoyl)-piperidin-4-yl)methyl)-4-oxo-4,5-dihydro-1*H*-pyrazolo[3,4-*d*]pyrimidin-1-yl)phenyl)-[4,4'-bipiperidine]-1-carbonyl)cubane-1-carboxamido)-3,3-dimethylbutanoyl)-*N*-(4-(4-methylthiazol-5-yl)benzyl)pyrrolidine-2-carboxamide (NK233)**

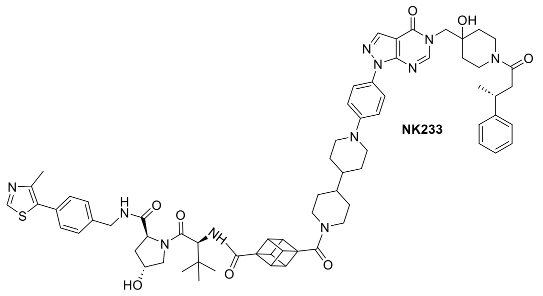

(2*S*,4*R*)-4-Hydroxy-1-((*S*)-2-(4-(1'-(4-(5-((4-hydroxy-1-((*R*)-3-phenylbutanoyl)-piperidin-4-yl)methyl)-4-oxo-4,5-dihydro-1*H*-pyrazolo[3,4-*d*]pyrimidin-1-yl)phenyl)-[4,4'-bipiperidine]-1-carbonyl)cubane-1-carboxamido)-3,3-dimethylbutanoyl)-*N*-(4-(4-methylthiazol-5-yl)benzyl)pyrrolidine-2-carboxamide (NK233) was synthesized as described for NK232 (Scheme 9, exchanging (*R*)-5-((4-hydroxy-1-(3-phenylbutanoyl)piperidin-4-yl)methyl)-1-(4-(piperazin-1-yl)phenyl)-1,5-dihydro-4*H*-pyrazolo[3,4-*d*]pyrimidin-4-one TFA salt (NK210) for (*R*)-1-(4-([4,4'-bipiperidin]-1-yl)phenyl)-5-((4-hydroxy-1-(3-phenylbutanoyl)piperidin-4-yl)methyl)-1,5-dihydro-4*H*-pyrazolo[3,4-*d*]pyrimidin-4-one TFA salt (NK212)).

HRMS (m/z) $[M+2H]^{2+}$ calculated for $C_{69}H_{83}N_{11}O_8S^{2+}$: 612.8068, found: 612.8059.

$^1$H NMR (600 MHz, DMSO-*d*$_6$) δ (ppm): 8.99 (s, 1H), 8.58 (t, *J* = 6.1 Hz, 1H), 8.30 (s, 1H), 8.28 (d, *J* = 14.2 Hz, 1H), 7.81 (d, *J* = 8.5 Hz, 2H), 7.70 (dd, *J* = 11.7, 9.5 Hz, 1H), 7.43 – 7.38 (m, 4H), 7.30 – 7.24 (m, 4H), 7.16 (ddd, *J* = 8.7, 6.3, 2.5 Hz, 3H), 4.61 (d, *J* = 9.3 Hz, 1H), 4.47 – 4.40 (m, 2H), 4.38 – 4.33 (m, 2H), 4.23 (dd, *J* = 15.8, 5.4 Hz, 1H), 4.15 – 4.07 (m, 6H), 4.06 – 3.97 (m, 2H), 3.94 (d, *J* = 4.1 Hz, 1H), 3.82 (d, *J* = 11.9 Hz, 2H), 3.70 – 3.62 (m, 4H), 3.23 – 3.14 (m, 3H), 3.04 (t, *J* = 12.5 Hz, 1H), 2.87 (t, *J* = 13.7 Hz, 1H), 2.75 (s, 1H), 2.66 – 2.57 (m, 2H), 2.45 (s, 3H), 2.09 – 1.96 (m, 2H), 1.91 (ddd, *J* = 12.9, 8.8, 4.6 Hz, 1H), 1.80 (s, 3H), 1.74 (d, *J* = 12.3 Hz, 1H), 1.44 – 1.23 (m, 10H), 1.21 (dd, *J* = 7.0, 1.5 Hz, 3H), 0.95 (s, 9H).

$^{13}$C NMR (151 MHz, DMSO-*d*$_6$) δ (ppm): 171.93, 170.34, 169.47, 169.12, 168.19, 158.05 ("d", *J* = 35.0 Hz), 151.91, 151.50, 150.80, 150.55, 147.72, 146.59, 139.50, 135.51, 132.35, 131.18, 129.65, 128.68, 128.22 ("d", *J* = 4.8 Hz), 127.45, 126.91 ("d", *J* = 4.1 Hz), 125.96 ("d", *J* = 6.1 Hz), 122.90, 117.01, 106.25, 69.12 ("d", *J* = 7.5 Hz), 68.88, 58.76, 57.21, 56.43, 56.18, 56.00, 53.01, 45.97 ("d", *J* = 11.2 Hz), 41.67, 41.48 ("d", *J* = 4.2 Hz), 40.44, 40.24 ("d", *J* = 5.5 Hz), 37.95, 36.93, 36.13 ("d", *J* = 31.6 Hz),

35.53, 35.04 (“d”, *J* = 21.7 Hz), 34.19 (“d”, *J* = 20.5 Hz), 33.41, 29.99, 29.71, 29.60, 28.69, 28.38, 26.44, 22.00 (“d”, *J* = 29.7 Hz), 15.95.

**Synthesis of (2*S*,4*R*)-4-hydroxy-1-((*S*)-2-(4-(7-(4-(5-((4-hydroxy-1-((*R*)-3-phenylbutanoyl)-piperidin-4-yl)methyl)-4-oxo-4,5-dihydro-1*H*-pyrazolo[3,4-*d*]pyrimidin-1-yl)phenyl)-2,7-diazaspiro[3.5]nonane-2-carbonyl)cubane-1-carboxamido)-3,3-dimethylbutanoyl)-*N*-(4-(4-methylthiazol-5-yl)benzyl)pyrrolidine-2-carboxamide (NK234)**

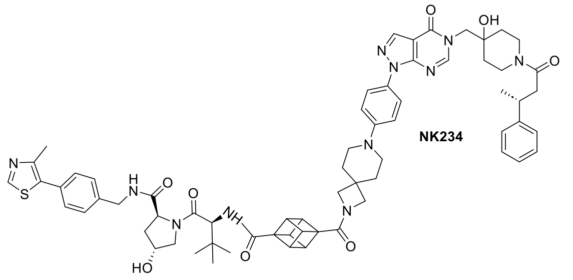

(2*S*,4*R*)-4-Hydroxy-1-((*S*)-2-(4-(7-(4-(5-((4-hydroxy-1-((*R*)-3-phenylbutanoyl)-piperidin-4-yl)methyl)-4-oxo-4,5-dihydro-1*H*-pyrazolo[3,4-*d*]pyrimidin-1-yl)phenyl)-2,7-diazaspiro[3.5]nonane-2-carbonyl)cubane-1-carboxamido)-3,3-dimethylbutanoyl)-*N*-(4-(4-methylthiazol-5-yl)benzyl)pyrrolidine-2-carboxamide (NK234) was synthesized as described for NK232 (Scheme 9, exchanging (*R*)-5-((4-hydroxy-1-(3-phenylbutanoyl)piperidin-4-yl)methyl)-1-(4-(piperazin-1-yl)phenyl)-1,5-dihydro-4*H*-pyrazolo[3,4-*d*]pyrimidin-4-one TFA salt (NK210) for (*R*)-1-(4-(2,7-diazaspiro[3.5]nonan-7-yl)phenyl)-5-((4-hydroxy-1-(3-phenylbutanoyl)piperidin-4-yl)methyl)-1,5-dihydro-4*H*-pyrazolo[3,4-*d*]pyrimidin-4-one TFA salt (NK220)).

HRMS (m/z) $[M+2H]^{2+}$ calculated for $C_{66}H_{77}N_{11}O_8S^{2+}$: 591.7833, found: 591.7835.

$^{1}H$ NMR (700 MHz, DMSO-$d_6$) δ (ppm): 8.99 (s, 1H), 8.56 (t, *J* = 6.1 Hz, 1H), 8.29 (d, *J* = 1.0 Hz, 1H), 8.28 (d, *J* = 16.5 Hz, 1H), 7.82 (d, *J* = 8.6 Hz, 2H), 7.65 (d, *J* = 9.4 Hz, 1H), 7.42 – 7.38 (m, 4H), 7.28 – 7.23 (m, 4H), 7.19 – 7.14 (m, 3H), 4.59 (d, *J* = 9.4 Hz, 1H), 4.44 – 4.40 (m, 2H), 4.35 (tt, *J* = 4.3, 2.4 Hz, 1H), 4.22 (dd, *J* = 15.8, 5.5 Hz, 1H), 4.16 (dd, *J* = 5.7, 4.1 Hz, 3H), 4.11 (dd, *J* = 5.7, 4.1 Hz, 3H), 4.07 – 3.93 (m, 3H), 3.88 (s, 2H), 3.69 – 3.61 (m, 5H), 3.25 (s, 4H), 3.22 – 3.13 (m, 2H), 2.90 – 2.84 (m, 2H), 2.63 – 2.52 (m, 2H), 2.44 (s, 3H), 2.07 – 2.02 (m, 1H), 1.90 (ddd, *J* = 12.9, 8.7, 4.6 Hz, 1H), 1.87 – 1.84 (m, 4H), 1.56 – 1.24 (m, 4H), 1.20 (dd, *J* = 7.0, 1.6 Hz, 3H), 0.94 (s, 9H).

$^{13}C$ NMR (176 MHz, DMSO-$d_6$) δ (ppm): 171.90, 170.50, 170.28, 169.45, 169.10 (“d”, *J* = 3.1 Hz), 156.91 (“d”, *J* = 9.4 Hz), 151.91, 151.50, 150.56, 148.54, 147.66, 146.63 (“d”, *J* = 19.1 Hz), 139.50, 135.61, 131.18, 129.63, 128.66, 128.20 (“d”, *J* = 5.8 Hz), 127.44, 126.89 (“d”, *J* = 4.8 Hz), 126.31, 125.94 (“d”, *J* = 6.2 Hz), 122.84, 116.24, 106.27, 69.12 (“d”, *J* = 8.4 Hz), 68.87, 58.74, 58.62, 57.91, 56.71, 56.41, 56.15, 56.08, 53.00, 46.30, 45.65, 41.66, 41.00 (“d”, *J* = 19.7 Hz), 40.43, 40.23 (“d”, *J* = 7.3 Hz), 37.94, 36.92, 36.11 (“d”, *J* = 36.2 Hz), 35.51, 34.90 (“d”, *J* = 19.5 Hz), 34.06 (“d”, *J* = 10.6 Hz), 26.42, 21.98 (“d”, *J* = 34.7 Hz), 15.91.

#### 3.10 Synthesis of enhanced PROTACs NK250 and NK266

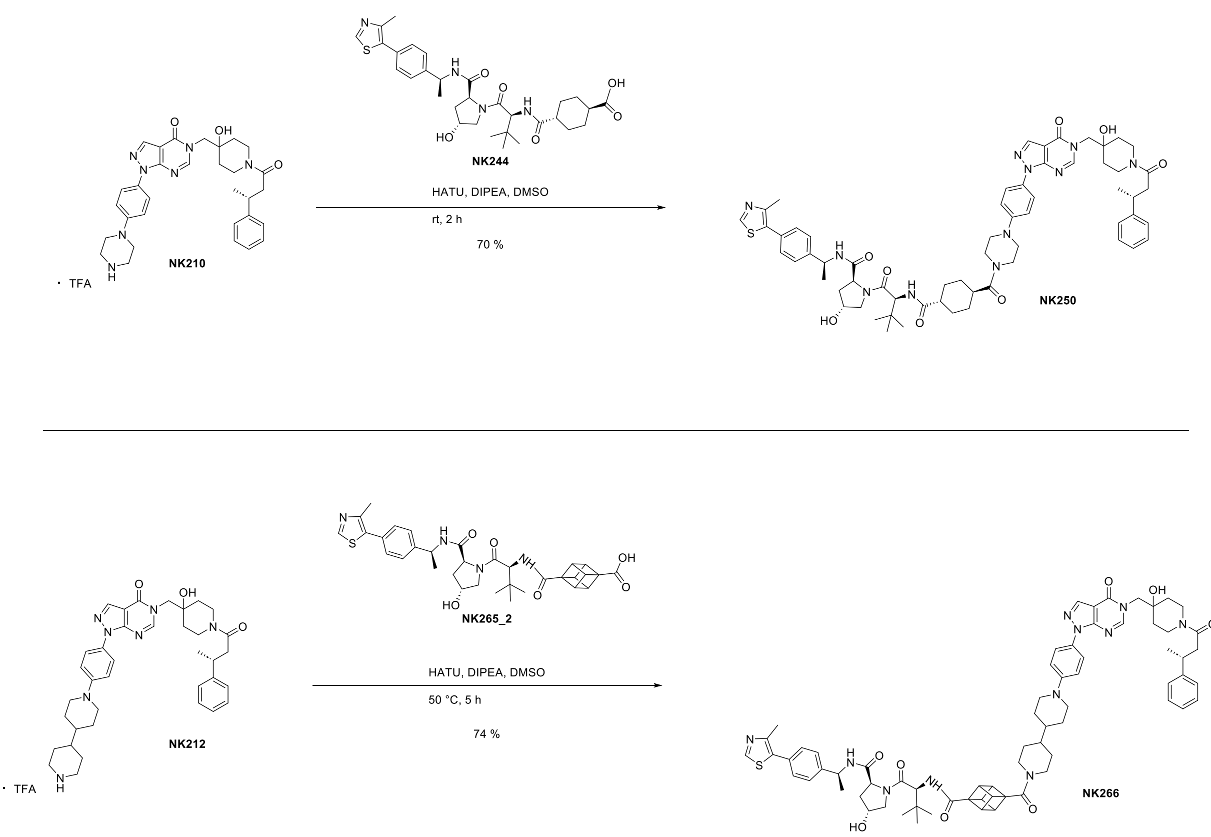

**Scheme 10.** Synthesis of enhanced PROTACs NK250 and NK266.

**Synthesis of (2*S*,4*R*)-4-hydroxy-1-((*S*)-2-((1*r*,4*S*)-4-(4-(4-(5-((4-hydroxy-1-((*R*)-3-phenylbutanoyl)-piperidin-4-yl)methyl)-4-oxo-4,5-dihydro-1*H*-pyrazolo[3,4-*d*]pyrimidin-1-yl)phenyl)piperazine-1-carbonyl)cyclohexane-1-carboxamido)-3,3-dimethylbutanoyl)-*N*-((*S*)-1-(4-(4-methylthiazol-5-yl)phenyl)ethyl)pyrrolidine-2-carboxamide (NK250)**

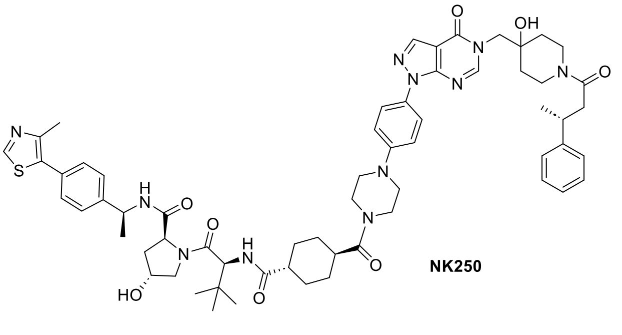

NK244 (19.7 mg, 0.03 mmol, 1.1 eq), HATU (14.8 mg, 0.04 mmol, 1.3 eq) and DIPEA (15.4 mg, 0.04 mmol, 4 eq) were dissolved in DMSO (1 mL) and stirred at rt for 20 min. To this was added NK210 (20.0 mg, 0.03 mmol, 1 eq) and the reaction was stirred for 2 h at rt. After completion, the crude product was directly purified by preparative reverse phase column chromatography (ACN/$H_2O$ + 0.1 % TFA). After evaporation of the solvents, (2*S*,4*R*)-4-Hydroxy-1-((*S*)-2-((1*r*,4*S*)-4-(4-(4-(5-((4-hydroxy-1-((*R*)-3-phenylbutanoyl)piperidin-4-yl)methyl)-4-oxo-4,5-dihydro-1*H*-pyrazolo[3,4-*d*]pyrimidin-1-yl)phenyl)piperazine-1-carbonyl)-cyclohexane-1-carboxamido)-3,3-dimethylbutanoyl)-*N*-((*S*)-1-(4-(4-methylthiazol-5-yl)phenyl)ethyl)pyrrolidine-2-carboxamide (NK250) (23.8 mg, 0.02 mmol, 70 % yield) was obtained as a white powder.

HRMS (m/z) $[M+2H]^{2+}$ calculated for $C_{62}H_{79}N_{11}O_8S^{2+}$: 568.7912, found: 568.7909.

$^1H$ NMR (600 MHz, DMSO-$d_6$) δ (ppm): 8.99 (s, 1H), 8.38 (d, *J* = 7.8 Hz, 1H), 8.30 (s, 1H), 8.28 (d, *J* = 14.2 Hz, 1H), 7.82 (d, *J* = 9.0 Hz, 2H), 7.69 (d, *J* = 9.3 Hz, 1H), 7.43 (d, *J* = 8.2 Hz, 2H), 7.38 (d, *J* = 8.3 Hz, 2H), 7.28 – 7.23 (m, 4H), 7.17 – 7.11 (m, 3H), 4.92 (t, *J* = 7.2 Hz, 1H), 4.50 (d, *J* = 9.3 Hz, 1H), 4.42 (t, *J* = 8.1 Hz, 1H), 4.28 (tt, *J* = 4.5, 2.5 Hz, 1H), 4.07 – 3.97 (m, 2H), 3.94 (d, *J* = 3.6 Hz, 1H), 3.70 – 3.60 (m, 6H), 3.57 (d, *J* = 10.8 Hz, 1H), 3.25 – 3.15 (m, 6H), 2.86 (q, *J* = 8.4, 5.1 Hz, 1H), 2.66 – 2.57

(m, 2H), 2.45 (s, 3H), 2.03 – 1.98 (m, 1H), 1.79 (ddd, *J* = 12.8, 8.4, 4.5 Hz, 2H), 1.70 (q, *J* = 14.8, 11.8 Hz, 3H), 1.57 – 1.29 (m, 13H), 1.20 (dd, *J* = 7.0, 1.3 Hz, 3H), 0.94 (s, 9H).

$^{13}$C NMR (151 MHz, DMSO-*d*$_6$) δ (ppm): 174.83, 172.81, 171.96, 169.65, 169.10 (“d”, *J* = 2.7 Hz), 156.91 (“d”, *J* = 8.0 Hz), 151.90 (“d”, *J* = 3.3 Hz), 151.48, 150.89, 150.57 (“d”, *J* = 2.4 Hz), 147.70, 146.63 (“d”, *J* = 16.4 Hz), 139.51, 135.61, 131.17, 129.63, 128.86, 128.64, 128.21 (“d”, *J* = 4.8 Hz), 128.08, 127.43, 126.90 (“d”, *J* = 4.0 Hz), 125.94 (“d”, *J* = 5.6 Hz), 122.88, 106.26, 69.12 (“d”, *J* = 7.2 Hz), 68.87, 58.71, 56.20 (“d”, *J* = 42.5 Hz), 52.99, 45.12, 42.72, 42.46, 41.64, 41.44, 41.00 (“d”, *J* = 16.5 Hz), 40.43, 40.23 (“d”, *J* = 6.0 Hz), 38.51, 37.94, 36.92, 36.12 (“d”, *J* = 31.1 Hz), 35.44, 35.38, 34.90 (“d”, *J* = 16.4 Hz), 34.18 (“d”, *J* = 19.5 Hz), 29.97, 28.86, 28.33, 26.36, 21.99 (“d”, *J* = 29.9 Hz), 15.97, 15.94.

##### Synthesis of (2*S*,4*R*)-4-hydroxy-1-((*S*)-2-(4-(1'-(4-(5-((4-hydroxy-1-((*R*)-3-phenylbutanoyl)-piperidin-4-yl)methyl)-4-oxo-4,5-dihydro-1*H*-pyrazolo[3,4-*d*]pyrimidin-1-yl)phenyl)-[4,4'-bipiperidine]-1-carbonyl)cubane-1-carboxamido)-3,3-dimethylbutanoyl)-*N*-((*S*)-1-(4-(4-methylthiazol-5-yl)phenyl)ethyl)pyrrolidine-2-carboxamide (NK266)

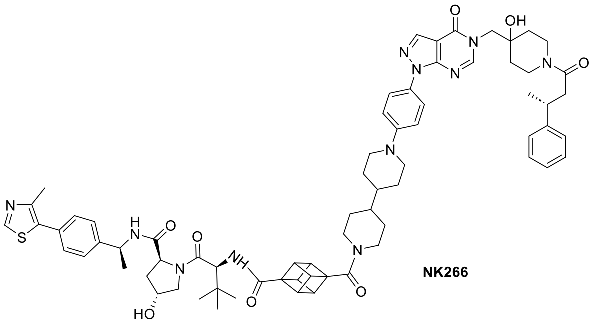

NK266

NK265_2 (8.8 mg, 0.01 mmol, 1.1 eq), HATU (6.6 mg, 0.02 mmol, 1.3 eq) and DIPEA (8.6 mg, 0.06 mmol, 5.0 eq) were dissolved in DMSO (1 mL) and stirred at 50 °C for 15 min. To this was added NK212 (10.0 mg, 0.01 mmol, 1.0 eq) and the mixture was stirred at 50 °C for 5 h. After completion, the crude product was directly purified by preparative reverse phase column chromatography (ACN/$H_2O$ + 0.1 % TFA). After evaporation of the solvents, (2*S*,4*R*)-4-hydroxy-1-((*S*)-2-(4-(1'-(4-(5-((4-hydroxy-1-((*R*)-3-phenylbutanoyl)-piperidin-4-yl)methyl)-4-oxo-4,5-dihydro-1*H*-pyrazolo[3,4-*d*]pyrimidin-1-yl)phenyl)-[4,4'-bipiperidine]-1-carbonyl)cubane-1-carboxamido)-3,3-dimethylbutanoyl)-*N*-((*S*)-1-(4-(4-methylthiazol-5-yl)phenyl)ethyl)pyrrolidine-2-carboxamide (NK266) (12.2 mg, 0.01 mmol, 74 % yield) was obtained as an off-white solid.

HRMS (m/z) [M+2H]$^{2+}$ calculated for $C_{70}H_{85}N_{11}O_8S^{2+}$: 619.8146, found: 619.8140.

$^{1}$H NMR (600 MHz, DMSO-*d*$_6$) δ (ppm): 8.98 (s, 1H), 8.39 (d, *J* = 7.8 Hz, 1H), 8.30 – 8.25 (m, 2H), 7.77 (d, *J* = 8.5 Hz, 2H), 7.58 (dd, *J* = 11.3, 9.3 Hz, 1H), 7.43 (d, *J* = 8.3 Hz, 2H), 7.38 (d, *J* = 8.3 Hz, 2H), 7.26 (dd, *J* = 8.6, 6.1 Hz, 4H), 7.15 (ddd, *J* = 8.6, 6.2, 2.6 Hz, 1H), 7.10 (d, *J* = 8.5 Hz, 2H), 4.95 – 4.86 (m, 2H), 4.57 (d, *J* = 9.2 Hz, 1H), 4.43 (t, *J* = 8.1 Hz, 1H), 4.35 (d, *J* = 12.6 Hz, 1H), 4.31 – 4.26 (m, 1H), 4.13 – 4.07 (m, 6H), 4.07 – 3.92 (m, 3H), 3.81 (d, *J* = 11.8 Hz, 2H), 3.69 – 3.56 (m, 3H), 3.44 (d, *J* = 11.4 Hz, 1H), 3.23 – 3.13 (m, 2H), 3.03 (t, *J* = 12.6 Hz, 1H), 2.86 (q, *J* = 8.6, 5.4 Hz, 1H), 2.69 (s, 2H), 2.65 – 2.51 (m, 2H), 2.45 (s, 3H), 2.05 – 2.00 (m, 1H), 1.79 (td, *J* = 8.0, 4.3 Hz, 4H), 1.73 (d, *J* = 12.4 Hz, 1H), 1.57 – 1.39 (m, 2H), 1.37 (d, *J* = 7.0 Hz, 3H), 1.33 – 1.22 (m, 6H), 1.20 (dd, *J* = 7.0, 1.4 Hz, 3H), 1.16 – 1.01 (m, 2H), 0.95 (s, 9H).

$^{13}$C NMR (151 MHz, DMSO-*d*$_6$) δ (ppm): 171.06, 170.71, 169.82, 169.58 (“d”, *J* = 2.8 Hz), 168.63, 157.40 (“d”, *J* = 7.8 Hz), 152.31, 151.96, 150.96, 148.22, 147.11 (“d”, *J* = 16.4 Hz), 145.08, 135.98, 131.58, 130.17, 129.30, 128.68 (“d”, *J* = 4.8 Hz), 127.37 (“d”, *J* = 3.8 Hz), 126.87, 126.41 (“d”, *J* = 5.6 Hz), 123.37, 116.07, 106.67, 69.59 (“d”, *J* = 7.1 Hz), 69.25, 59.12, 57.73, 56.82, 56.67, 56.47, 53.46, 49.32, 48.13, 46.41 (“d”, *J* = 15.4 Hz), 45.15, 41.94, 41.47 (“d”, *J* = 16.7 Hz), 40.94, 40.71 (“d”, *J* = 6.0 Hz), 38.18, 37.39, 36.59 (“d”, *J* = 31.2 Hz), 36.03, 35.37 (“d”, *J* = 17.1 Hz), 34.65 (“d”, *J* = 20.2 Hz), 30.18, 29.02 (“d”, *J* = 36.3 Hz), 26.97, 22.90, 22.46 (“d”, *J* = 29.8 Hz), 16.45.

#### 3.11 Synthesis of a VHL inhibitor

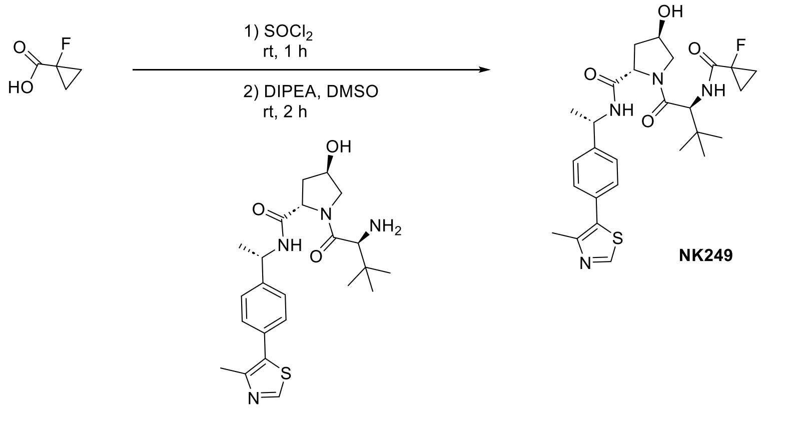

76 %

**Scheme 11.** Synthesis of VHL inhibitor NK249.

**Synthesis of (2*S*,4*R*)-1-((*S*)-2-(1-fluorocyclopropane-1-carboxamido)-3,3-dimethylbutanoyl)-4-hydroxy-*N*-((*S*)-1-(4-(4-methylthiazol-5-yl)phenyl)ethyl)pyrrolidine-2-carboxamide (NK249)**

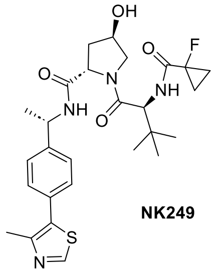

**NK249**

1-Fluorocyclopropane-1-carboxylic acid (5.6 mg, 0.05 mmol, 1.3 eq), was dissolved in $SOCl_2$ (1 mL) and stirred at rt for 1 h. After completion, the solvent was evaporated under reduced pressure. The crude product was dissolved in DMSO (1 mL) and to this were added (2*S*,4*R*)-1-((*S*)-2-amino-3,3-dimethylbutanoyl)-4-hydroxy-*N*-((*S*)-1-(4-(4-methylthiazol-5-yl)phenyl)ethyl)-pyrrolidine-2-carboxamide HCl salt (20.0 mg, 0.04 mmol, 1.0 eq) and DIPEA (58.1 mg, 0.45 mmol, 10.0 eq), and the mixture was stirred at rt for 2 h. After completion, the crude product was directly purified by preparative reverse phase column chromatography (ACN/$H_2O$ + 0.1 % TFA). After evaporation of the solvents under reduced pressure, the (2*S*,4*R*)-1-((*S*)-2-(1-fluorocyclopropane-1-carboxamido)-3,3-dimethylbutanoyl)-4-hydroxy-*N*-((*S*)-1-(4-(4-methylthiazol-5-yl)phenyl)ethyl)pyrrolidine-2-carboxamide (NK249) (18.2 mg, 0.03 mmol, 76 % yield) was obtained as a white powder.

HRMS (m/z) $[M+H]^+$ calculated for $C_{27}H_{36}FN_4O_4S^+$: 531.2436, found: 531.2442.

$^1H$ NMR (500 MHz, DMSO-$d_6$) δ (ppm): 8.99 (s, 1H), 8.47 (d, *J* = 7.8 Hz, 1H), 7.46 – 7.42 (m, 2H), 7.37 (d, *J* = 8.4 Hz, 2H), 7.25 (dd, *J* = 9.2, 2.9 Hz, 1H), 4.91 (p, *J* = 7.1 Hz, 1H), 4.58 (dd, *J* = 9.3, 1.2 Hz, 1H), 4.48 (dd, *J* = 8.9, 7.8 Hz, 1H), 4.28 (dt, *J* = 4.1, 2.0 Hz, 1H), 3.62 – 3.54 (m, 2H), 2.46 (s, 3H), 2.07 (ddt, *J* = 12.6, 7.8, 1.8 Hz, 1H), 1.77 (ddd, *J* = 13.2, 9.0, 4.5 Hz, 1H), 1.40 – 1.33 (m, 5H), 1.23 – 1.20 (m, 2H), 0.97 (s, 9H).

$^{13}C$ NMR (126 MHz, DMSO-$d_6$) δ (ppm): 170.86, 169.23, 168.47 (d, *J* = 20.4 Hz), 151.98, 148.19, 145.20, 131.61, 130.15, 129.32, 126.81, 78.61 (d, *J* = 232.5 Hz), 69.28, 59.11, 57.11, 57.03, 48.23, 38.19, 36.58, 26.70, 22.97, 16.45, 13.44 (d, *J* = 10.1 Hz), 13.16 (d, *J* = 10.1 Hz).

#### 4. NMR spectra of compounds

$^1H$ NMR spectra of NK188.

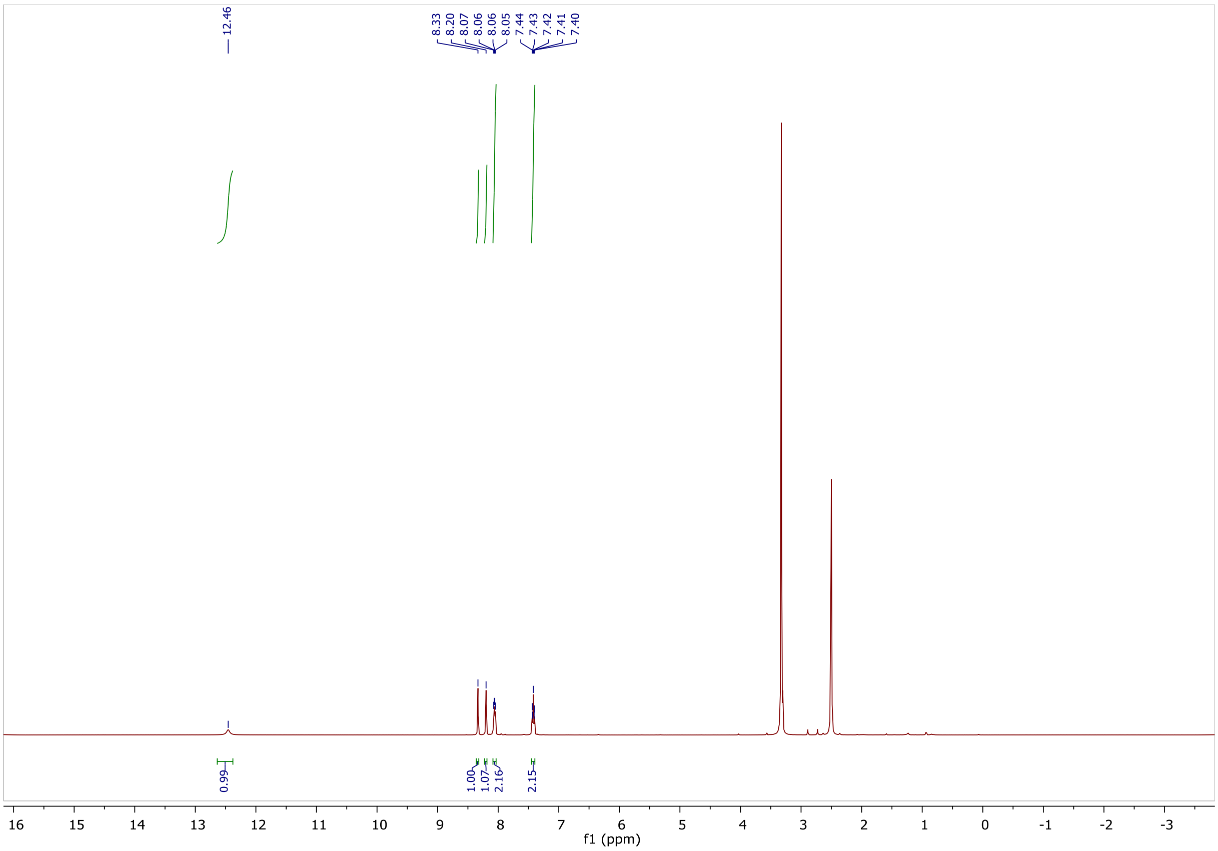

$^{13}C$ NMR spectra of NK188.

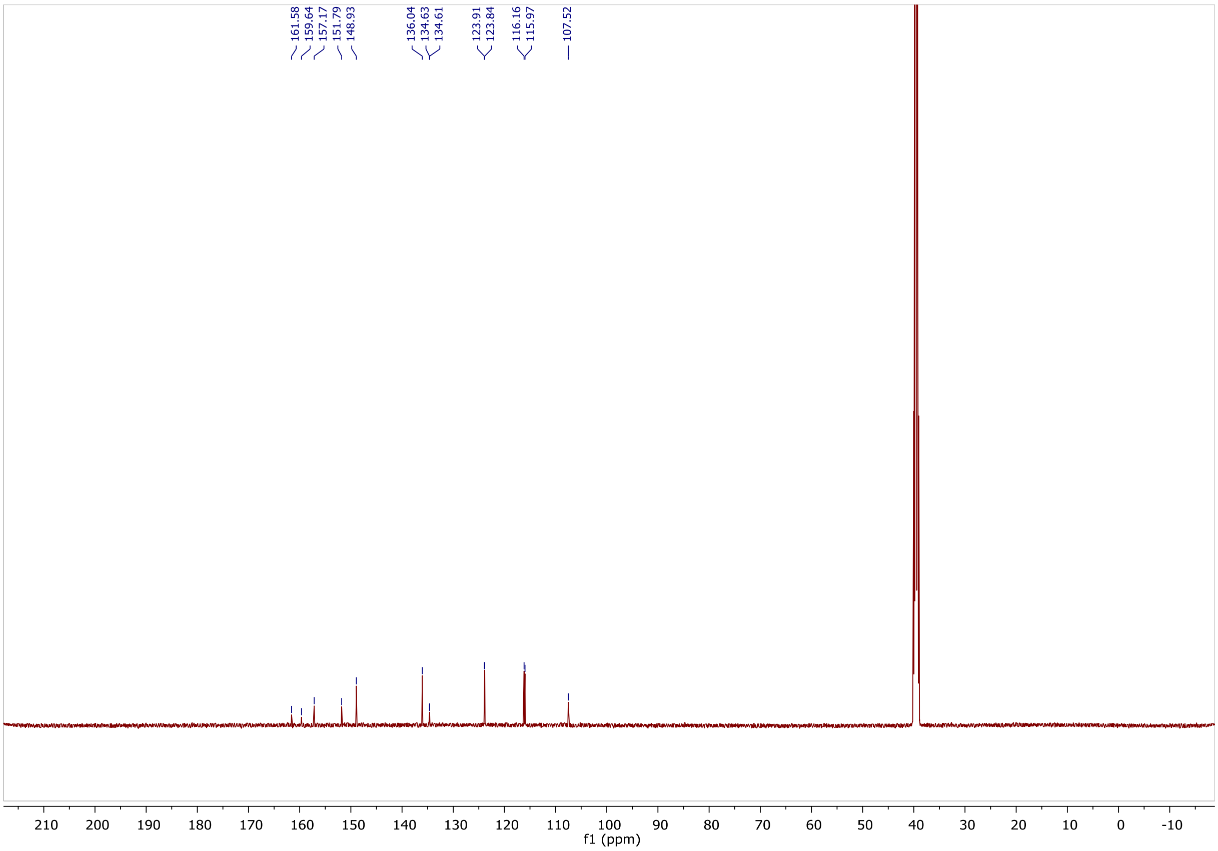

$^{1}H$ NMR spectra of NK189.

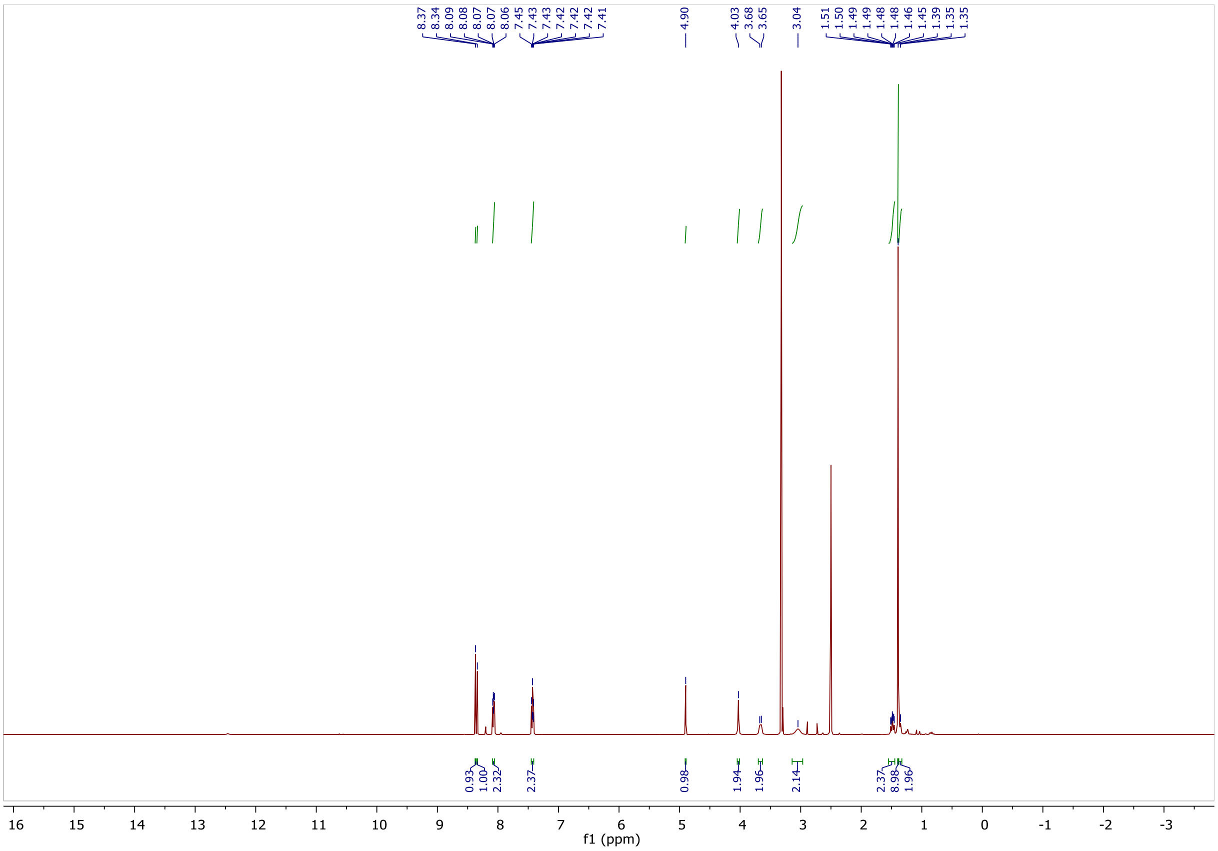

$^{13}C$ NMR spectra of NK189.

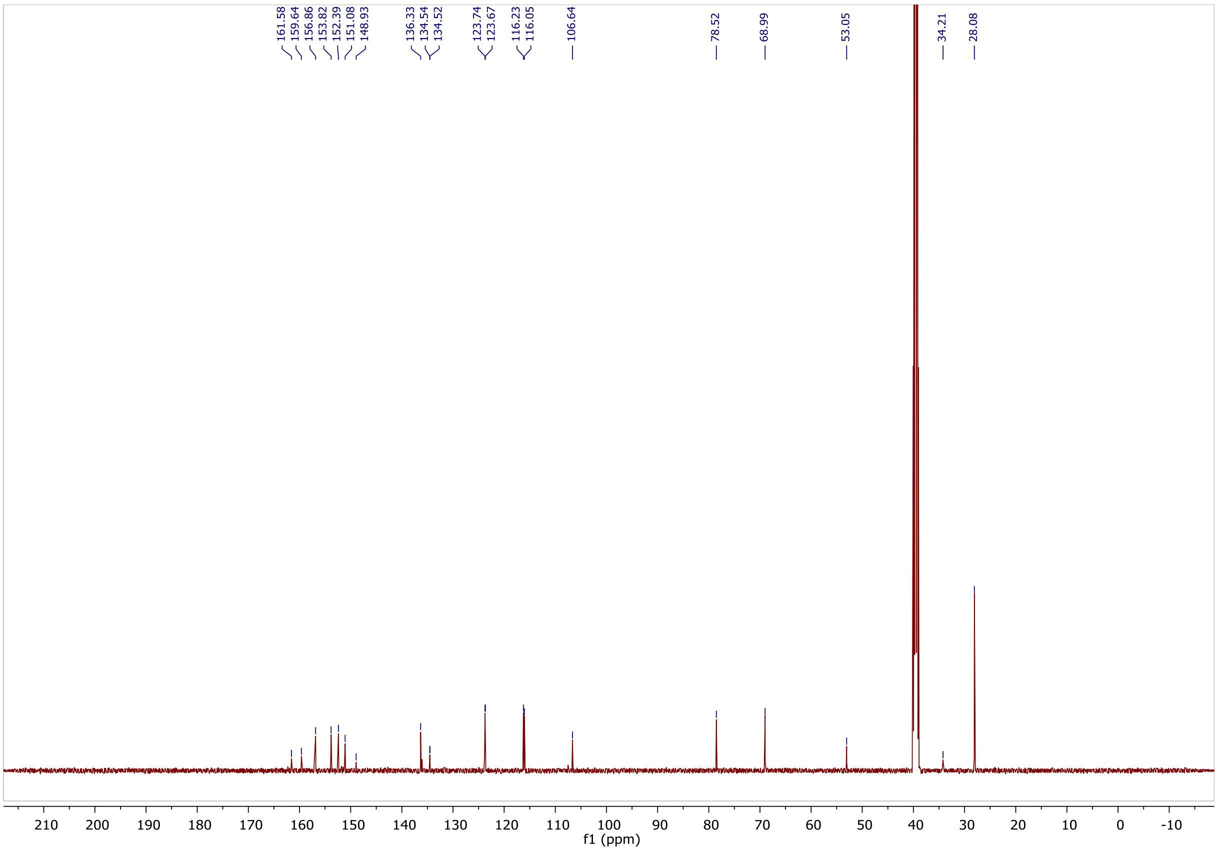

$^{1}H$ NMR spectra of NK192.

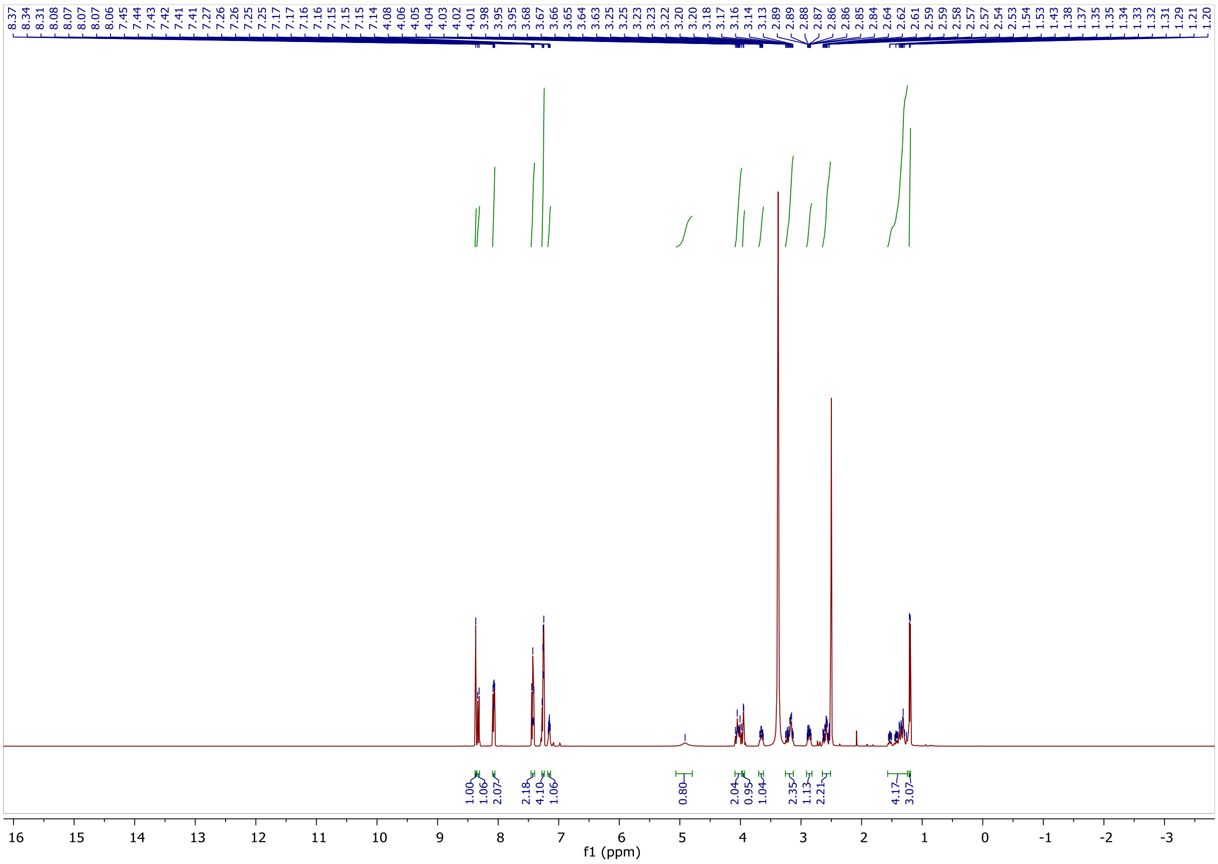

$^{13}C$ NMR spectra of NK192.

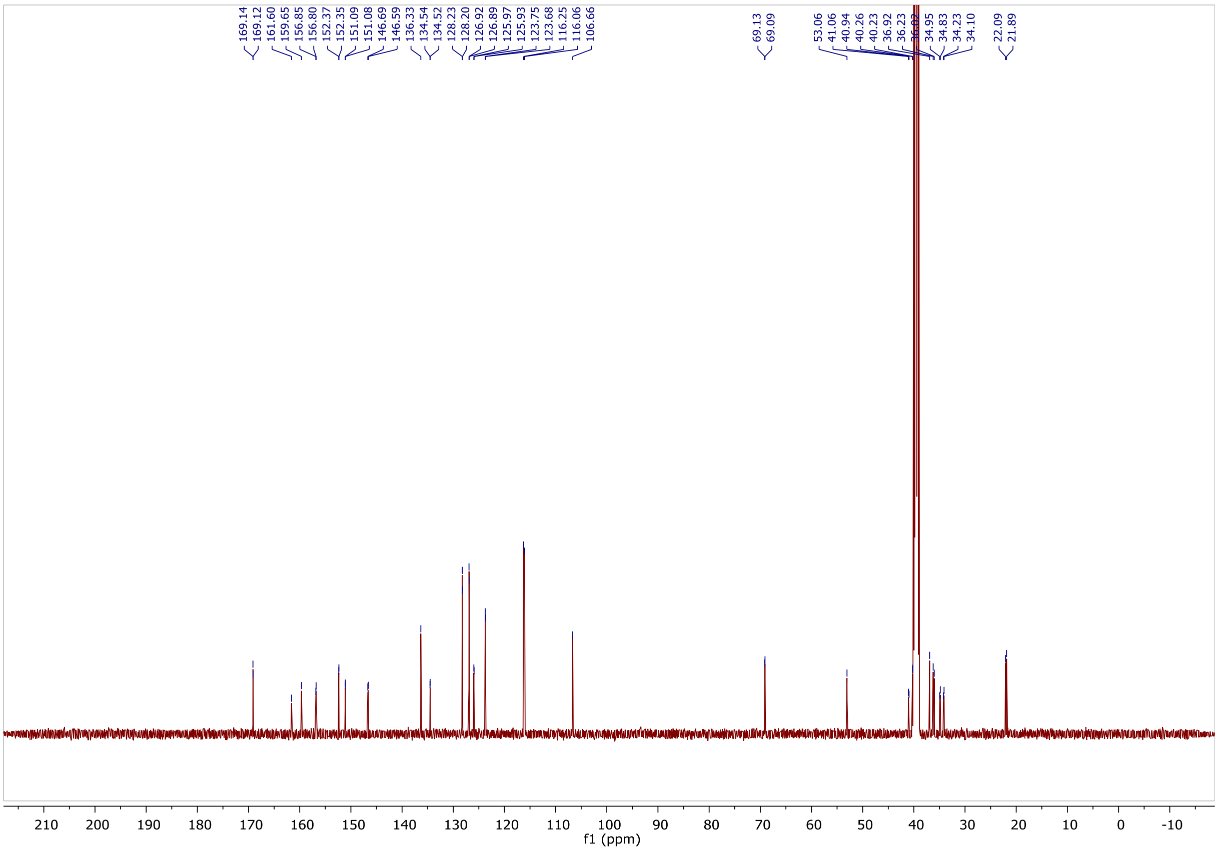

[1]H NMR spectra of NK135.

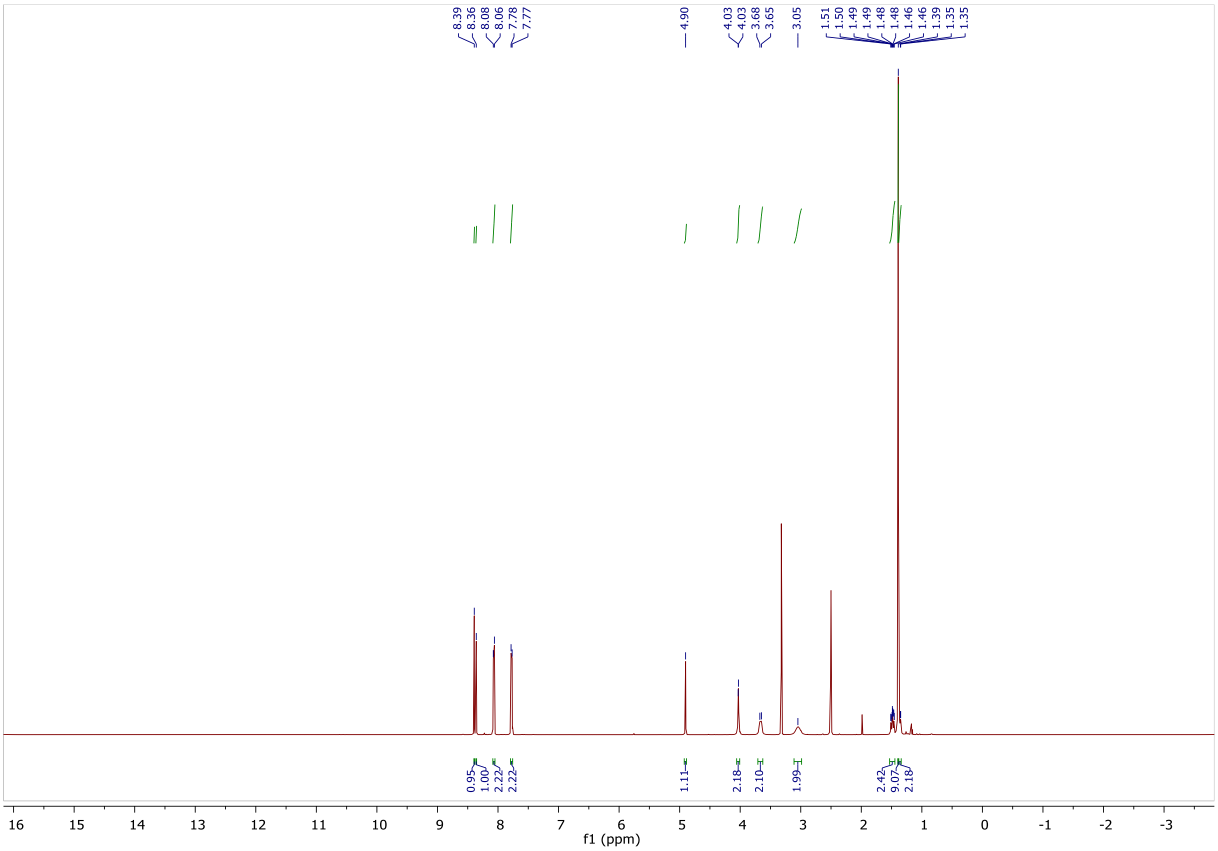

[13]C NMR spectra of NK135.

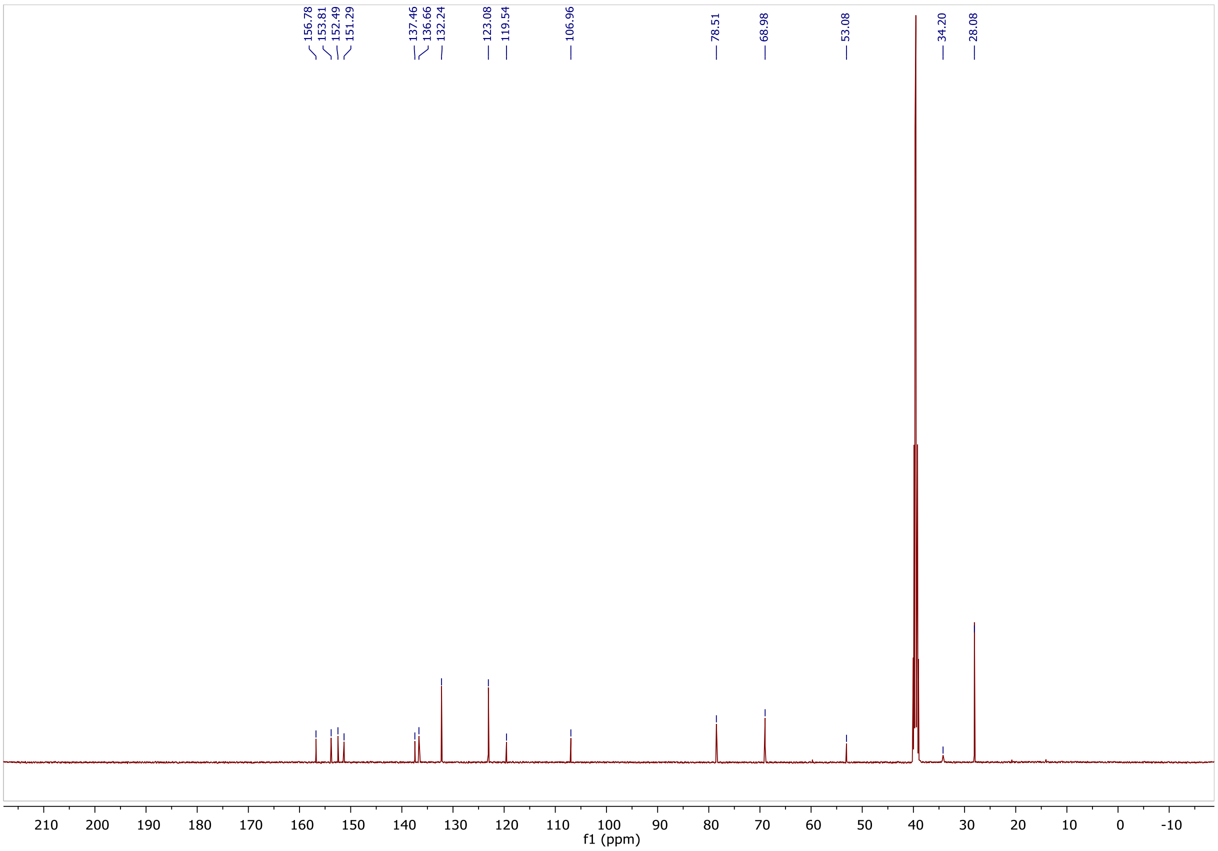

$^1H$ NMR spectra of NK137.

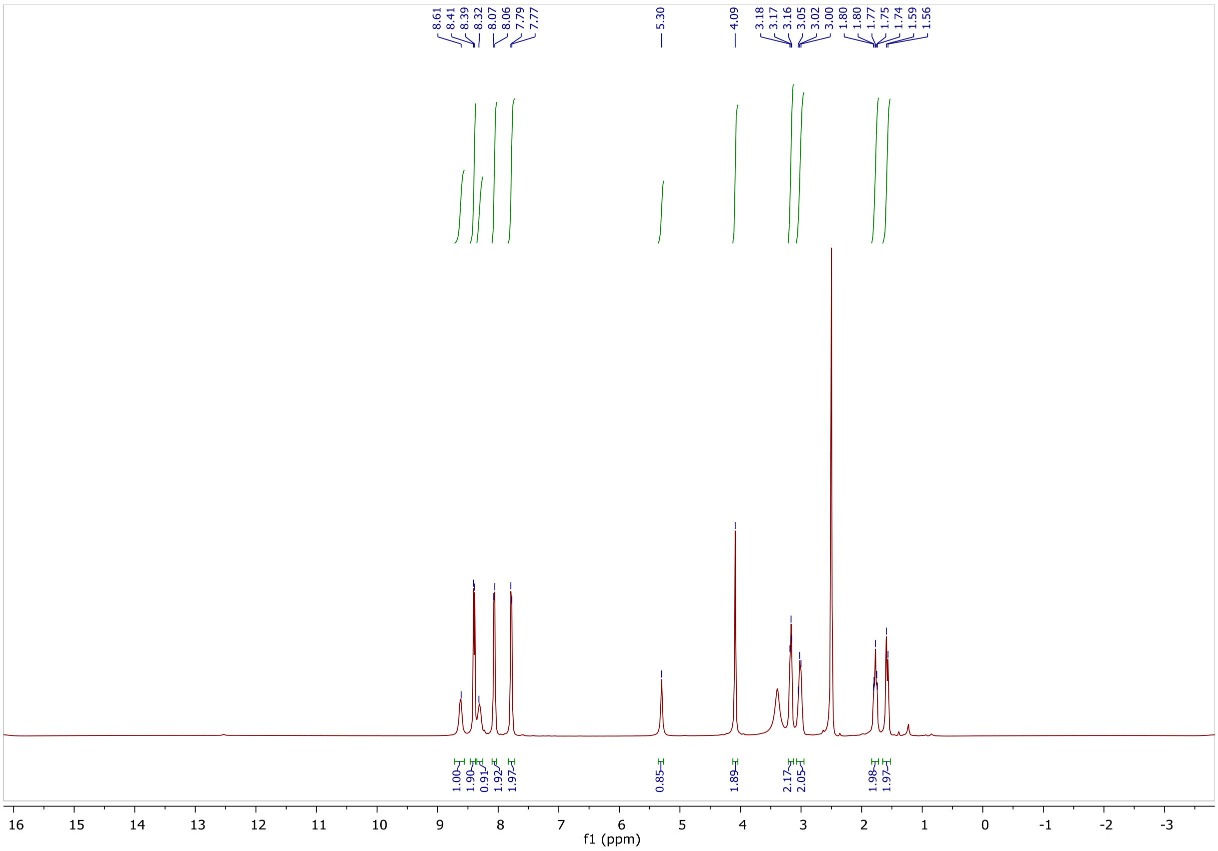

$^{13}C$ NMR spectra of NK137.

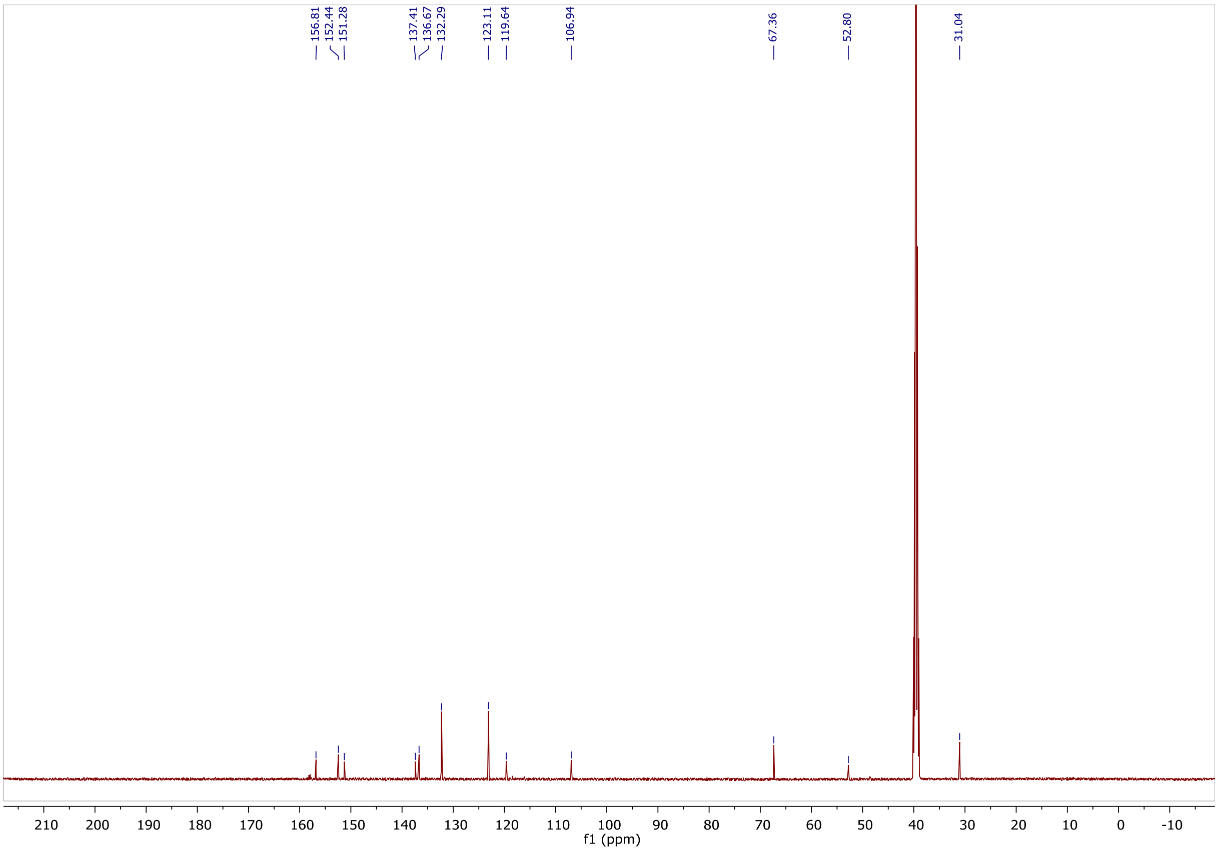

#### $^1$H NMR spectra of NK195.

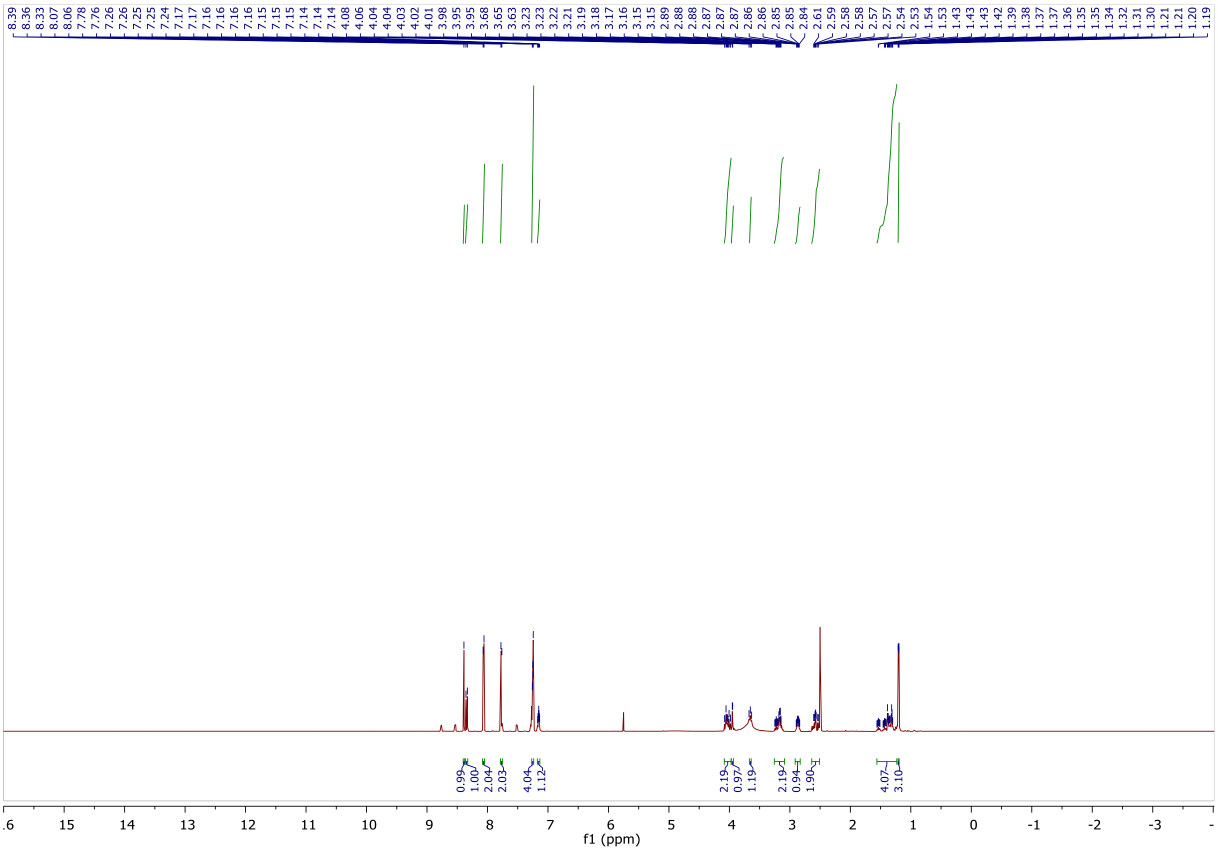

#### $^{13}$C NMR spectra of NK195.

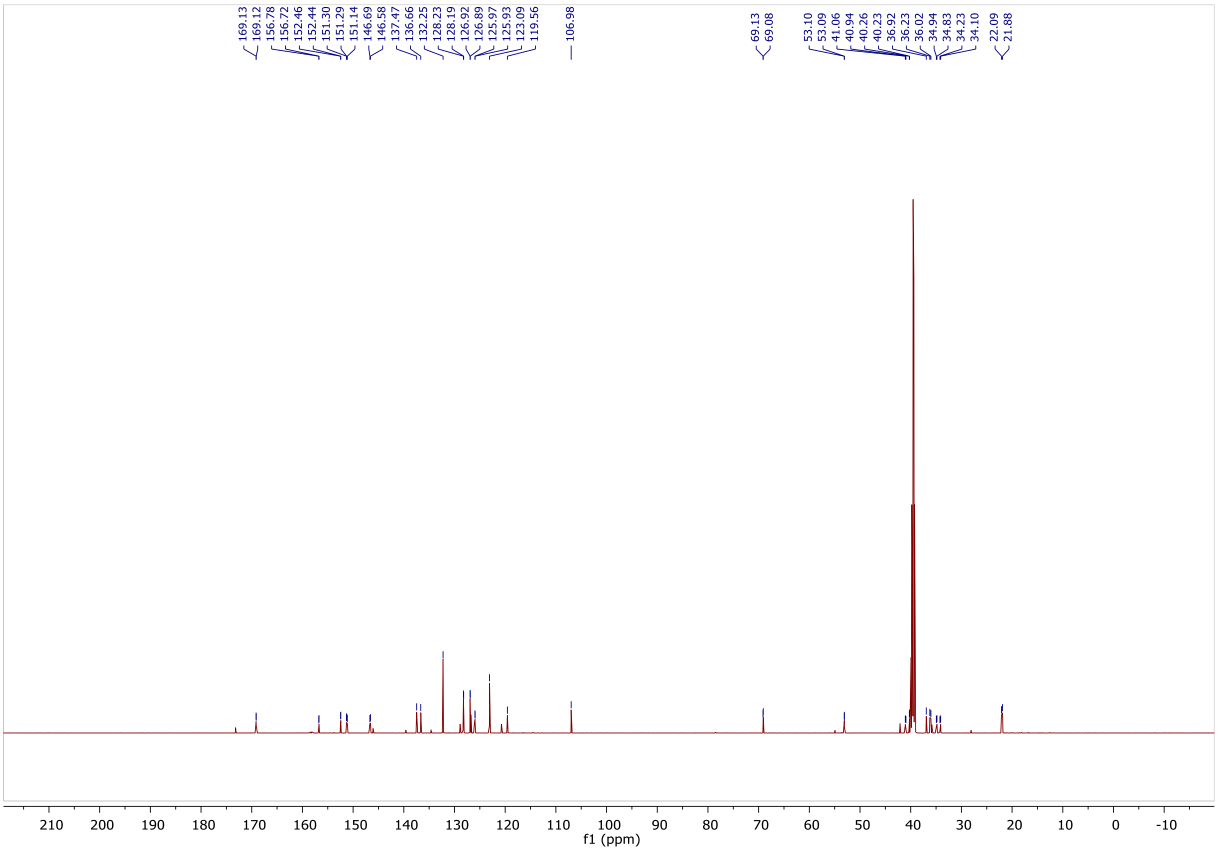

$^{1}H$ NMR spectra of NK210.

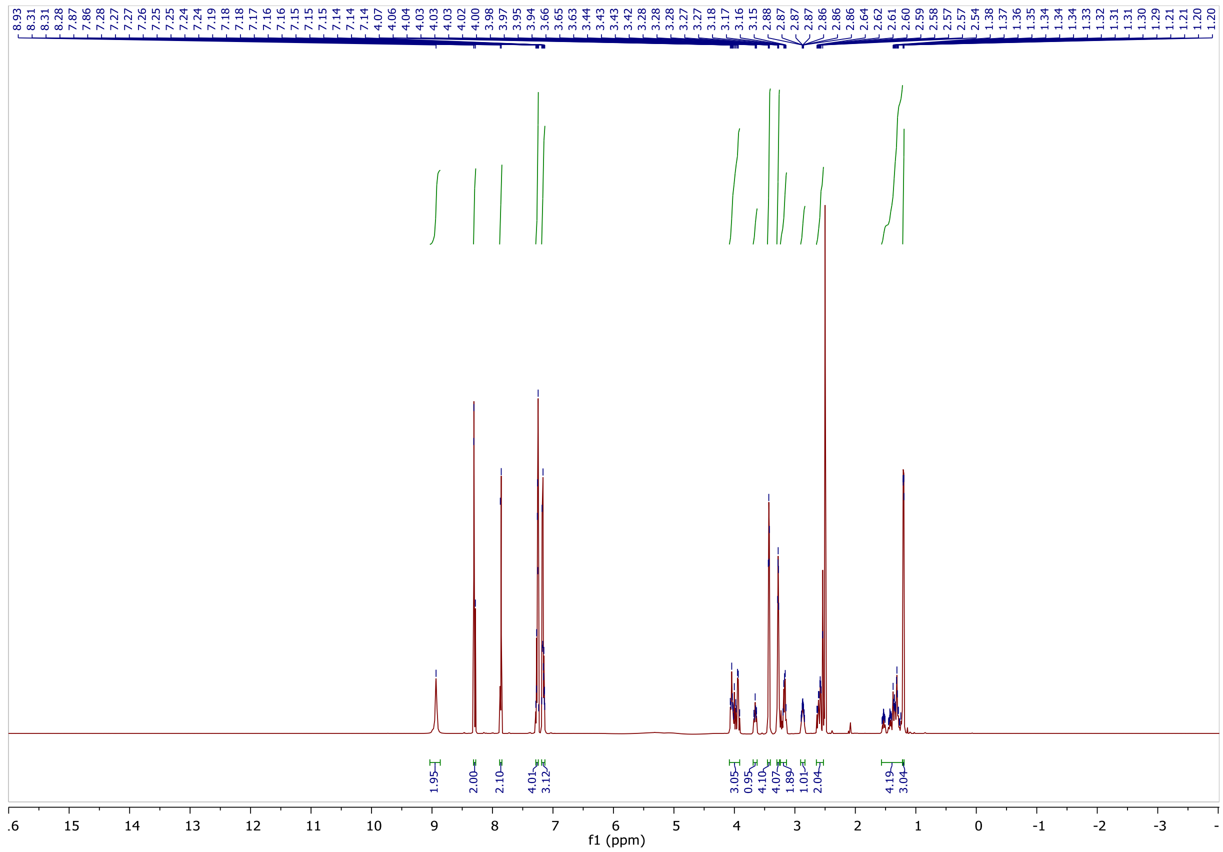

$^{13}C$ NMR spectra of NK210.

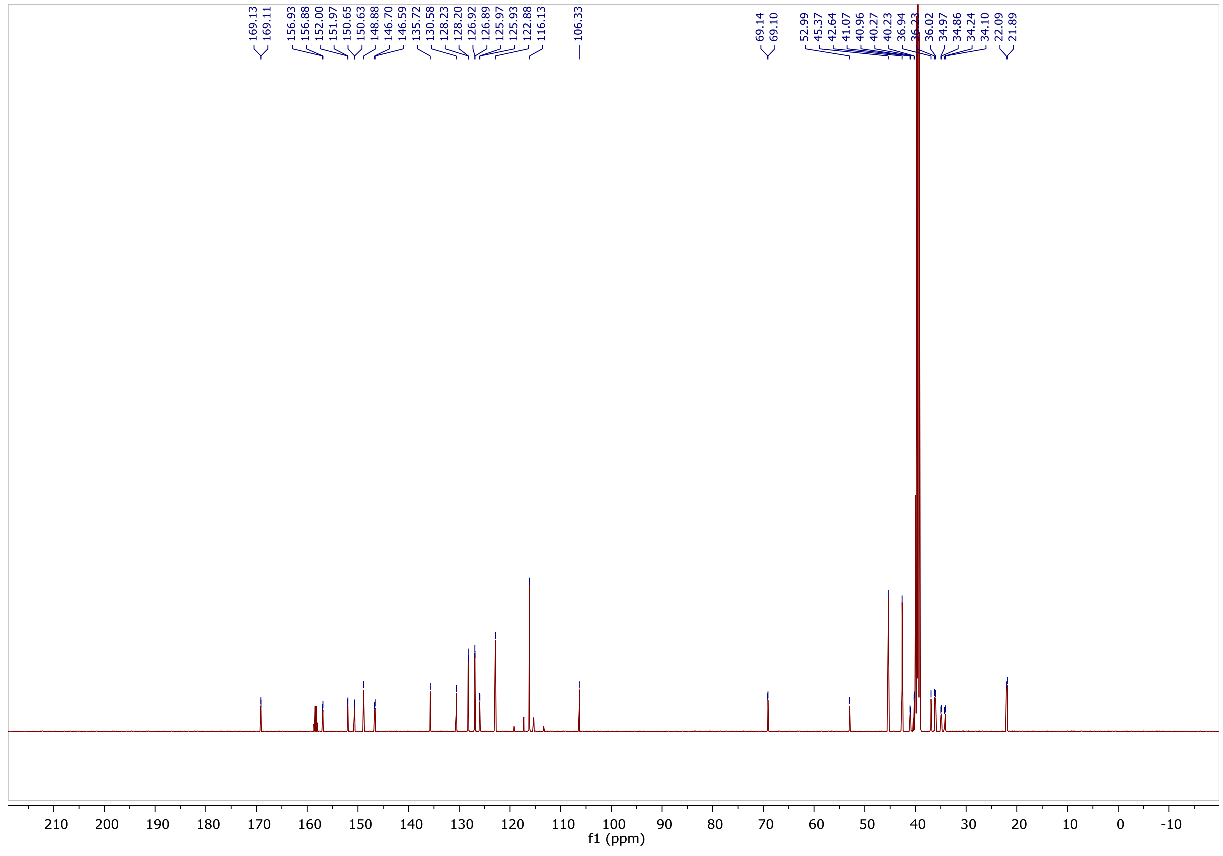

[1]H NMR spectra of NK212.

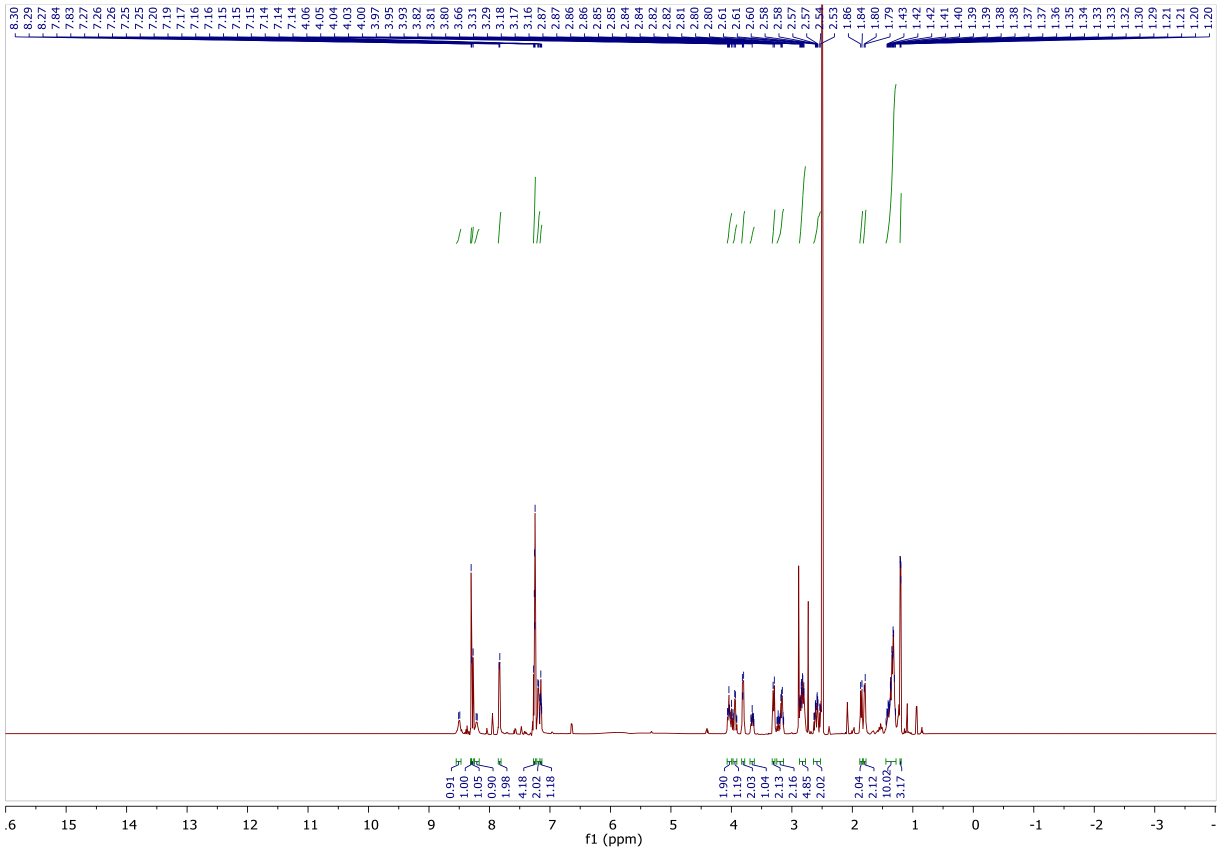

[13]C NMR spectra of NK212.

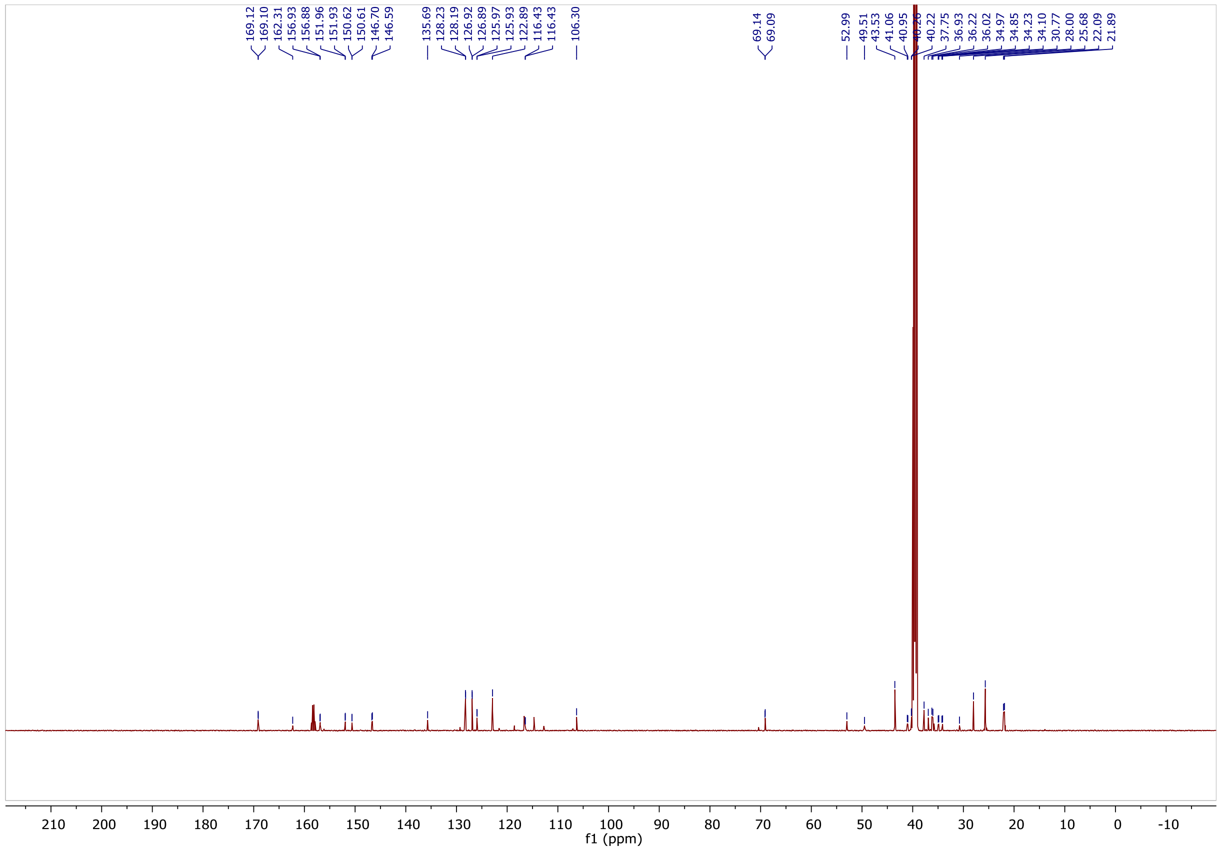

### [1]H NMR spectra of NK220.

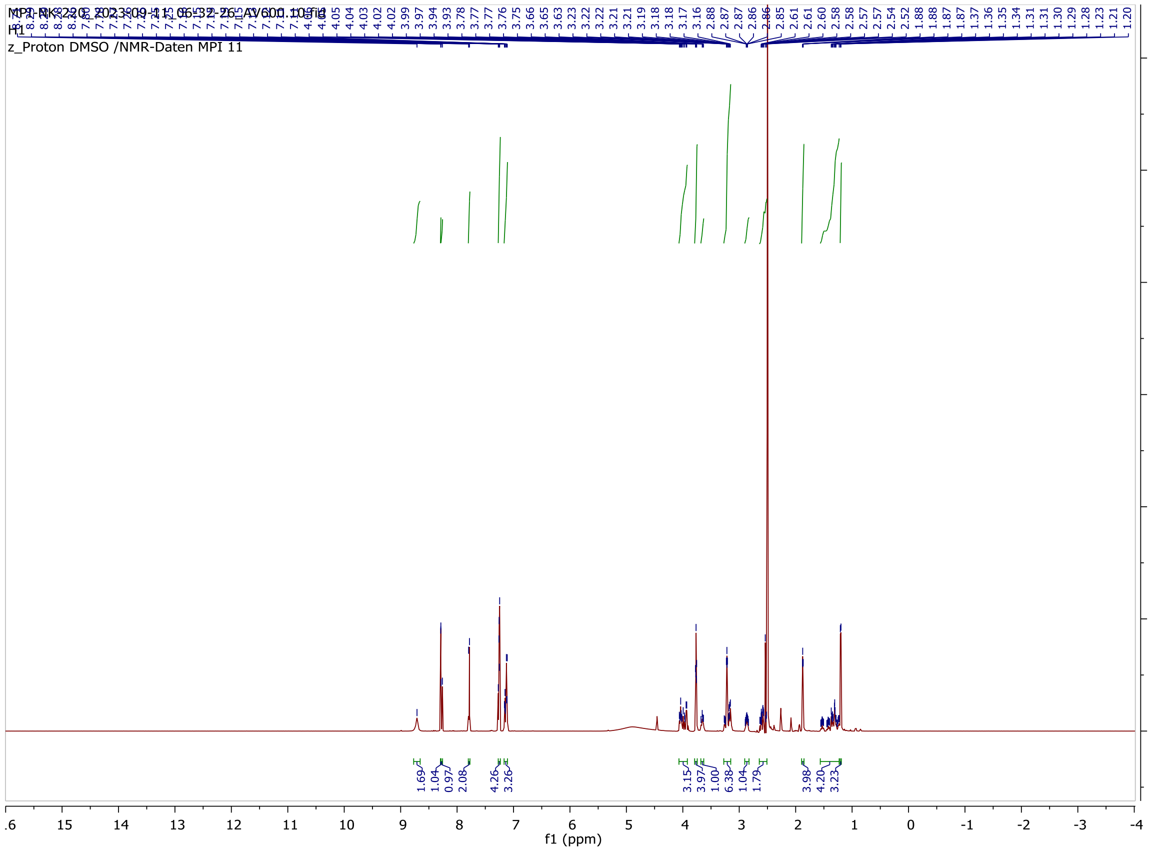

### [13]C NMR spectra of NK220.

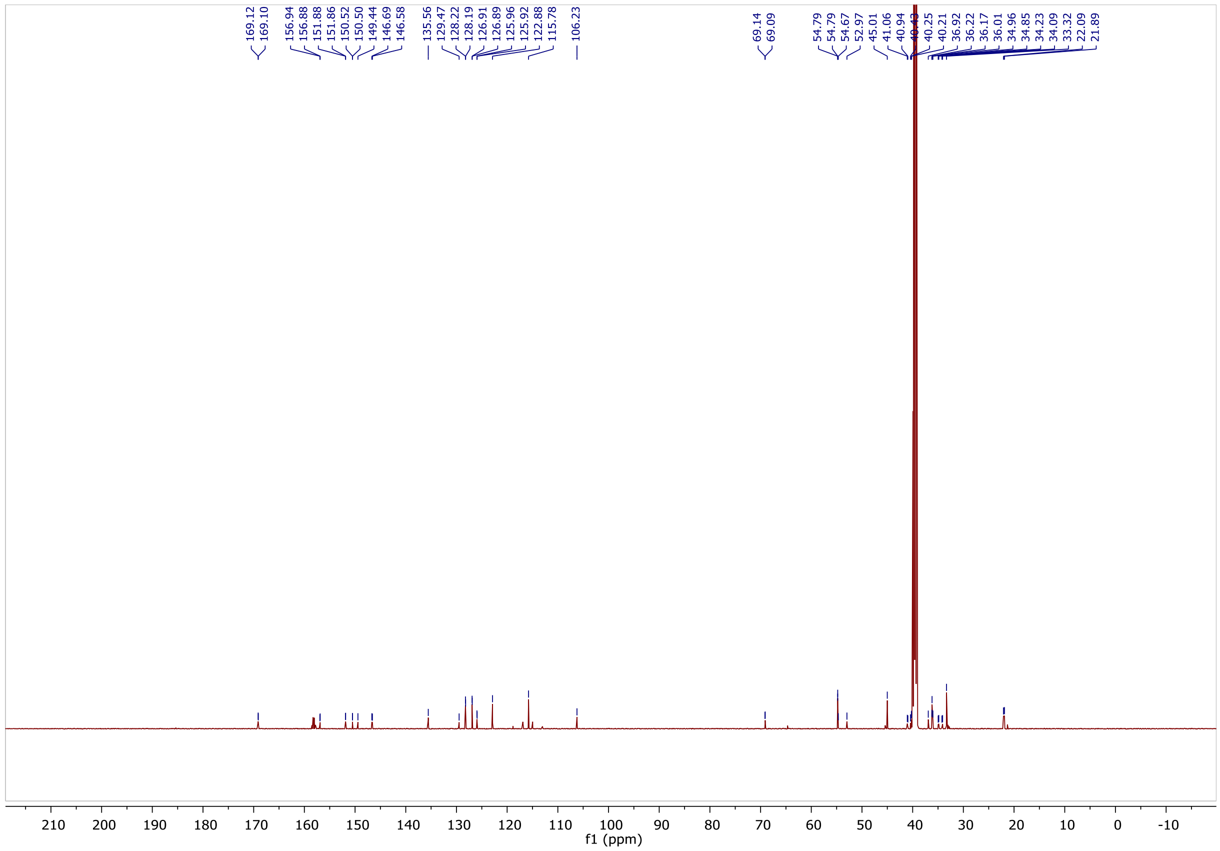

$^{1}$H NMR spectra of NK226.

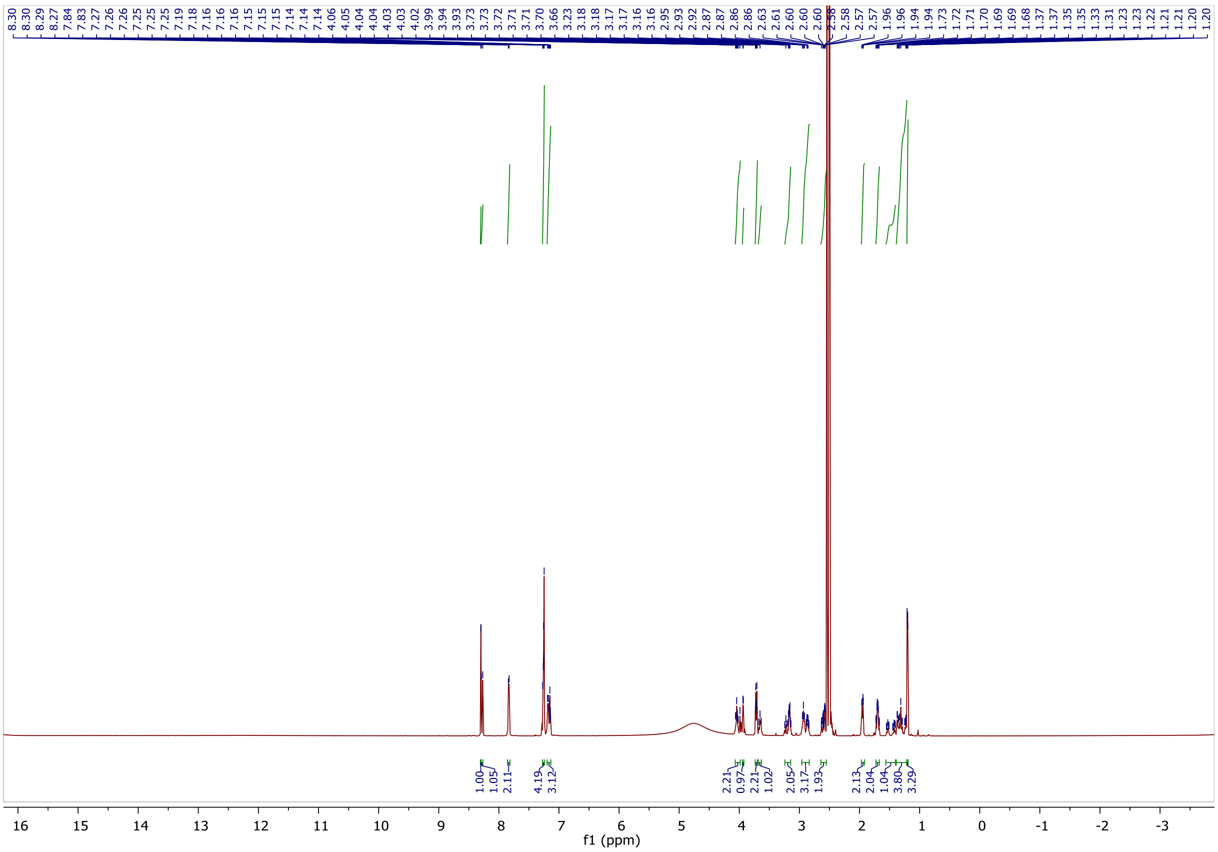

$^{13}$C NMR spectra of NK226.

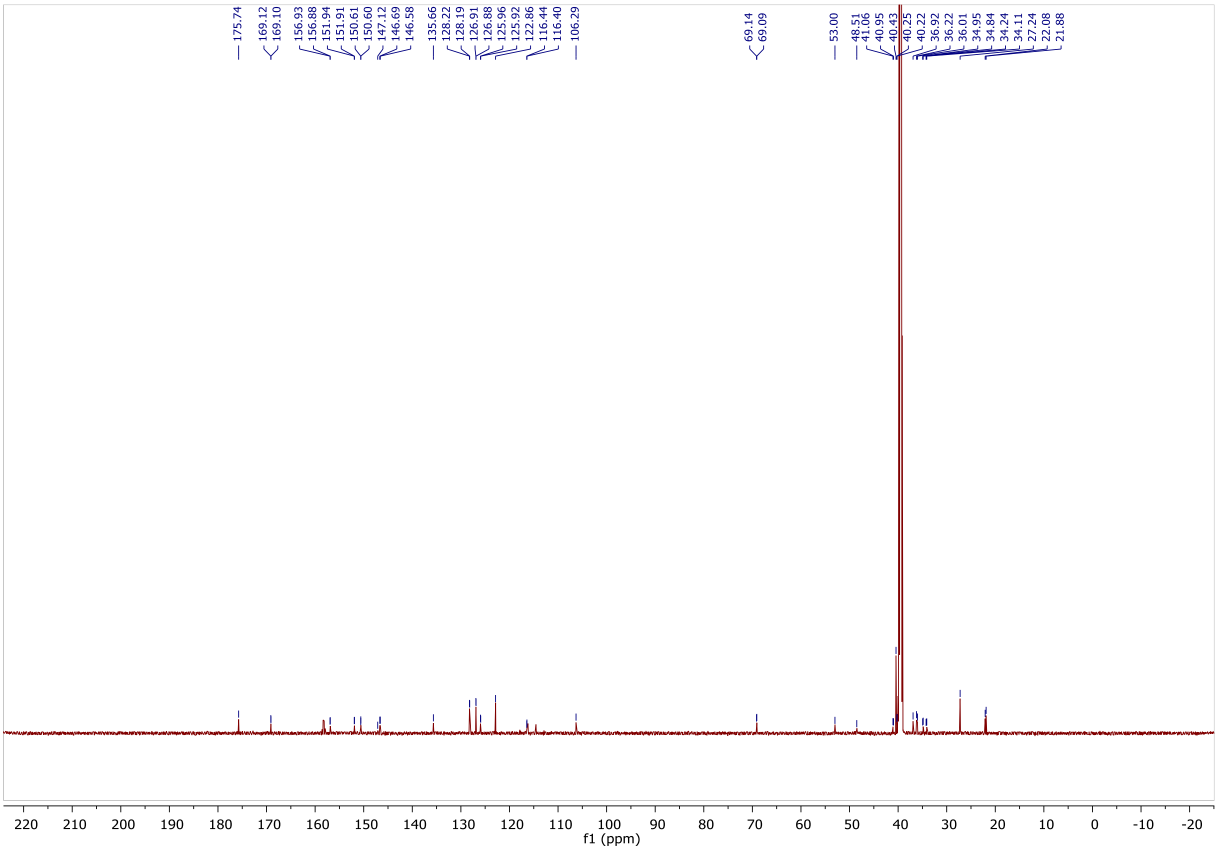

$^1H$ NMR spectra of NK231.

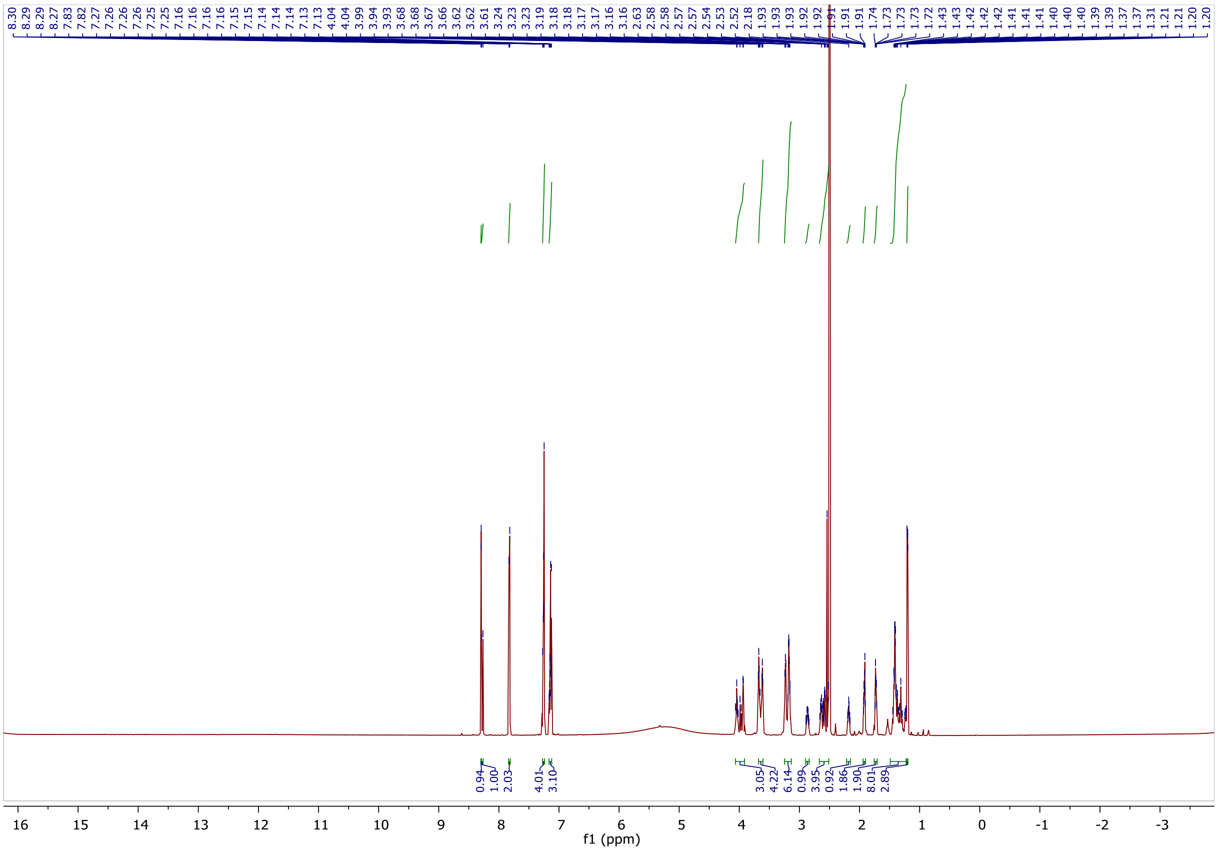

$^{13}C$ NMR spectra of NK231.

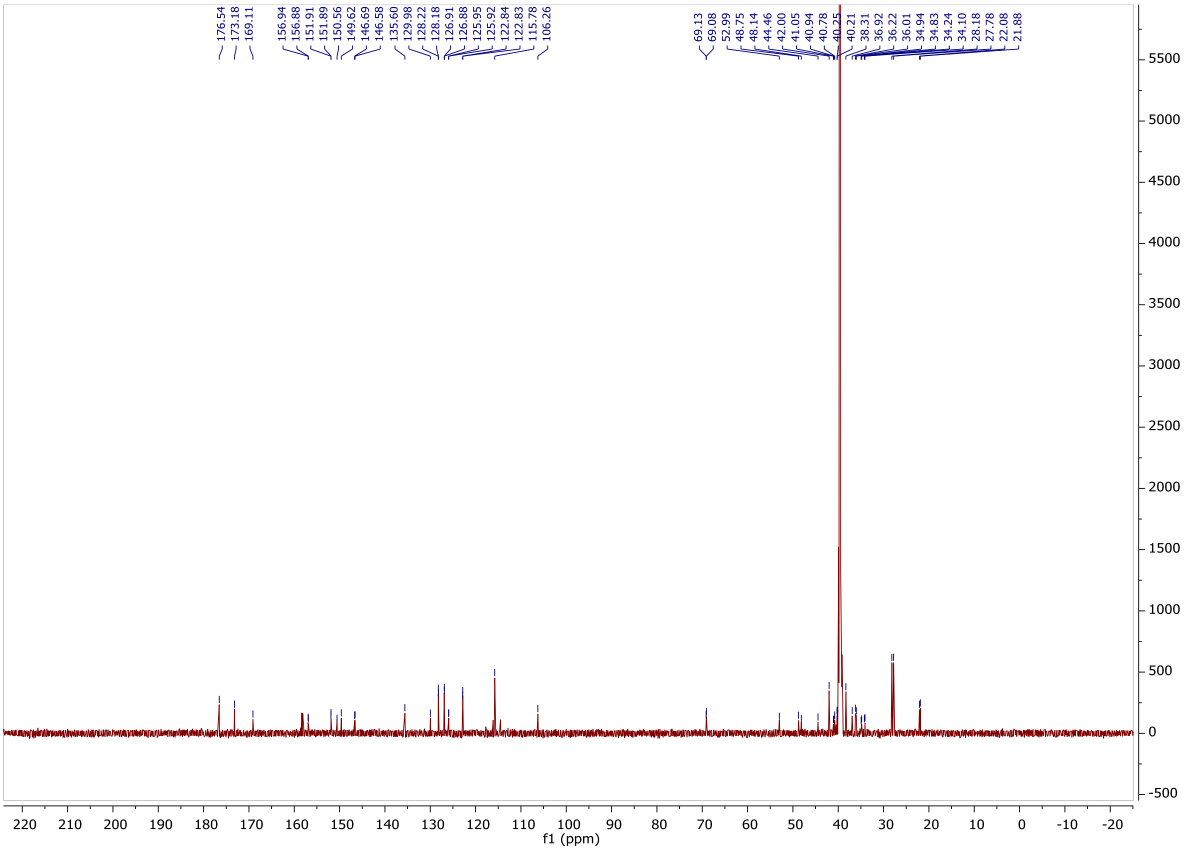

$^{1}H$ NMR spectra of NK267.

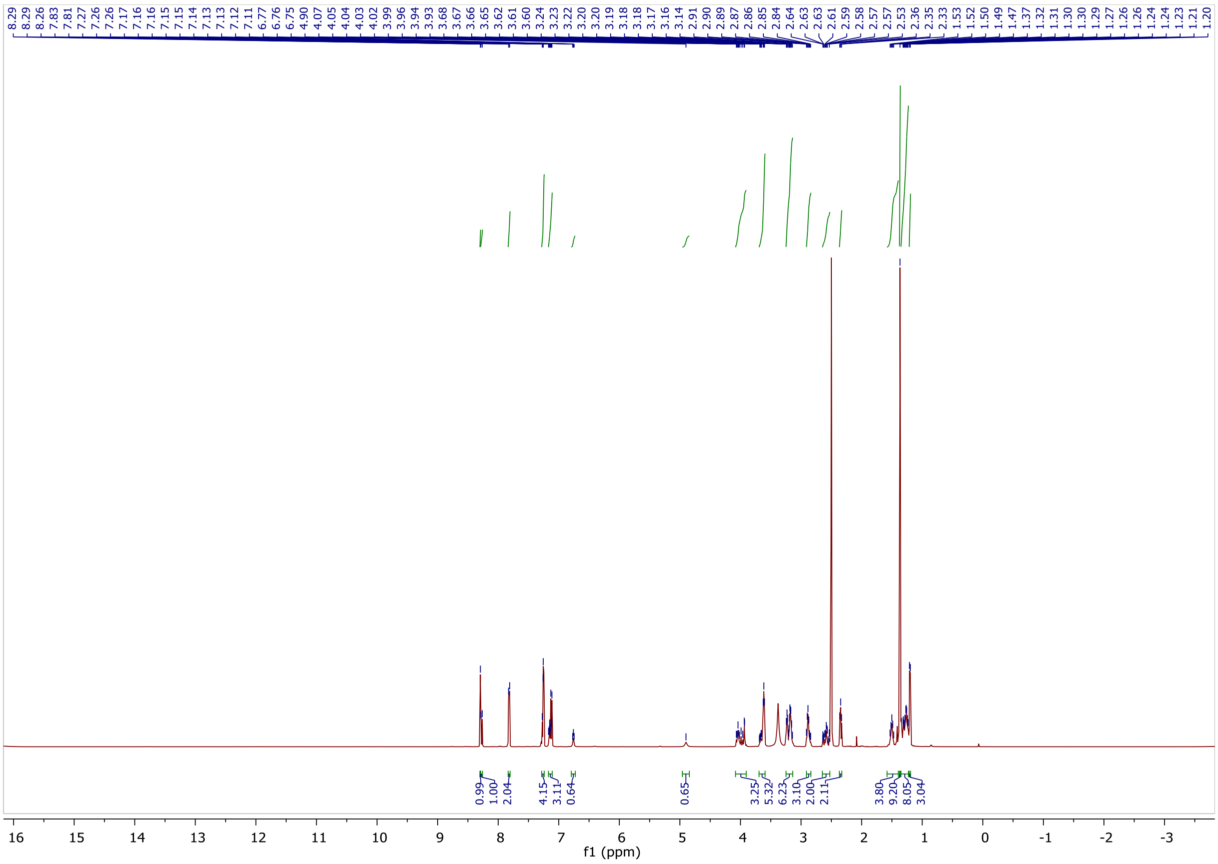

$^{13}C$ NMR spectra of NK267.

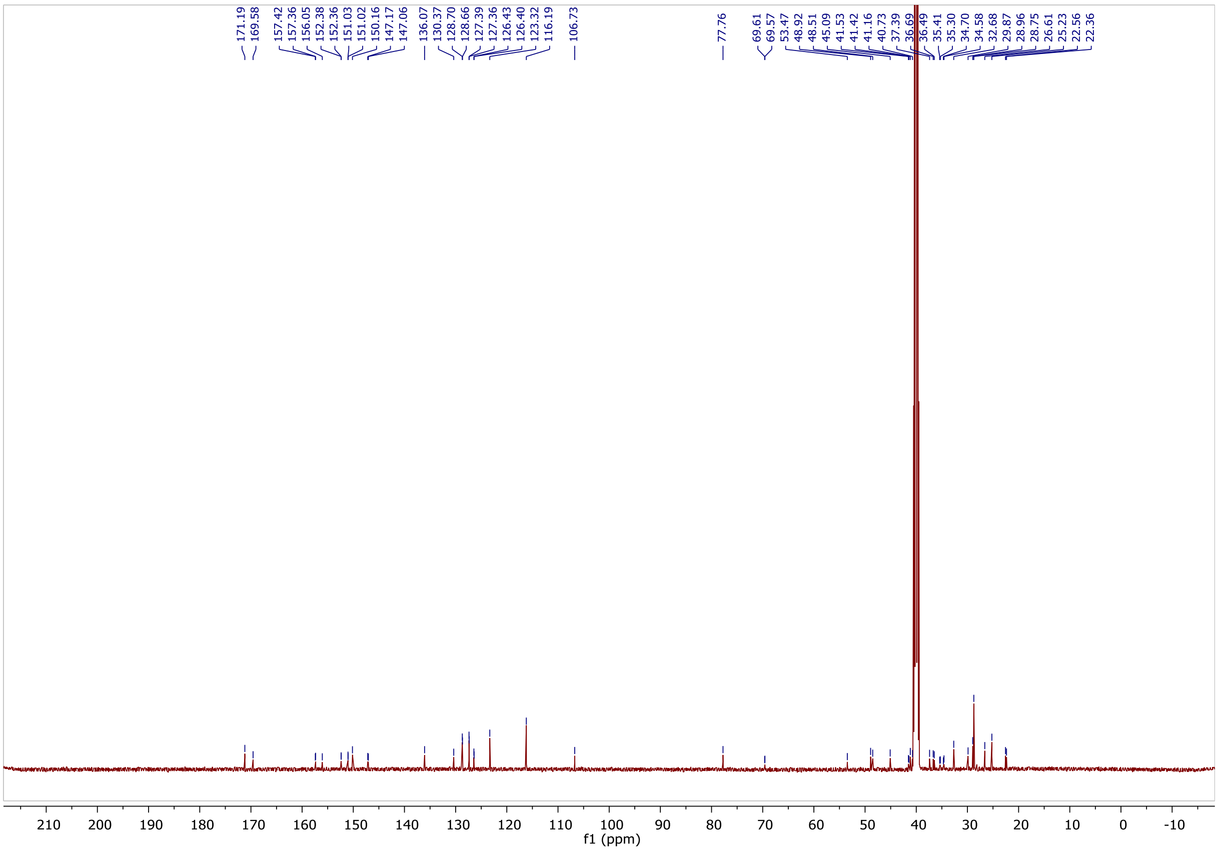

$^{1}$H NMR spectra of NK264.

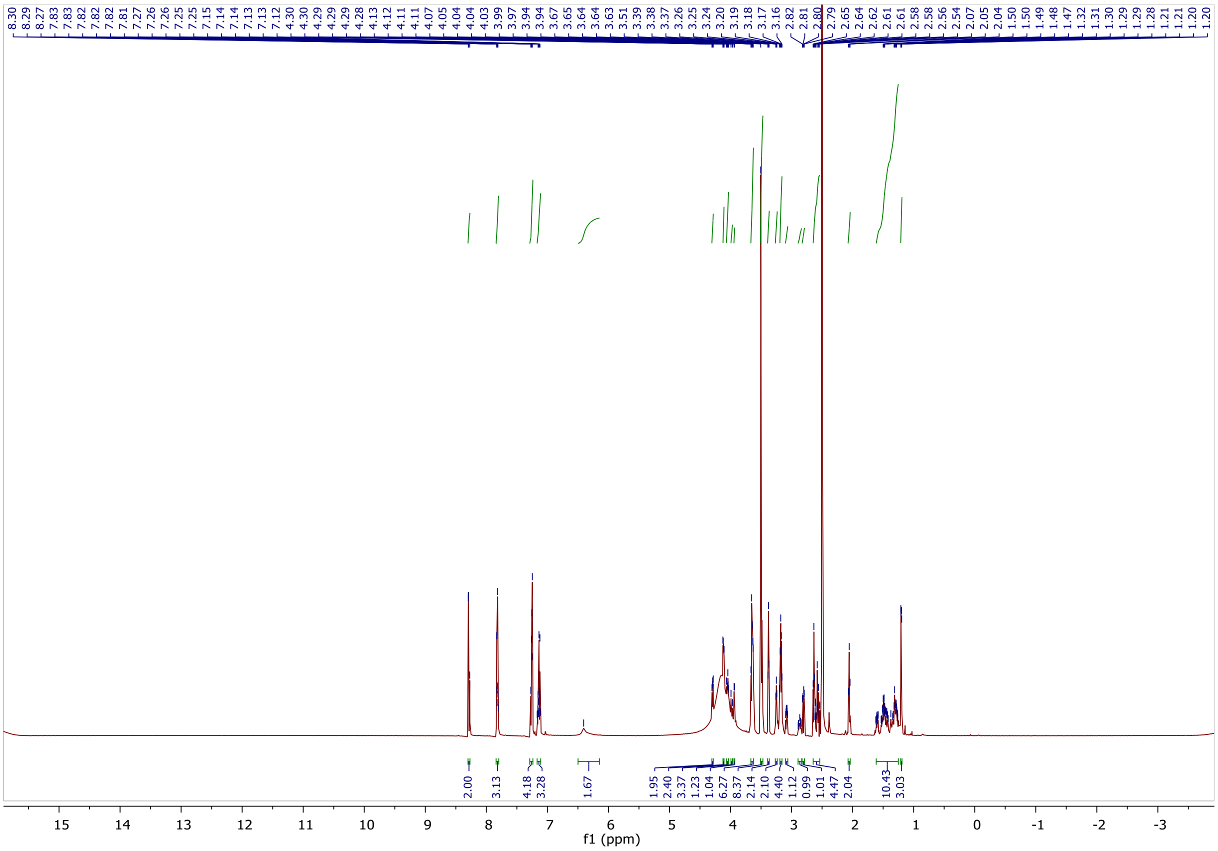

$^{13}$C NMR spectra of NK264.

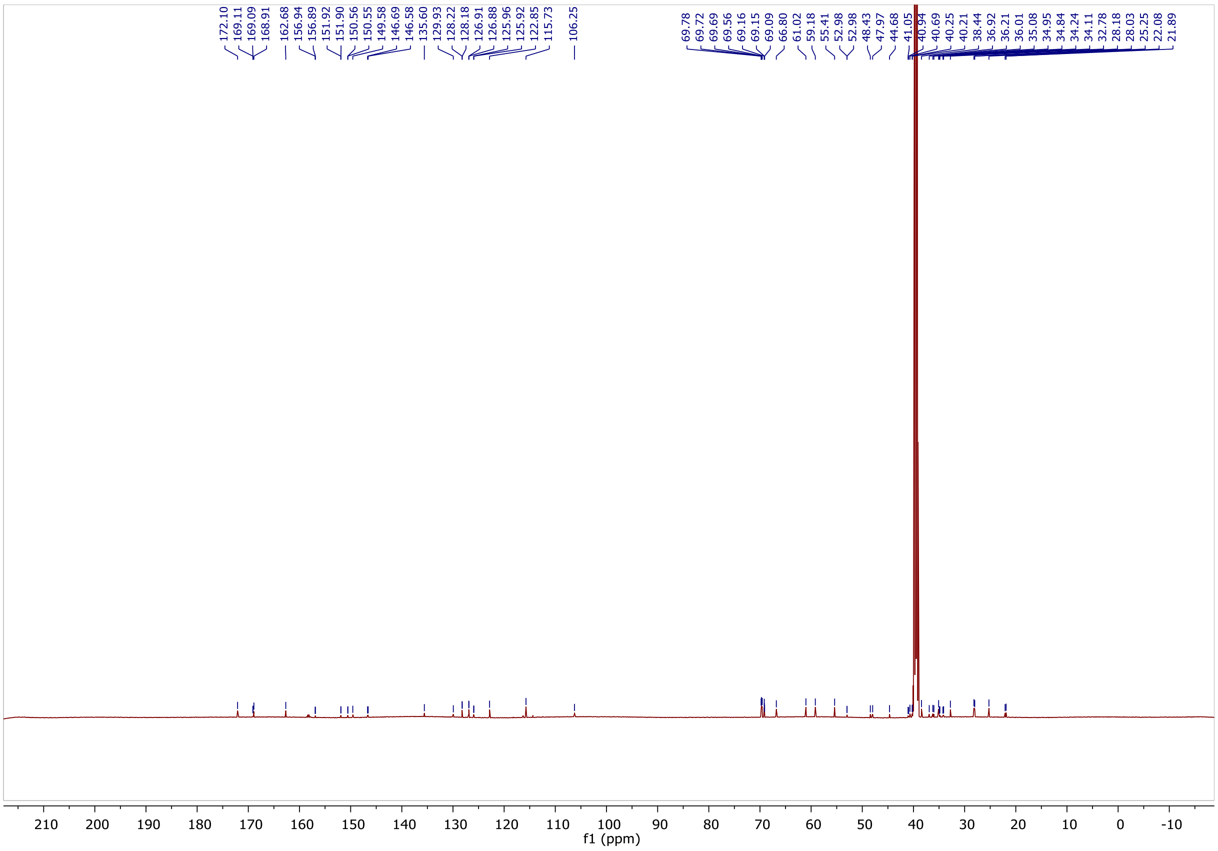

$^{1}H$ NMR spectra of NK206.

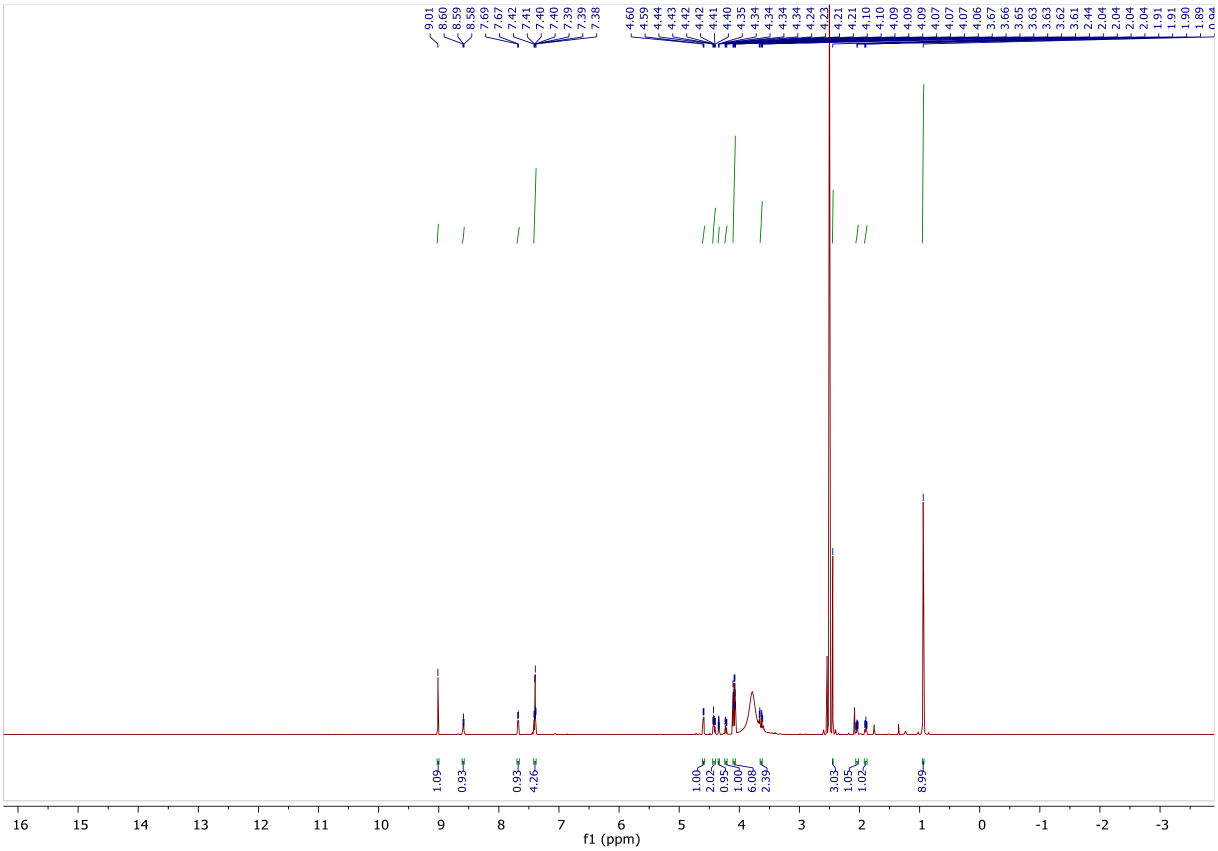

$^{13}C$ NMR spectra of NK206.

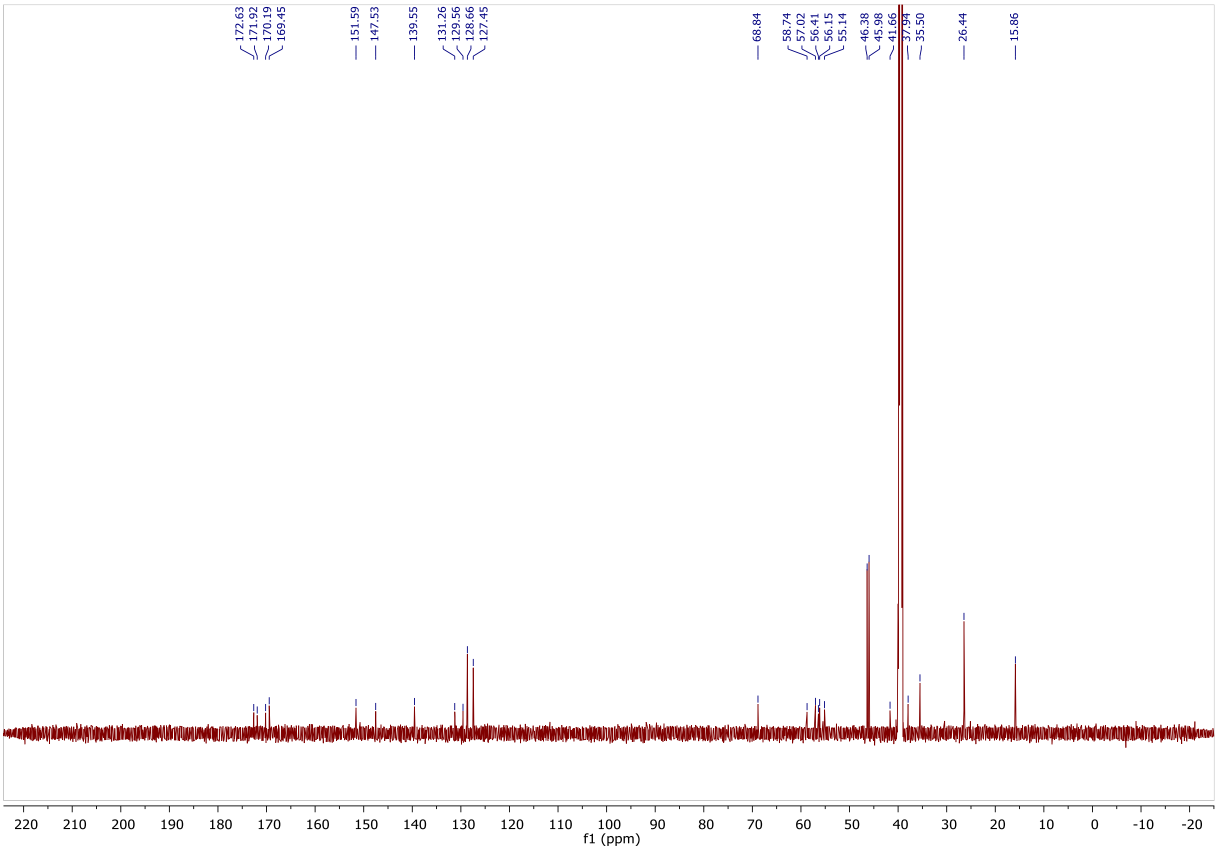

$^1H$ NMR spectra of NK265_2.

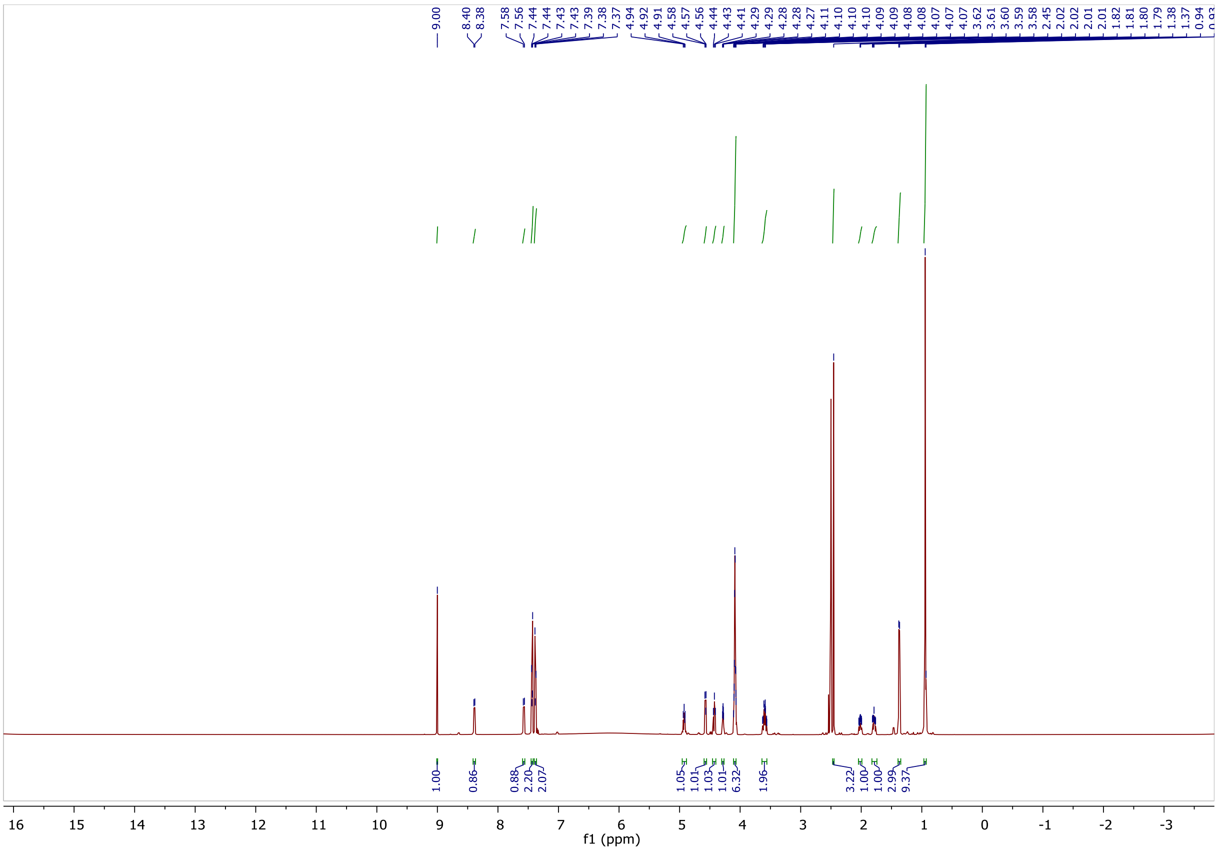

$^{13}C$ NMR spectra of NK265_2.

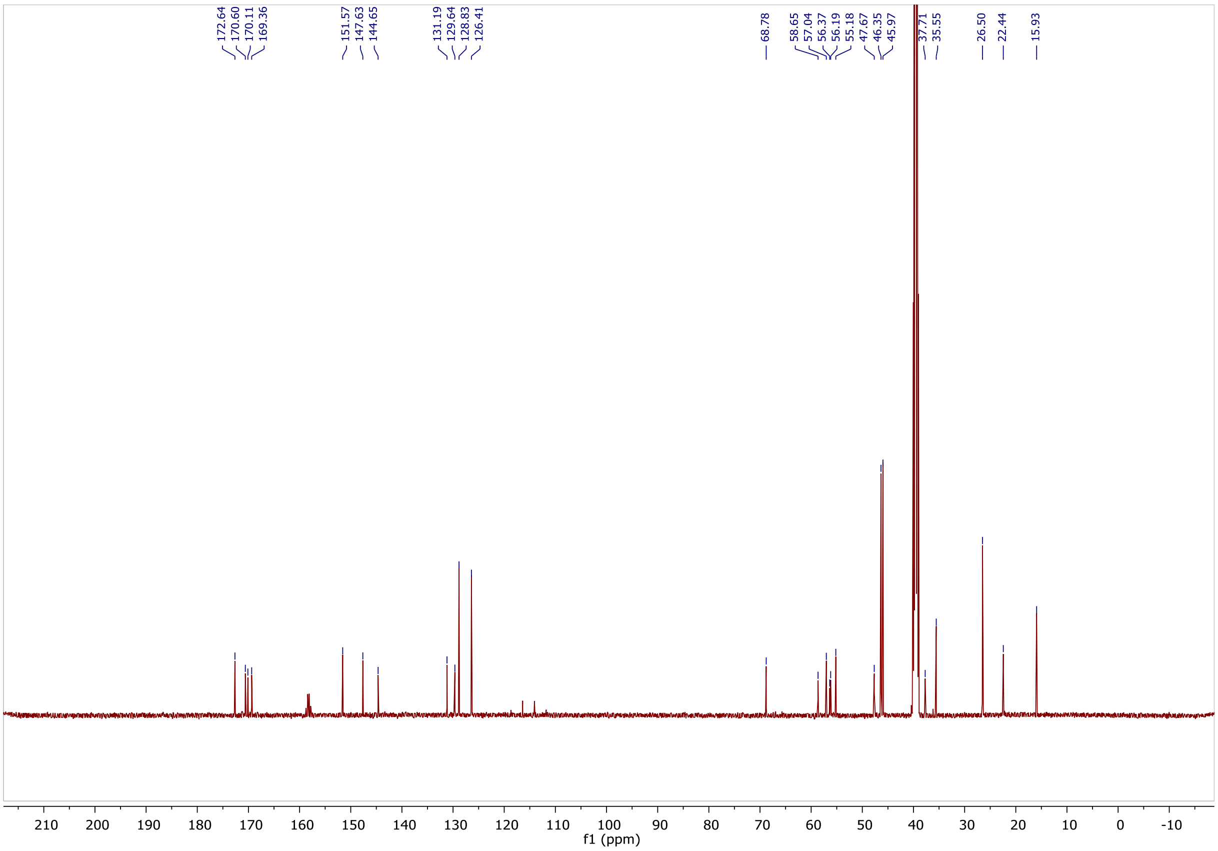

$^{1}H$ NMR spectra of NK213.

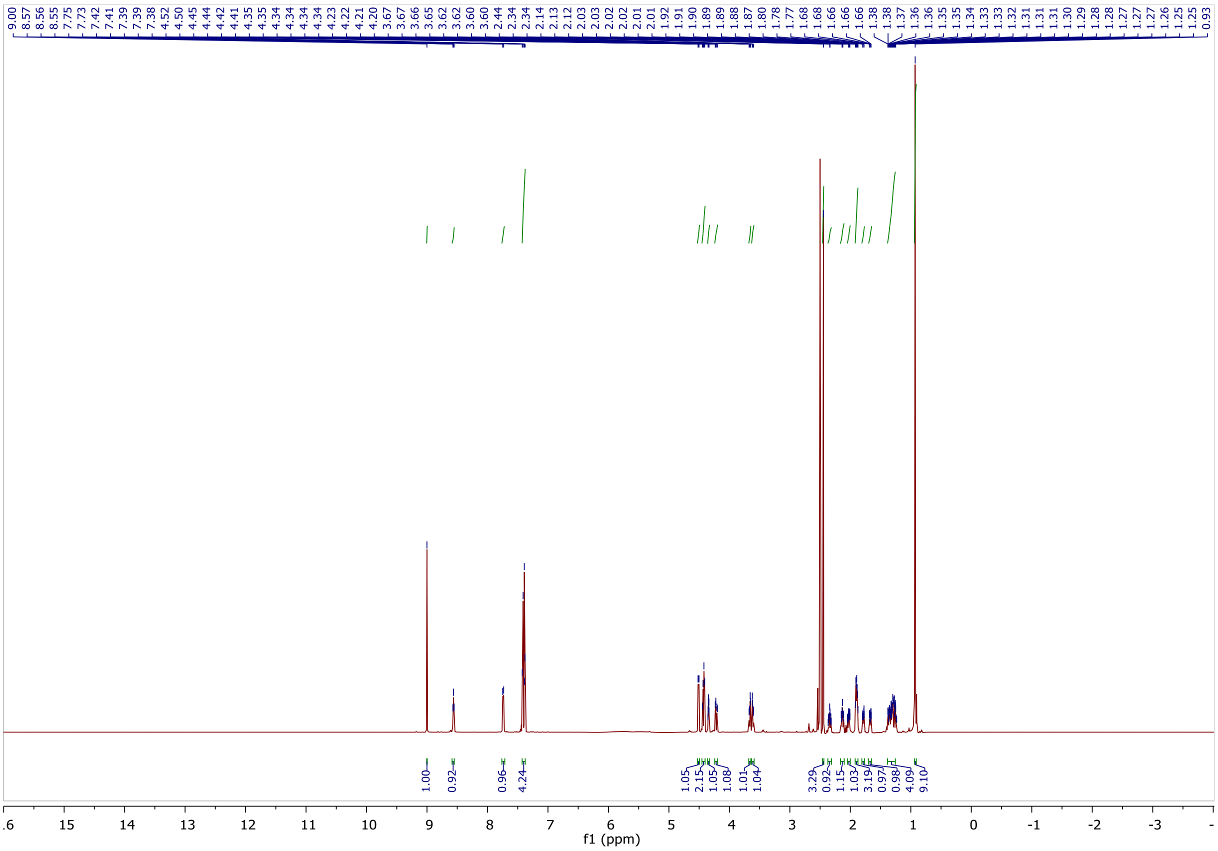

$^{13}C$ NMR spectra of NK213.

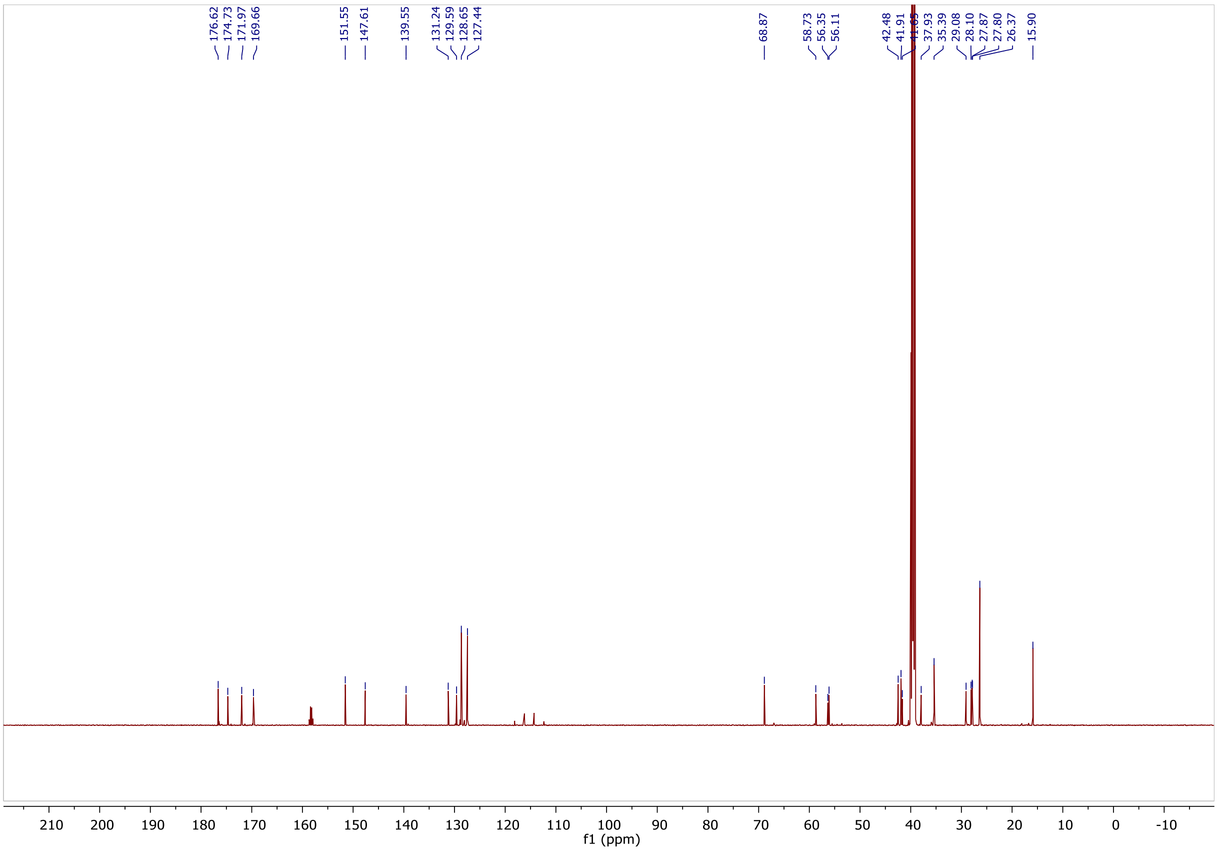

$^{1}H$ NMR spectra of NK244.

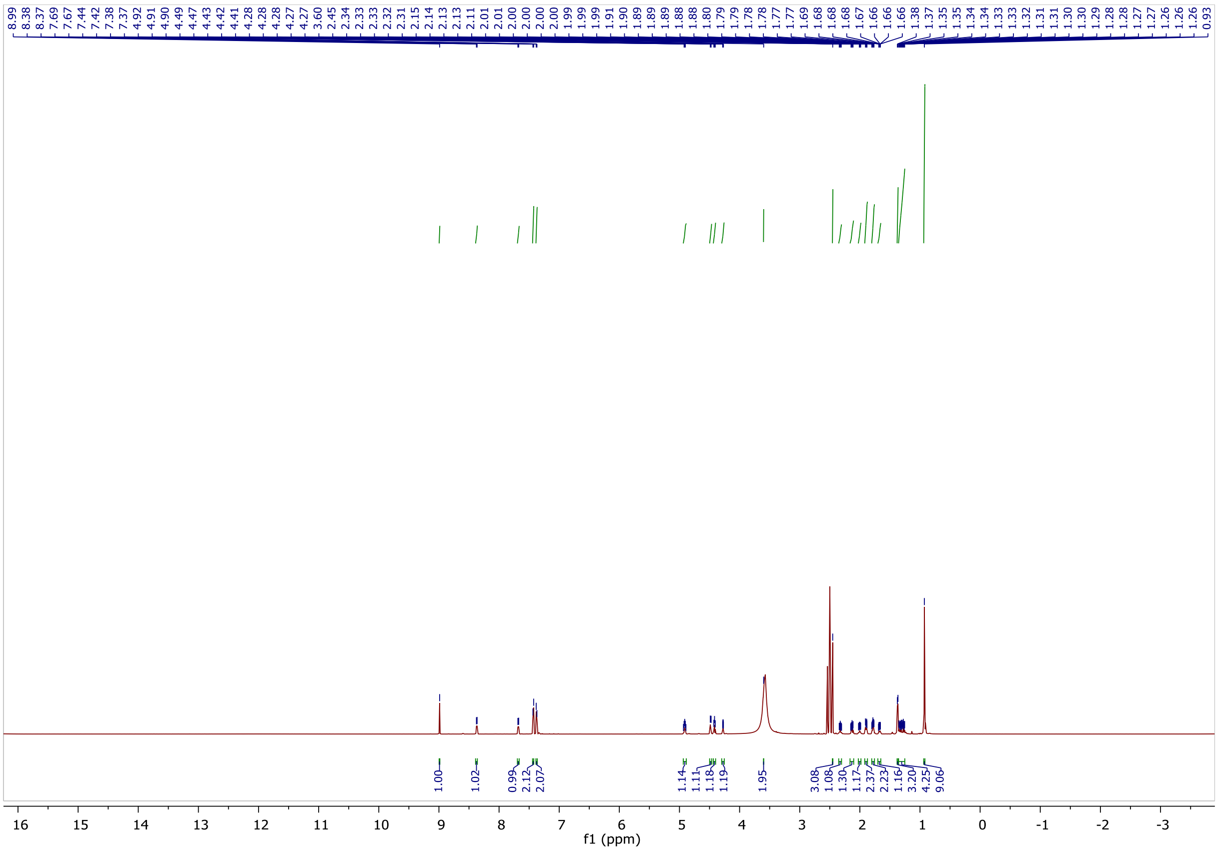

$^{13}C$ NMR spectra of NK244.

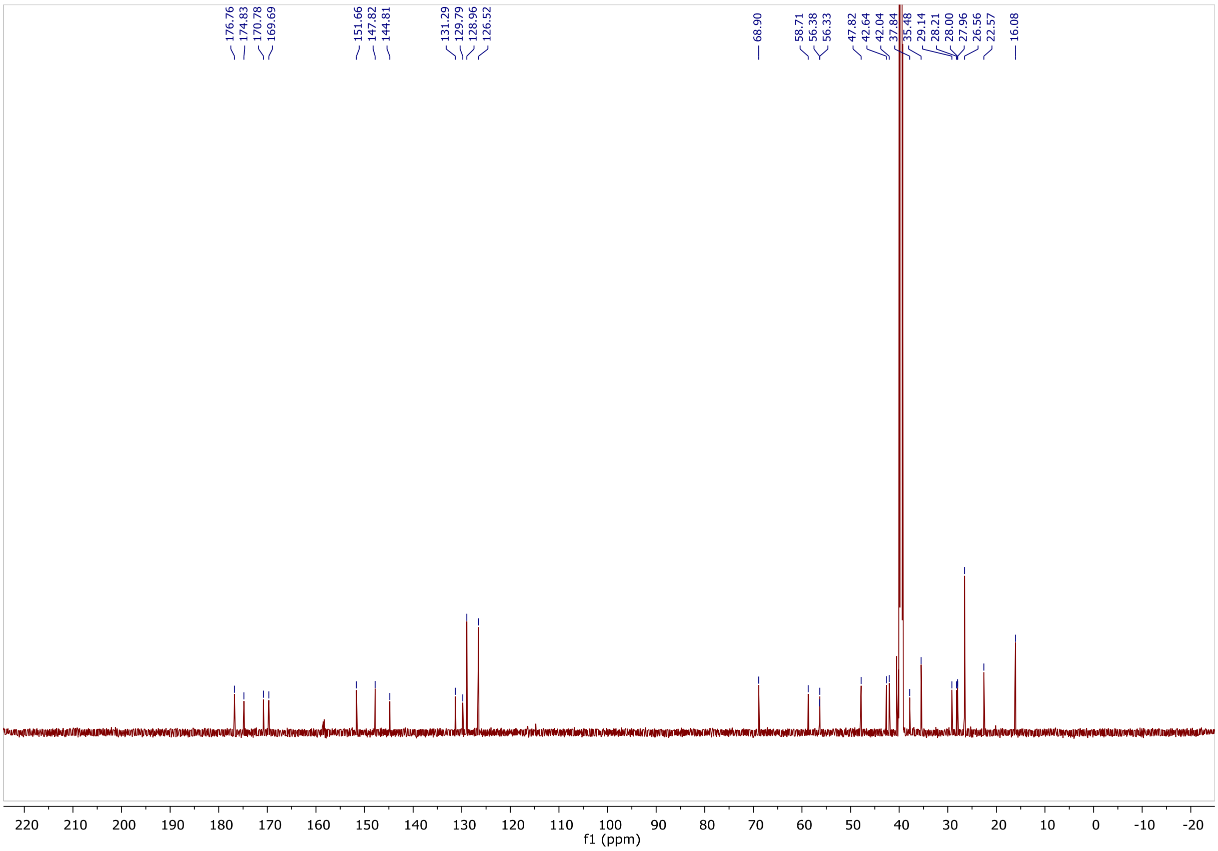

$^{1}H$ NMR spectra of NK209.

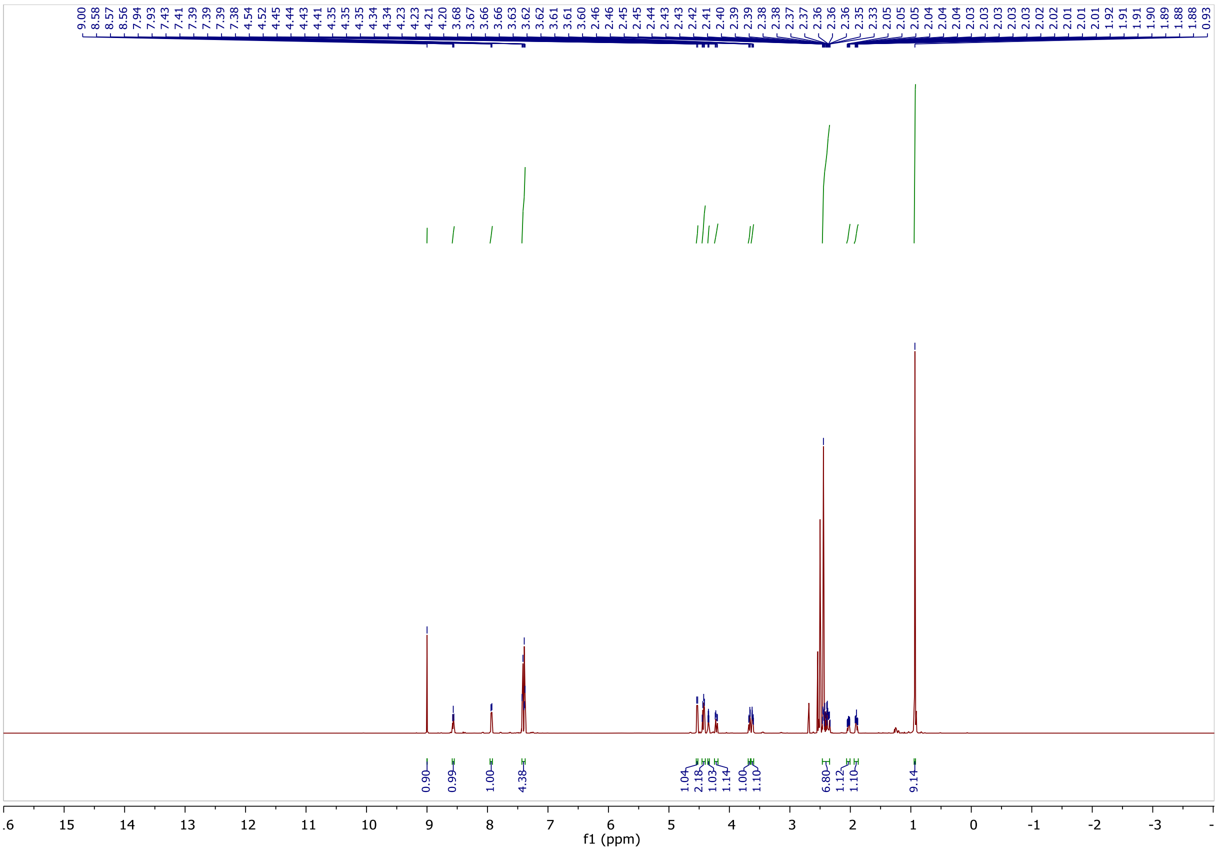

$^{13}C$ NMR spectra of NK209.

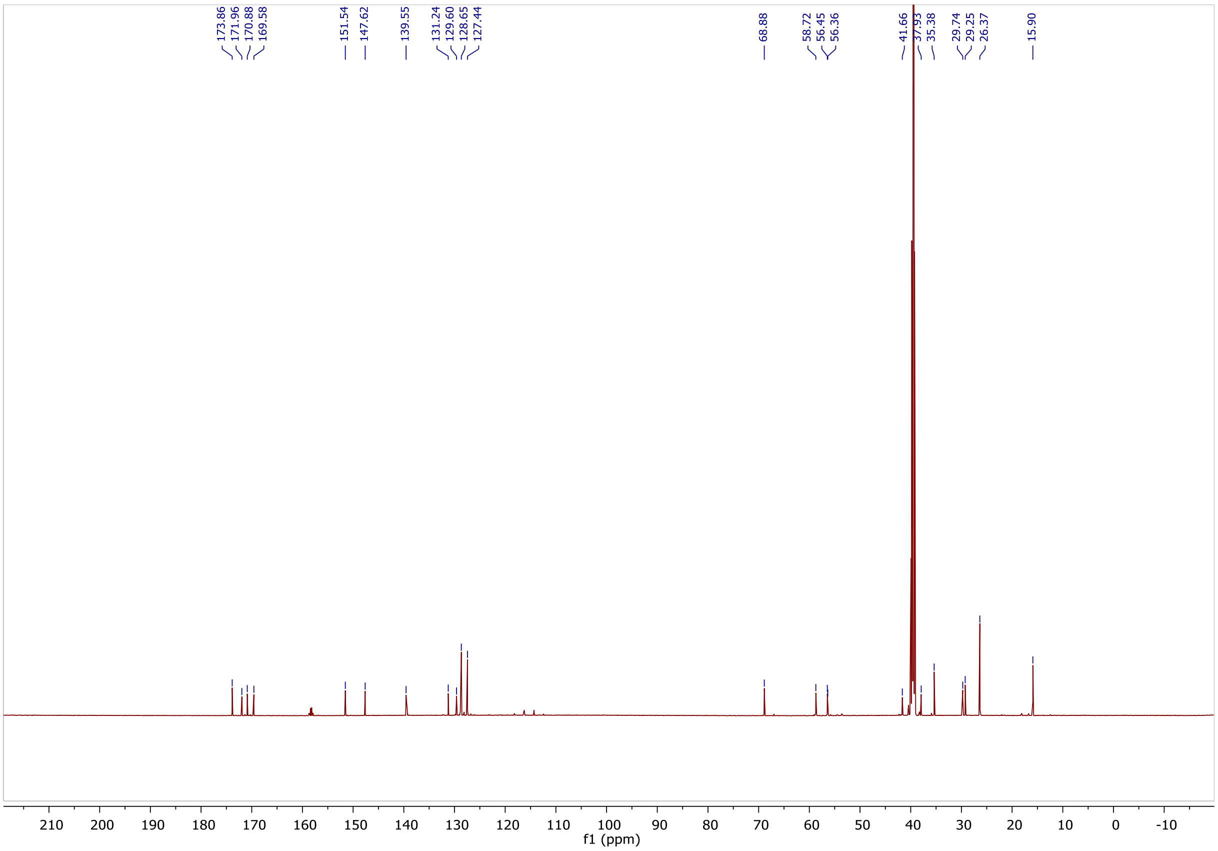

$^{1}H$ NMR spectra of NK208.

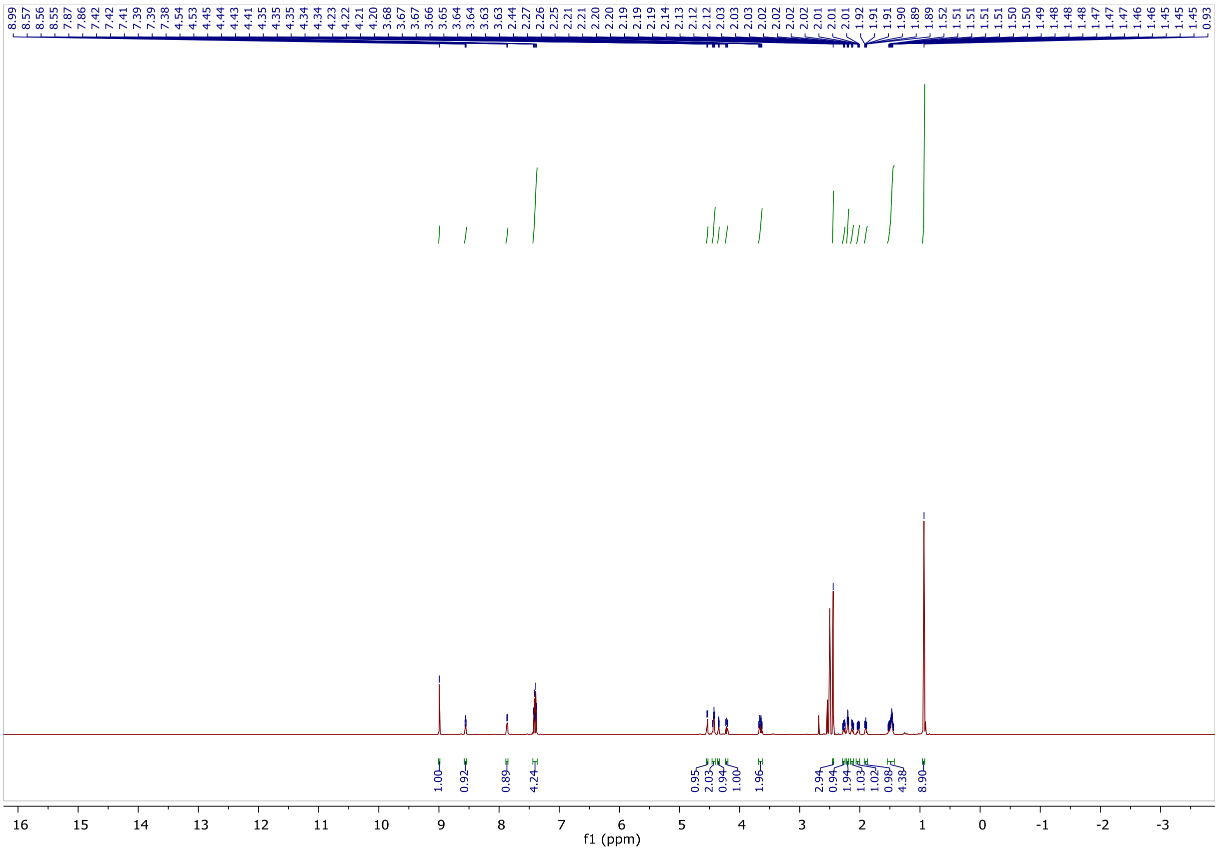

$^{13}C$ NMR spectra of NK208.

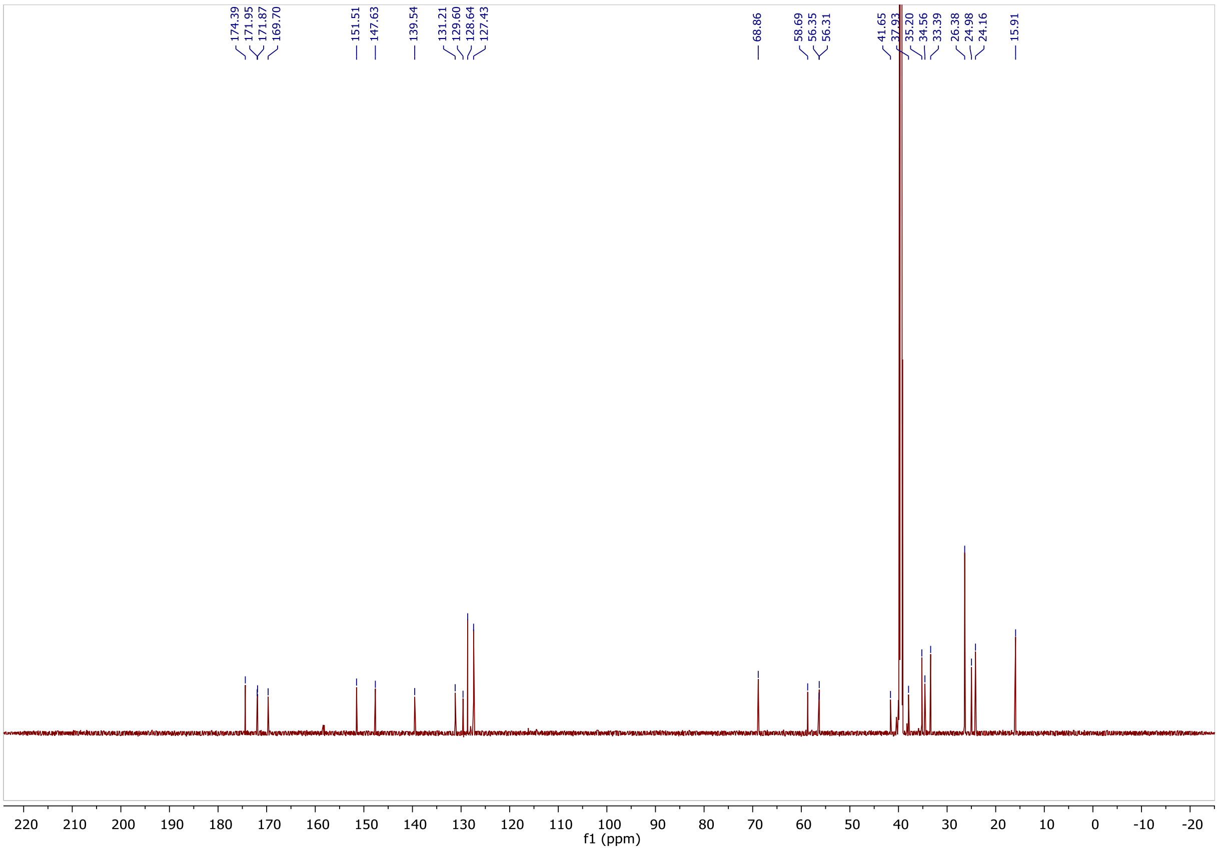

<sup></sup>

$^{1}$H NMR spectra of NK215.

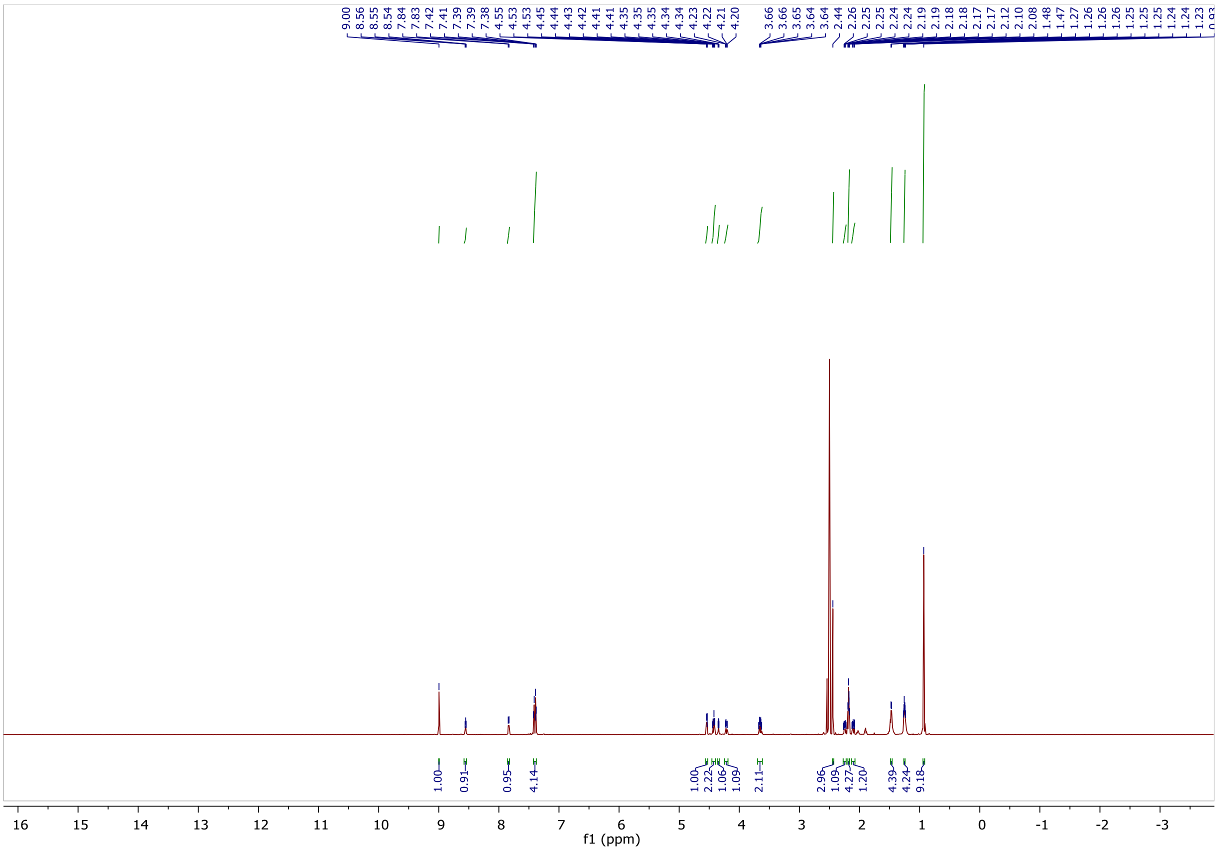

$^{13}$C NMR spectra of NK215.

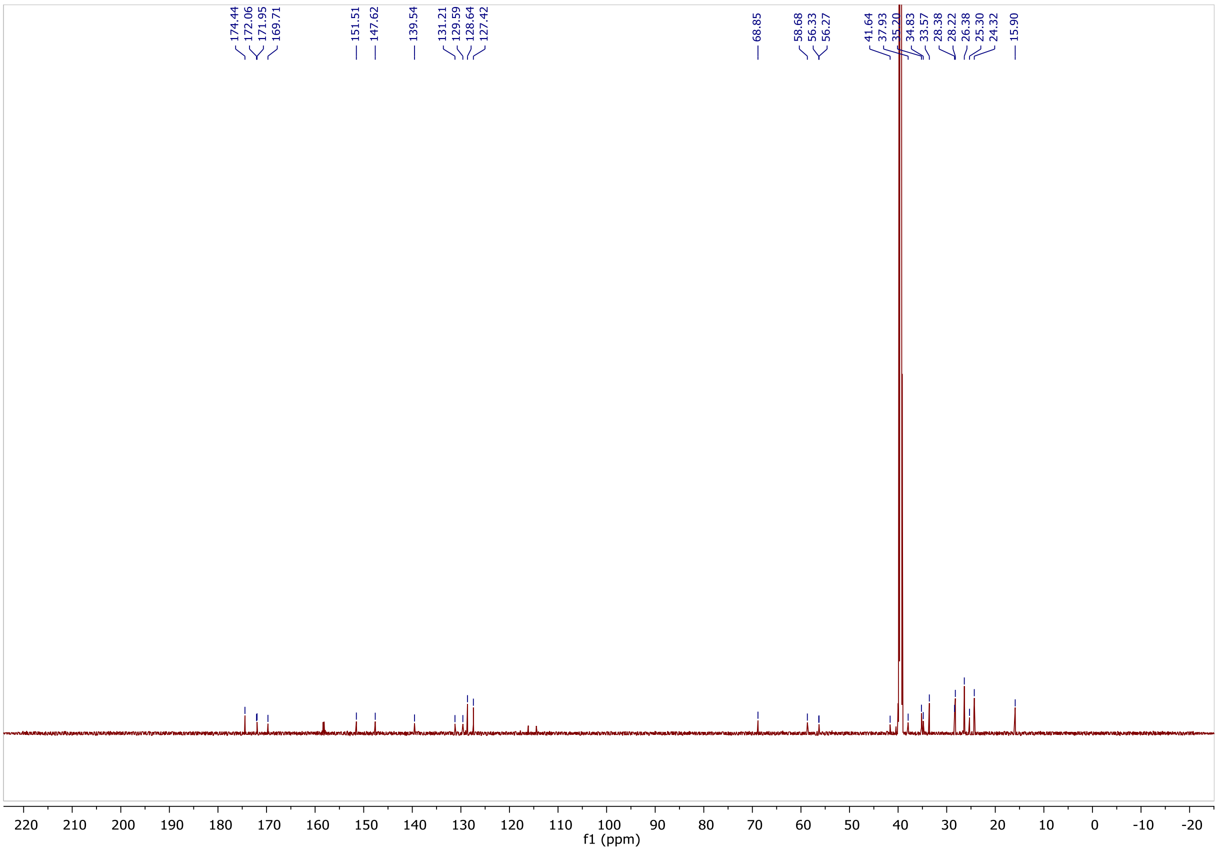

$^1$H NMR spectra of NK207.

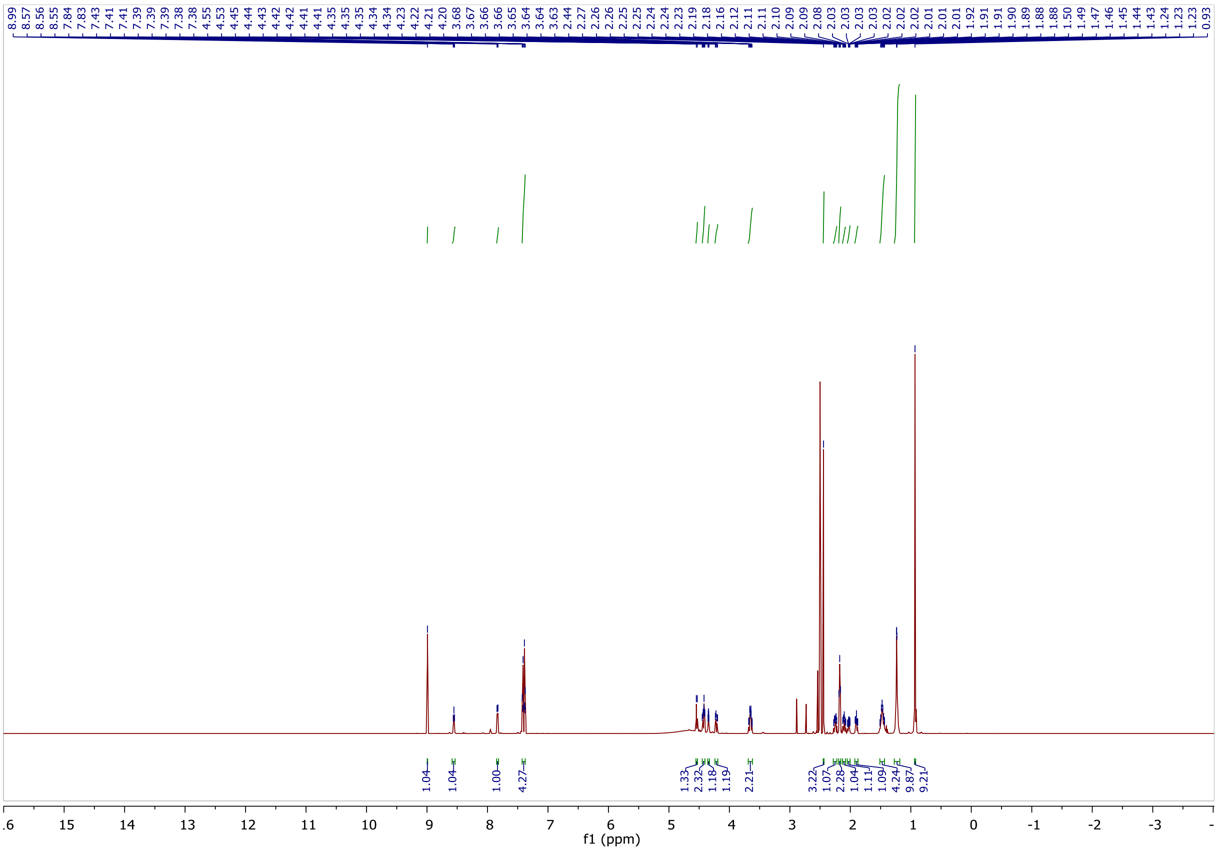

$^{13}$C NMR spectra of NK207.

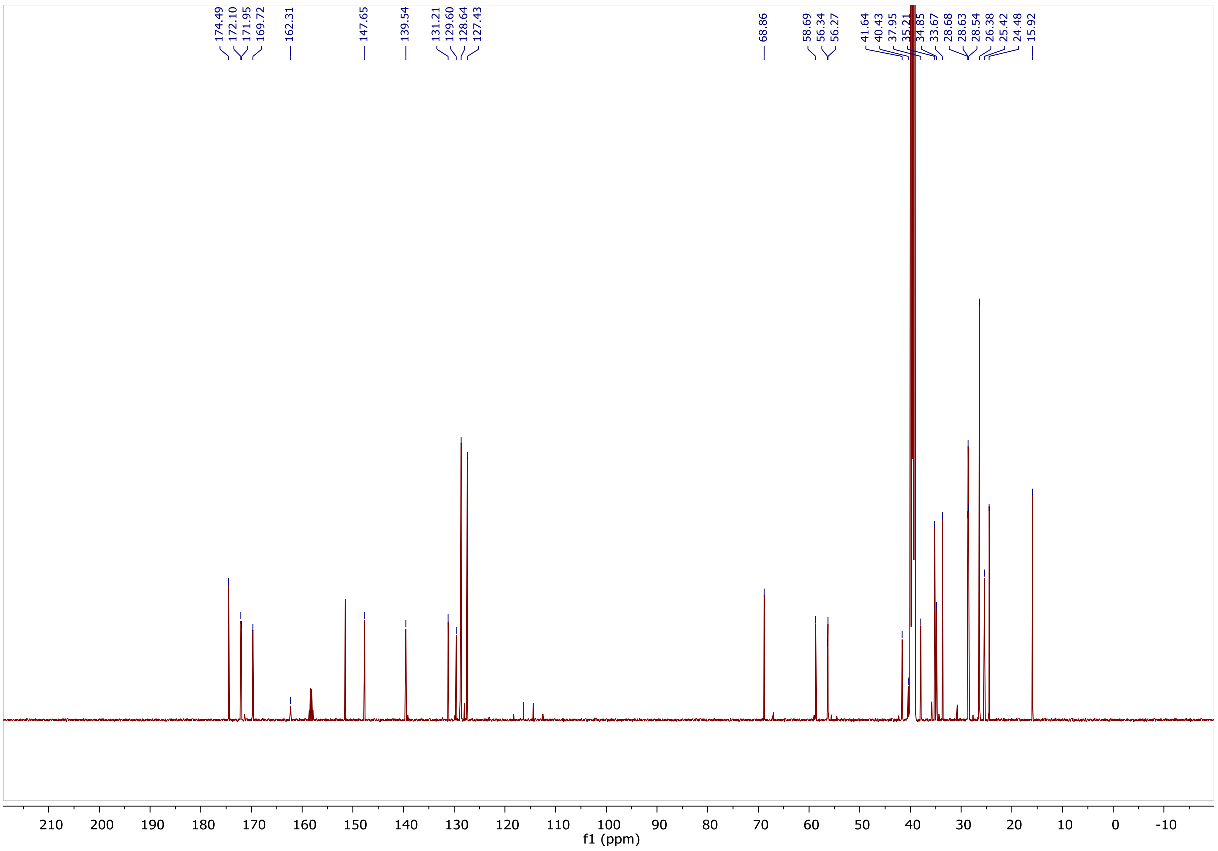

[1]H NMR spectra of NK216.

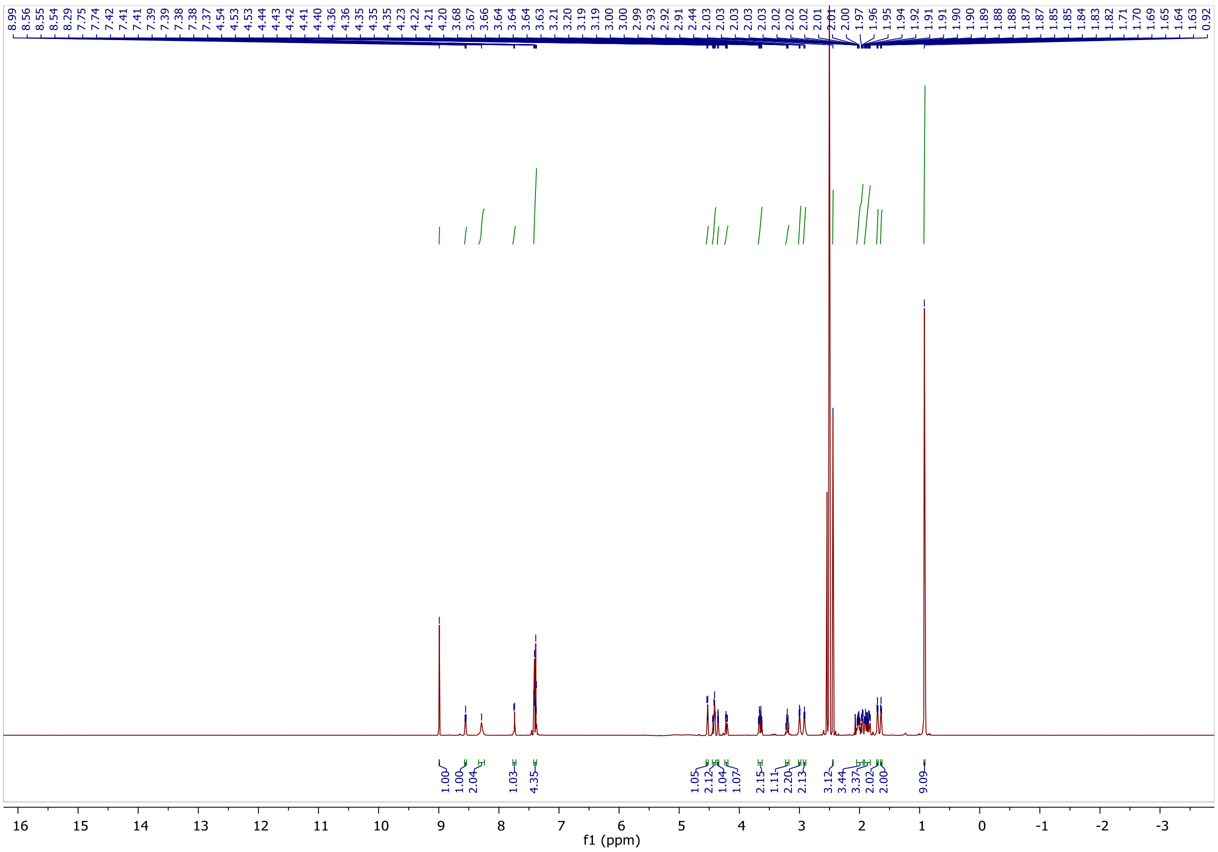

[13]C NMR spectra of NK216.

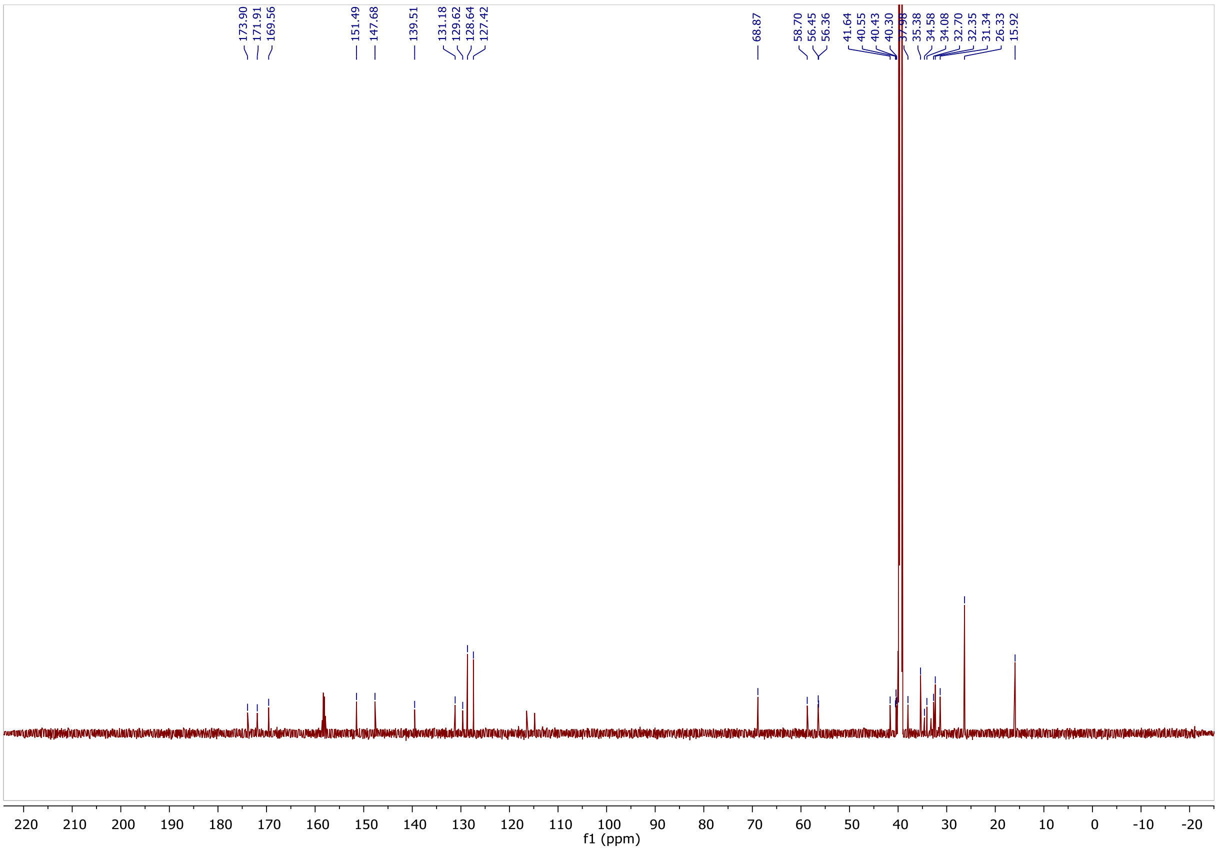

$^{1}H$ NMR spectra of NK239.

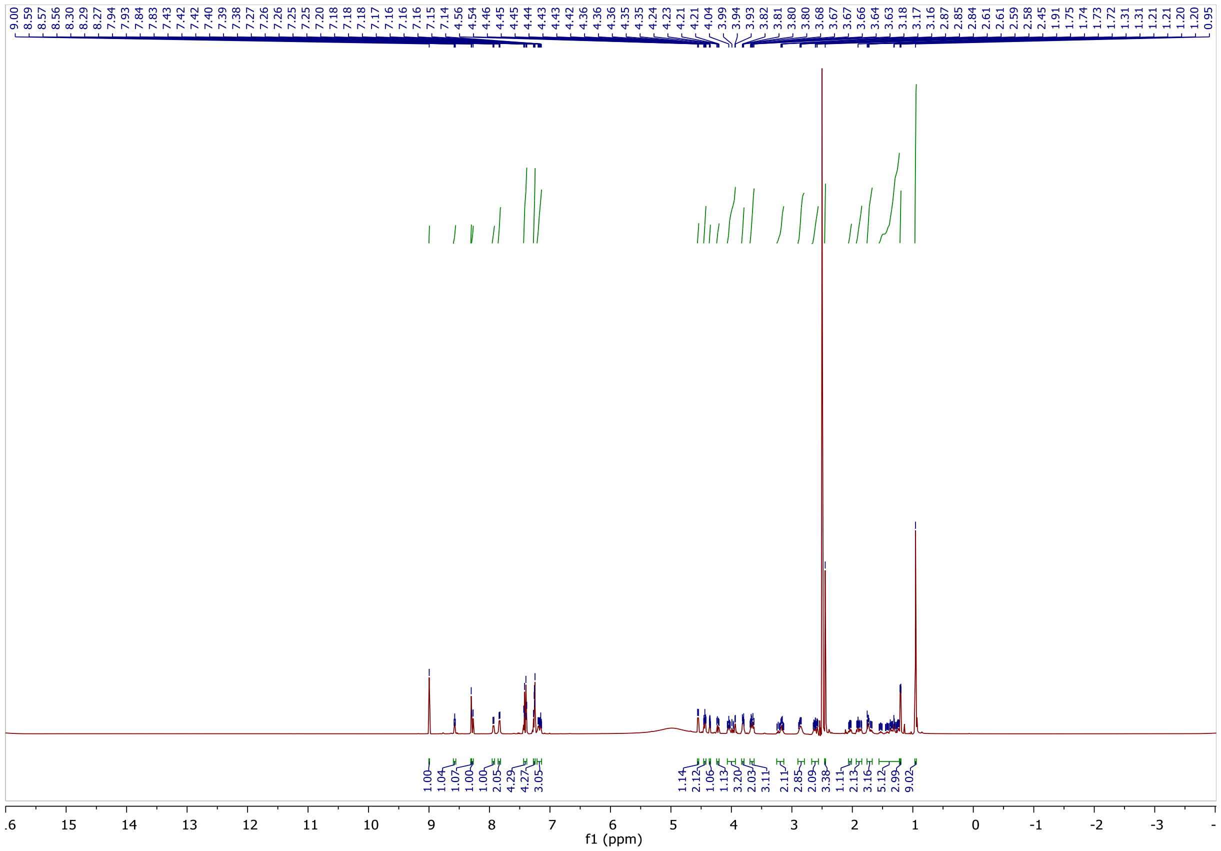

$^{13}C$ NMR spectra of NK239.

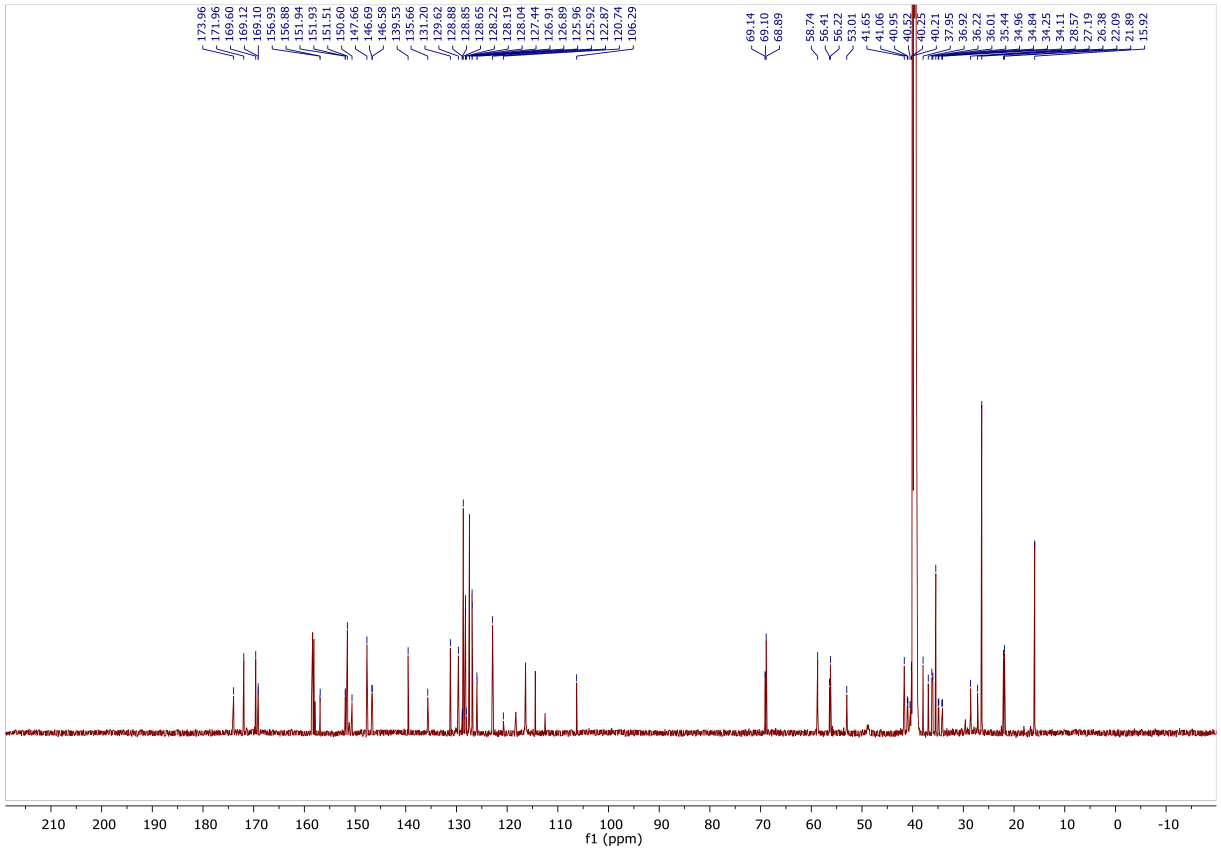

$^{1}H$ NMR spectra of NK240.

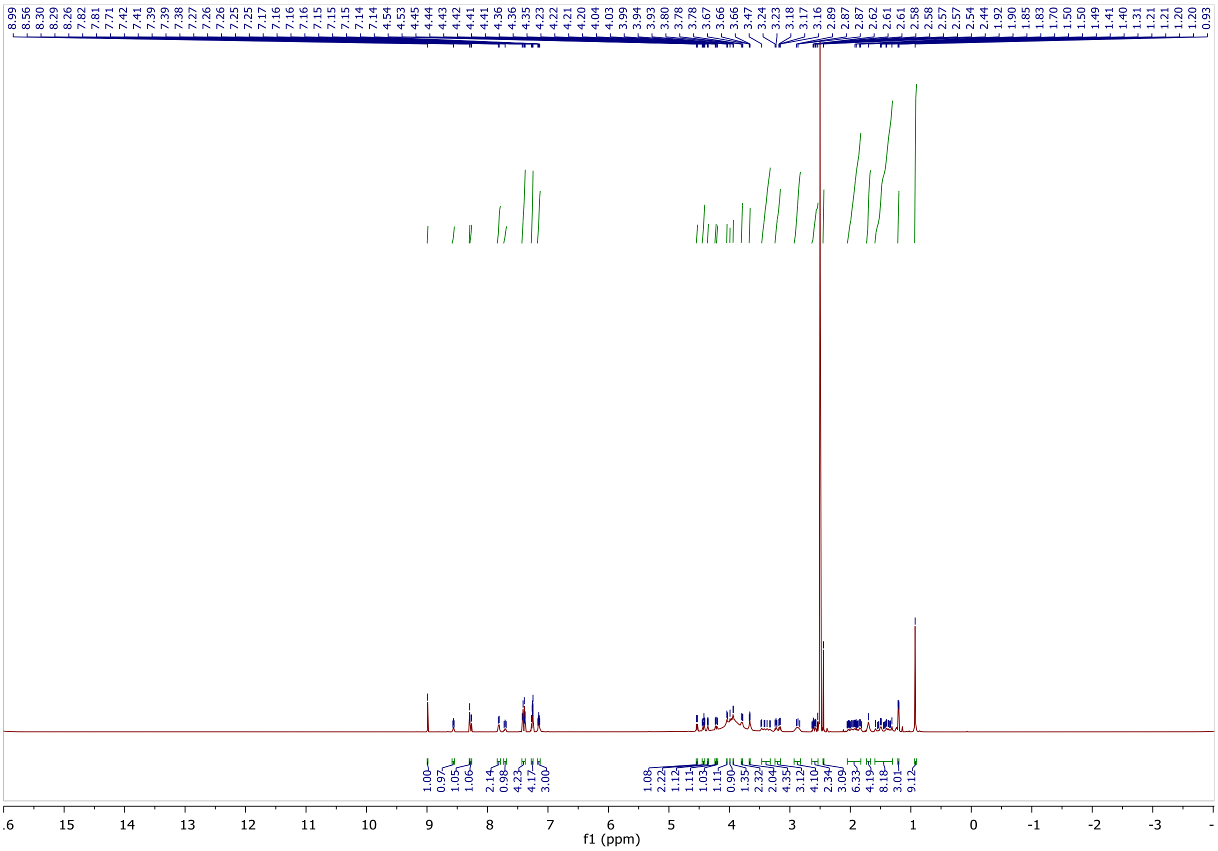

$^{13}C$ NMR spectra of NK240.

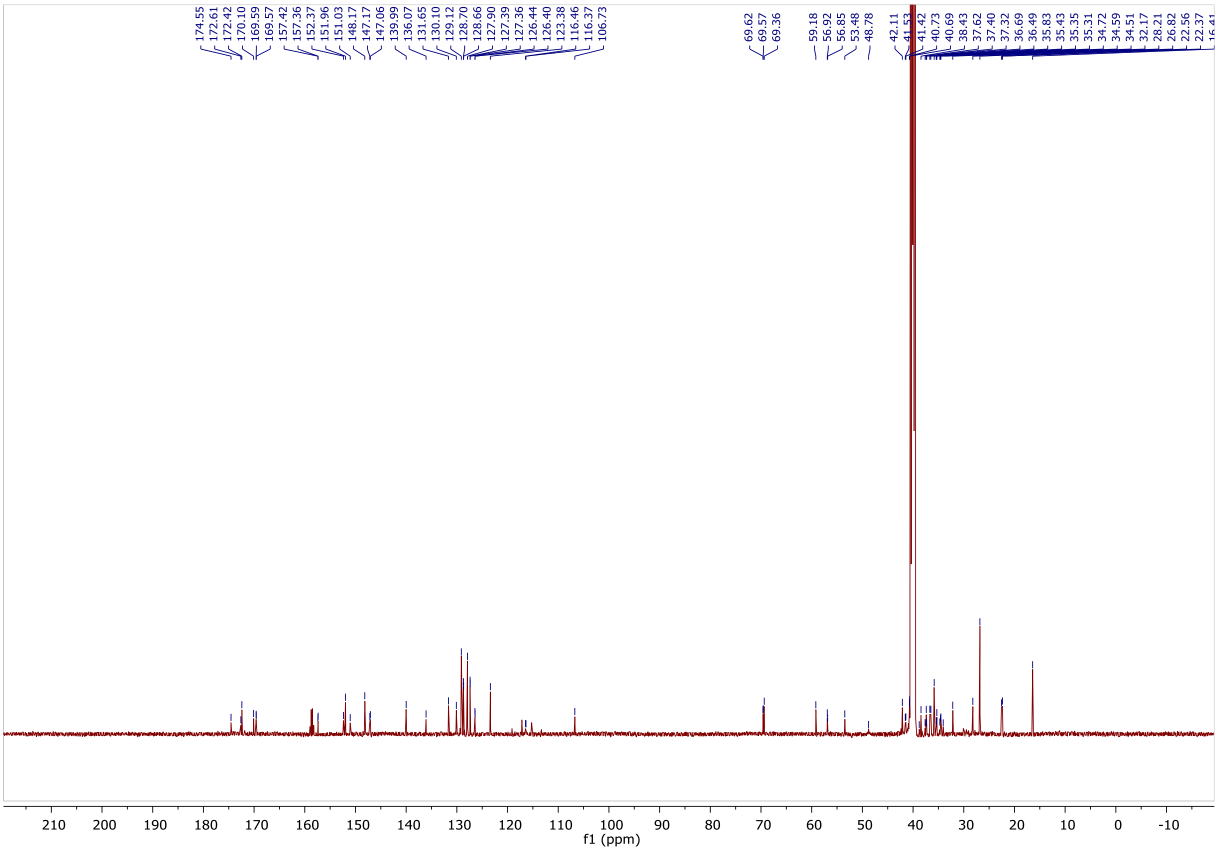

$^{1}H$ NMR spectra of NK237.

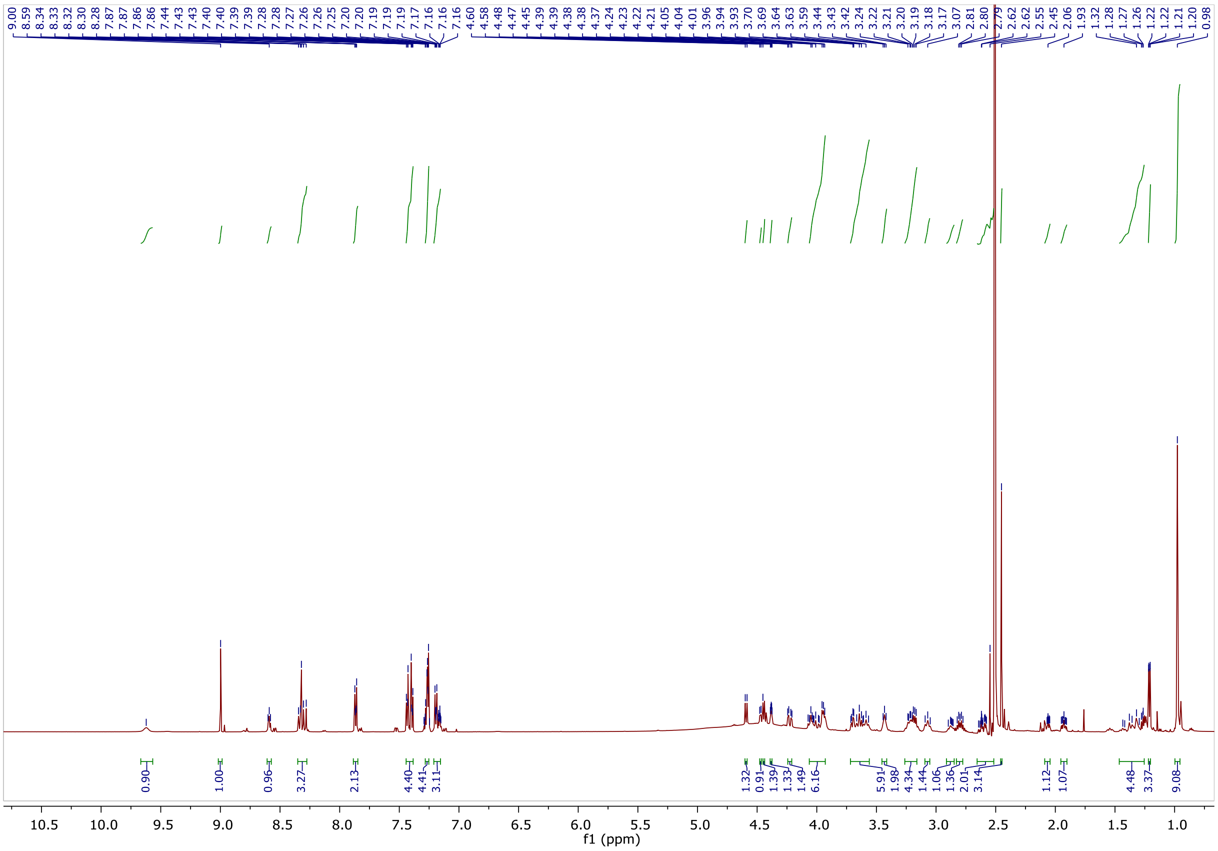

$^{13}C$ NMR spectra of NK237.

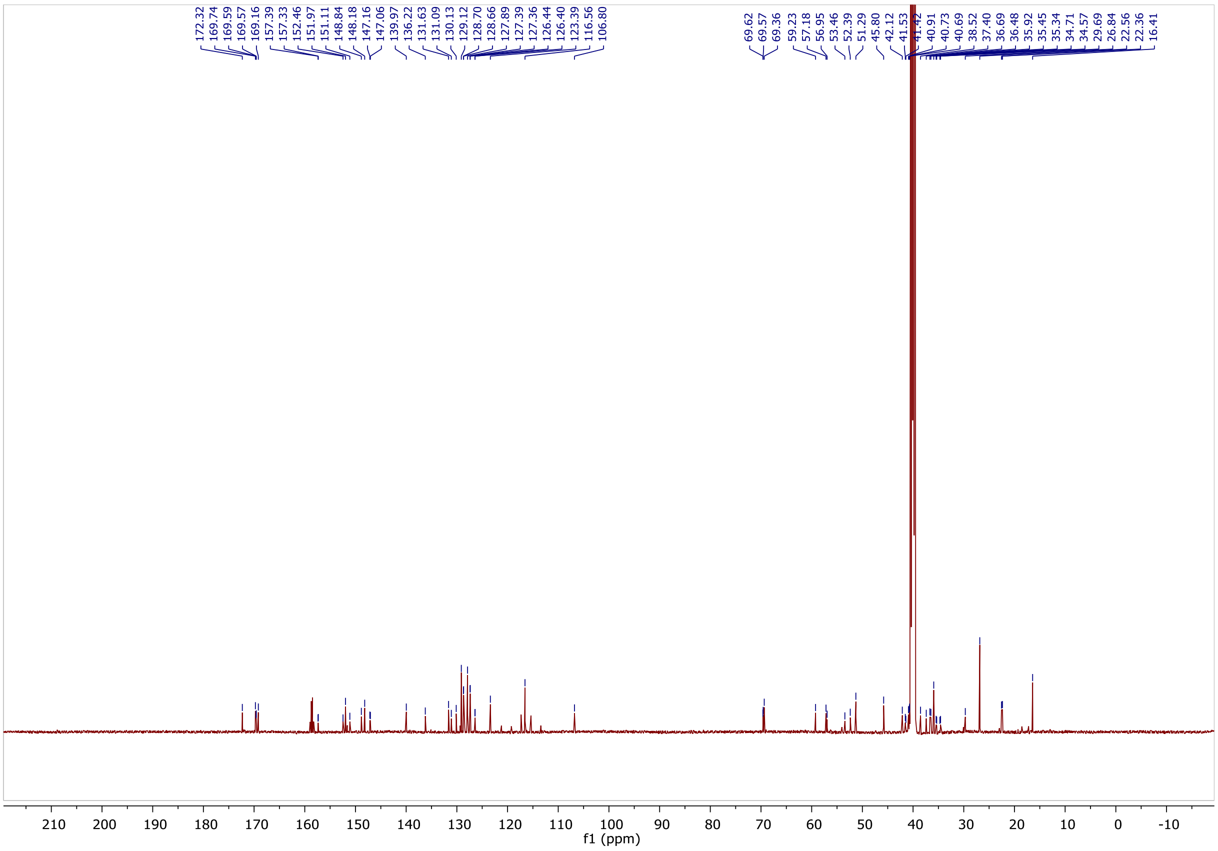

$^{1}H$ NMR spectra of NK238.

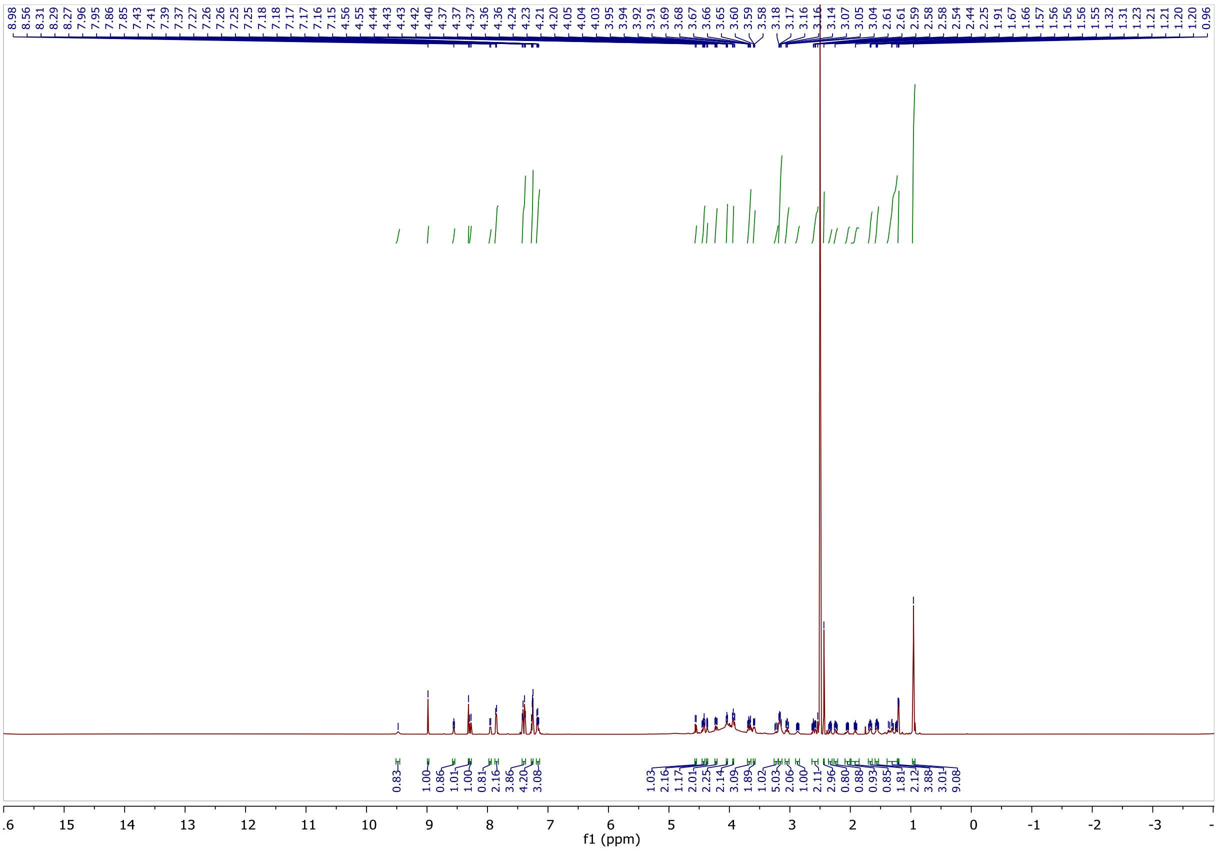

$^{13}C$ NMR spectra of NK238.

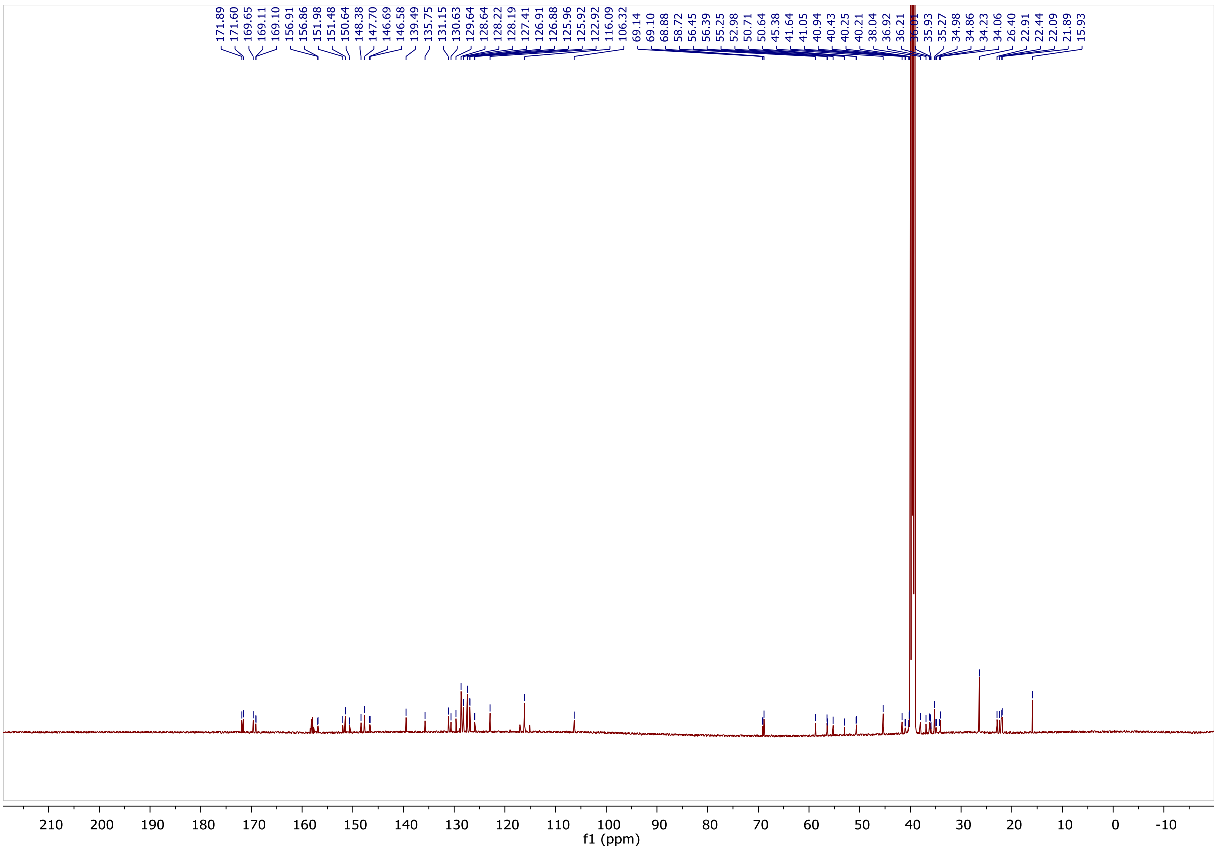

[1]H NMR spectra of NK242.

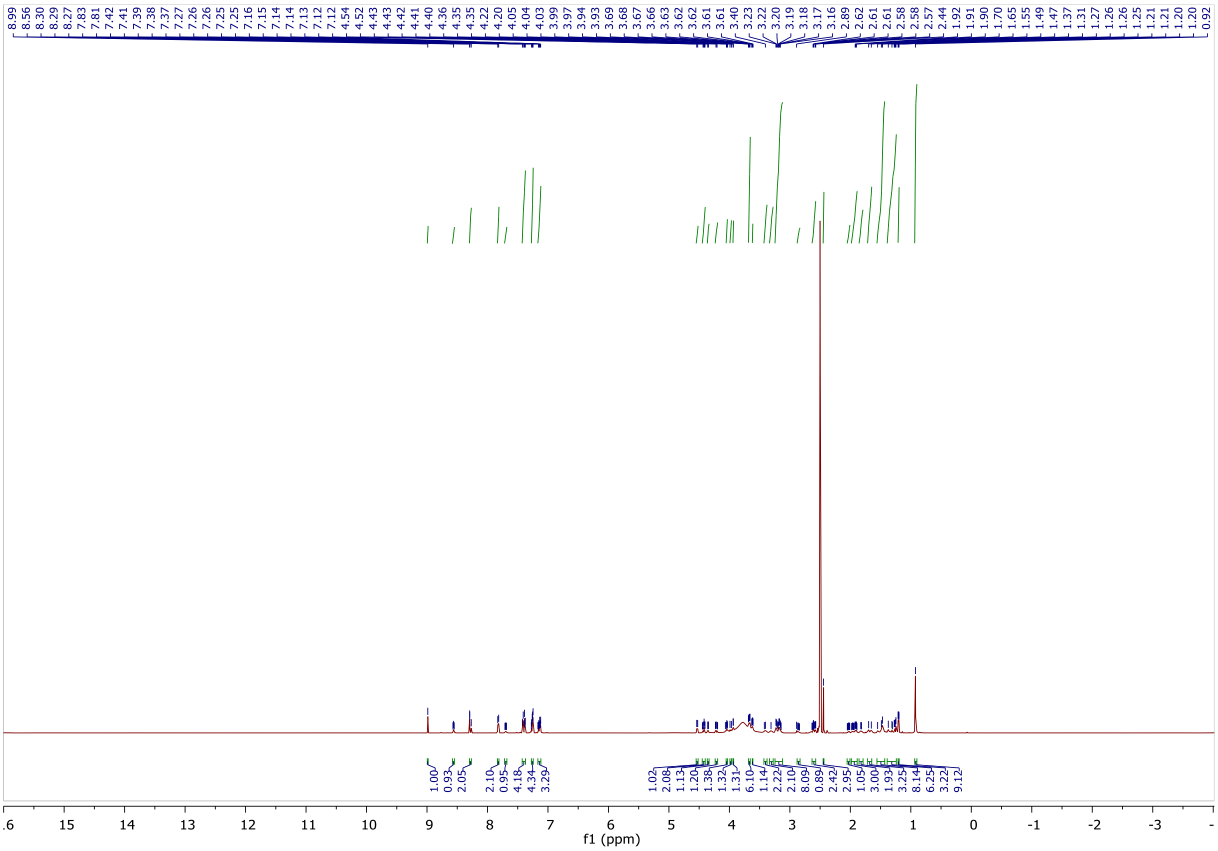

[13]C NMR spectra of NK242.

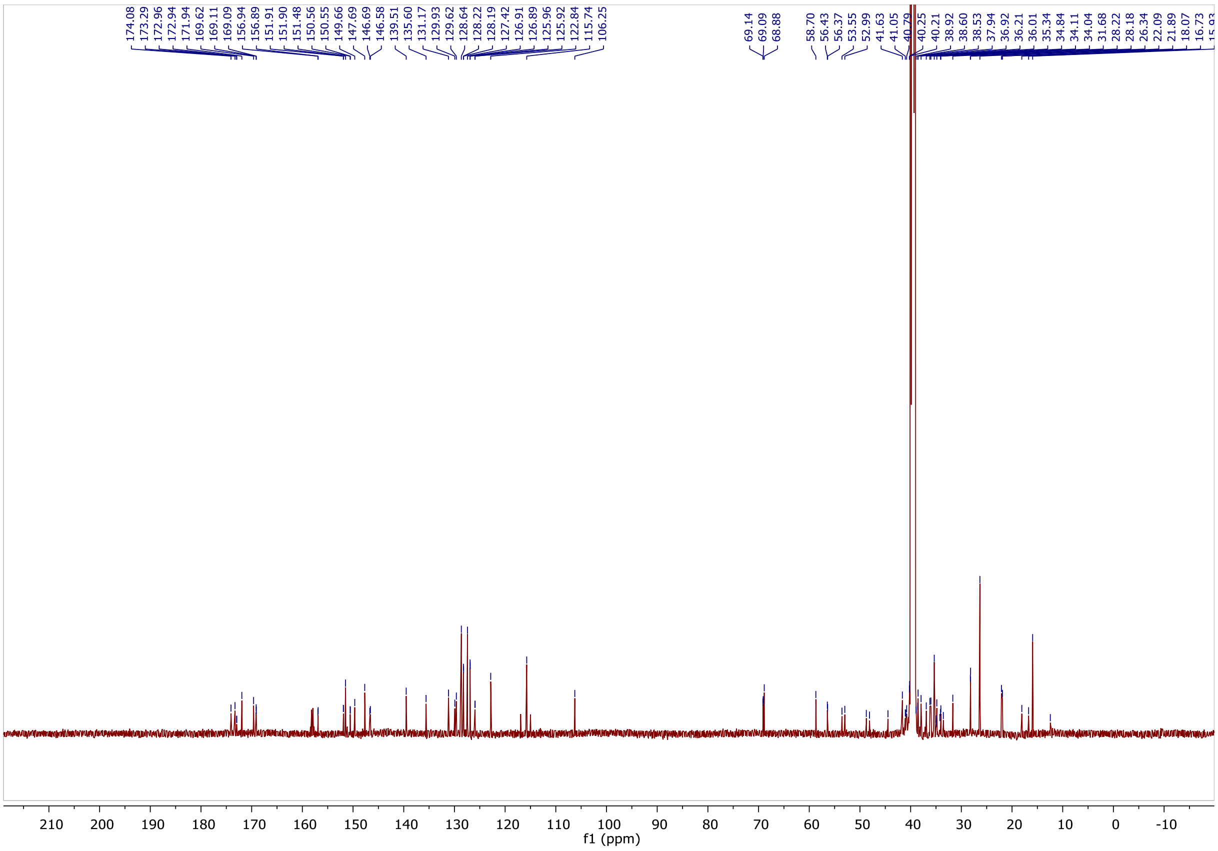

$^{1}H$ NMR spectra of NK245.

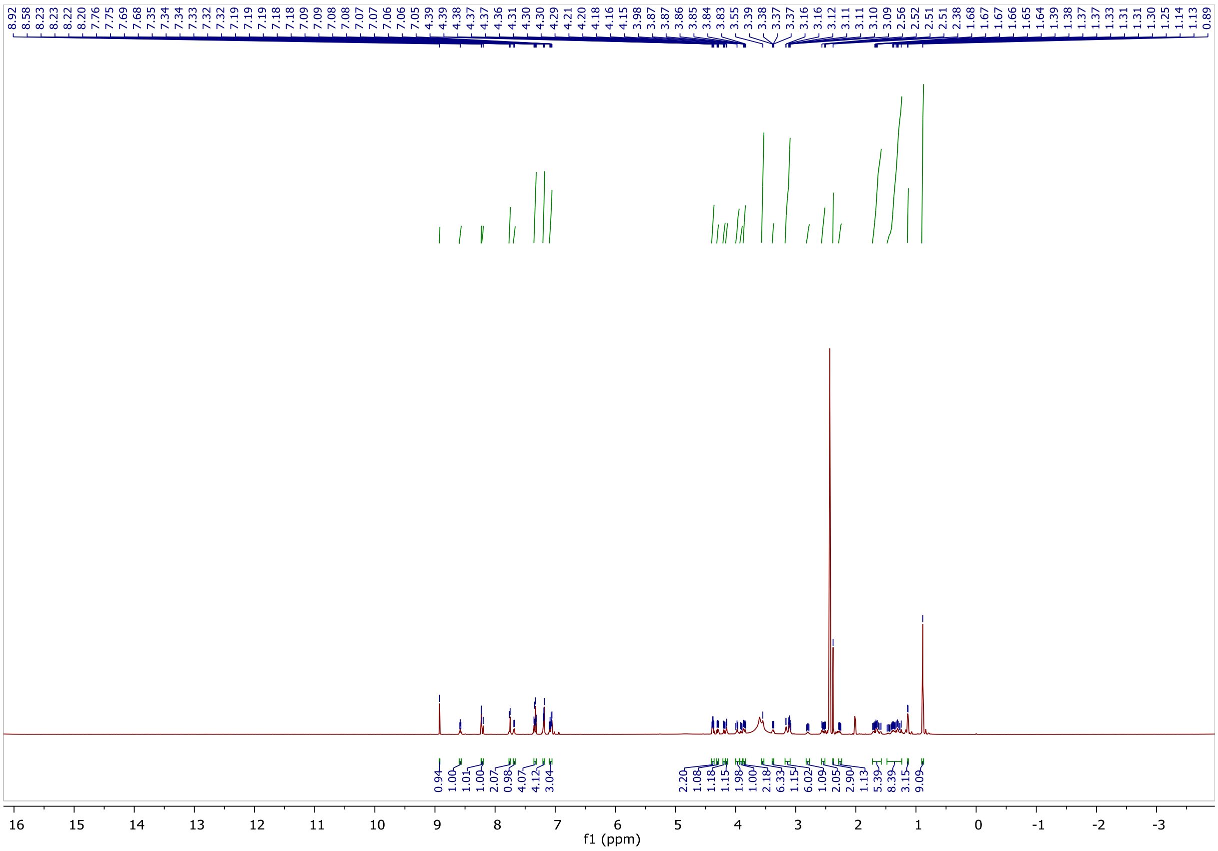

$^{13}C$ NMR spectra of NK245.

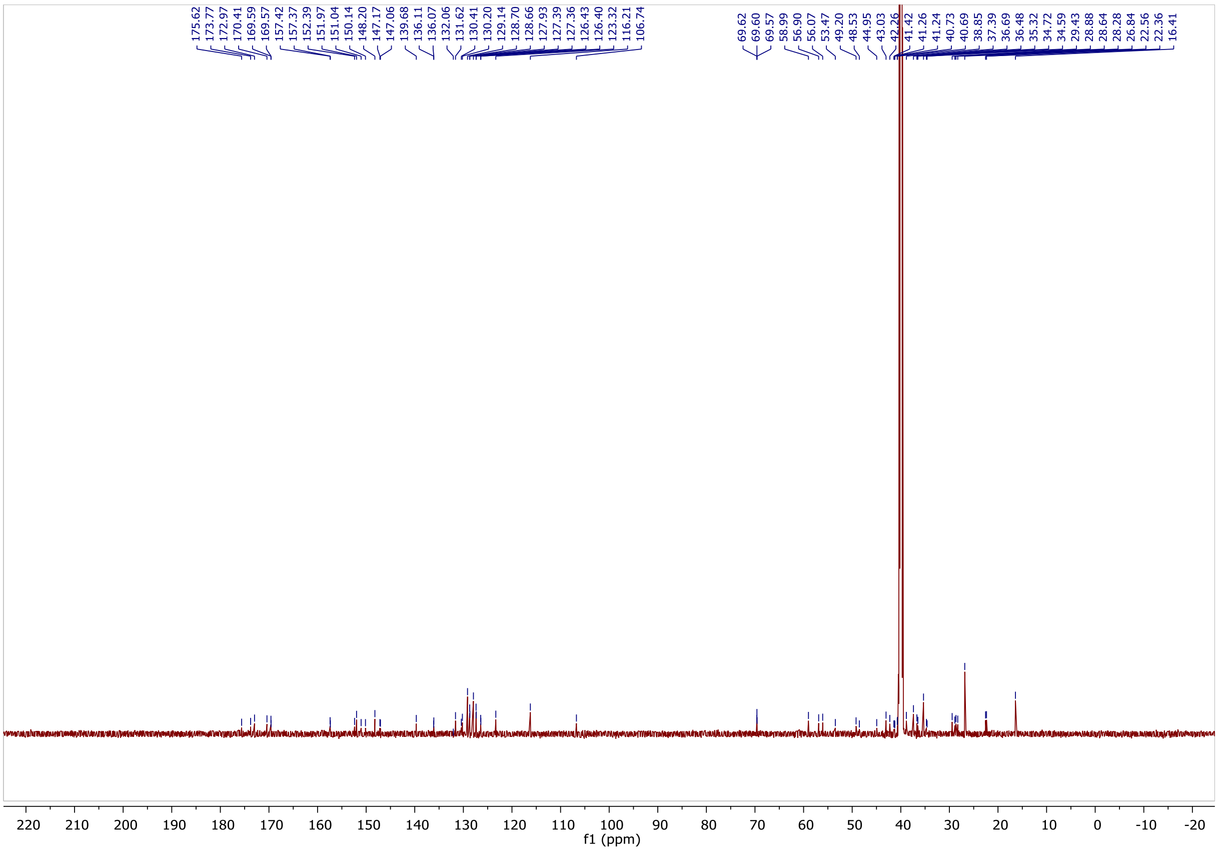

$^{1}H$ NMR spectra of NK221.

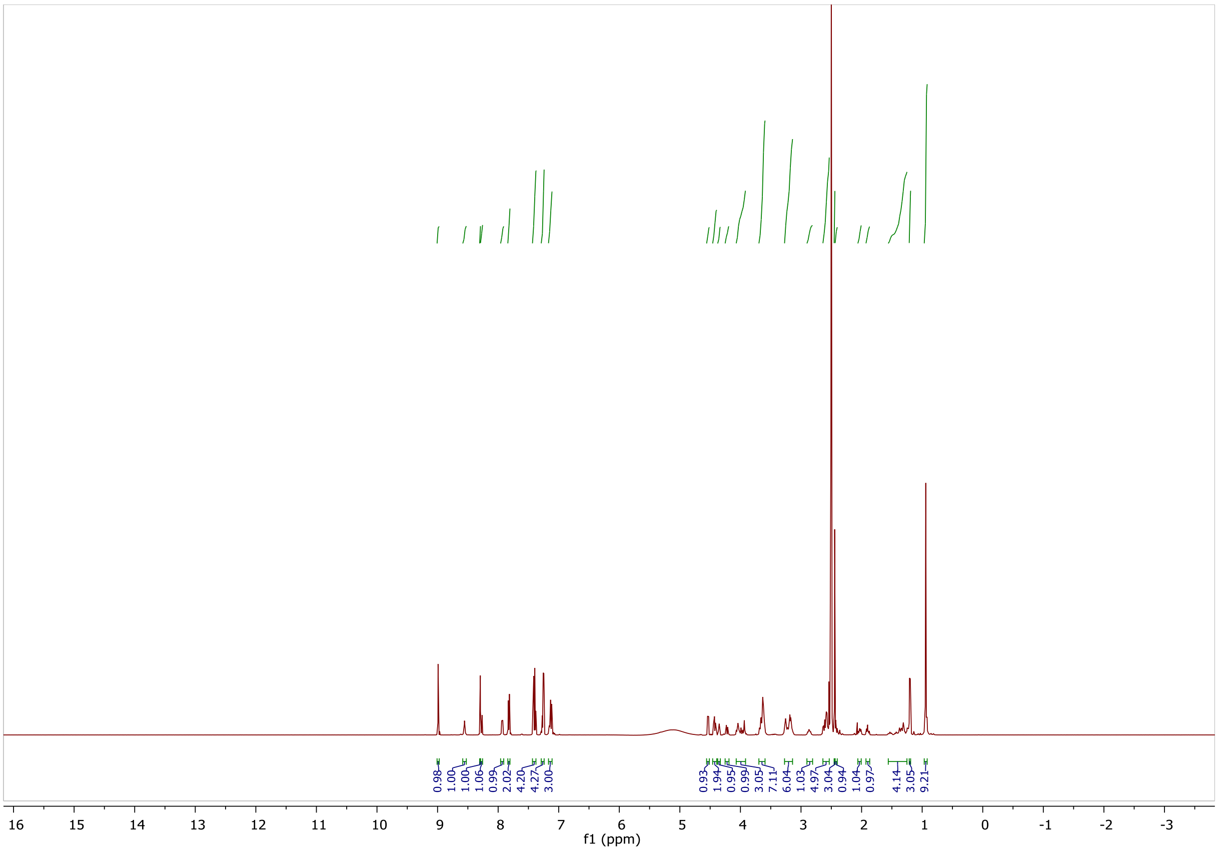

$^{13}C$ NMR spectra of NK221.

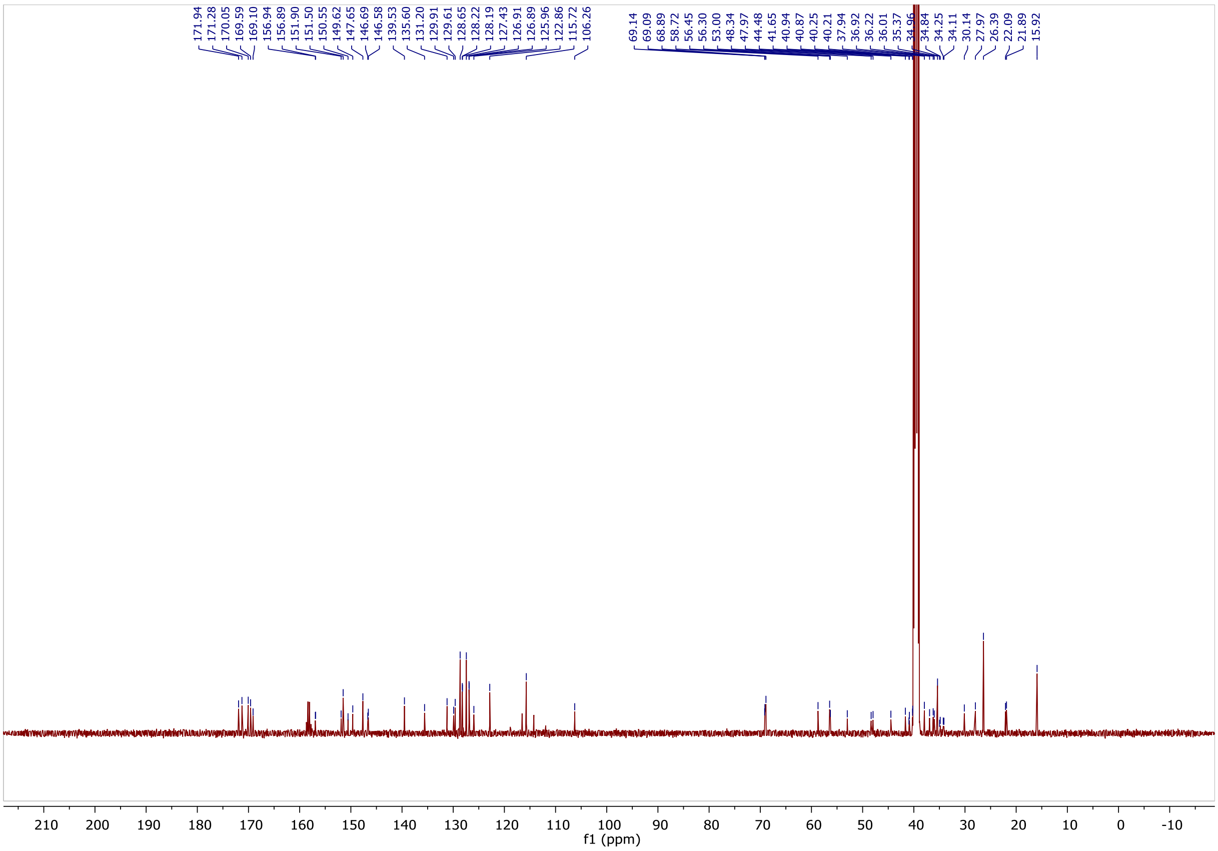

#### $^{1}$H NMR spectra of NK222.

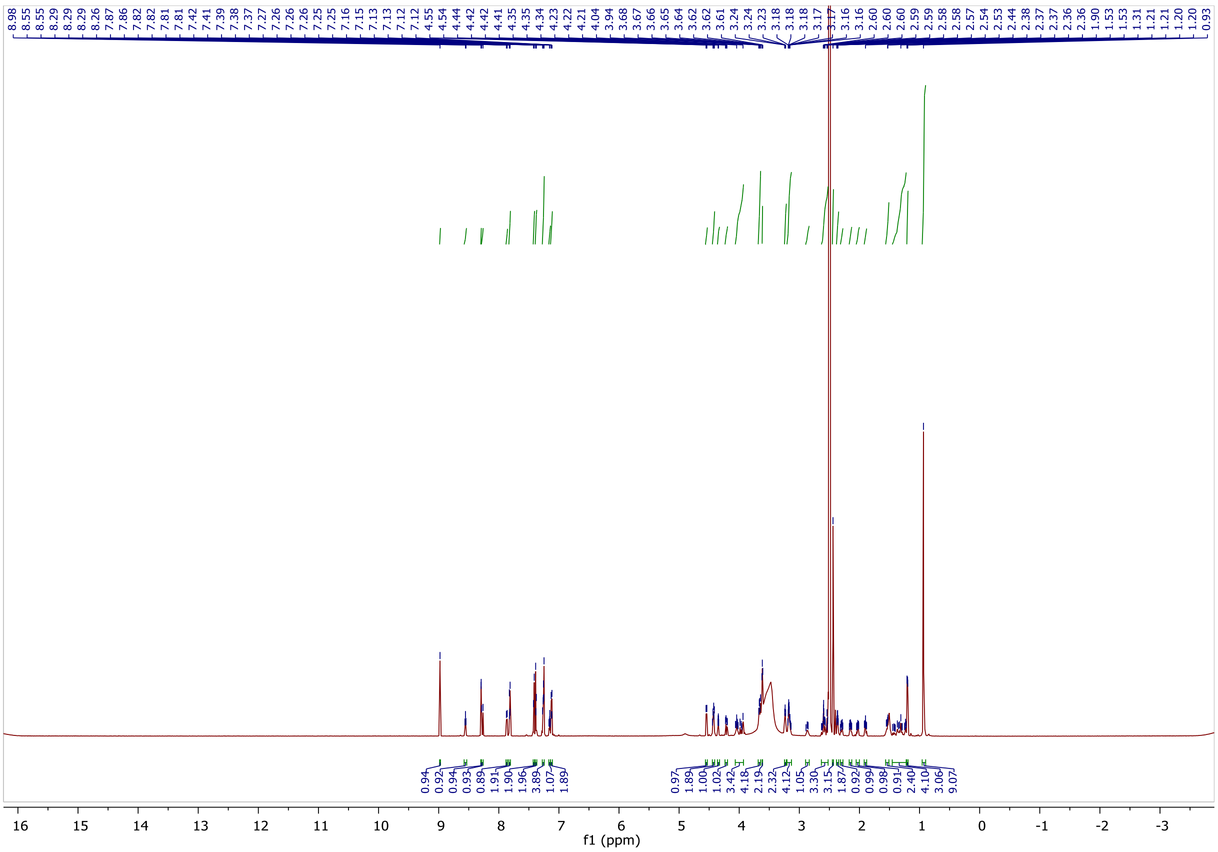

#### $^{13}$C NMR spectra of NK222.

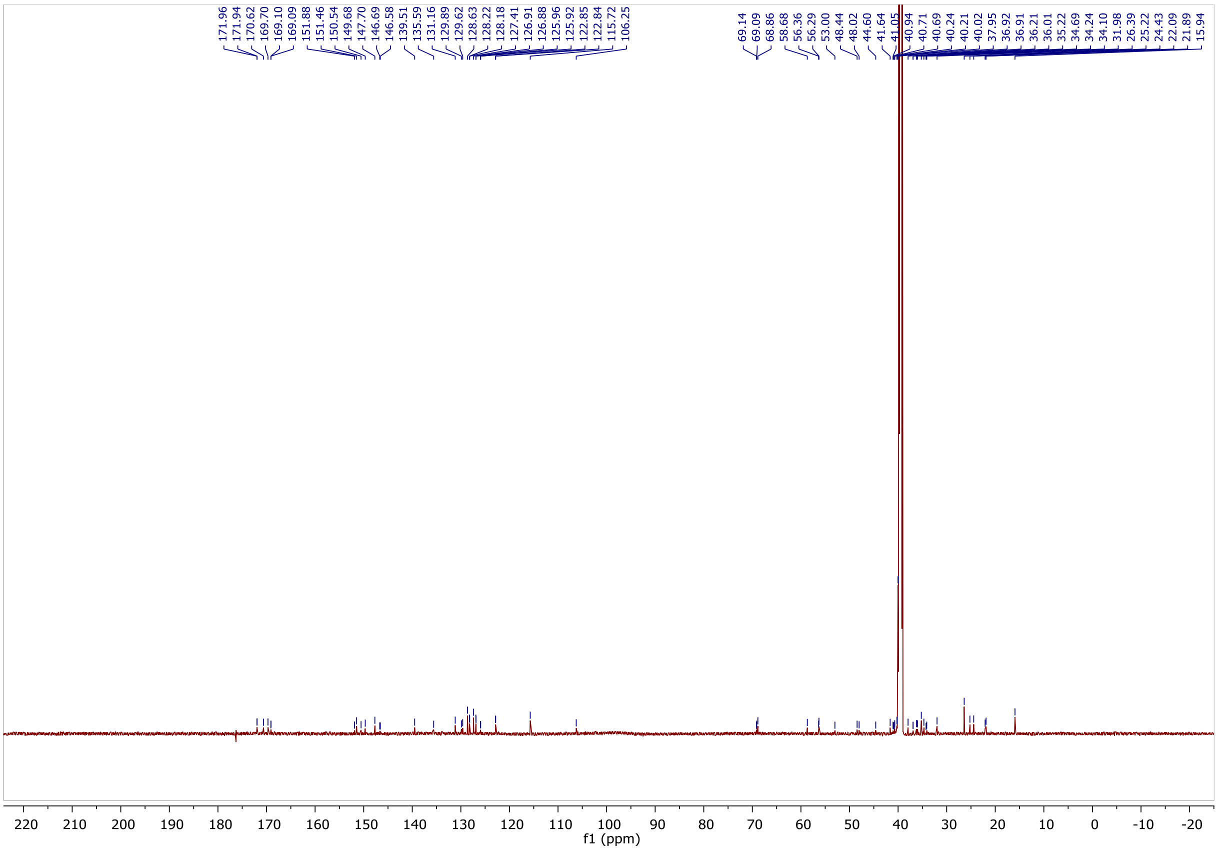

$^{1}H$ NMR spectra of NK223.

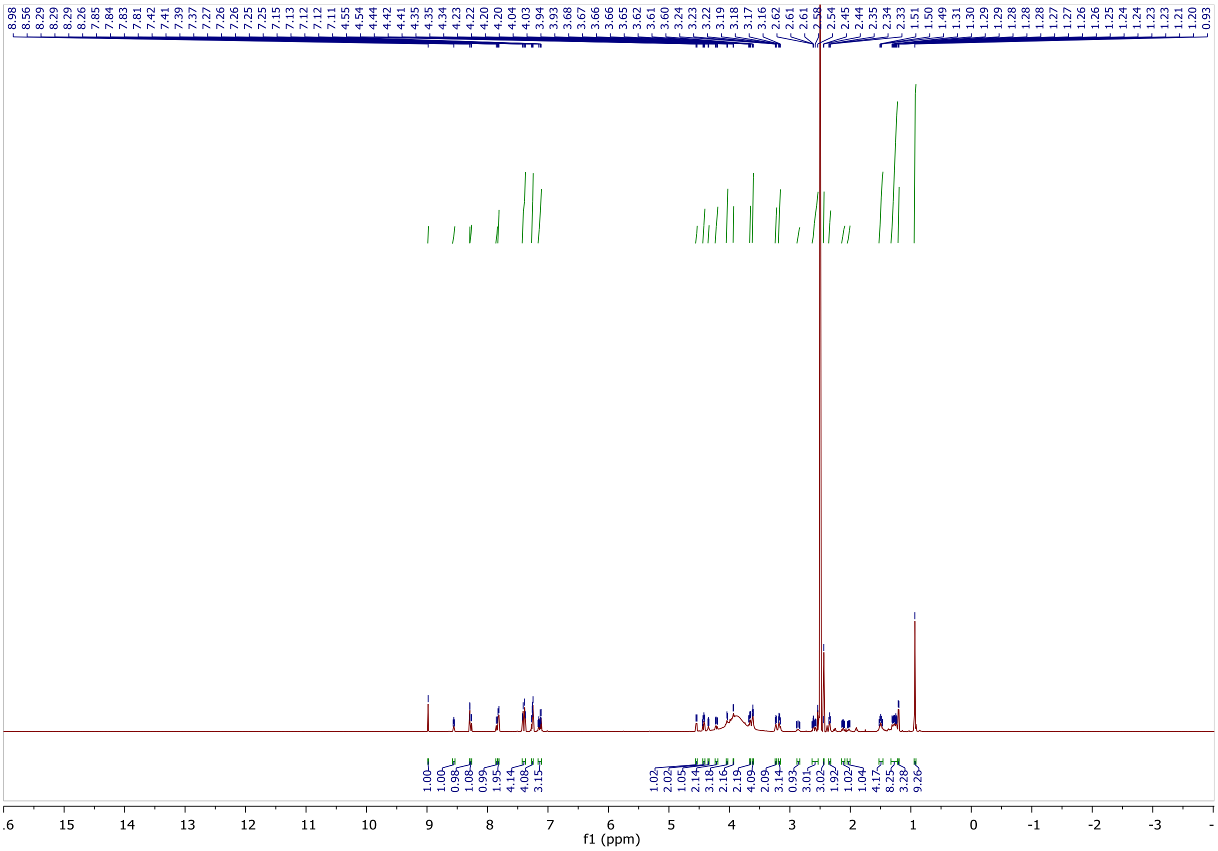

$^{13}C$ NMR spectra of NK223.

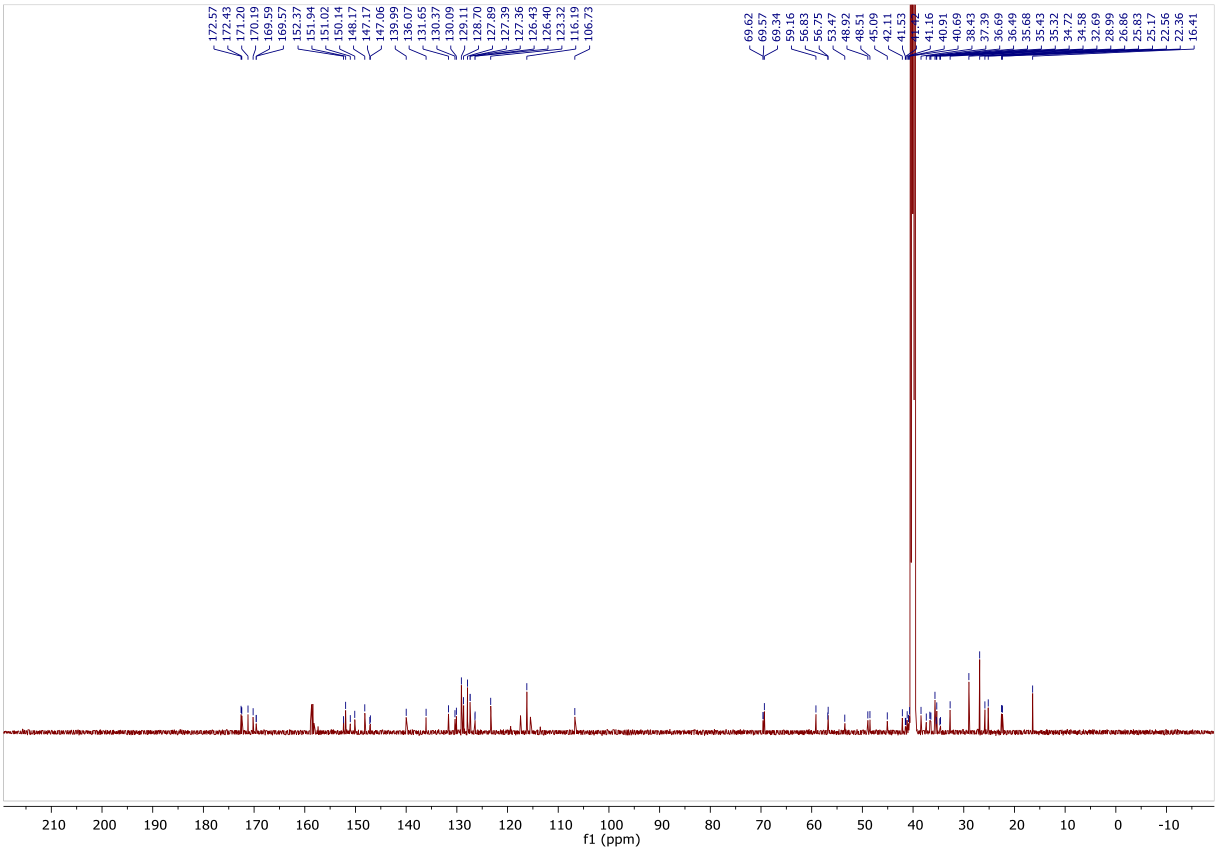

<sup></sup>

$^{1}$H NMR spectra of NK224.

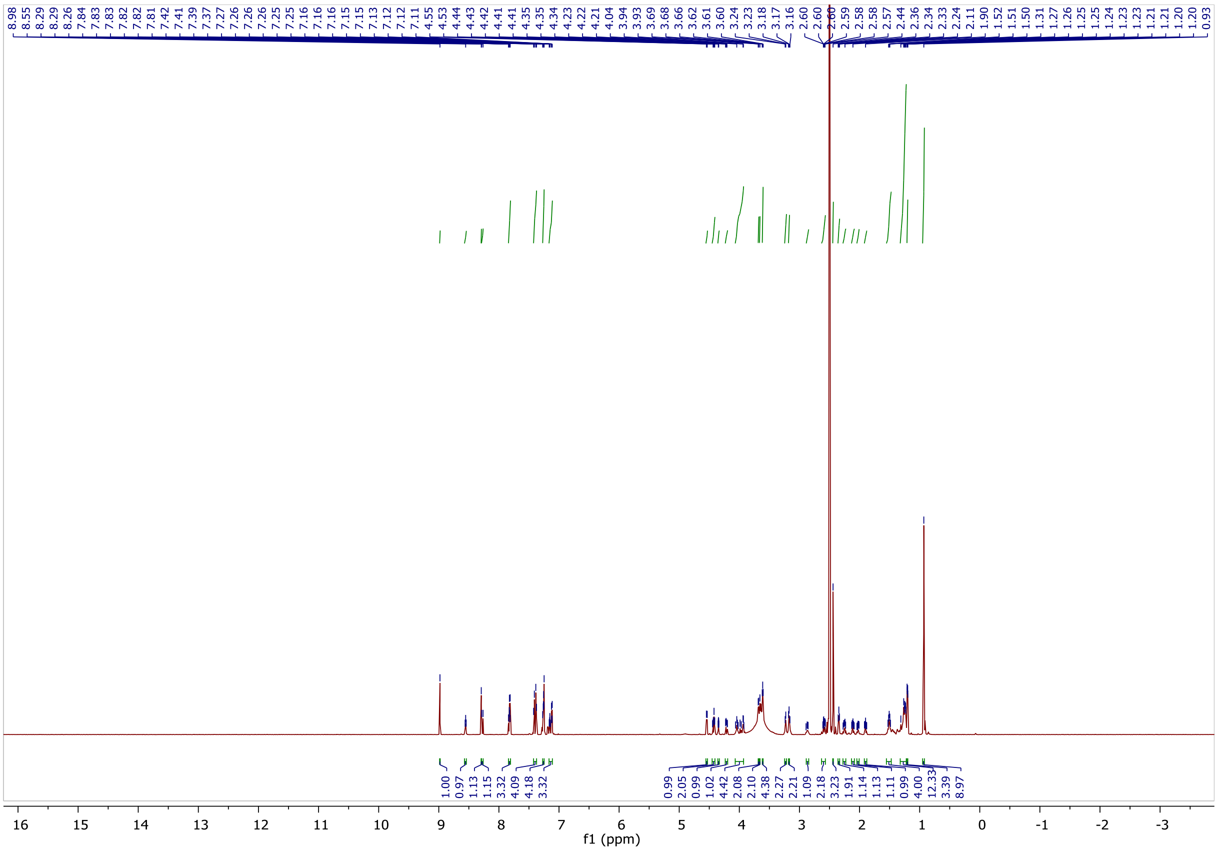

$^{13}$C NMR spectra of NK224.

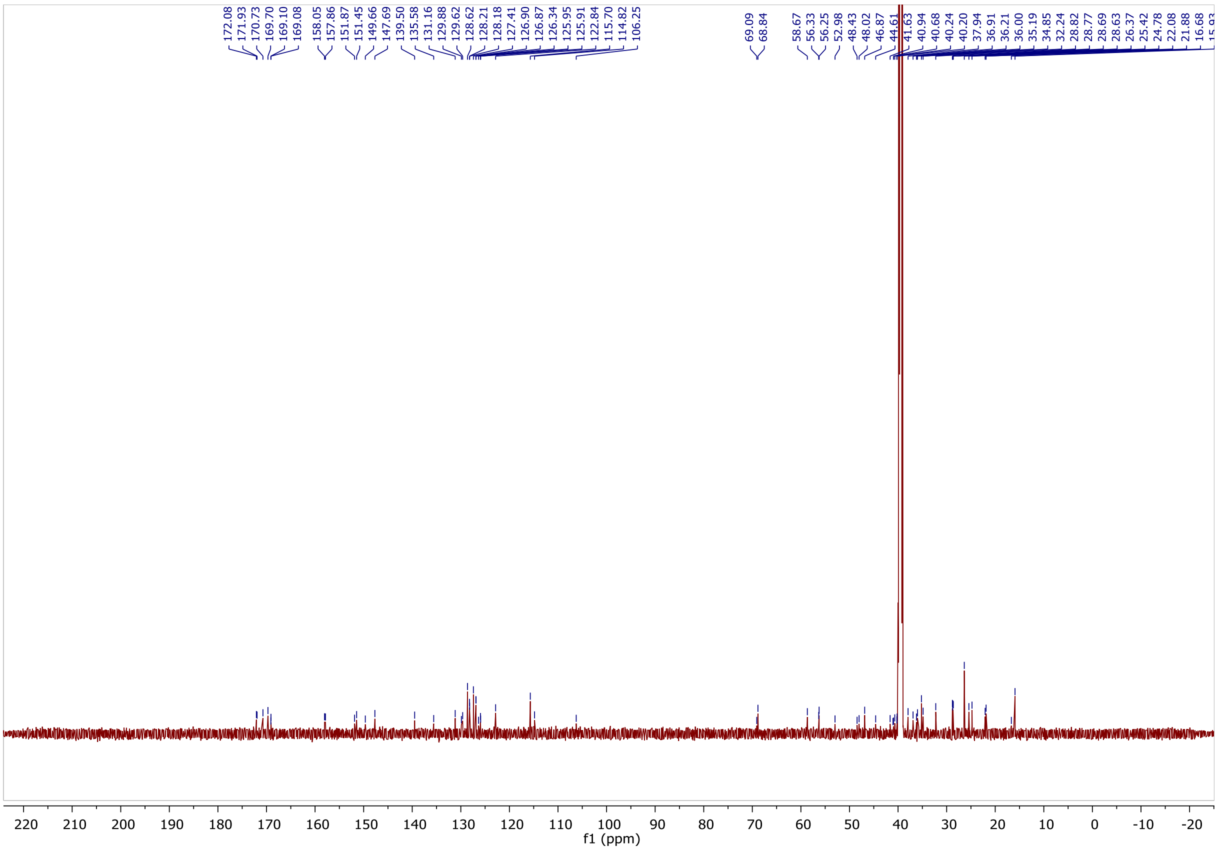

[1]H NMR spectra of NK225.

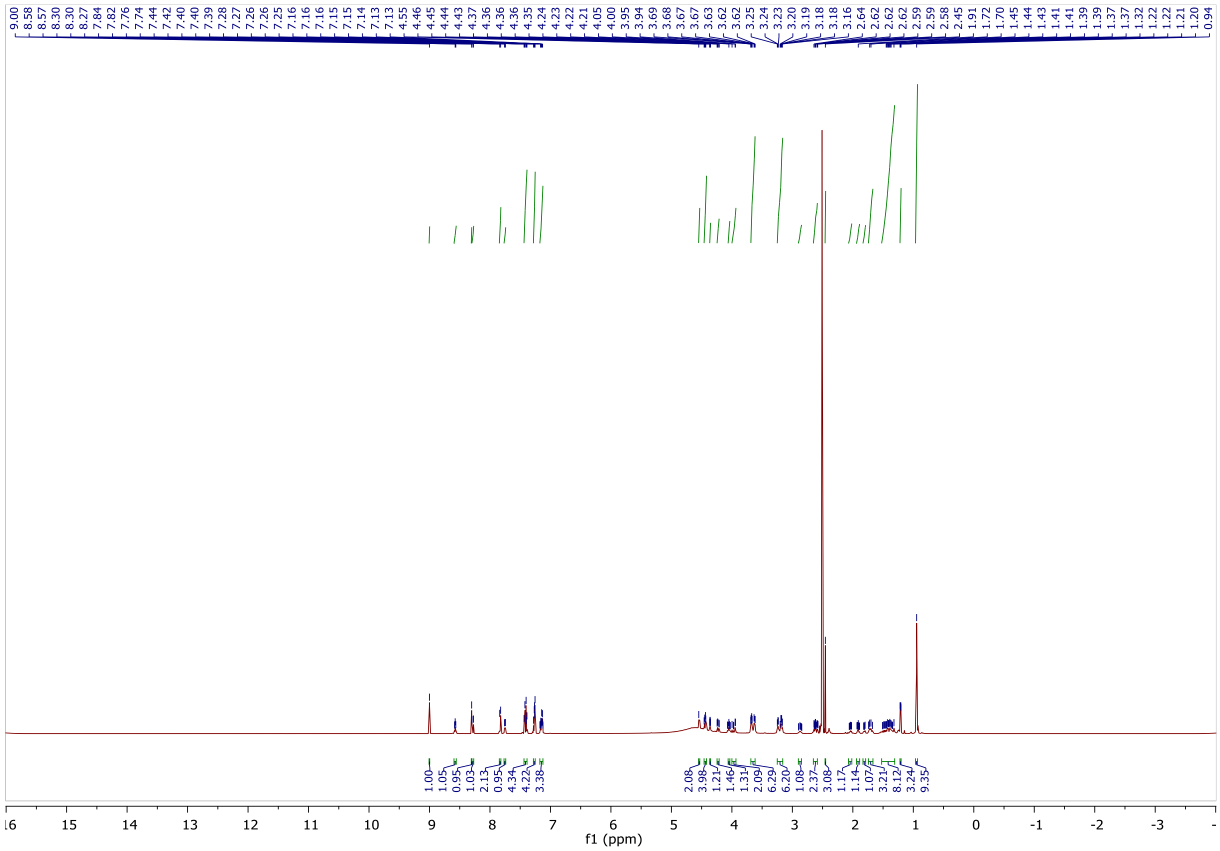

[13]C NMR spectra of NK225.

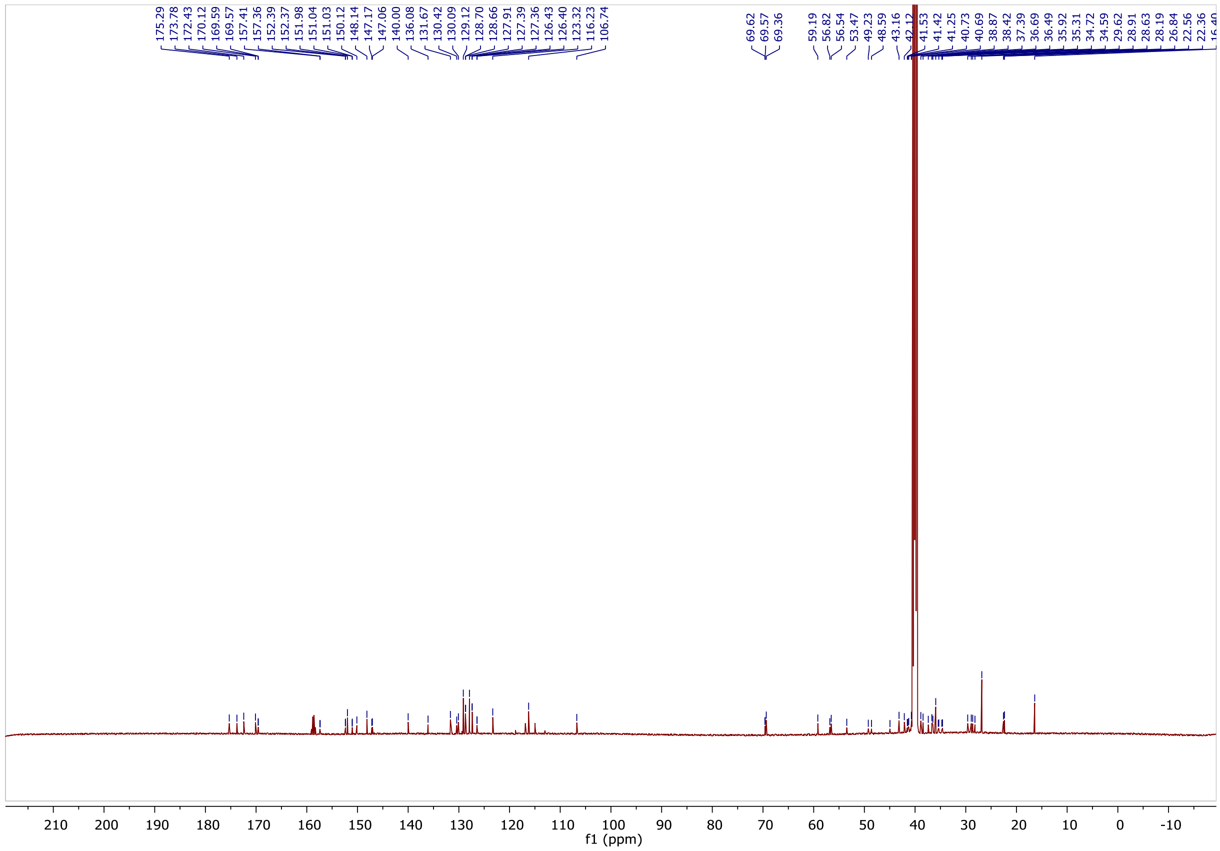

$^{1}H$ NMR spectra of NK228.

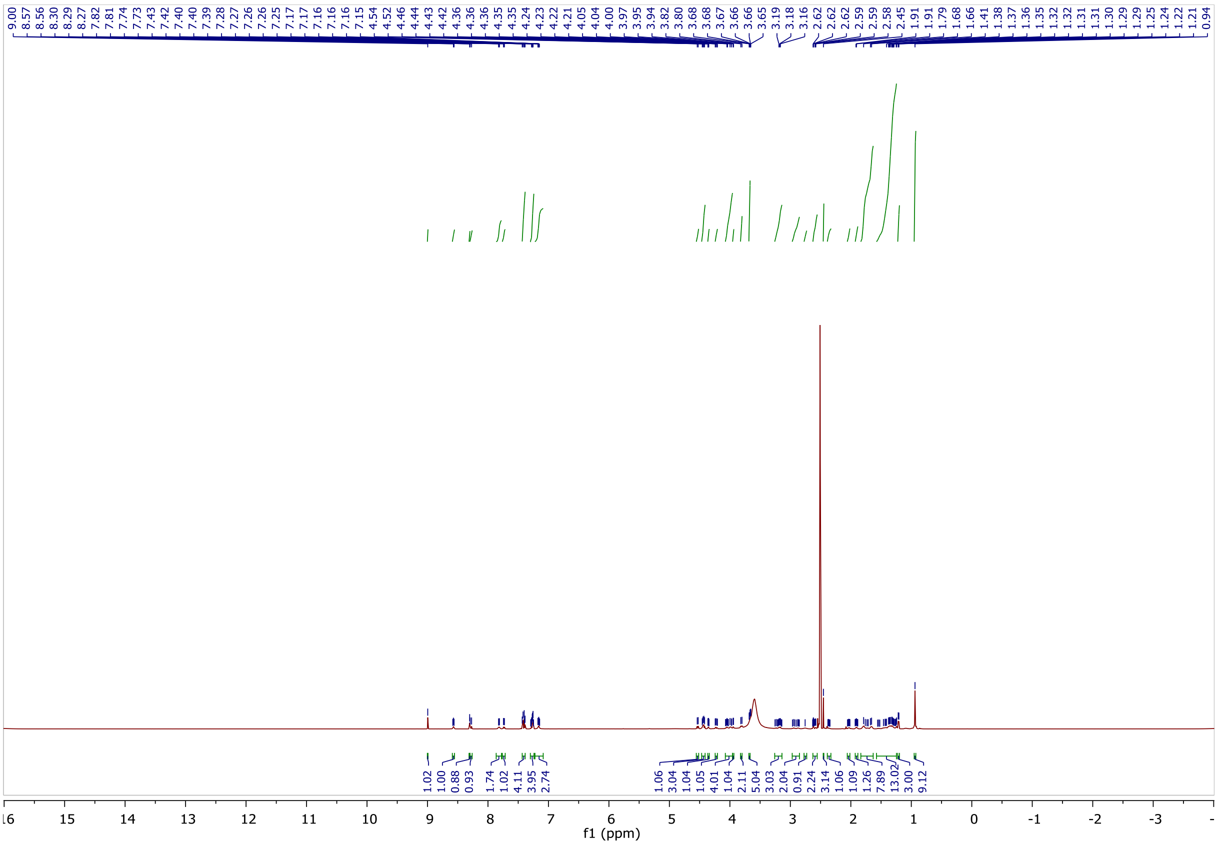

$^{13}C$ NMR spectra of NK228.

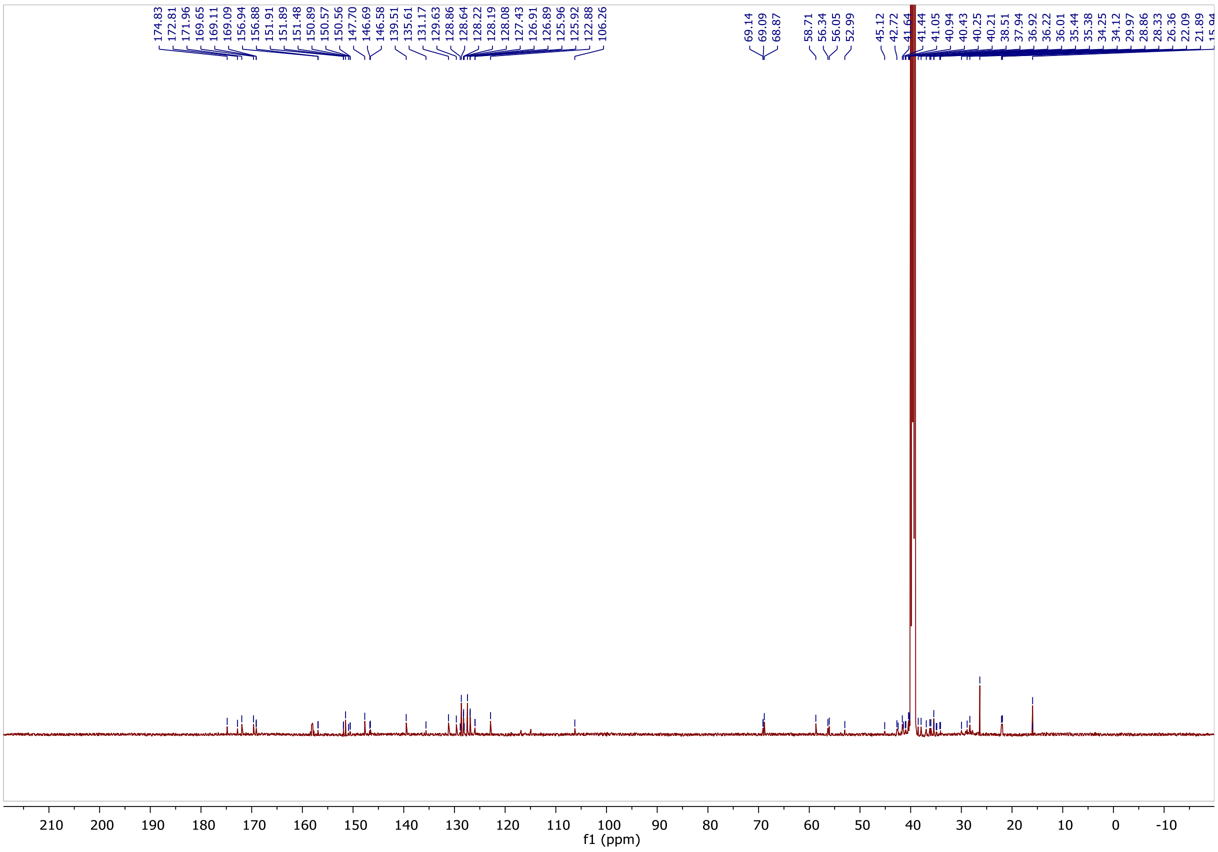

[1]H NMR spectra of NK230.

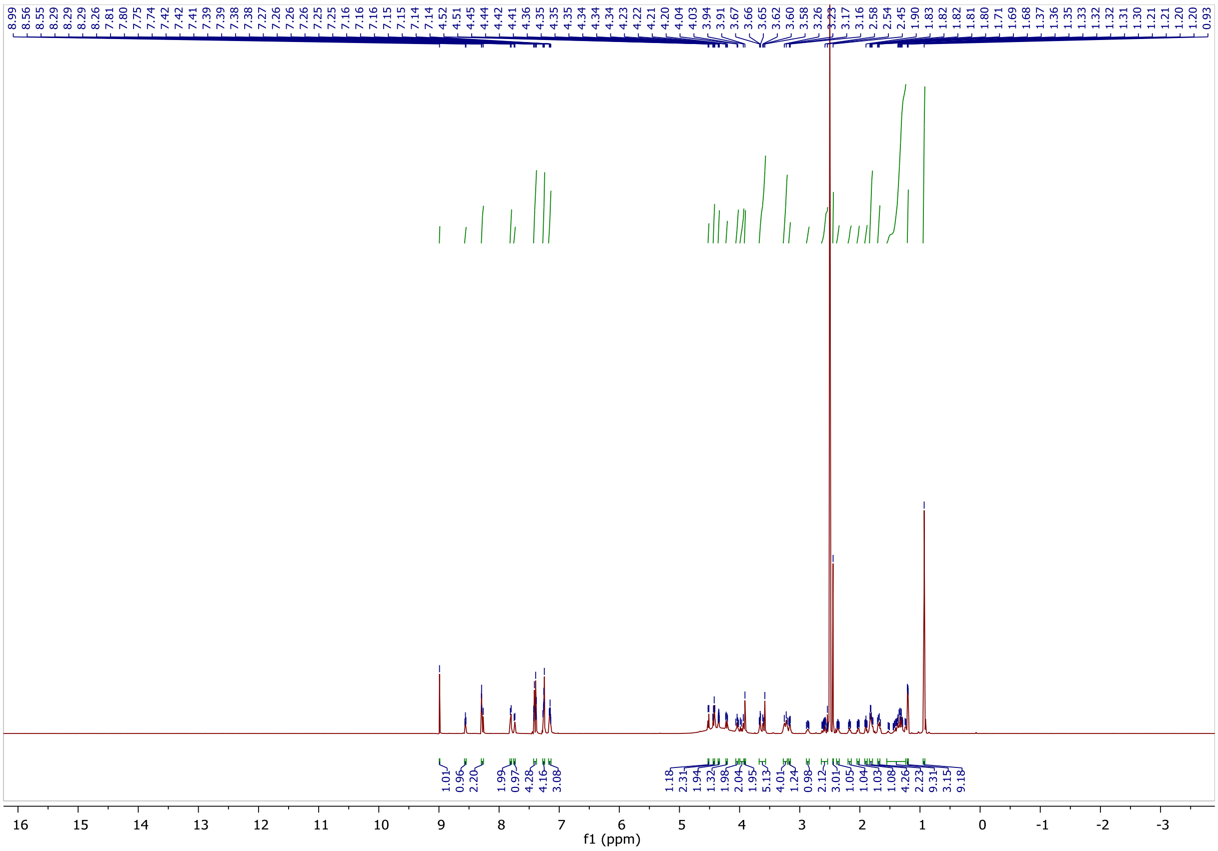

[13]C NMR spectra of NK230.

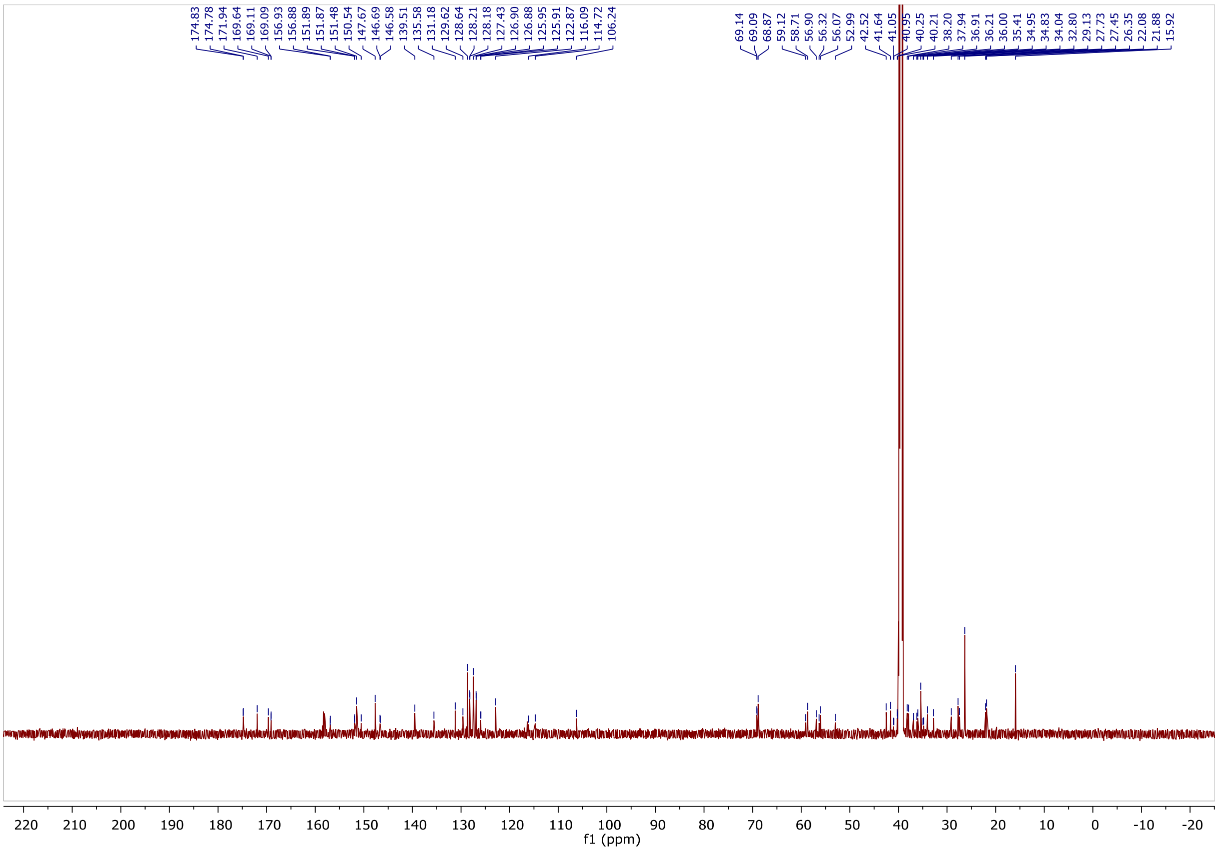

$^1$H NMR spectra of NK232.

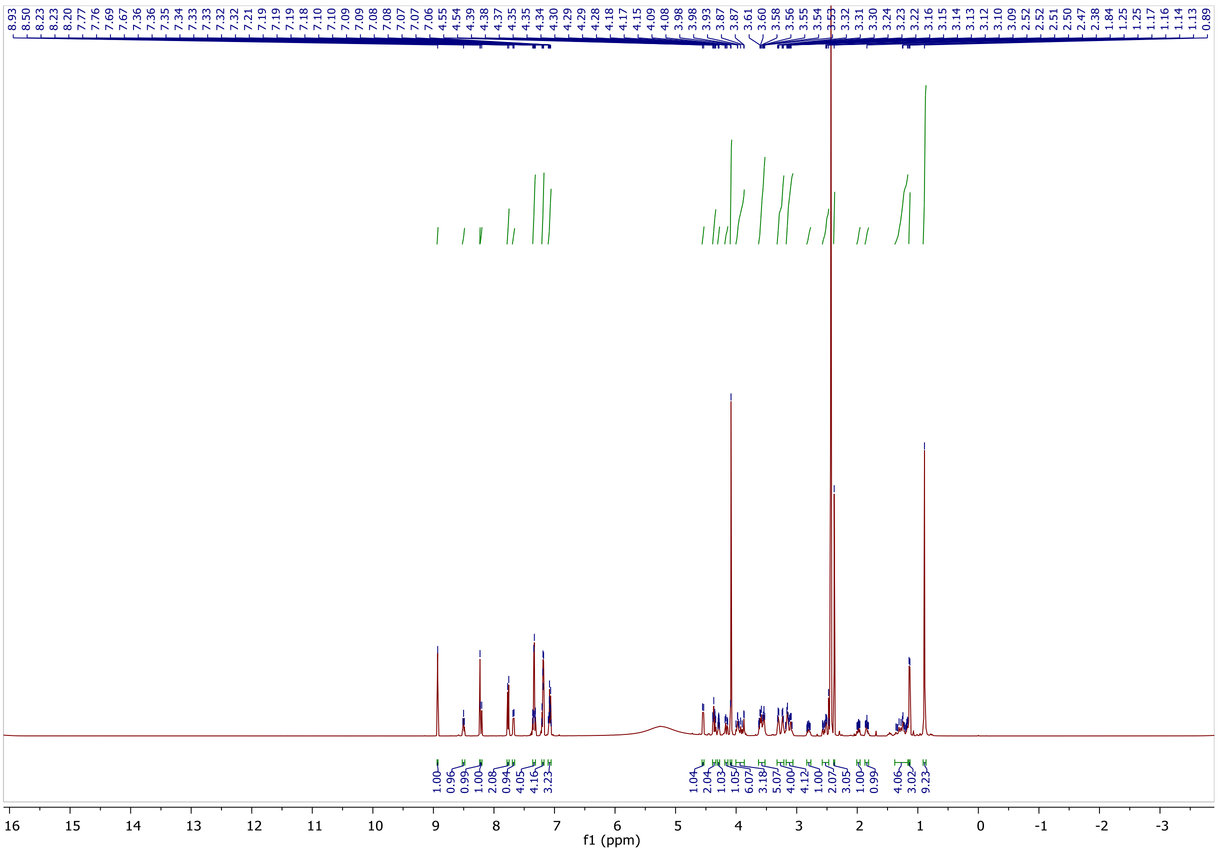

$^{13}$C NMR spectra of NK232.

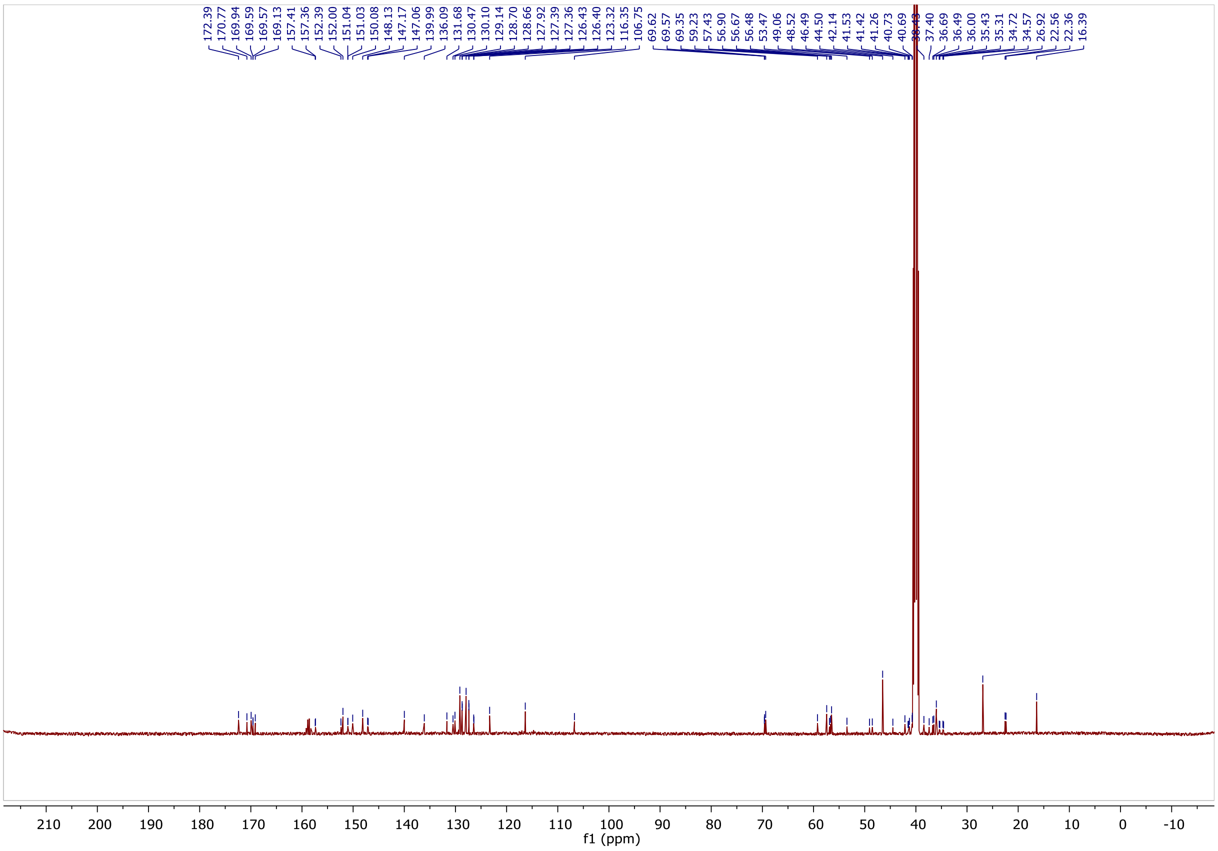

$^{1}H$ NMR spectra of NK233.

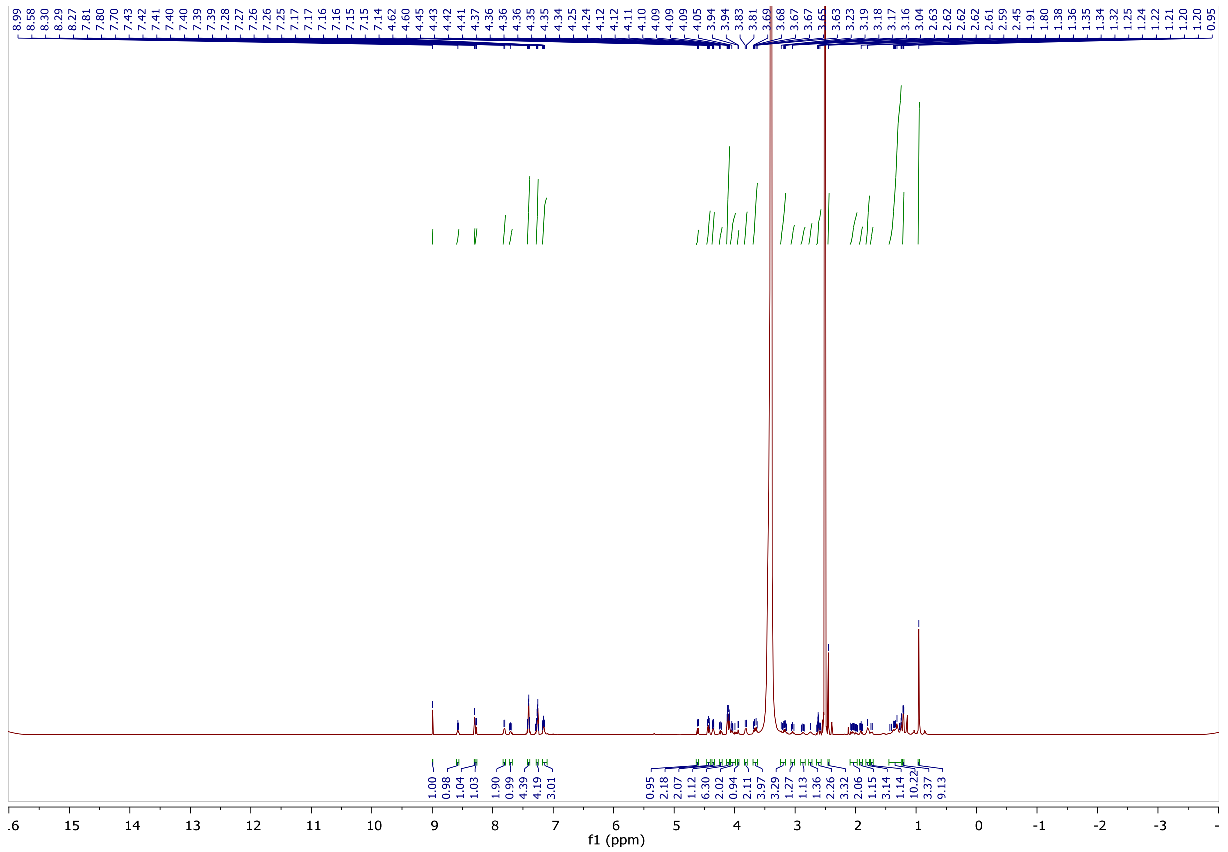

$^{13}C$ NMR spectra of NK233.

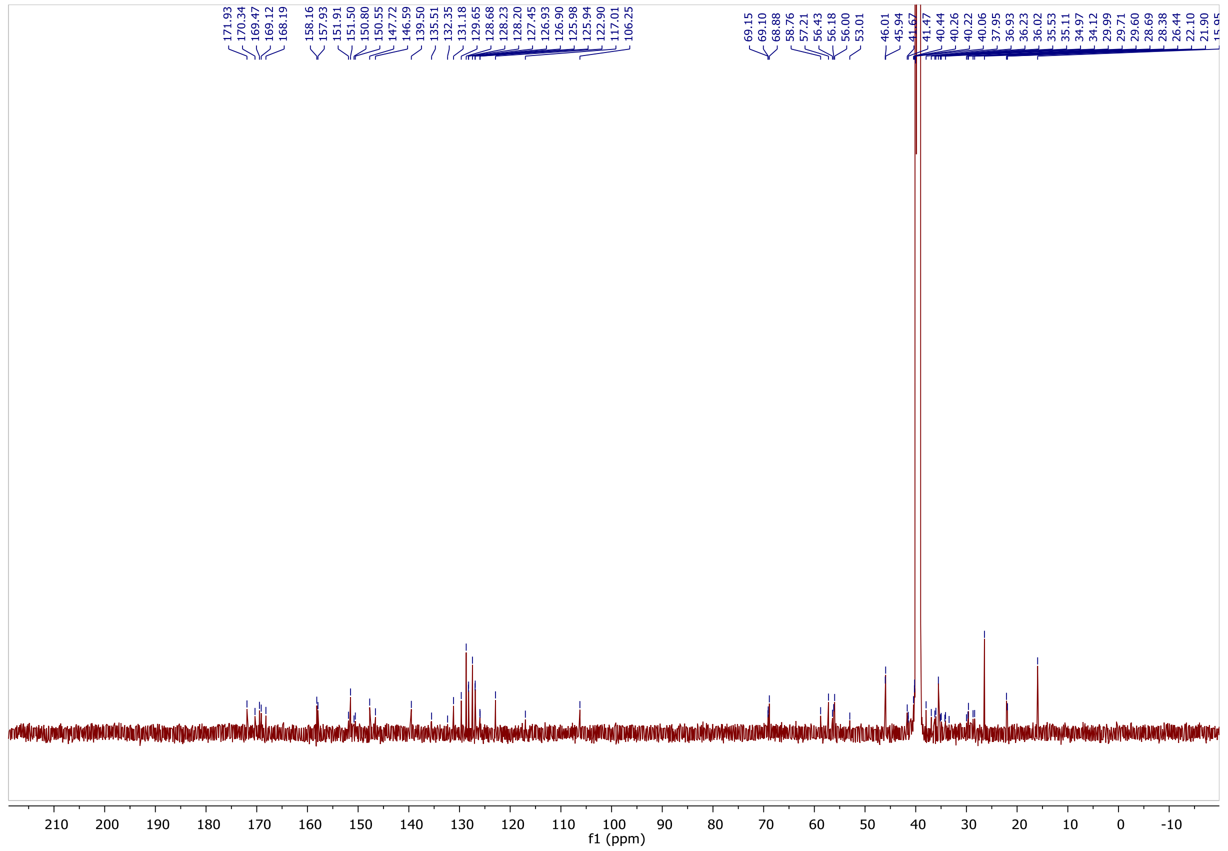

$^{1}H$ NMR spectra of NK234.

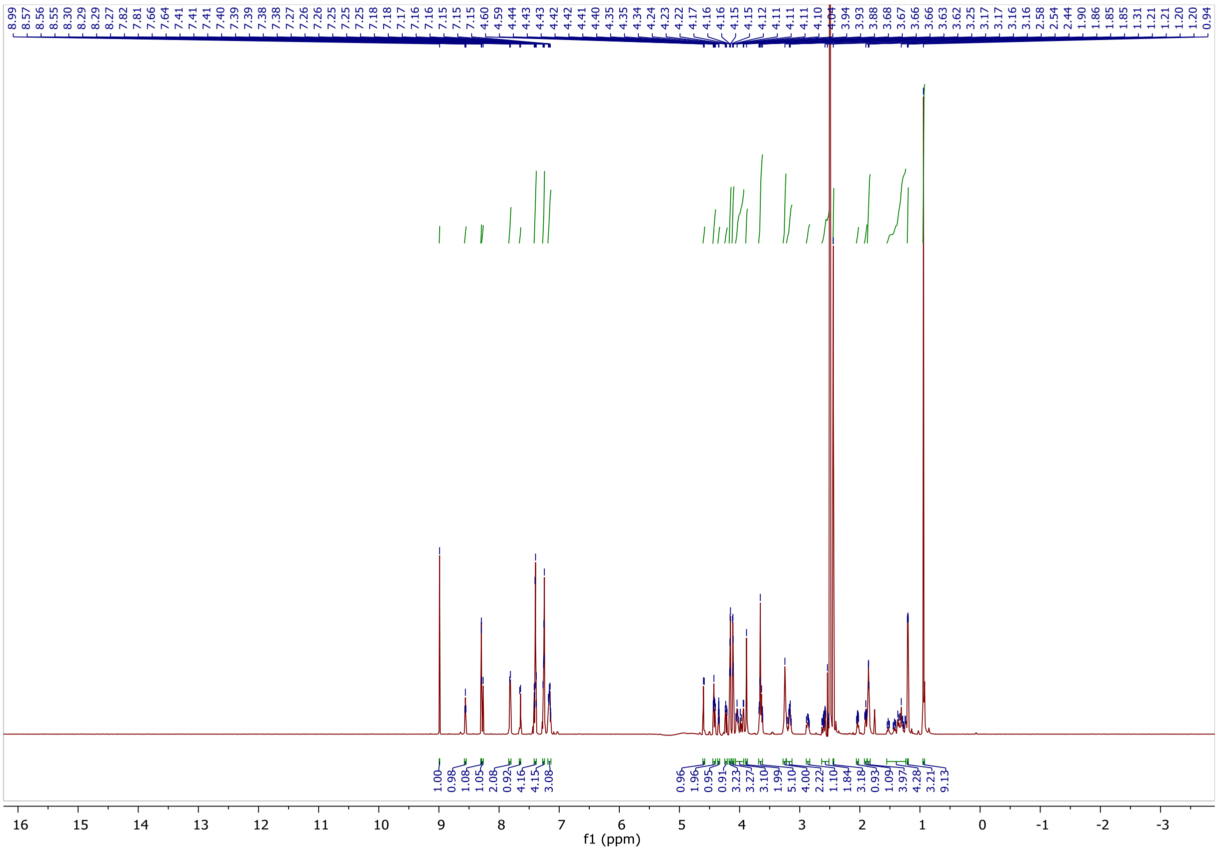

$^{13}C$ NMR spectra of NK234.

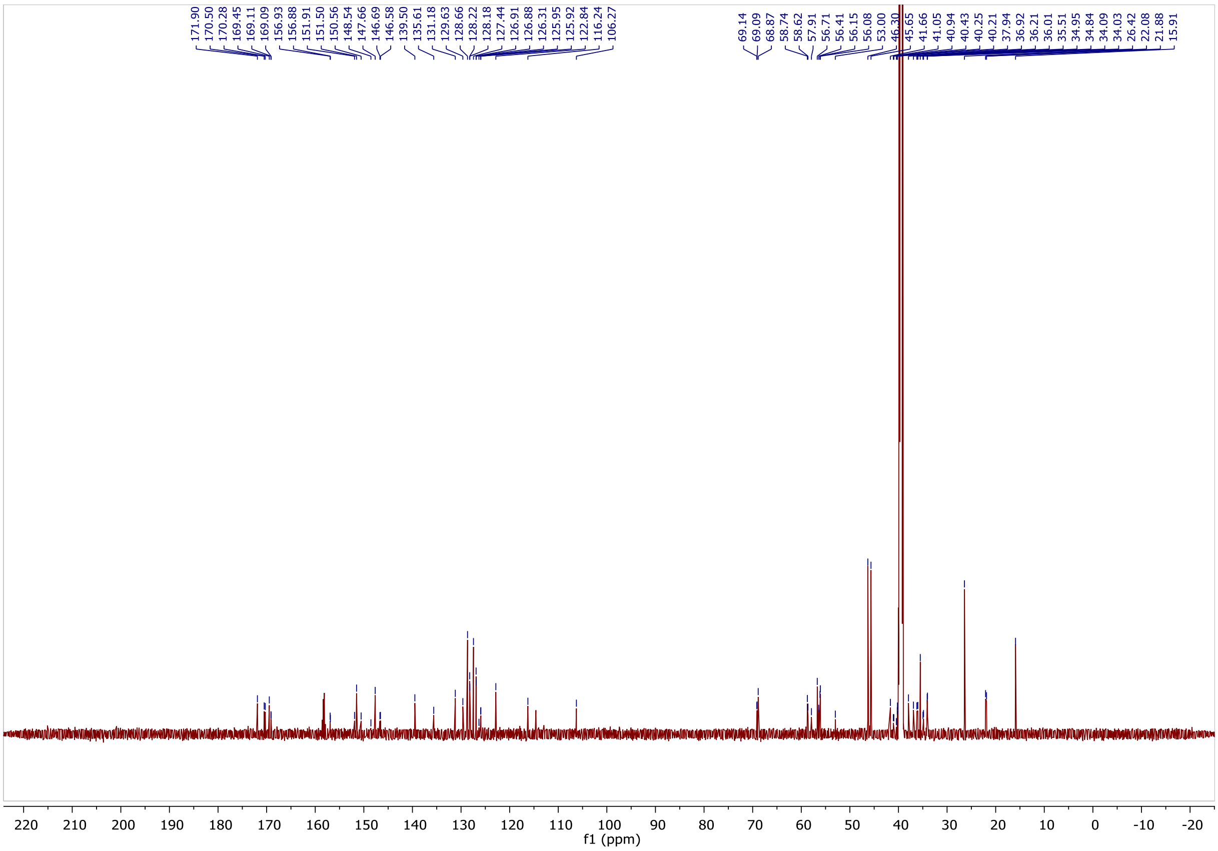

[1]H NMR spectra of NK250.

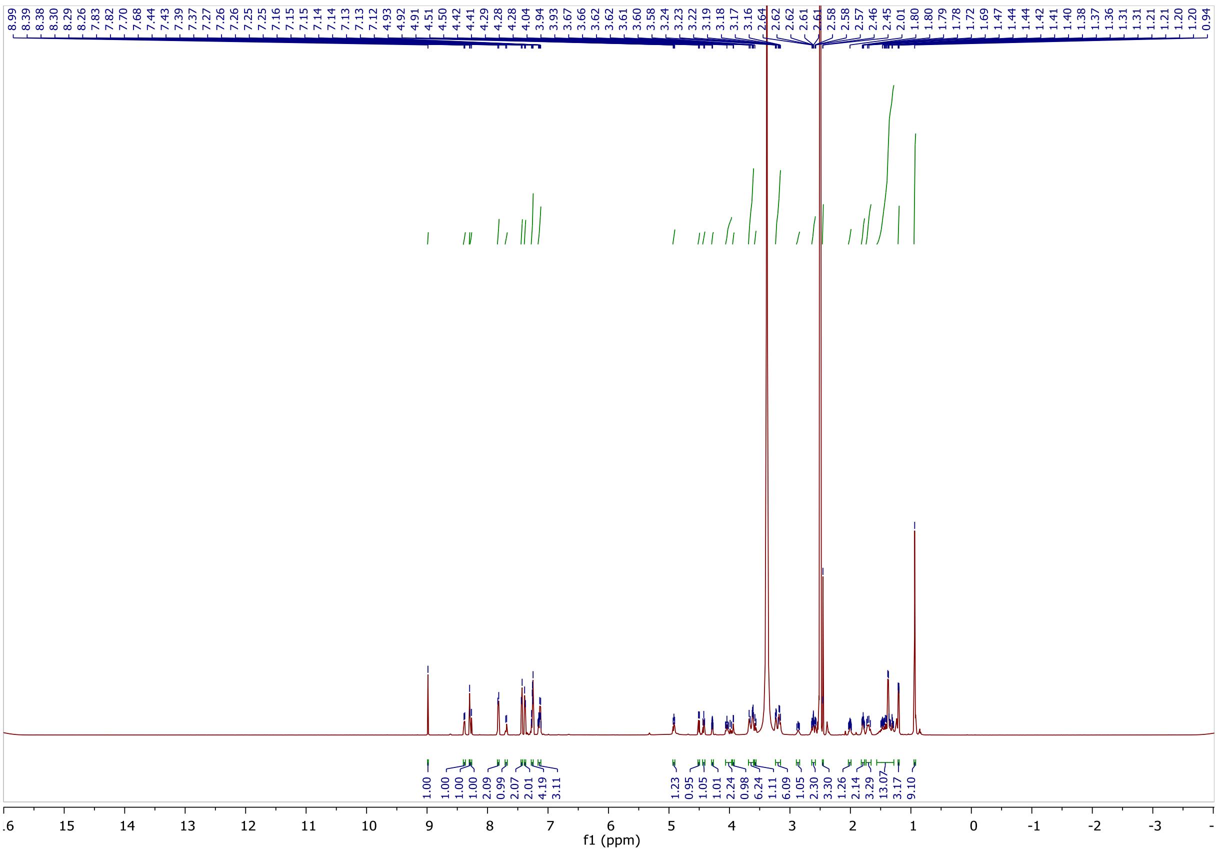

[13]C NMR spectra of NK250.

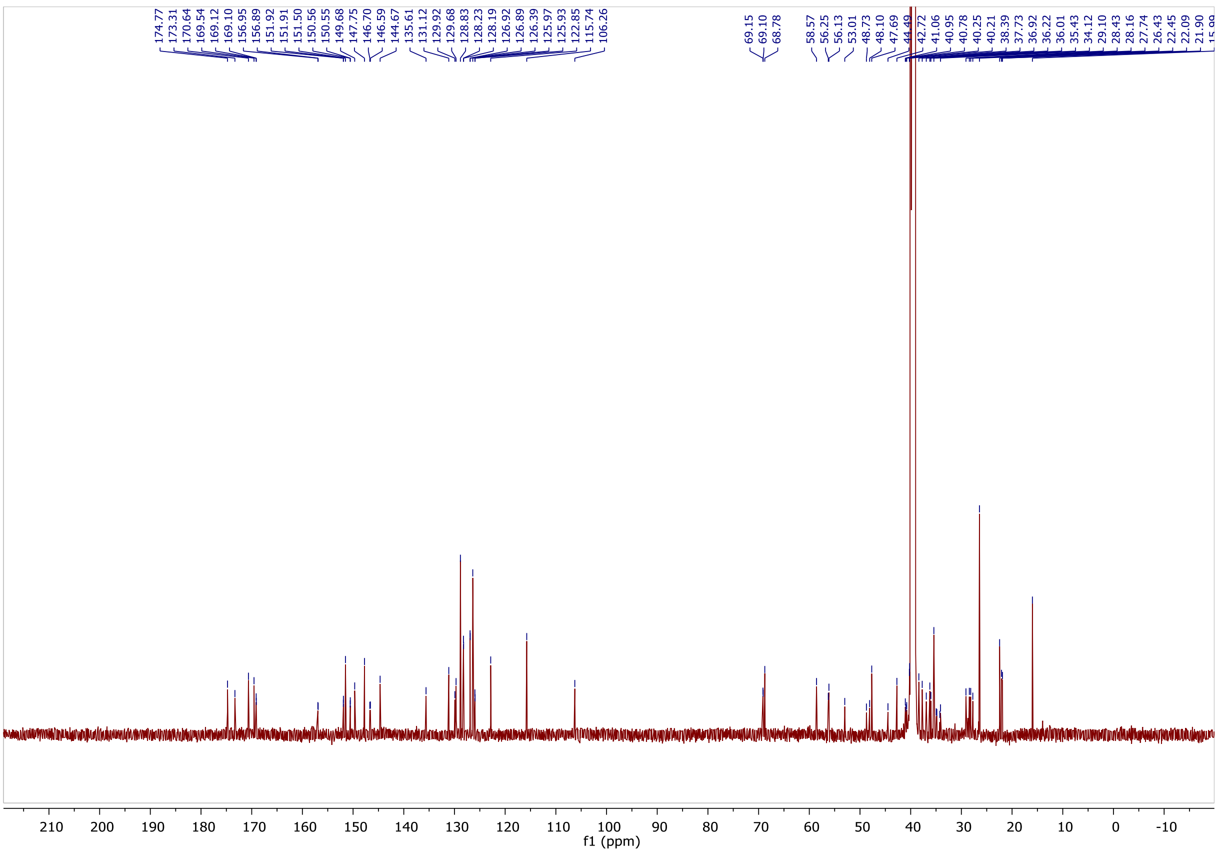

$^{1}H$ NMR spectra of NK266.

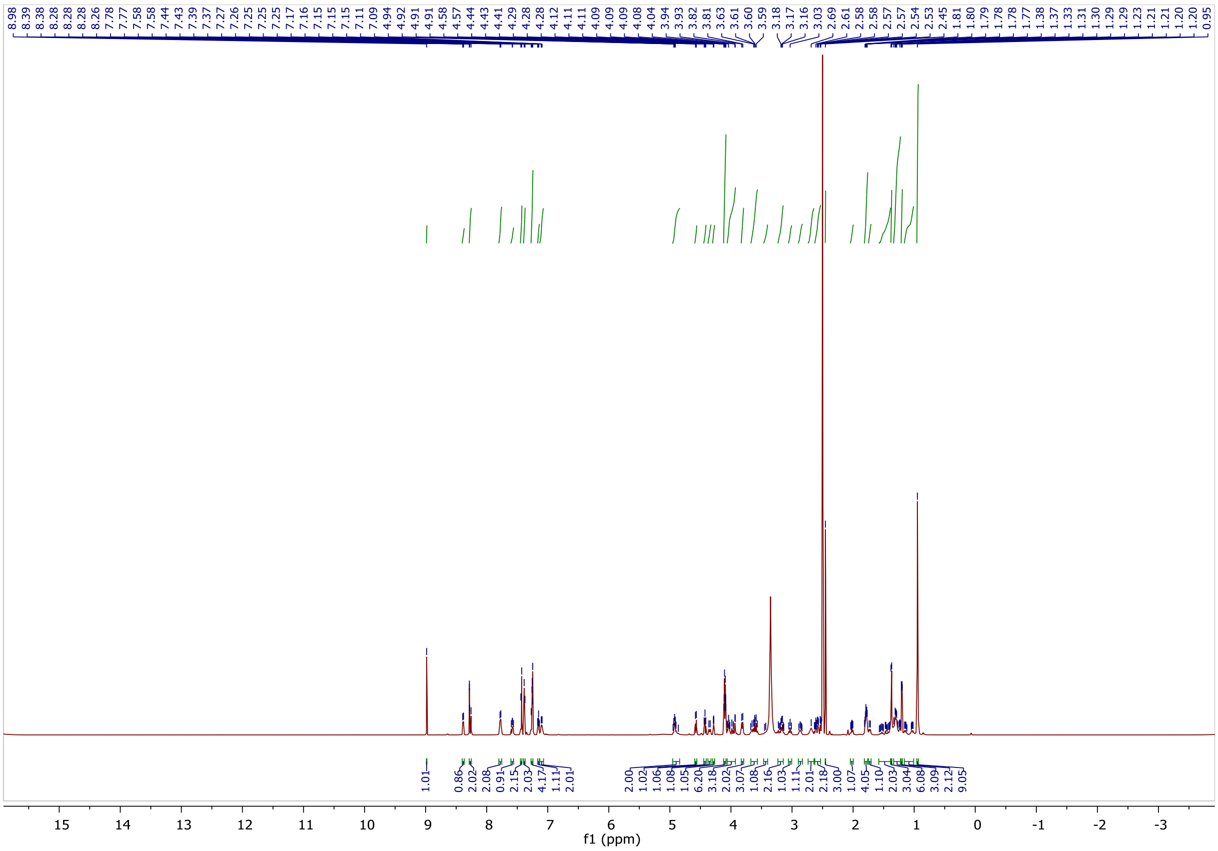

$^{13}C$ NMR spectra of NK266.

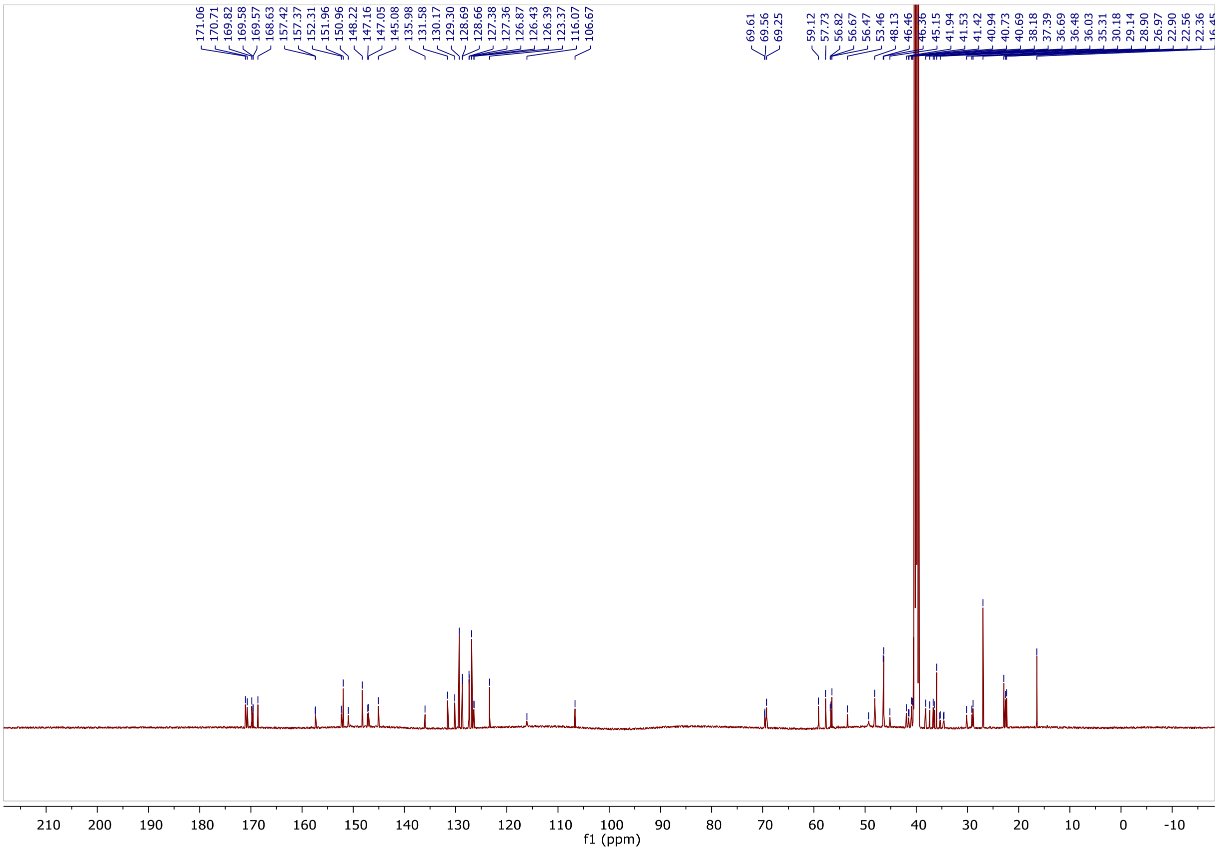

$^{1}H$ NMR spectra of NK249.

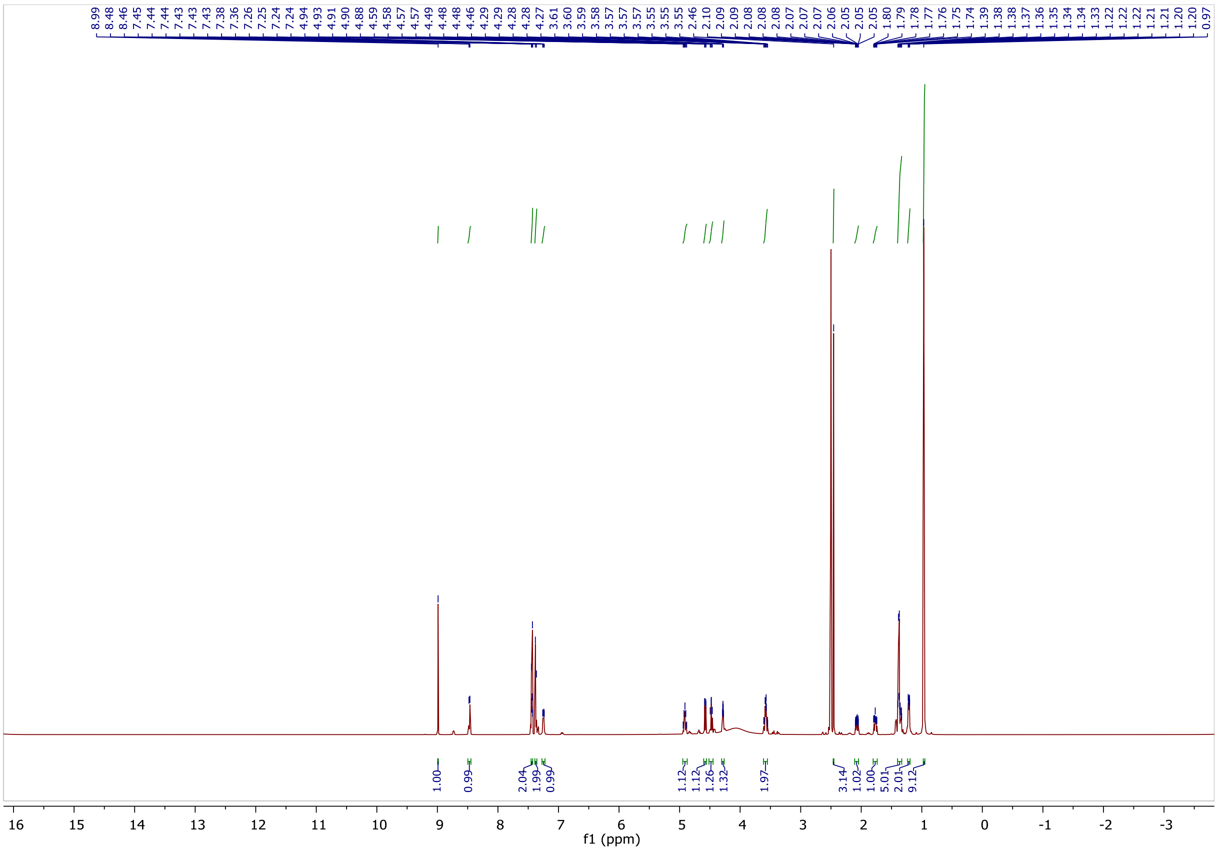

$^{13}C$ NMR spectra of NK249.

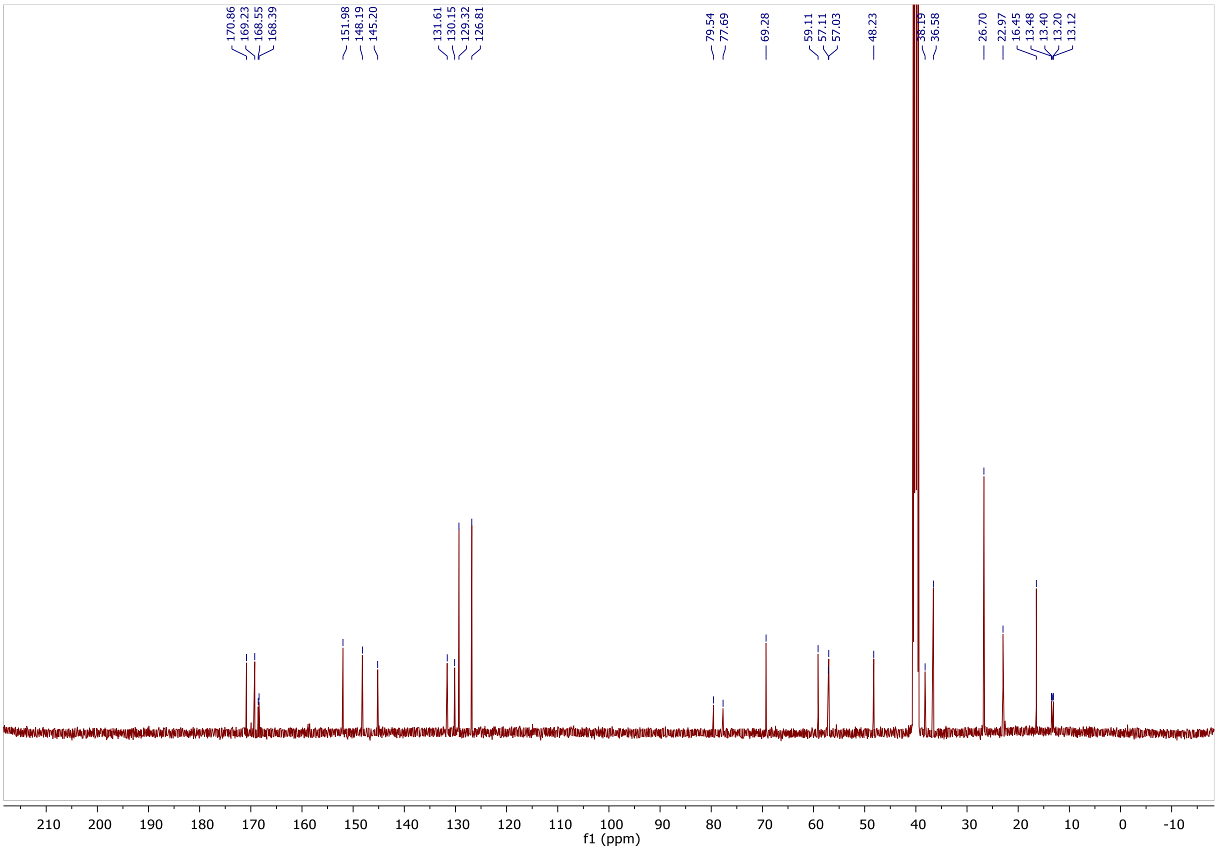

#### 5. Uncropped gels and blots

### Fig. 2c

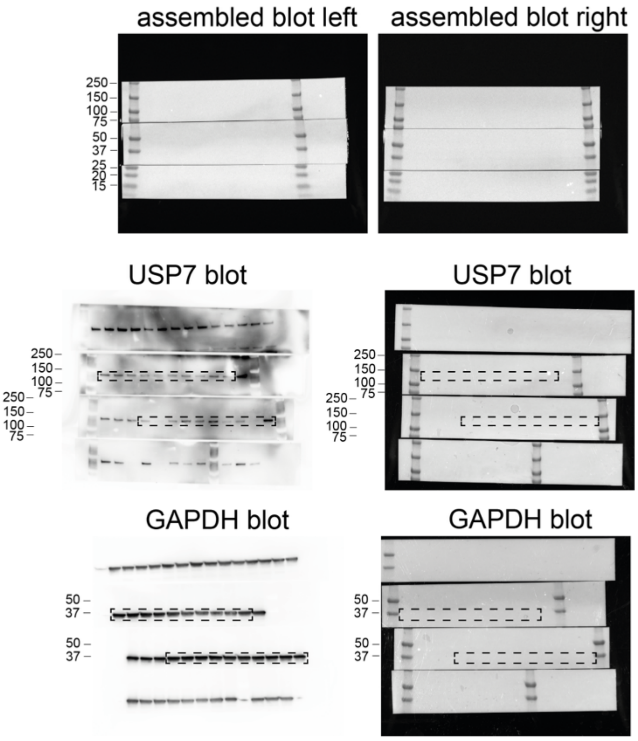

### Fig. 2d

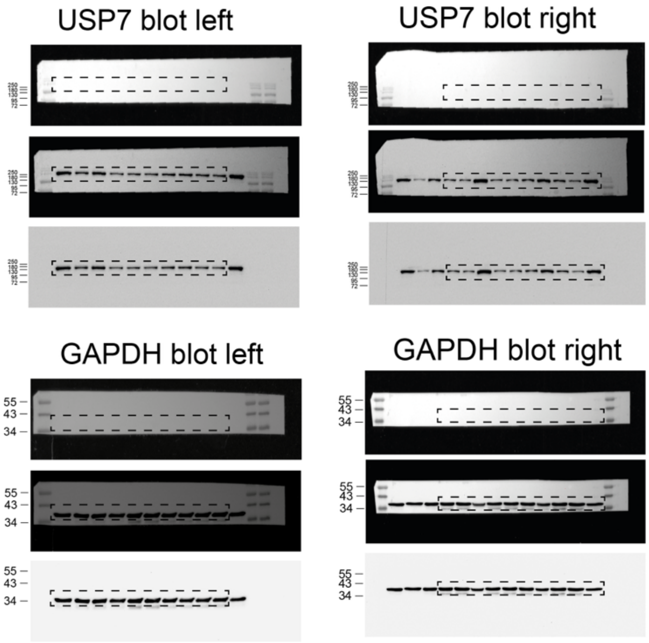

#### Fig. 3b

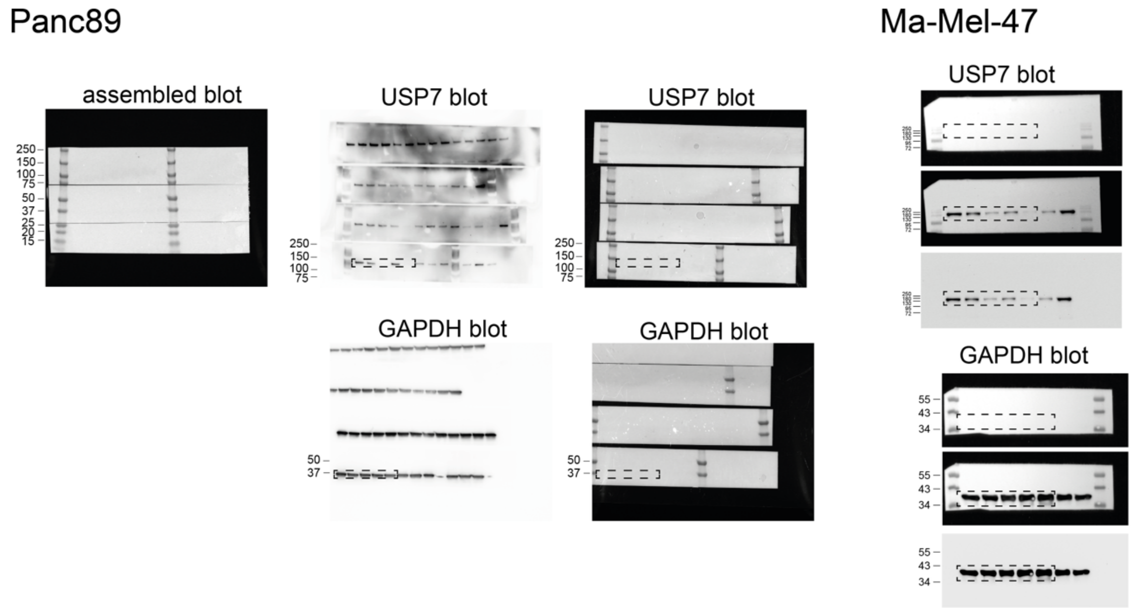
Panc89
Ma-Mel-47
assembled blot
USP7 blot
USP7 blot
USP7 blot
GAPDH blot
GAPDH blot
GAPDH blot
250
150
100
75
50
37
25
20
15
55
43
34

#### Fig. 3c

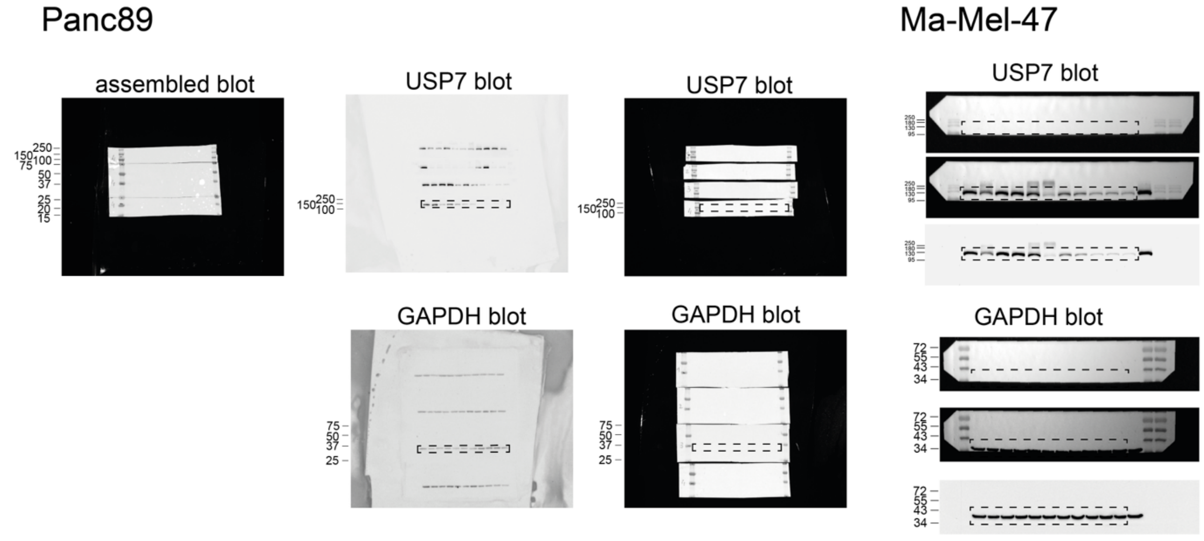
Panc89
Ma-Mel-47
assembled blot
USP7 blot
USP7 blot
USP7 blot
GAPDH blot
GAPDH blot
GAPDH blot
250
150
100
75
50
37
25
20
15
72
55
43
34

### Fig. 3d

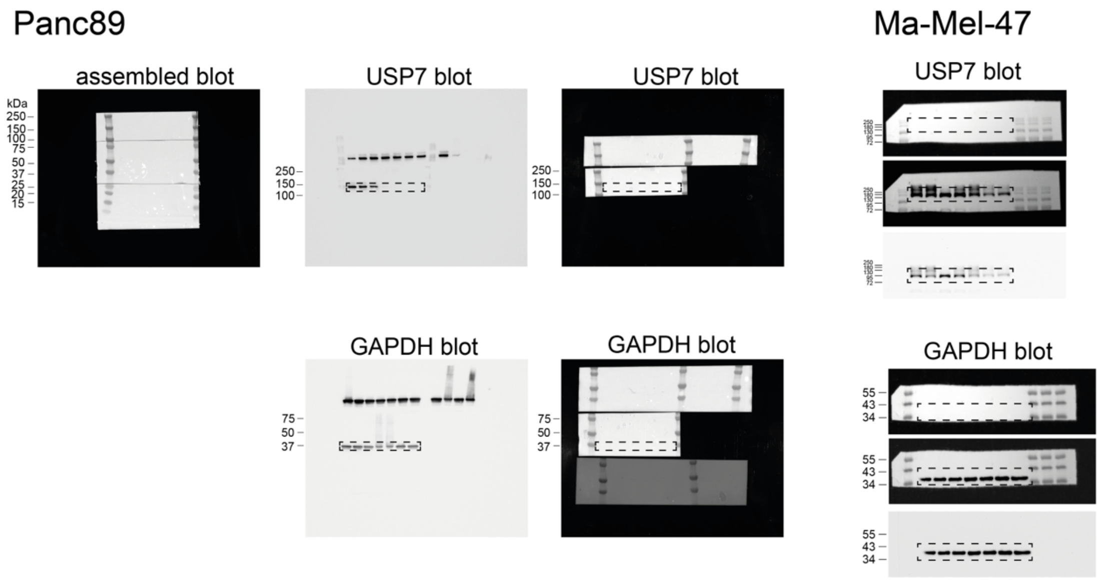
Panc89
Ma-Mel-47
assembled blot
USP7 blot
USP7 blot
USP7 blot
GAPDH blot
GAPDH blot
GAPDH blot
kDa
250
150
100
75
50
37
25
20
15
55
43
34

### Fig. 3e

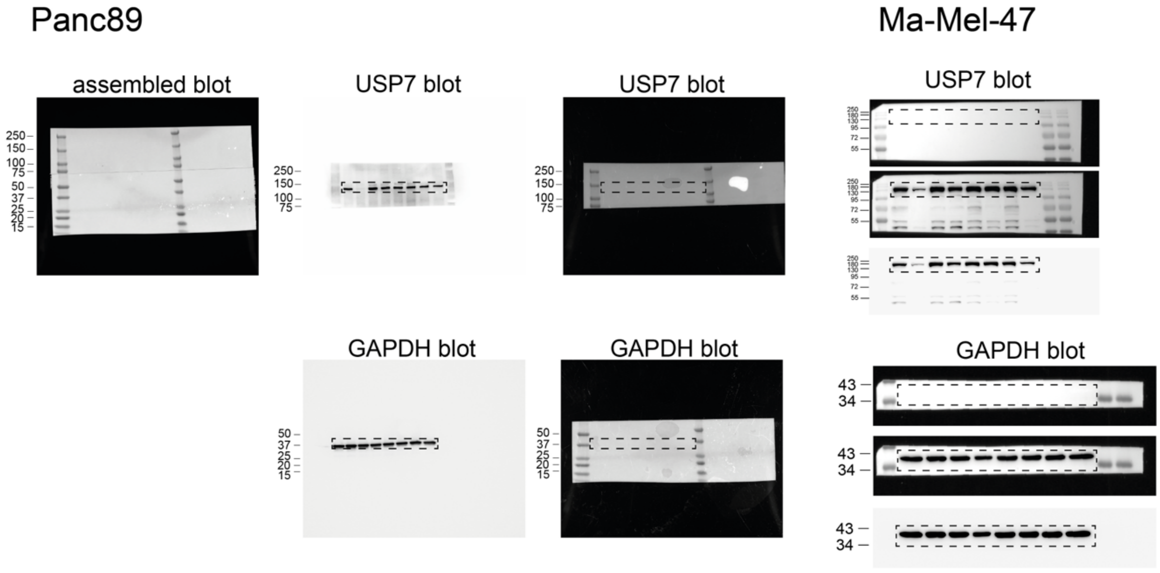
Panc89
Ma-Mel-47
assembled blot
USP7 blot
USP7 blot
USP7 blot
GAPDH blot
GAPDH blot
GAPDH blot
250
150
100
75
50
37
25
20
15
43
34

**Fig. 6b**

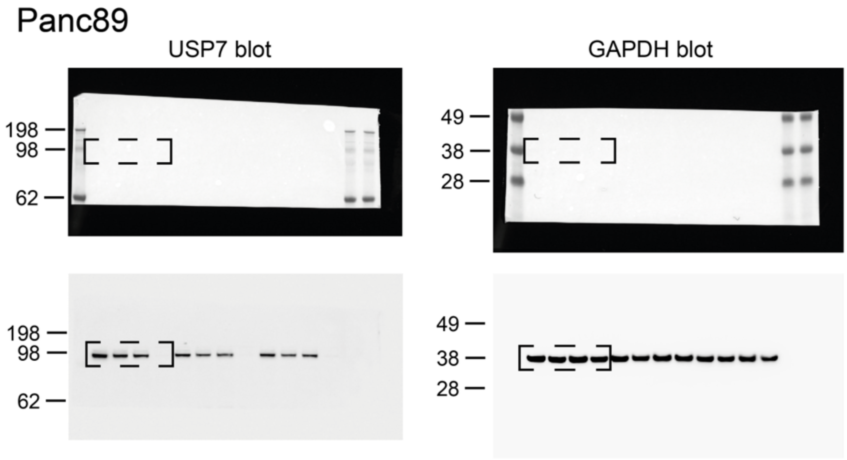

#### Supporting Fig. 2a

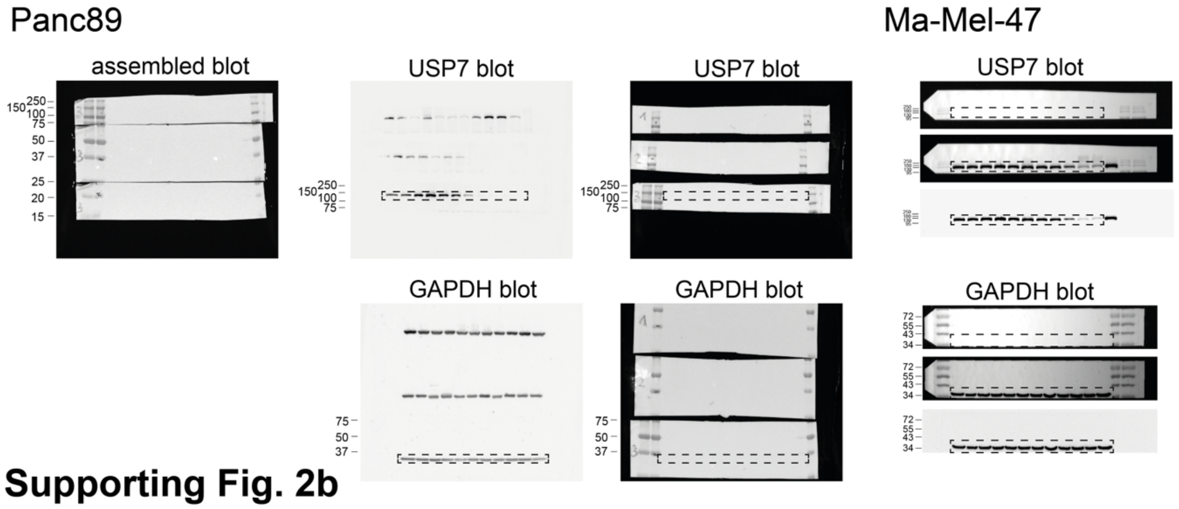

#### Supporting Fig. 2b

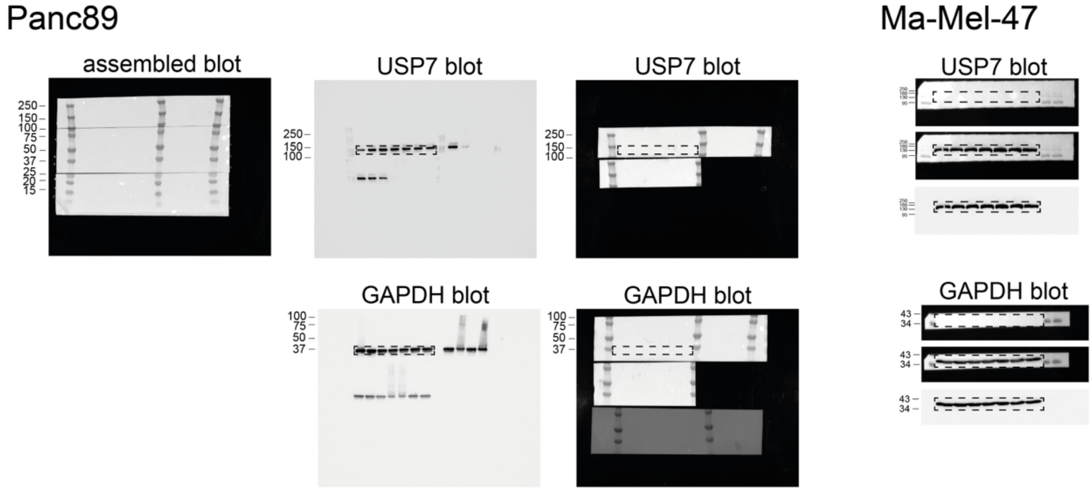

#### Supporting Fig. 6d

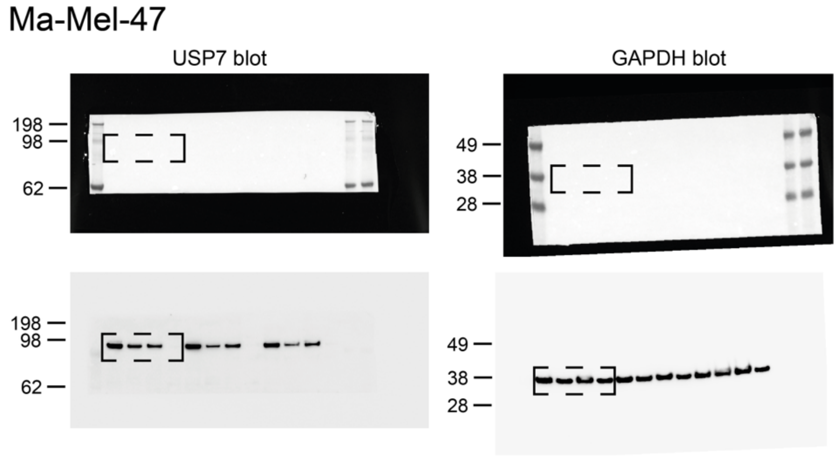

#### 6. Supporting References

1. Gavory, G., O'Dowd, C.R., Helm, M.D., Flasz, J., Arkoudis, E., Dossang, A., Hughes, C., Cassidy, E., McClelland, K., Odrzywol, E., Page, N., Barker, O., Miel, H. & Harrison, T. Discovery and characterization of highly potent and selective allosteric USP7 inhibitors. *Nat Chem Biol* **14**, 118-125 (2018).

2. Turnbull, A.P., Ioannidis, S., Krajewski, W.W., Pinto-Fernandez, A., Heride, C., Martin, A.C.L., Tonkin, L.M., Townsend, E.C., Buker, S.M., Lancia, D.R., Caravella, J.A., Toms, A.V., Charlton, T.M., Lahdenranta, J., Wilker, E., Follows, B.C., Evans, N.J., Stead, L., Alli, C., Zarayskiy, V.V., Talbot, A.C., Buckmelter, A.J., Wang, M., McKinnon, C.L., Saab, F., McGouran, J.F., Century, H., Gersch, M., Pittman, M.S., Marshall, C.G., Raynham, T.M., Simcox, M., Stewart, L.M.D., McLoughlin, S.B., Escobedo, J.A., Bair, K.W., Dinsmore, C.J., Hammonds, T.R., Kim, S., Urbe, S., Clague, M.J., Kessler, B.M. & Komander, D. Molecular basis of USP7 inhibition by selective small-molecule inhibitors. *Nature* **550**, 481-486 (2017).
